# Automating microbial taxonomy workflows with PHANTASM: PHylogenomic ANalyses for the TAxonomy and Systematics of Microbes

**DOI:** 10.1101/2022.10.18.512716

**Authors:** Joseph S. Wirth, Eliot C. Bush

## Abstract

The nucleotide sequences of 16S ribosomal RNA (rRNA) genes have been used to inform the taxonomic placement of prokaryotes for several decades. Whole-genome approaches can better resolve evolutionary relationships of organisms, but these analyses often require computational proficiencies that are uncommon among microbiologists. PHANTASM is a new tool capable of automating these workflows. This tool was designed to work for a wide range of prokaryotes and is the first example of an automated reconciliation of NCBI’s Taxonomy database with that of the List of Prokaryotic names with Standing in Nomenclature (LPSN). In this study, we describe the workflow of PHANTASM and provide several examples of results generated by it. The source code is freely-available on GitHub. In order to facilitate the ease-of-access for researchers, PHANTASM is also available as a Docker image. While other tools exist to facilitate starting points for these analyses, PHANTASM provides users with a greater degree of control and produces outputs that can be used to make publication-quality figures.

## Introduction

### 16S ribosomal RNA genes in prokaryotic taxonomy

The nucleotide sequences of 16S ribosomal RNA (rRNA) genes have been used to inform the taxonomic placement of prokaryotes for several decades. Research has demonstrated that whole-genome approaches can better resolve evolutionary relationships of organisms, especially when the genetic distances are quite low (eg. strains of the same species) (1-5). However, the vast number of publicly available 16S rRNA gene sequences make this gene useful for obtaining a rough estimate of the phylogenetic position of a given taxon. Because of this, many studies rely on 16S rRNA gene sequences to determine which reference taxa to include when reconstructing the phylogeny of a group with the goal of justifying the taxonomic placement of an organism. Unfortunately, the unreliability of 16S rRNA as a phylogenetic marker can cause closely-related organisms to be omitted.

### Existing tools for conducting phylogenomic analyses for taxonomic purposes

Three existing tools that aim to assist researchers with the taxonomic placement of novel isolates are available: the Microbial Genome Atlas (MiGA) (6,7), the Type strain Genome Server (TYGS) (8), and the Genome Taxonomy Database Toolkit (GTDB-Tk) (9,10). MiGA (http://microbial-genomes.org/) identifies the closest relative of the query genome by leveraging the average amino acid identities (AAI) and average nucleotide identities (ANI) calculated for the query against a set of representative genomes. Once the closest-related representative genome is identified, it compares the query genome to the cluster of genomes which was represented by the best-matching representative genome (6,7). This method works well, however, it relies on the NCBI Taxonomy database (see below). Because of this discrepancy, MiGA may inadvertently omit genomes that should be included or include genomes that may not be useful.

The Type strain Genome Server (TYGS) is another tool developed with the purpose of facilitating taxonomic analyses based on whole-genome sequences and is owned by the DSMZ (8). This web-tool works by using 16S rRNA gene sequences as a proxy for taxonomic placement, and then performs phylogenomic analyses based on this initial placement. This workflow is attractive, but there are major drawbacks to TYGS revolving around the lack of free access. The code base is not open-source nor provided in a way that would allow offline use. This means that all genomes must be run through their online portal either directly or via a remote connection on the command line. Second, TYGS relies on a proprietary database of curated genome sequences, and this database cannot be examined in any meaningful way. Third, TYGS does not calculate common metrics such as average amino acid identity (AAI) and instead uses the DSMZ’s proprietary *in silico* approaches such as digital DNA-DNA hybridization (dDDH), genome-to-genome distance calculator (GGDC), and genome blast distance phylogeny (GBDP).

The Genome Taxonomy Database Toolkit, or GTDB-Tk, is another tool designed to help classify microbes using genomic data (9,10). This tool works by leveraging 120 bacterial or 122 archaeal marker genes. Using alignments of these genes, the input genome(s) is assigned to a domain and placed into a precalculated, domain-specific reference tree. GTDB-Tk then refines its classification using relative evolutionary divergence (RED) values (11) and ANI values to reference genomes. This method works well and its curated reference data helps promote accurate taxonomic assignments. However, the developers of GTDB-Tk v2 discourage people from using the trees produced by their workflow in publications and suggest inferring a tree using marker genes and methods suitable for the user’s data (personal communication).

GTDB-Tk v2 is designed to run on a server with multiple CPUs and ≥ 55 GB of RAM. Obtaining and maintaining such devices is not trivial as servers are expensive, need dedicated infrastructure for their upkeep and maintenance, and require expertise in administrating Linux systems. While GTDB-Tk can calculate ANI values for close relatives of the input genome(s), it does not offer a way of calculating AAI values. In addition, this program requires downloading more than 66 GB of reference data before the software can run. Taken together, this program is suited for identifying related genomes to serve as a starting point of phylogenomic analyses. Ultimately, GTDB-Tk has many merits but it was not designed for microbiologists with little to no computational experience.

These shortcomings, namely the potential for inaccurate taxonomic information used by MiGA, the non-open-source nature of TYGS, and the computational proficiencies required to use GTDB-Tk need to be addressed in any future tools developed for automating the taxonomic placement of prokaryotes.

### NCBI Taxonomy

NCBI Taxonomy is one of the best, free-to-use, publicly-available, taxonomic databases due to its sophisticated tools for automated queries and the fact that its entries are linked to NCBI’s other databases, such as the Assembly database, which houses thousands of high-quality, whole-genome assemblies. The taxonomy provided by NCBI is roughly accurate, but the NCBI Taxonomy database contains erroneous entries, especially when it comes to the accurate designation of type material (12-14). This is due, in part, to the fluid nature of taxonomy as a whole. While NCBI has steps in place to prevent the inaccurate classification of new additions to its Taxonomy database, it does not have automated ways for resolving conflicts that arose from the reclassification of existing species. In particular, NCBI lacks the ability to maintain accurate type material designations in its database, especially when dealing with changes to existing taxa such as one species being reclassified as two or more distinct species (14). These incongruencies of the NCBI Taxonomy database with the current state of the literature mean that NCBI Taxonomy is not an authoritative source of taxonomic information for *Archaea* and *Bacteria*.

### The List of Prokaryotic names with Standing in Nomenclature (LPSN) as an authoritative taxonomic database

Since its establishment in 1997 by Professor Jean Euzéby, the List of Prokaryotic names with Standing in Nomenclature (LPSN) has served the microbiological community by providing a centralized resource containing the most up-to-date taxonomic information for both *Archaea* and *Bacteria* (15-18). This website has been invaluable to microbial taxonomists and its content is accessed millions of times a year by researchers across the globe (17). Originally, this database was self-funded, but in recent years it has become part of the DSMZ (18). Importantly, this database is regularly updated and its taxonomic data is considered to be the most congruent with the official taxonomy recognized by the International Committee on Systematics of Prokaryotes. Since its acquisition by the DSMZ, a copyright agreement explicitly prohibits automated downloads of any data on its website without prior approval (https://lpsn.dsmz.de/text/copyright). Only a single text file that contains data for a subset of the validly published species and genera is provided for download (https://lpsn.dsmz.de/downloads), and there is no such resource for higher taxonomic ranks (ie. family, order, class, phylum, and domain). While the LPSN is in the process of developing an API, at the time of writing this tool is in its infancy as it takes hours to process large queries and does not provide access to any data that are not already available in the text file available for download.

### The importance of sequence selection for phylogenomic analyses

The conclusions of any phylogeny-based approach are dependent on the selection of reference sequences used for the analyses. Omitting close-relatives can lead to inaccurate classifications and the proposals of novel taxa that should instead be classified under previously described taxa. Because of this, it is important for phylogenomic analyses to make an effort to include a robust set of reference taxa in their phylogenetic reconstructions.

Wirth and Whitman (2018) resolved this issue by employing a novel technique to identify phylogenetic markers whose evolution was highly correlated with the inferred phylogeny of a group of *Rhodobacteraceae* (19). In that study, 16S rRNA gene sequences were used to initially determine which taxa to include in the analysis. However, this gene was poorly correlated with the phylogeny of this group. To address this problem, the researchers built a phylogenetic tree based on the core genes and then examined how well correlated each of the core genes were with the resulting phylogeny (19). This method identified an alternative phylogenetic marker, RpoC, as being highly correlated with the evolution of this group. Wirth and Whitman used RpoC sequence similarities to identify genomes that were closely-related to the taxa in their analysis but had not been detected by traditional 16S rRNA gene sequence similarities. This allowed for the inclusion of taxa that otherwise would have been omitted due to the poor resolution 16S rRNA gene sequences afforded in this group. Furthermore, this method prevented what would have been the erroneous proposals of several novel genera and instead facilitated the transfer of taxa to pre-existing genera (19). This methodology has proven useful for assessing the taxonomic positions of other taxonomic groups, such as the *Staphylococcaceae* (20) and the *Sphingomonadaceae* (21), and can be applied to a wide range of microbial taxa as it allows phylogenetic markers to be evaluated in a clade-specific manner.

In the present study, a new tool for automating this workflow is described: <u>PH</u>ylogenomic <u>AN</u>alyses for the <u>TA</u>xonomy and <u>S</u>ystematics of <u>M</u>icrobes (PHANTASM). This tool offers several improvements to the existing tools discussed above. It addresses the taxonomic inconsistencies found in NCBI Taxonomy by reconciling it with the taxonomy provided by the LPSN. It also applies the methodology of Wirth and Whitman (2018) to refine the genome selection via clade-specific phylogenetic markers (19). Furthermore, both the data used and code base are freely accessible (https://github.com/dr-joe-wirth/phantasm) which enables transparent analyses that are easily repeated and/or expanded upon. It does not require large downloads and can be run with the default settings on devices with as little as 8 GB of RAM. In order to facilitate ease-of-access for microbiologists, this software is also available as a Docker image on DockerHub (https://hub.docker.com/r/jwirth/phantasm). The Docker image contains all of the dependencies pre-installed, and needs only to be mounted as a container on a user’s machine. This means that Docker (https://www.docker.com/products/docker-desktop) is the only software that must be installed on a user’s machine.

## Materials and Methods

### Sequence selection for test cases

The whole genome sequences used to test PHANTASM were downloaded from NCBI (https://www.ncbi.nlm.nih.gov/) on February 15, 2022. A complete set of accession numbers and other important information can be found in **Table S1**.

### Obtaining taxonomic data from the LPSN

Data found on the List of Prokaryotic names with Standing in Nomenclature (LPSN) (https://lpsn.dsmz.de/) (18) website was critical for building an accurate taxonomy. Because DSMZ’s API was of limited usefulness (only provided access to a subset of the genera and species and was computationally inefficient on large queries), it was necessary to download the data as a csv file directly from the LPSN website (https://lpsn.dsmz.de/downloads).

The genus and species data were downloaded as a csv on January 5, 2023. Because the LPSN specifically prohibits automated downloads, the webpages for each parental genus, family, order, class, and phylum were manually visited between January 5, 2023 and January 8, 2023 and saved locally as html files. Custom python scripts were then used to scrape the relevant data from these html files in order to build the necessary data structures required for PHANTASM to function. **Table S2** provides a complete list of all the webpages accessed during this time. The generated text files can be found in the code repository on GitHub (https://github.com/dr-joe-wirth/phantasm) and within the Docker image hosted on DockerHub (https://hub.docker.com/r/jwirth/phantasm). In accordance with the LPSN copyright agreement (https://lpsn.dsmz.de/text/copyright), all entries in these text files contain the web addresses to the original LPSN webpages.

### Description of PHANTASM’s workflow

Because phylogenetic markers often vary in a clade-specific manner, it was important for PHANTASM to identify markers that could ensure a suitable set of genomes was selected for phylogenomic analyses. To this end, PHANTASM employs the following strategy (**Fig. 1**). First, the 16S rRNA gene sequences are extracted from the user’s input genome(s) and are used to query the NCBI targeted loci database (22) in order to identify the taxonomic orders with hits to the input’s 16S rRNA gene sequences. Alternatively, the user may provide a list of NCBI Taxonomy IDs indicating which families, genera, and/or species the input genome(s) are most closely related to. This can be particularly useful for genome assemblies whose 16S rRNA gene sequences were not recruited and annotated. Next, the taxonomic data for these orders are downloaded from NCBI Taxonomy and stored in a custom data structure. This taxonomic structure is then reconciled with that of the List of Prokaryotic names with Standing in Nomenclature (LPSN) in an effort to minimize the use of inaccurate and/or redundant names (16-18). Once reconciled, the records are linked to NCBI Assembly which contains the relevant data for publicly-available whole-genome sequences.

**Figure 1.**
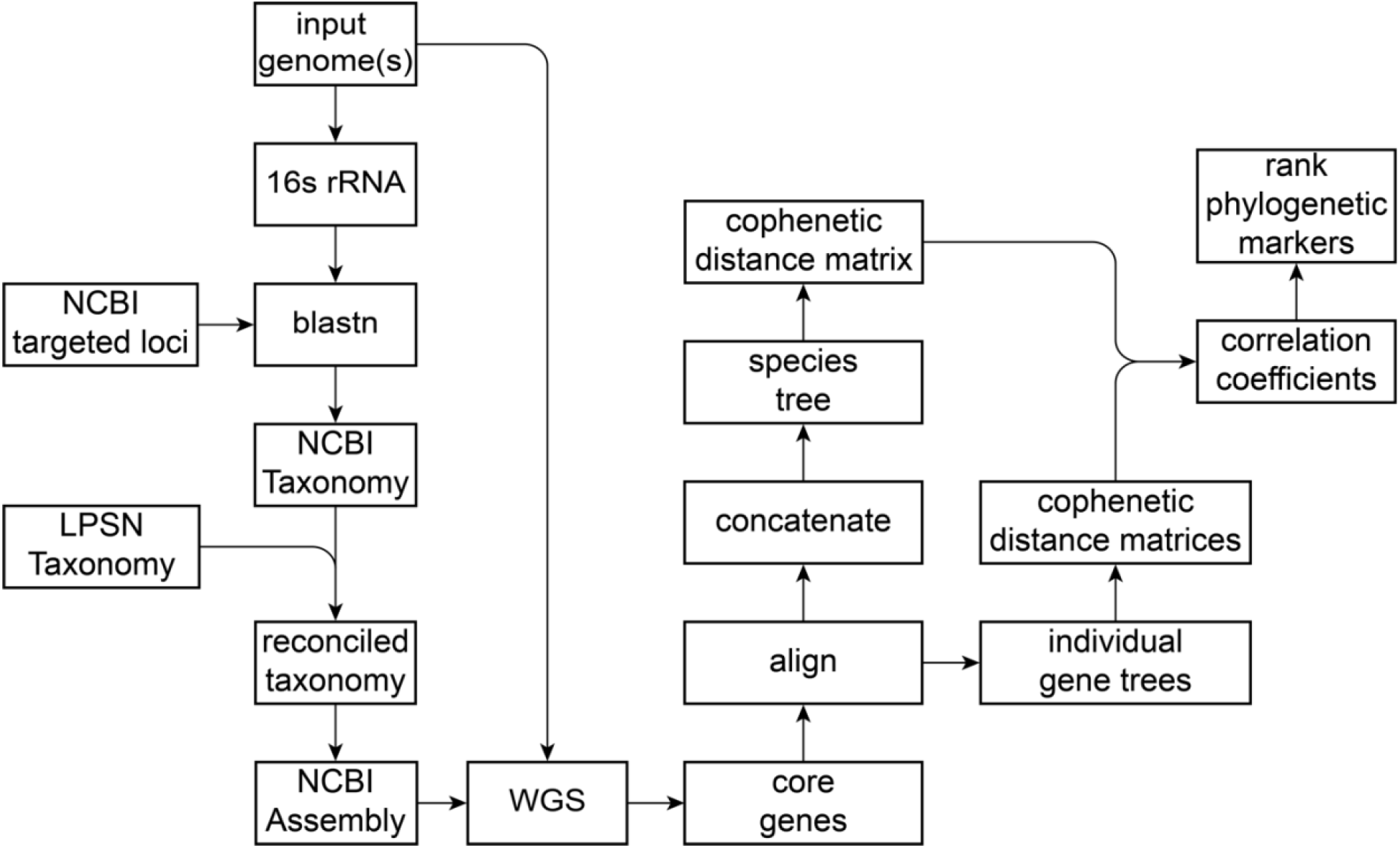
Workflow to identify a suitable phylogenomic marker for the input genome. One or more annotated genomes in GenBank format are provided to PHANTASM as input. The 16S rRNA gene sequences from these genomes are used to identify a taxonomic space (eg. a taxonomic order). If 16S rRNA gene sequences are unavailable, the user may specify a list of NCBI Taxonomy IDs of related taxa instead. The taxonomic data is initially built from the NCBI Taxonomy database and is reconciled with the taxonomic structure of the List of Prokaryotic names with Standing in Nomenclature (LPSN). PHANTASM downloads whole genome sequences (WGS) to maximize the taxonomic breadth represented. It uses these genomes to calculate the core genes shared between all genomes. An individual gene tree is constructed for each core gene, the alignments are concatenated, and the concatenated alignment is used to reconstruct a phylogenomic tree. Finally, a correlation coefficient of the cophenetic distances is calculated for each core gene tree as it compares to the species tree. These cophenetic correlation coefficients are then used to rank each of the core genes for their suitability as a phylogenetic marker. This step can be skipped if the user has already identified a phylogenetic marker(s) and/or a set of reference genomes.

Next, an ingroup and outgroup are identified based on both taxonomic data and sequence homology. This strategy allows PHANTASM to maximize the breadth of taxonomic space represented which helps to counteract any biases present in the 16S rRNA gene sequences. Once a set of suitable genomes is identified, PHANTASM automatically downloads them from NCBI’s Assembly database.

Using these genomes, a core genome is calculated (see below). These core genes are individually aligned using MUSCLE v5.1 (23), the alignments are concatenated, and the concatenated alignment is used to construct a species tree using FastTree v2.1.11 (24,25). Each core gene alignment is also used to create individual gene trees using FastTree v2.1.11. A Pearson correlation coefficient for each core gene is calculated by comparing the cophenetic distances of its gene tree to the cophenetic distances of the species tree (19,26). These cophenetic correlation coefficients are then used to rank the core genes for their suitability as a phylogenetic marker for the genomes of interest, and the results are saved to a text file for the user to examine. This step can be skipped if the user has already identified one or more suitable phylogenetic markers for their input genome(s). Additionally, users can choose to exclude specific taxa from their analysis.

After ranking the core genes, a user must select one or more core genes to be used as phylogenetic markers. Using this gene, PHANTASM refines the taxa included in its phylogeny (**Fig. 2**). The protein sequence(s) for the selected gene(s) in the input genome(s) is used to query the non-redundant protein database on NCBI. If this query is too computationally expensive, then it will instead query the RefSeq Protein database on NCBI. Based on these results, PHANTASM automatically selects genomes in order to maximize the relevant taxonomic space. To do this, the following criteria are used: one genome (preferably the type strain) for each species from the most closely-related genus is added to the set of genomes, and the minimum bitscore of this genus is saved. Any genus containing one or more species whose bitscores are higher than the aforementioned minimum bitscore also have a single genome for each of their children species added to the set of genomes. For all other genera identified in the blast results, only a single species is selected with a preference for the type strain of the type species where possible. Finally, an outgroup is selected from outside the targeted taxonomic space. This step can be skipped if the user has already identified a suitable set of reference genomes.

**Figure 2.**
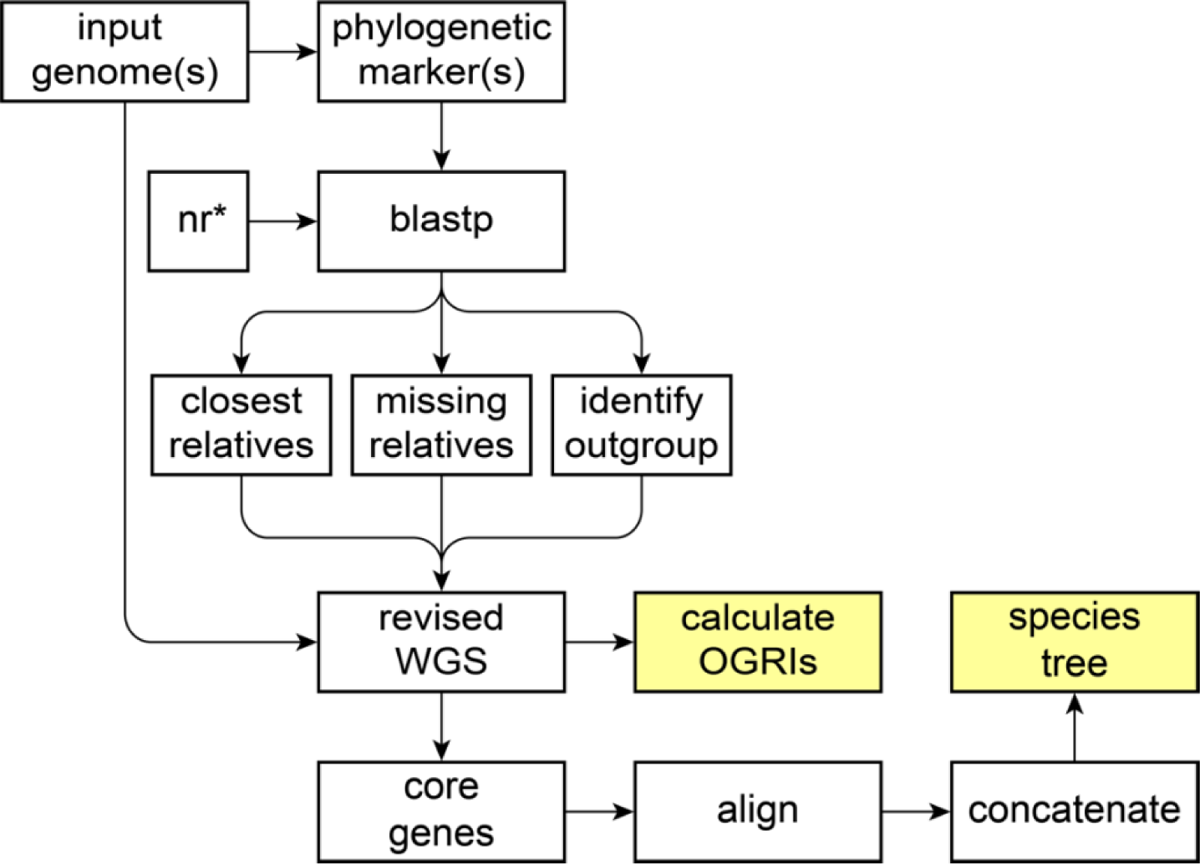
Workflow to refine the phylogeny and perform phylogenomic analyses. Using one or more phylogenetic markers identified by the user, PHANTASM performs a BLASTp against the non-redundant protein database (nr) and uses the results to identify a suitable set of genomes in an effort maximize the relevant taxonomic space. These genomes are downloaded, and the core genes are calculated based on BLASTp scores. The core genes are aligned and concatenated, and this concatenated alignment is used to construct a species tree. The overall genome related indices (OGRIs), specifically average amino acid identity (AAI) and average nucleotide identity (ANI) matrices, are calculated and used to build heatmaps of the AAI and ANI values. If a suitable set of reference genomes are already known, users can skip reference genome selection and instead run only the analyses starting at the “revised WGS” step. *: If a BLASTp against the non-redundant protein database (nr) is too computationally expensive, then PHANTASM will query the RefSeq Protein database instead.

After identifying a suitable set of reference genomes informed by sequence homology of a given phylogenetic marker(s), PHANTASM downloads the selected genomes from NCBI’s Assembly database and constructs a species tree from a concatenated alignment of the core genes as described above. Users are given the option of generating bootstrapped trees. This setting is off by default due to the significant increase in run times. If users elect to produce bootstrapped trees, then PHANTASM employs IQTree with automatic model selection (27,28). In addition, PHANTASM calculates the average amino acid identity (AAI) and the average nucleotide identity (ANI) (see below) and generates heatmaps based on the values. Together, this provides the user with much of the information necessary to taxonomically assign their microbe(s) of interest. Because all of the resulting data are freely available, the user can choose to perform additional analyses using the genomes identified by PHANTASM.

### Calculation of core genes

Core genes are calculated using xenoGI as previously described (29). Briefly, an all-vs-all BLASTp is performed for all pairwise combinations of genomes. The resulting BLAST tables are then used to identify orthologs with exactly one copy in each genome using the all-around best reciprocal hit method. Core genes are also required to have an evalue of at least 10^−8^, a percent identity of at least 35 %, and a query alignment coverage of at least 65 %. Users can opt to reduce the number of core genes used to construct the “species tree” by excluding proteins whose alignments contain more than 5% gaps for one or more taxa (19-21,30).

### Phylogenetic and phylogenomic tree construction

A multiple sequence alignment is made for each core gene using MUSCLE v5.1 (23) with the default settings, and these alignments are used as input for FastTree v2.1.11 (24,25) with the default settings to construct individual “gene trees”. The alignments are also concatenated, and this concatenated alignment is used as input for FastTree v2.1.11 with the default settings to construct a “species tree”. Alternatively, IQTree v1.6.12 can be used to construct the “species tree” if bootstrap supports are required.

### Ranking phylogenetic markers

Phylogenetic markers are ranked as previously described with some modifications (19,20,26). Briefly, cophenetic distance matrices are obtained from each of the gene trees as well as from the species tree. The cophenetic distance matrix of each gene tree is compared to that of the species tree and a Pearson correlation coefficient is calculated. These coefficients range from 0 to 1 and are used to rank genes as putative phylogenetic markers. Higher correlation coefficients are indicative of genes whose phylogenetic reconstructions are better correlated with the inferred phylogeny based on the concatenated alignment of all core genes. Such genes may constitute good phylogenetic markers, but ultimately the user must decide which gene(s) should be employed as a phylogenetic marker(s).

### Calculation of average amino acid identity

For each pairwise comparison of genomes, exactly two BLASTp tables are generated. From these two tables, the best reciprocal hits for the respective pair of genomes are identified, and the percent identities of these proteins as calculated by BLASTp are obtained. The average amino acid identity (AAI) is calculated using xenoGI (29) for each pairwise combination of genomes by determining the mean percent identity weighted by the length of the alignment using the following equation:

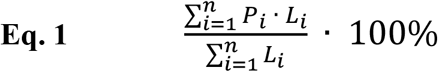

where *n* is the number of shared proteins as determined by BLASTp, *P* is the percent identity of a protein shared between two genomes (ie. the percent identity in a BLASTp alignment), and *L* is the alignment length of the shared protein.

### Calculation of average nucleotide identity

Average nucleotide identity (ANI) is calculated using pyani v.0.2.11 (31) with the mummer method (ANIm). Null values produced when pyani fails for a given pairwise combination are omitted from the resulting plots.

## Results and Discussion

### Using Ruegeria pomeroyi as a test case

Because the *Alphaproteobacteria* have undergone several major taxonomic revisions in recent years (19,32-34), a genome from this group was a good example for demonstrating PHANTASM’s functionality. To this end, the whole genome sequence for *Ruegeria pomeroyi* DSS-3^T^ (GCF_000011965.2) was used as input to PHANTASM. The initial phylogeny that was reconstructed by PHANTASM is shown in **Figure 3**. As can be seen, PHANTASM accurately identified the input genome as a member of the order *Rhodobacterales*, and attempted to maximize the taxonomic breadth of the taxonomic space by including a diverse set of genomes within the *Alphaproteobacteria*. PHANTASM identified several suitable phylogenetic markers in these organisms (**Table S3**), including *rpoC* and *gyrB*, both of which had cophenetic correlation coefficients of 0.98. Using *rpoC* (SPO_RS17765) and *gyrB* (SPO_RS00795), phylogenetic refinement was performed followed by phylogenomic analyses (**Fig. 4, Fig. 5**). PHANTASM accurately identified *Ruegeria, Phaeobacter, Leisingera*, and *Tritonibacter* as the most closely-related genera to the input, and fully populated the species for these genera. In addition, PHANTASM included a single genome for several other closely-related genera. The AAI (**Fig. 4**) and ANI (**Fig. 5**) heatmaps also show that the input genome is identical to that of *Ruegeria pomeroyi* DSS-3^T^, which was expected. **Figures 4** and **5** are not the raw outputs from PHANTASM, but were generated by combining the species tree in Newick format with a pdf of the AAI and ANI heatmaps, respectively. Examples of raw outputs can be found in the supplementary material (**Fig. S1-S24**).

**Figure 3.**
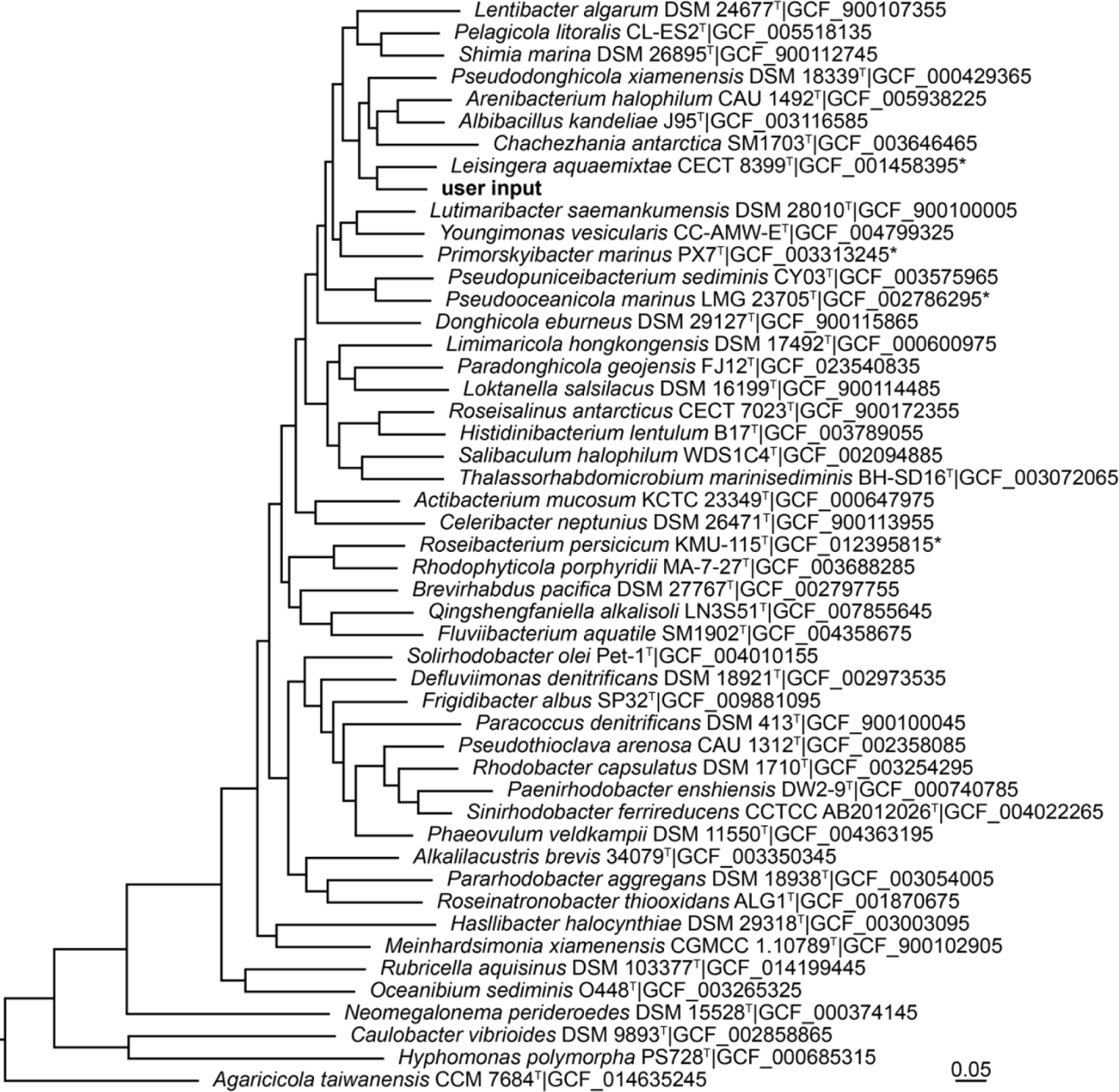
Initial phylogeny constructed by PHANTASM when using the whole genome sequence of *Ruegeria pomeroyi* DSS-3 as input. The tree was constructed from a concatenated alignment of the core genes as described in Materials and Methods. The input genome (GCF_000011965.2) is bolded. All other genomes were automatically identified by PHANTASM. The tree was rooted on *Acidithiobacillus thiooxidans* ATCC 19377^T^ (GCF_009662475.1), which has been omitted to allow better resolution of the relevant phylogenomic relationships. The scale bar indicates the number of substitutions per site. All genomes, including the outgroup, represent the type strain of their respective species. Unless indicated with an asterisk, all genomes represent the type species of their respective genus.

**Figure 4.**
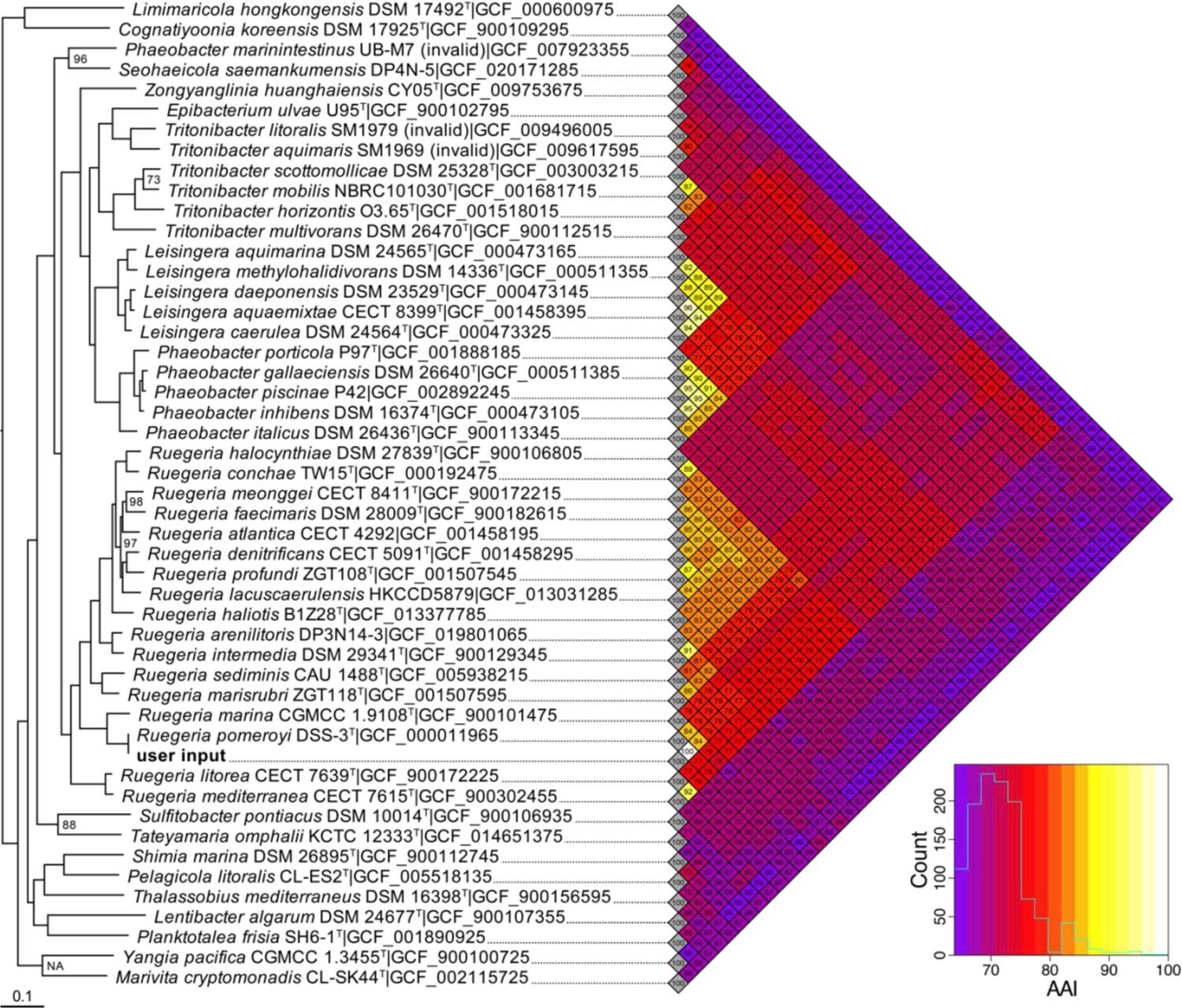
Phylogenomic tree and heatmap of average amino acid identities produced by PHANTASM when using the whole genome sequence of *Ruegeria pomeroyi* DSS-3 as input. The tree was constructed from a concatenated alignment of the core genes using IQTree with automatic model selection as described in Materials and Methods. The input genome (GCF_000011965.2) is bolded. All other genomes were automatically identified by PHANTASM. The tree was rooted on *Acidithiobacillus thiooxidans* ATCC 19377^T^ (GCF_009662475.1), which has been omitted to allow better resolution of the relevant phylogenomic relationships. The scale bar indicates the number of substitutions per site. Invalidly published species are indicated. Genomes representing the type strain of their respective species are indicated with a superscript T. Unless specified, all nodes had 100% bootstrap support after 100 parametric bootstrap replicates. The distribution of AAI values is shown in the bottom right corner. The heatmap (pdf) and species tree (newick) were combined to produce this figure. Examples of raw output can be found in the supplementary material.

**Figure 5.**
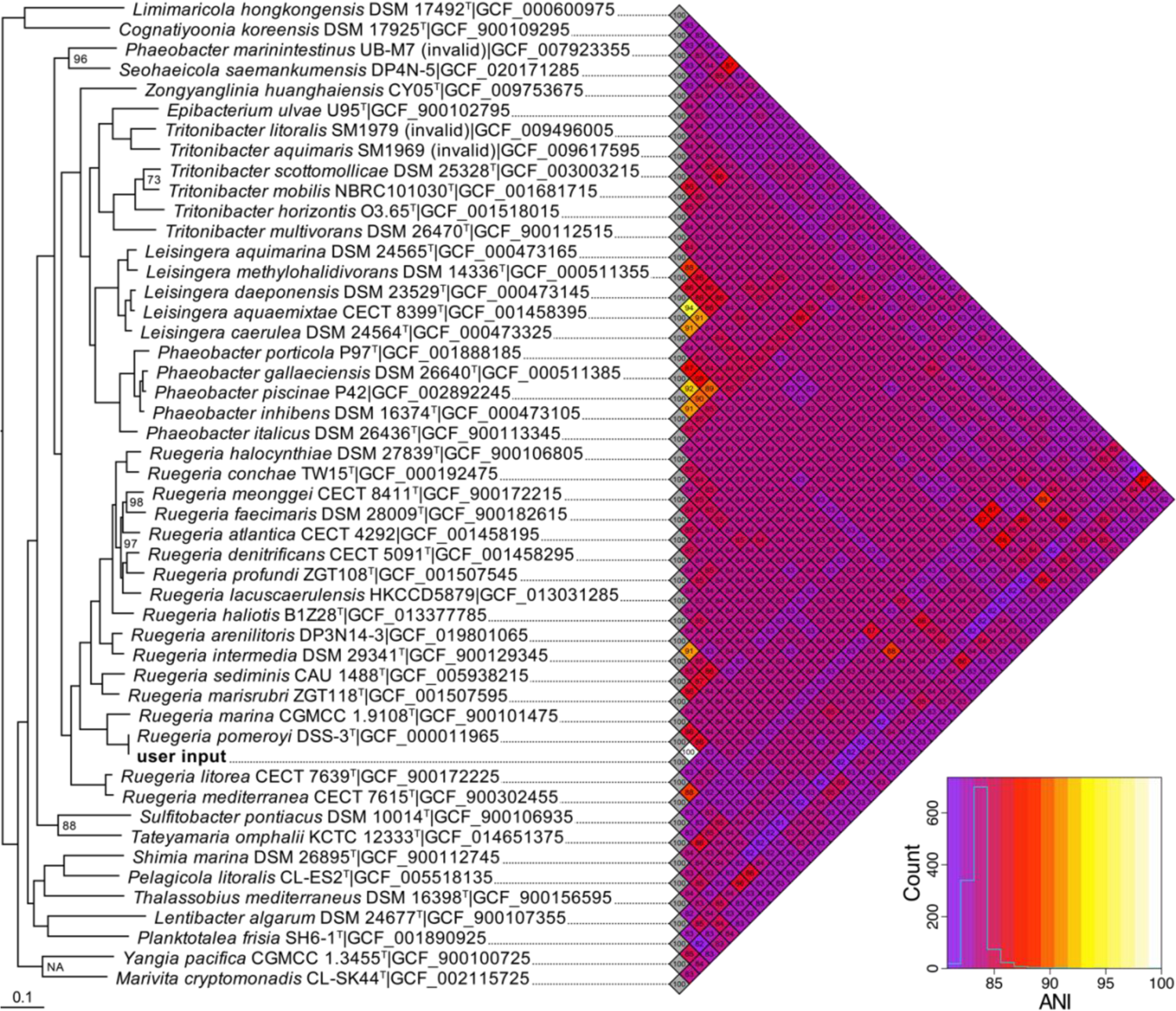
Phylogenomic tree and heatmap of average nucleotide identities produced by PHANTASM when using the whole genome sequence of *Ruegeria pomeroyi* DSS-3 as input. The tree was constructed from a concatenated alignment of the core genes as described in Materials and Methods. The input genome (GCF_000011965.2) is bolded. All other genomes were automatically identified by PHANTASM. The tree was rooted on *Acidithiobacillus thiooxidans* ATCC 19377^T^ (GCF_009662475.1), which has been omitted to allow better resolution of the relevant phylogenomic relationships. The scale bar indicates the number of substitutions per site. Invalidly published species are indicated. Genomes representing the type strain of their respective species are indicated with a superscript T. Unless specified, all nodes had 100% bootstrap support after 100 parametric bootstrap replicates. The distribution of ANI values is shown in the bottom right corner. The heatmap (pdf) and species tree (newick) were combined to produce this figure. Examples of raw output can be found in the supplementary material.

### Comparing phylogenetic marker calling by PHANTASM and the GET_HOMOLOGUES + GET_PHYLOMARKERS pipeline

The phylogenetic markers automatically identified for the *Ruegeria pomeroyi* test case were in agreement with the GET_HOMOLOGUES v 16092021 and GET_PHYLOMARKERS v 2.4.6_20nov2022 pipeline (35,36). These tools can be used to identify a core genome and suitable phylogenetic markers, respectively. For this test case, GET_HOMOLOGUES found 121 of the 125 core genes identified by PHANTASM, and PHANTASM found 121 of the 182 core genes identified by GET_HOMOLOGUES. By default, GET_PHYLOMARKERS only identifies the top 60 phylogenetic markers. 47 of these 60 markers were also identified as core genes by PHANTASM. However, the performance of GET_PHYLOMARKERS was not as robust as PHANTASM. The GET_HOMOLOGUES + GET_PHYLOMARKERS pipeline took over 9 hours to run on a Linux machine with 50 CPUs while PHANTASM took less than half the time to accomplish this task on the same machine.

Unlike PHANTASM, GET_PHYLOMARKERS does not numerically score individual genes for their suitability as phylogenetic markers, so it was not possible to know the ranking of the 60 phylogenetic markers identified with this software package. However, these genes were manually scored using the same method employed by PHANTASM. PHANTASM identified 40 of the 43 core genes with cophenetic correlation coefficients greater than or equal to 0.95, while GET_PHYLOMARKERS only found 21 of these genes (**Table S3**). Additionally, PHANTASM identified 89 of the 95 core genes with cophenetic correlation coefficients greater than or equal to 0.90 (**Table S3**). Of the six genes that were missed by PHANTASM, only *dnaJ* has been previously published as a suitable phylogenetic marker for the *Alphaproteobacteria* (37). GET_PHYLOMARKERS also identified 18 genes whose cophenetic correlation coefficients were below 0.90, including *fumC*, a gene with a very low cophenetic correlation coefficient (0.42). While PHANTASM calculated the cophenetic correlation coefficient for all the core genes it identified, the documentation specifically discourages the use of genes whose values are below 0.90. Taken together, these results suggest that automated phylogenetic marker calling in PHANTASM is functioning properly.

### PHANTASM produces results that are congruent with existing tools

The results generated by PHANTASM for *Ruegeria pomeroyi* are congruent with those obtained from existing software. The phylogeny generated by TYGS (**Fig. 6**) is in agreement with that generated by PHANTASM. However, the results generated by PHANTASM have several advantages: 1) the tree has more phylogenomic context by including a wider distribution of genera, 2) the tree does not have redundant strains, 3) the number of tips in the tree can be adjusted by the user, 4) all data are freely available including the accession numbers for the reference genomes, and 5) the AAI and ANI matrices are automatically calculated.

**Figure 6.**
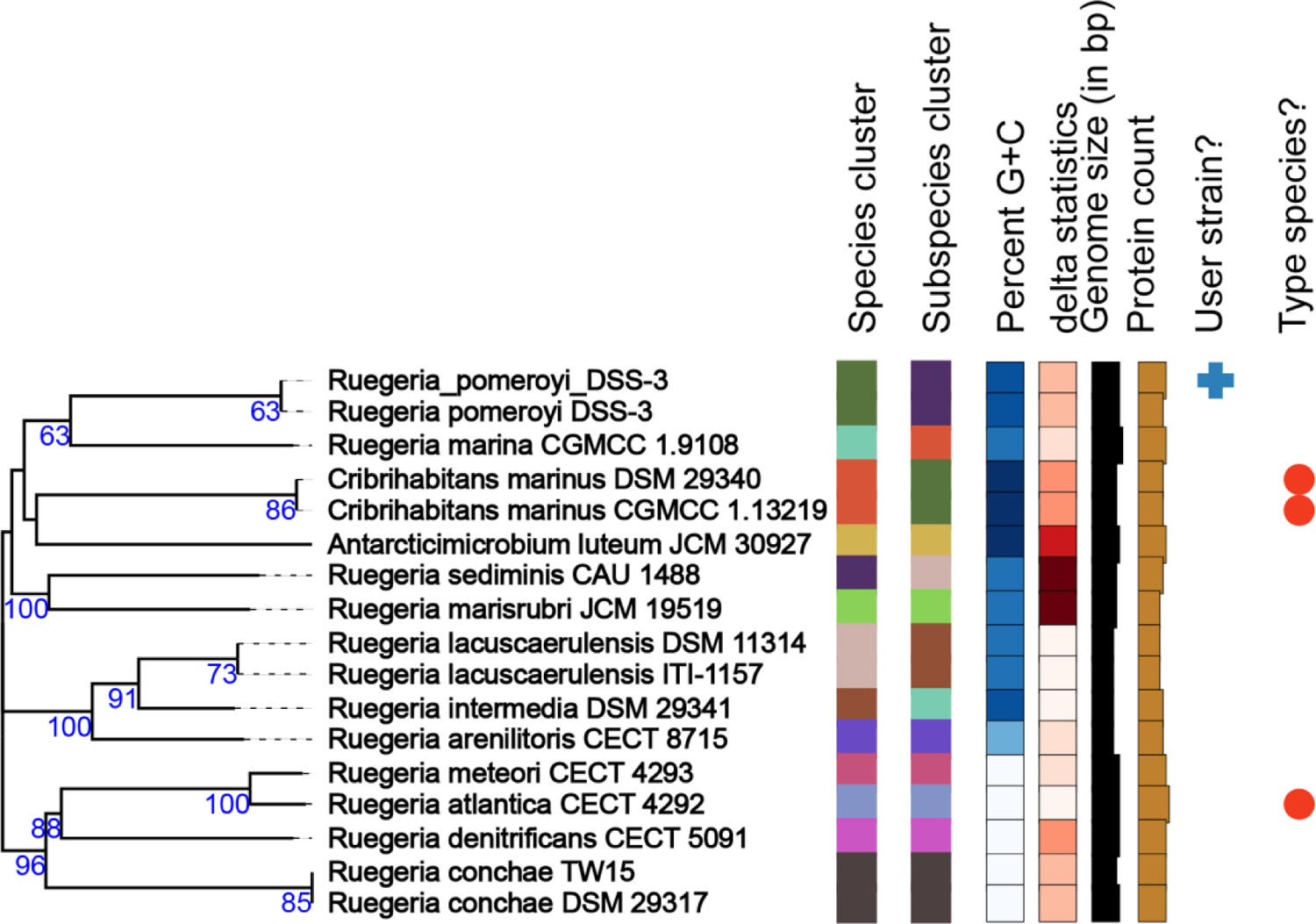
Phylogenomic analysis by TYGS when using the whole genome sequence of *Ruegeria pomeroyi* DSS-3 as input. Figure was generated by the Type Strain Genome Server (TYGS) (https://tygs.dsmz.de/) and no modifications have been made. The input genome (GCF_000011965.2) is indicated with a blue cross.

Similarly, the results generated by PHANTASM agree with those generated by MiGA, although some values differ due to the estimated AAI method employed by MiGA (data available online; see below). However, both PHANTASM and MiGA accurately identified the input genome as an existing strain of *Ruegeria pomeroyi* and MiGA indicated that the input had 100% average nucleotide identity (ANI) with *Ruegeria pomeroyi* DSS-3^T^ as was expected. MiGA did not offer a way to export figures, but the data can be accessed directly (http://microbial-genomes.org/query_datasets/M:SZJ_XNS) by logging in with the username and using the password “password”. MiGA did not produce a phylogenomic tree, and the AAI values were less accurate than those produced by PHANTASM due to the utilization of the “heuristic AAI” method used by MiGA (6).

PHANTASM also generated results that were congruent with GTDB-Tk v2 (9,10). The summary file output by this program accurately assigned the genome to *Ruegeria pomeroyi* and provided the accession number for the reference sequence of the type strain’s whole genome. However, interpreting the phylogenetic tree produced by GTDB-Tk v2 was quite difficult as this tree had 10,527 tips. In addition, the tree produced by GTDB-Tk v2 had no taxonomic names and instead listed only the accession numbers for the RefSeq (RS) and GenBank (GB) whole genome sequences (**Fig. 7**). GTDB-Tk v2 indicated that the input genome shared 100 % ANI with *Ruegeria pomeroyi* DSS-3 but did not output any additional ANI values. In contrast, PHANTASM produces trees with taxonomic names as well as the accession numbers of the genomes. PHANTASM also calculates AAI and ANI for each pairwise combination of genomes.

**Figure 7.**
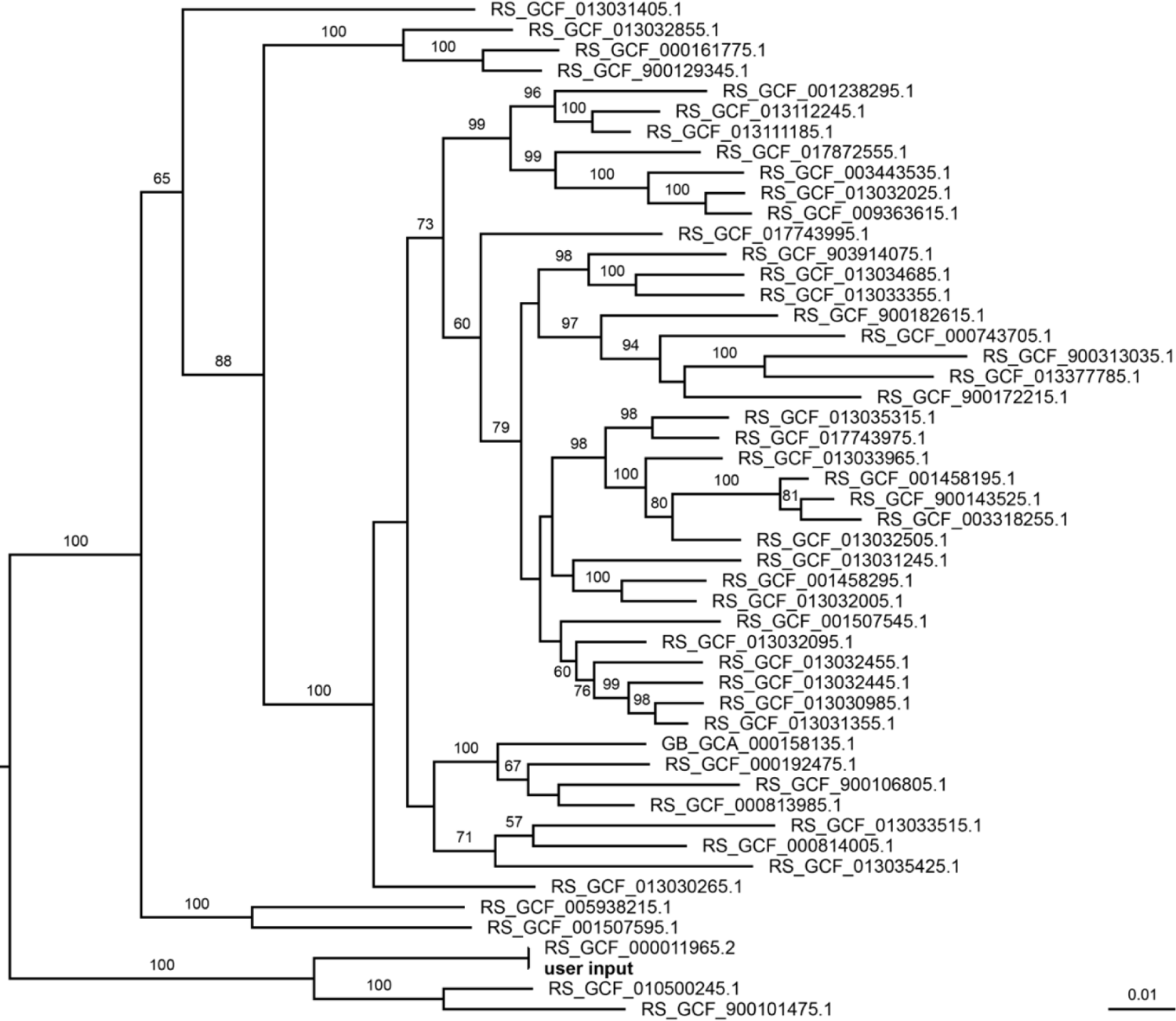
Maximum likelihood phylogenetic tree produced by GTDB-Tk v2. The input genome is bolded. The tree represents a subtree pruned from a tree containing 10,527 tips. Branch labels indicate statistical support after 100 non-parametric bootstrap replicates; values less than 50 % have been omitted. The scale bar indicates substitutions per site.

### PHANTASM works with a wide range of Archaeal and Bacterial taxa

PHANTASM has been tested on members of several other taxonomic groups outside of the *Alphaproteobacteria* including *Firmicutes* (**Fig. S1-S4**), *Gammaproteobacteria* (**Fig. S5-S8**), *Bacteroidetes* (**Fig. S9-S12**), *Euryarchaeota* (**Fig. S13-S20**), and *Crenarchaeota* (**Fig. S21-S24**). PHANTASM also works with *Cyanobacteria* (data not shown). However, reliance on the text files available for download at the LPSN has significantly reduced the ability of PHANTASM to accurately work with cyanobacterial genomes as the cyanobacterial nomenclature is largely omitted from the file. While most of the genomes used to test PHANTASM are complete (**Table S1**), this is not a requirement for PHANTASM to work. In fact, one genome tested had 196 contigs. The only requirement for PHANTASM to run successfully is that the coding sequences are annotated. In fact, PHANTASM has been shown to work with genomic data obtained via single-cell sequencing, and these sequences contained between 64 and 194 contigs (data not shown).

### PHANTASM provides the first description of an open-source and automated way to reconcile NCBI’s Taxonomy with that of the LPSN

While the List of Prokaryotic names with Standing in Nomenclature (LPSN) has existed for 25 years, PHANTASM provides the first example of automatically reconciling the data found in NCBI’s Taxonomy database with that of the LPSN. We intend to update the text files containing the data from the LPSN annually and hope that the LPSN adds to their suite of tools for automatically accessing the data on their website so that this reconciliation can be fully automated in the future.

## Data Availability

All genomes presented in this study are publicly available via NCBI, and the relevant data for these genomes have been provided in **Table S1**. The source code for PHANTASM is freely available on GitHub (https://github.com/dr-joe-wirth/phantasm). PHANTASM is also available as a Docker image on DockerHub (https://hub.docker.com/r/jwirth/phantasm).

## Acknowledgements

We would like to thank Harvey Mudd College undergraduates Michelle L. Johnson and Tonatiuh Gonzalez for their helpful discussions and technical advice. We would also like to thank Helen Dukes at the University of Georgia for her help with testing PHANTASM on single-cell genome sequences.

