## Supplemental Material for "Automating microbial taxonomy workflows with PHANTASM: PHylogenomic ANalyses for the TAxonomy and Systematics of Microbes"

### Supplementary Material

Joseph S. Wirth, Eliot C. Bush

**Table S1. Genomes used as inputs to PHANTASM.** All genomes were downloaded on February 15, 2022. The phylogenetic marker refers to the gene used to refine the phylogeny using PHANTASM. "*Sulfolobus islandicus*" is not validly published.

| Organism | Strain | RefSeq Accession | Assembly level | Contigs | Phylogenetic marker(s) | |
|---|---|---|---|---|---|---|
| *Ruegeria pomeroyi* | DSS-3$^{T}$ | GCF_000011965.2 | complete | 2 | *rpoC*, *gyrB* | SPO_RS17765, SPO_RS00795 |
| *Staphylococcus aureus* | DSM 20231$^{T}$ | GCF_001027105.1 | complete | 2 | *gyrA* | AA076_RS00030 |
| *Salmonella enterica* | LT2$^{T}$ | GCF_000006945.2 | complete | 2 | *dnaE*, *gyrB* | STM0231, STM3835 |
| *Williamwhitmania taraxaci* | A7P-90m$^{T}$ | GCF_900096565.1 | scaffold | 196 | *rpsC* | BLS65_RS03755 |
| *Methanosarcina mazei* | S-6$^{T}$ | GCF_000970205.1 | complete | 1 | DNA topoisomerase I | MSMAS_RS08570 |
| *Pyrococcus furiosus* | DSM 3638$^{T}$ | GCF_008245085.1 | complete | 1 | lysine-tRNA ligase | PFDSM3638_RS00615 |
| "*Sulfolobus islandicus*" | Y.N.15.51 | GCF_000022485.1 | complete | 2 | DNA polymerase I | YN1551_RS06105 |

**Table S3. Comparison of the phylogenetic marker identification capability of PHANTASM and the GET_HOMOLOGUES + GET_PHYLOMARKERS pipeline (35,36).** PHANTASM and GET_HOMOLOGUES version 16092021 (GET_HOMS) were used to determine the core genes. PHANTASM and GET_PHYLOMARKERS v2.4.6_20nov2022 (GET_PHYLO) were used to identify phylogenetic markers. PHANTASM documentation discourages the used of genes with cophenetic correlation coefficients ($r_{CCC}$) below 0.90 as phylogenetic markers. All cophenetic correlation coefficients were calculated using the PHANTASM software package. A plus sign (+) indicates that the software found a gene and a minus sign (–) indicates that the software did not find a gene. Minus signs in the gene column indicates CDS without annotated gene names.

| | | | **core gene?** | | **phylogenetic marker?** | |
|---|---|---|---|---|---|---|
| **locus tag** | **gene** | ***rCCC*** | **PHANTASM** | **GET_HOMS** | **PHANTASM** | **GET_PHYLO** |
| SPO_RS17765 | *rpoC* | 0.98 | + | + | + | + |
| SPO_RS08905 | *rpoD* | 0.98 | – | + | – | + |
| SPO_RS11250 | *uvrA* | 0.98 | + | + | + | + |
| SPO_RS16870 | *rnr* | 0.97 | + | + | + | – |
| SPO_RS01455 | *pyrB* | 0.97 | + | + | + | – |
| SPO_RS05055 | *clpX* | 0.97 | + | + | + | – |
| SPO_RS16950 | *rlmN* | 0.97 | + | + | + | – |
| SPO_RS04045 | *rpsA* | 0.97 | + | + | + | – |
| SPO_RS18205 | *pheS* | 0.97 | + | + | + | – |
| SPO_RS01630 | *ispB* | 0.96 | + | + | + | + |
| SPO_RS18425 | *uvrC* | 0.96 | + | + | + | – |
| SPO_RS00045 | *hrcA* | 0.96 | + | + | + | + |
| SPO_RS15980 | *ribF* | 0.96 | + | + | + | + |
| SPO_RS16030 | *atpA* | 0.96 | + | + | + | + |
| SPO_RS15385 | *ask* | 0.96 | + | + | + | – |
| SPO_RS15235 | *msbA* | 0.96 | + | + | + | + |
| SPO_RS10280 | – | 0.96 | + | + | + | – |
| SPO_RS18490 | *hemE* | 0.96 | + | + | + | – |
| SPO_RS15890 | *ileS* | 0.96 | + | + | + | + |
| SPO_RS02750 | *uvrB* | 0.96 | + | + | + | + |
| SPO_RS02455 | *rpsC* | 0.96 | + | + | + | – |
| SPO_RS08455 | *frr* | 0.95 | + | + | + | – |
| SPO_RS10305 | *typA* | 0.95 | + | + | + | – |
| SPO_RS17165 | *pnp* | 0.95 | + | + | + | – |
| SPO_RS00220 | *dnaJ* | 0.95 | – | + | – | + |
| SPO_RS02450 | *rplV* | 0.95 | + | + | + | + |
| SPO_RS10320 | *recA* | 0.95 | + | + | + | – |
| SPO_RS13115 | *prfB* | 0.95 | + | + | + | – |
| SPO_RS19705 | *hslU* | 0.95 | + | + | + | + |
| SPO_RS19470 | *truB* | 0.95 | + | + | + | + |
| SPO_RS08920 | *nrdR* | 0.95 | + | + | + | – |
| SPO_RS00300 | *argJ* | 0.95 | + | + | + | – |
| SPO_RS10930 | *gltX* | 0.95 | + | + | + | + |
| SPO_RS02125 | *mreB* | 0.95 | + | + | + | + |
| SPO_RS19145 | – | 0.95 | + | + | + | + |

| | | | | | | |
|---|---|---|---|---|---|---|
| SPO_RS06720 | *purA* | 0.95 | + | + | + | + |
| SPO_RS13670 | *prfC* | 0.95 | + | + | + | – |
| SPO_RS01700 | *lysA* | 0.95 | + | + | + | + |
| SPO_RS02675 | *yihA* | 0.95 | – | + | – | + |
| SPO_RS11155 | *coaD* | 0.95 | + | + | + | – |
| SPO_RS02580 | *rpoA* | 0.95 | + | + | + | + |
| SPO_RS04075 | *trpB* | 0.95 | + | + | + | – |
| SPO_RS14455 | *gatB* | 0.95 | + | + | + | – |
| SPO_RS19375 | – | 0.94 | – | + | – | + |
| SPO_RS08580 | *rplM* | 0.94 | + | + | + | – |
| SPO_RS13270 | *tgt* | 0.94 | + | + | + | – |
| SPO_RS19090 | *map* | 0.94 | + | + | + | – |
| SPO_RS14080 | *nuoH* | 0.94 | + | + | + | – |
| SPO_RS06770 | *gor* | 0.93 | + | + | + | + |
| SPO_RS06955 | *glmM* | 0.93 | + | + | + | + |
| SPO_RS13935 | – | 0.93 | + | + | + | – |
| SPO_RS06905 | *glyQ* | 0.93 | + | + | + | + |
| SPO_RS02585 | *rplQ* | 0.93 | + | + | + | – |
| SPO_RS17795 | *rplA* | 0.93 | + | + | + | – |
| SPO_RS04575 | *lgt* | 0.93 | – | + | – | + |
| SPO_RS19400 | *rplW* | 0.93 | + | + | + | – |
| SPO_RS18140 | *nth* | 0.93 | + | + | + | – |
| SPO_RS06095 | *ddlB* | 0.93 | + | + | + | – |
| SPO_RS13930 | *ycfH* | 0.93 | + | + | + | + |
| SPO_RS09495 | *grxD* | 0.92 | + | + | + | – |
| SPO_RS00745 | *ubiE* | 0.92 | + | + | + | – |
| SPO_RS02535 | *rpsE* | 0.92 | + | + | + | + |
| SPO_RS15770 | *tolQ* | 0.92 | + | – | + | – |
| SPO_RS17815 | *nusG* | 0.92 | + | + | + | + |
| SPO_RS13085 | *ilvN* | 0.92 | + | + | + | + |
| SPO_RS17375 | *leuS* | 0.92 | + | + | + | – |
| SPO_RS14050 | *nuoK* | 0.92 | + | + | + | – |
| SPO_RS02445 | *rpsS* | 0.92 | + | + | + | – |
| SPO_RS02440 | *rplB* | 0.92 | + | + | + | + |
| SPO_RS02525 | *rplF* | 0.92 | + | + | + | + |
| SPO_RS19310 | *smpB* | 0.92 | + | + | + | + |
| SPO_RS17455 | *mutY* | 0.92 | – | + | – | + |
| SPO_RS19740 | *dnaQ* | 0.92 | + | + | + | – |
| SPO_RS14130 | *nuoD* | 0.92 | + | + | + | + |
| SPO_RS07190 | *rpoH* | 0.91 | + | + | + | + |
| SPO_RS06530 | *thrS* | 0.91 | + | – | + | – |
| SPO_RS20525 | *murA* | 0.91 | + | + | + | – |
| SPO_RS16020 | *atpD* | 0.91 | + | + | + | + |
| SPO_RS02500 | *rplN* | 0.91 | + | + | + | – |

| | | | | | | |
|---|---|---|---|---|---|---|
| SPO_RS14070 | *nuoI* | 0.91 | + | + | + | – |
| SPO_RS02510 | *rplE* | 0.91 | + | + | + | – |
| SPO_RS18080 | *recR* | 0.91 | + | + | + | + |
| SPO_RS00750 | *mutM* | 0.91 | + | + | + | + |
| SPO_RS10555 | *hfq* | 0.91 | + | + | + | – |
| SPO_RS02520 | *rpsH* | 0.91 | + | + | + | – |
| SPO_RS06880 | *metZ* | 0.91 | + | + | + | – |
| SPO_RS02575 | *rpsK* | 0.90 | + | + | + | – |
| SPO_RS16025 | *atpG* | 0.90 | + | + | + | + |
| SPO_RS15735 | *ftsH* | 0.90 | + | – | + | – |
| SPO_RS02460 | *rplP* | 0.90 | + | + | + | + |
| SPO_RS06970 | *ilvC* | 0.90 | + | + | + | – |
| SPO_RS09590 | *purC* | 0.90 | + | + | + | + |
| SPO_RS00030 | *hemW* | 0.90 | + | + | + | – |
| SPO_RS15035 | *rpmG* | 0.90 | + | + | + | – |
| SPO_RS04950 | *lepA* | 0.90 | + | + | + | – |
| SPO_RS01985 | *trpS* | 0.89 | – | + | – | + |
| SPO_RS02530 | *rplR* | 0.89 | + | + | – | – |
| SPO_RS10905 | *trpC* | 0.89 | + | + | – | + |
| SPO_RS10995 | *purM* | 0.89 | + | + | – | – |
| SPO_RS10855 | *pdxH* | 0.88 | + | + | – | – |
| SPO_RS02505 | *rplX* | 0.88 | – | + | – | + |
| SPO_RS01095 | *leuD* | 0.88 | + | + | – | – |
| SPO_RS16095 | *hisC* | 0.88 | + | + | – | – |
| SPO_RS15055 | *gatC* | 0.88 | + | + | – | – |
| SPO_RS05895 | *hisB* | 0.88 | + | + | – | – |
| SPO_RS20160 | *infA* | 0.87 | + | + | – | – |
| SPO_RS20165 | *yhdE* | 0.87 | + | + | – | + |
| SPO_RS07025 | *carA* | 0.87 | + | + | – | – |
| SPO_RS01595 | *greA* | 0.87 | + | + | – | – |
| SPO_RS00040 | *rph* | 0.87 | + | + | – | – |
| SPO_RS04895 | *rpmB* | 0.87 | + | + | – | – |
| SPO_RS19695 | *hslV* | 0.86 | + | + | – | – |
| SPO_RS05440 | *yebC/pmpR* | 0.86 | + | + | – | + |
| SPO_RS08945 | *ribE* | 0.86 | + | + | – | – |
| SPO_RS11570 | *rpsR* | 0.86 | + | + | – | – |
| SPO_RS19770 | *mnmE* | 0.86 | – | + | – | + |
| SPO_RS15290 | *metF* | 0.85 | – | + | – | + |
| SPO_RS05870 | *hisA* | 0.85 | + | + | – | – |
| SPO_RS07090 | *lipB* | 0.85 | + | + | – | – |
| SPO_RS12415 | *ndk* | 0.85 | + | + | – | + |
| SPO_RS19415 | *rpsJ* | 0.84 | + | + | – | – |
| SPO_RS01830 | *sdhA* | 0.83 | – | + | – | + |
| SPO_RS19410 | *rplC* | 0.83 | + | – | – | – |

| | | | | | | |
|---|---|---|---|---|---|---|
| SPO_RS01505 | *yqgE/algH* | 0.83 | + | + | – | + |
| SPO_RS11635 | *glnA* | 0.83 | + | + | – | – |
| SPO_RS18530 | *clpS* | 0.82 | + | + | – | – |
| SPO_RS15540 | – | 0.81 | – | + | – | + |
| SPO_RS02655 | *argB* | 0.80 | + | + | – | + |
| SPO_RS05080 | *aat* | 0.80 | + | + | – | + |
| SPO_RS19745 | *coaE* | 0.80 | + | + | – | + |
| SPO_RS13295 | *tpiA* | 0.80 | + | + | – | + |
| SPO_RS16050 | *gloB* | 0.80 | + | + | – | + |
| SPO_RS17800 | *rplK* | 0.79 | + | + | – | + |
| SPO_RS10090 | *rpmA* | 0.78 | + | + | – | – |
| SPO_RS08955 | *ribB* | 0.77 | + | + | – | – |
| SPO_RS05050 | *clpP* | 0.75 | + | + | – | – |
| SPO_RS19485 | *rpsO* | 0.74 | + | + | – | – |
| SPO_RS09675 | *fumC* | 0.42 | – | + | – | + |

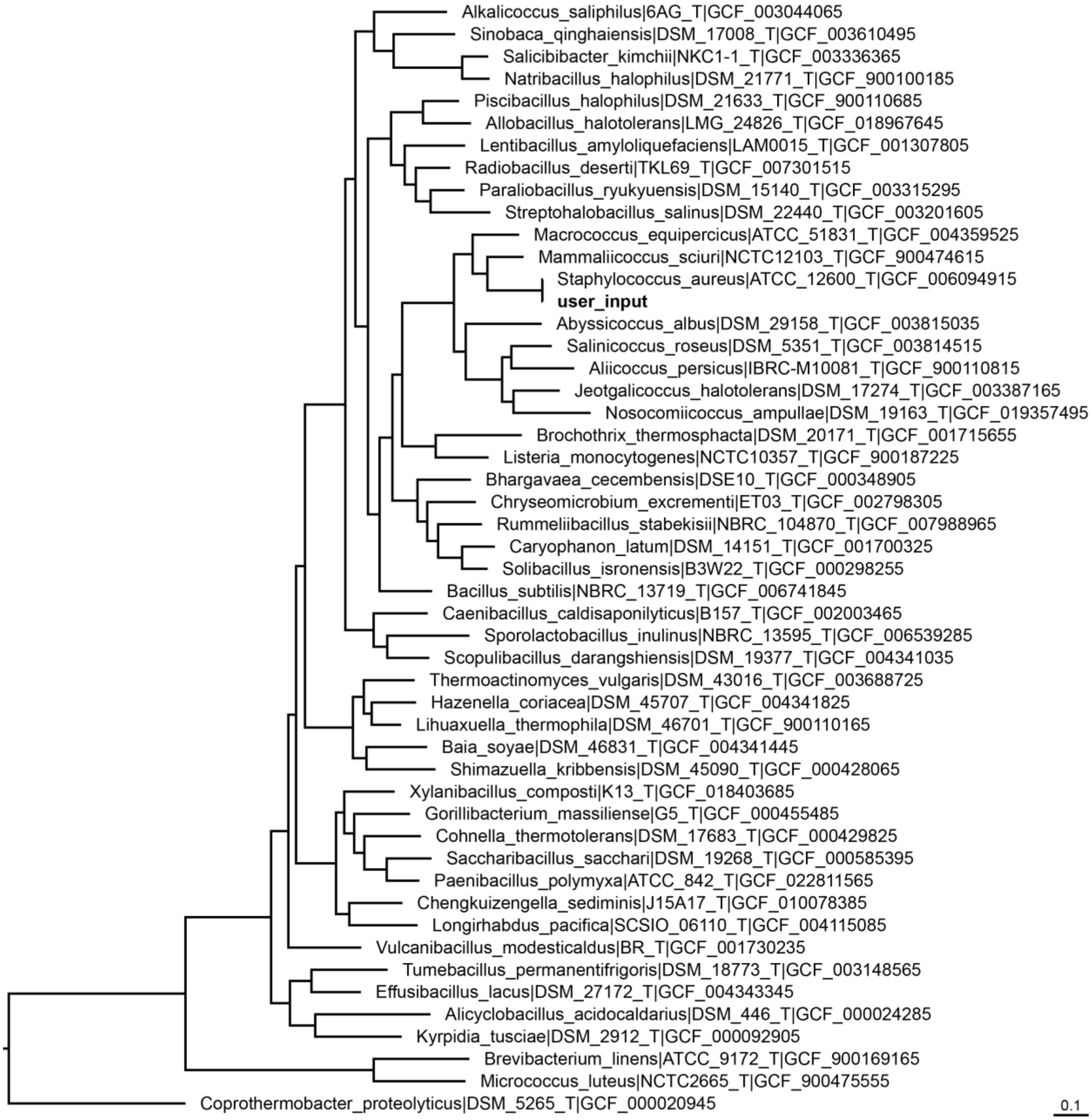


**Figure S1. Initial phylogeny constructed by PHANTASM when using the whole genome sequence of *Staphylococcus aureus* DSM 20231$^T$ as input.** The tree was constructed from a concatenated alignment of the core genes as described in Materials and Methods. The input genome (GCF_001027105.1) is bolded. All other genomes were automatically identified by PHANTASM. The tree was rooted on *Coprothermobacter proteolyticus* DSM 5265$^T$ (GCF_000020945). The scale bar indicates the number of substitutions per site. The tree was visualized with FigTree v1.4.4 (https://github.com/rambaut/figtree/releases) and is a graphical representation of the raw output of PHANTASM.

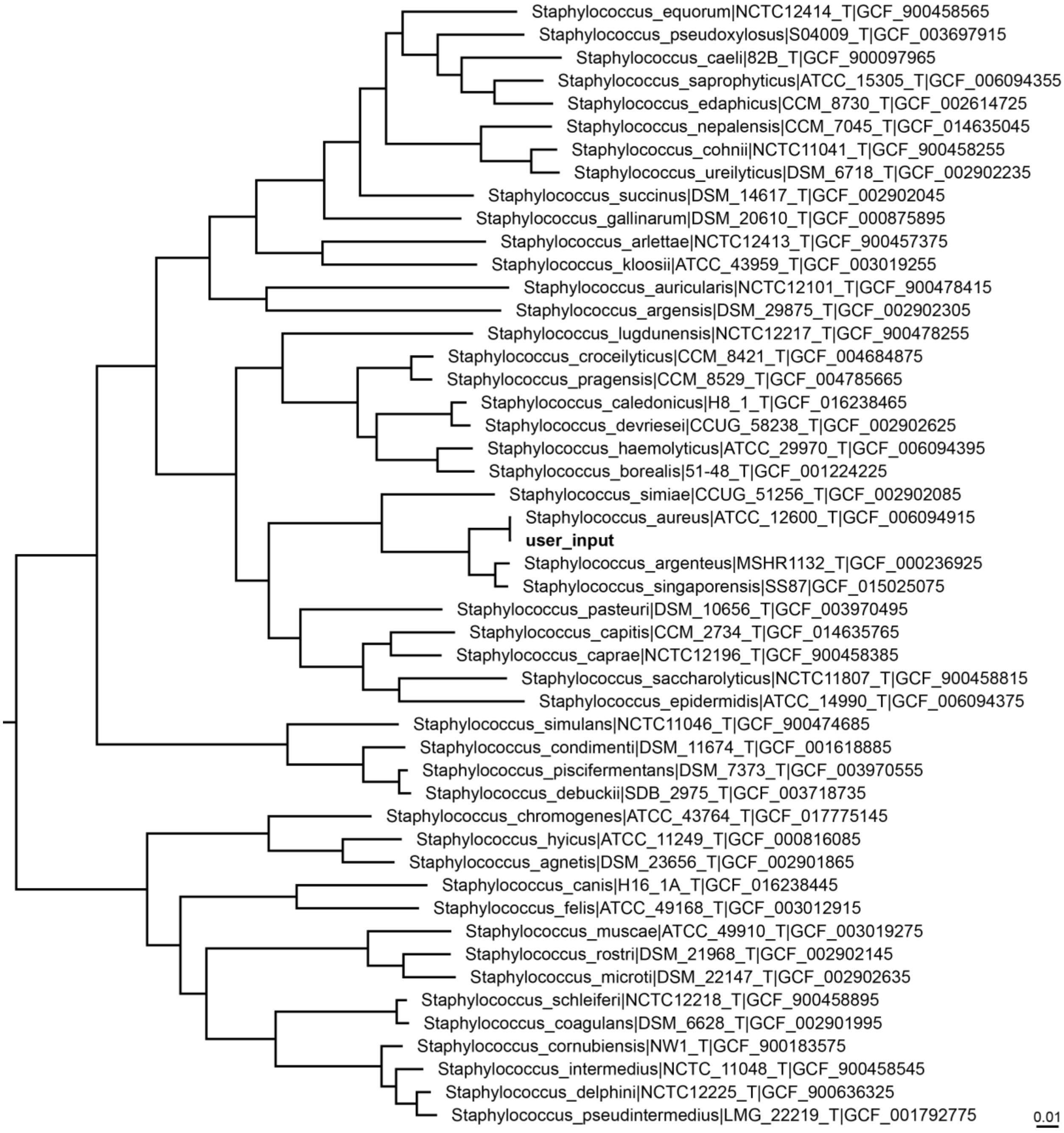


**Figure S2. Final phylogenomic reconstruction by PHANTASM when using the whole genome sequence of *Staphylococcus aureus* DSM 20231$^{T}$ as input.** The tree was constructed from a concatenated alignment of the core genes as described in Materials and Methods. The input genome (GCF_001027105.1) is bolded. All other genomes were automatically identified by PHANTASM. The tree was rooted on *Coprothermobacter proteolyticus* DSM 5265$^{T}$ (GCF_000020945), which was omitted. The scale bar indicates the number of substitutions per site. The tree was visualized with FigTree v1.4.4 (https://github.com/rambaut/figtree/releases) and is a graphical representation of the raw output of PHANTASM.

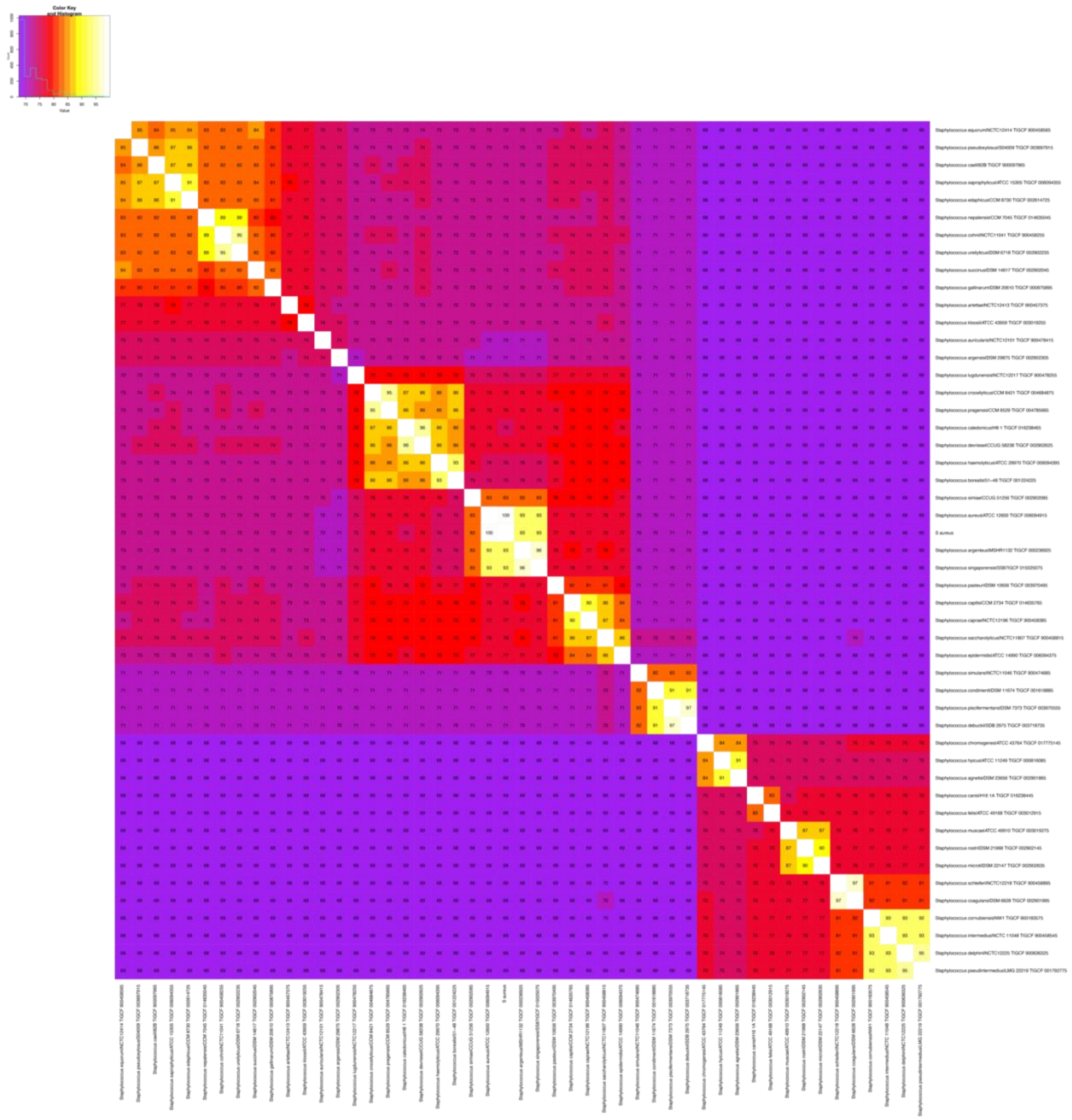

**Figure S3. Average amino acid identity heatmap automatically produced by PHANTASM when using the whole genome sequence of *Staphylococcus aureus* DSM 20231$^{T}$ as input.** The figure shows the raw output generated by PHANTASM without any modifications. Taxa are listed in the same order as is found in the tree in **Figure S2**. The distribution of values is shown in the top left corner. The input genome used was GCF_001027105.1.

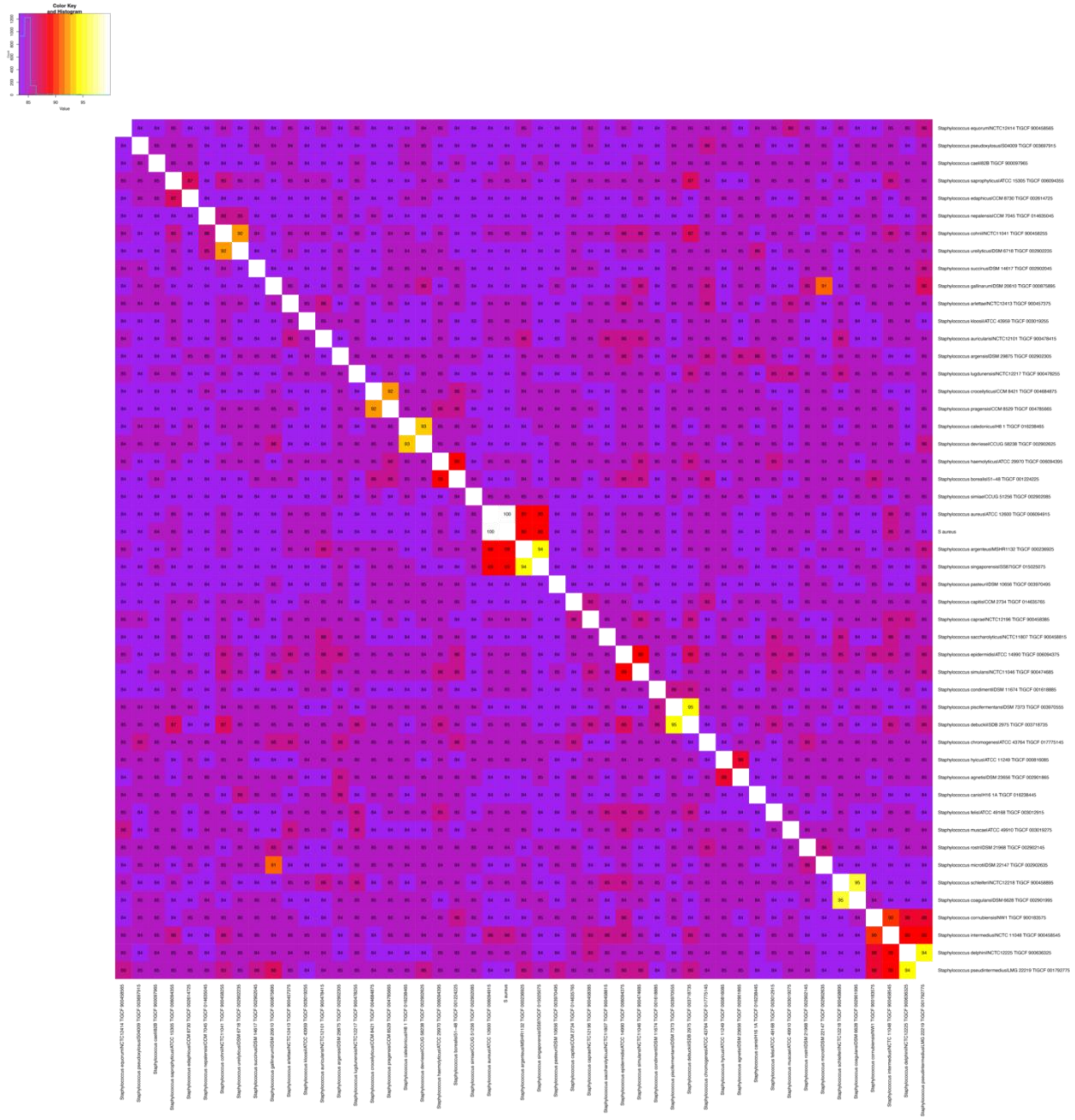

**Figure S4. Average nucleotide identity heatmap automatically produced by PHANTASM when using the whole genome sequence of *Staphylococcus aureus* DSM 20231$^{T}$ as input.** The figure shows the raw output generated by PHANTASM without any modifications. Taxa are listed in the same order as is found in the tree in **Figure S2**. The distribution of values is shown in the top left corner. The input genome used was GCF_001027105.1.

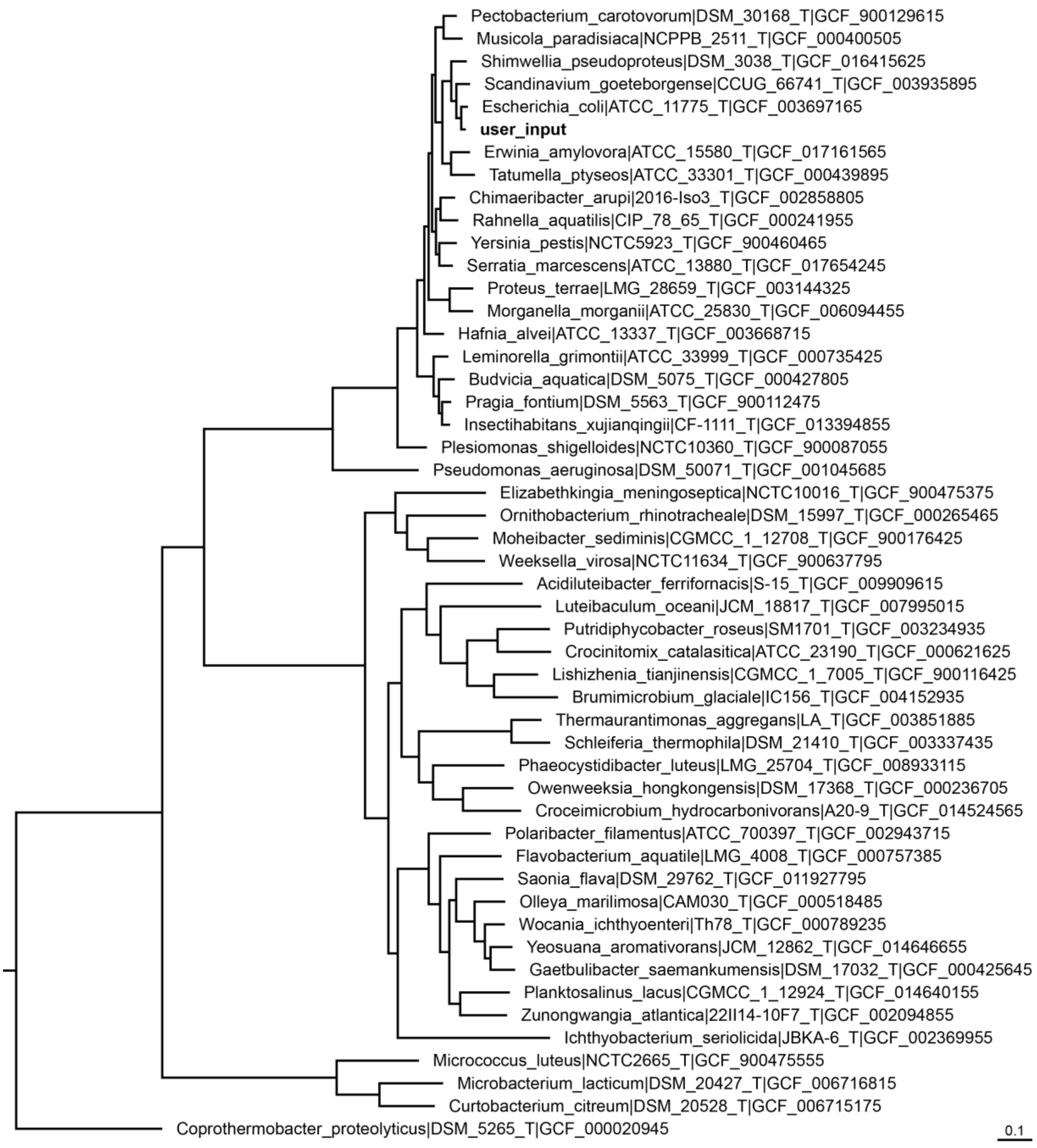


**Figure S5. Initial phylogeny constructed by PHANTASM when using the whole genome sequence of *Salmonella enterica* LT2$^{T}$ as input.** The tree was constructed from a concatenated alignment of the core genes as described in Materials and Methods. The input genome (GCF_000006945.2) is bolded. All other genomes were automatically identified by PHANTASM. The tree was rooted on *Coprothermobacter proteolyticus* DSM 5265$^{T}$ (GCF_000020945). The scale bar indicates the number of substitutions per site. The tree was visualized with FigTree v1.4.4 (https://github.com/rambaut/figtree/releases) and is a graphical representation of the raw output of PHANTASM.

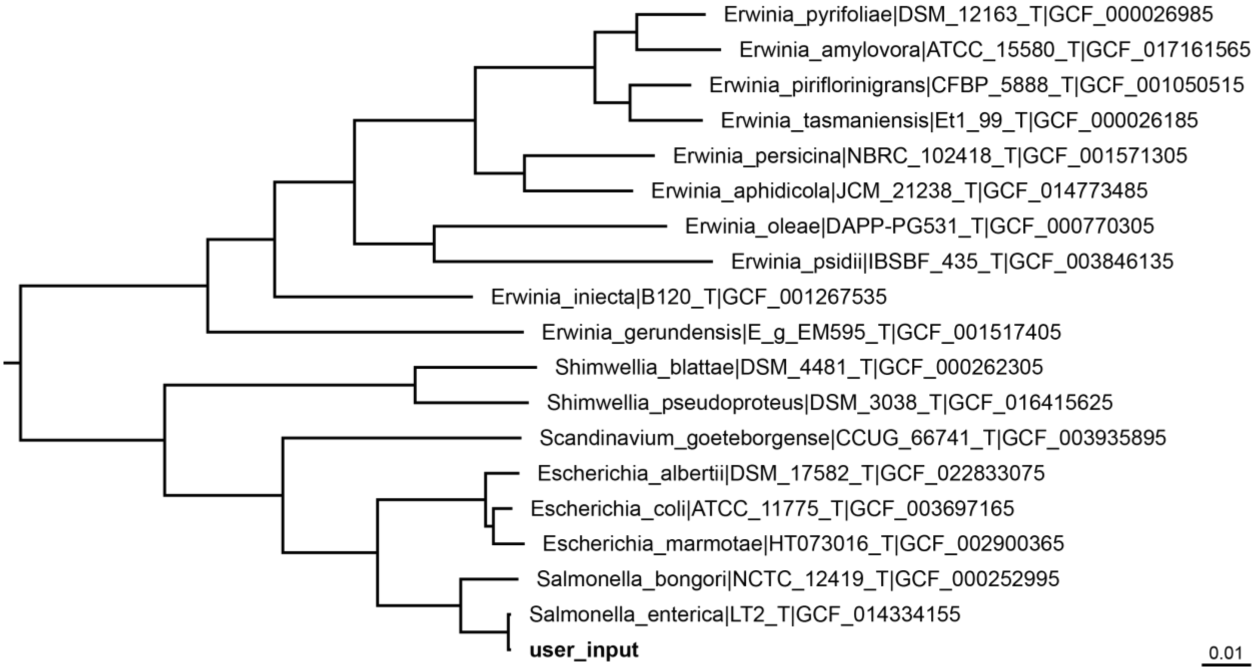


**Figure S6. Final phylogenomic reconstruction by PHANTASM when using the whole genome sequence of *Salmonella enterica* LT2$^{T}$ as input.** The tree was constructed from a concatenated alignment of the core genes as described in Materials and Methods. The input genome (GCF_000006945.2) is bolded. All other genomes were automatically identified by PHANTASM. The tree was rooted on *Coprothermobacter proteolyticus* DSM 5265$^{T}$ (GCF_000020945), which was omitted. The scale bar indicates the number of substitutions per site. The tree was visualized with FigTree v1.4.4 (https://github.com/rambaut/figtree/releases) and is a graphical representation of the raw output of PHANTASM.

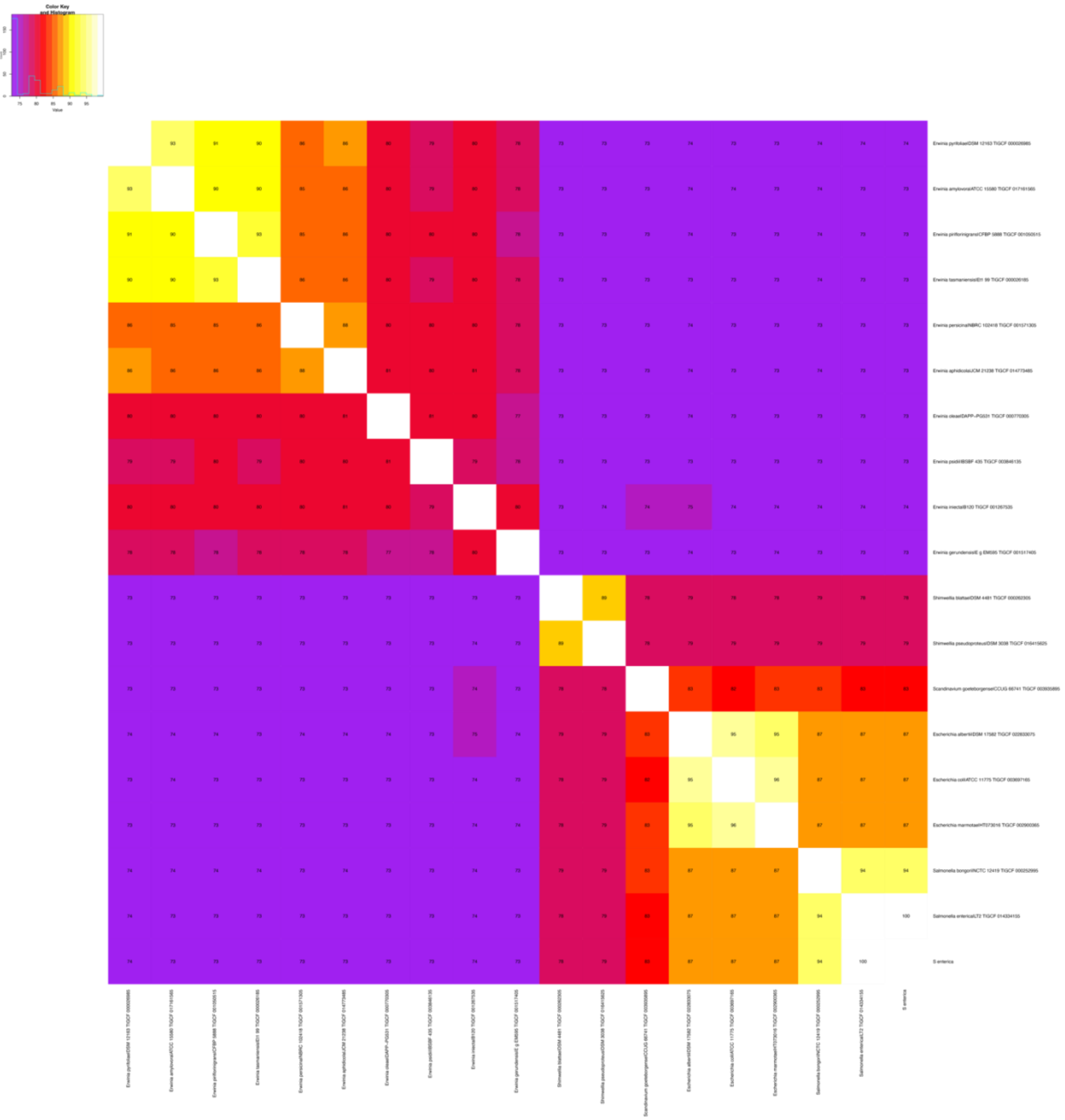

**Figure S7. Average amino acid identity heatmap automatically produced by PHANTASM when using the whole genome sequence of *Salmonella enterica* LT2$^{T}$ as input.** The figure shows the raw output generated by PHANTASM without any modifications. Taxa are listed in the same order as is found in the tree in **Figure S6**. The distribution of values is shown in the top left corner. The input genome used was GCF_000006945.2.

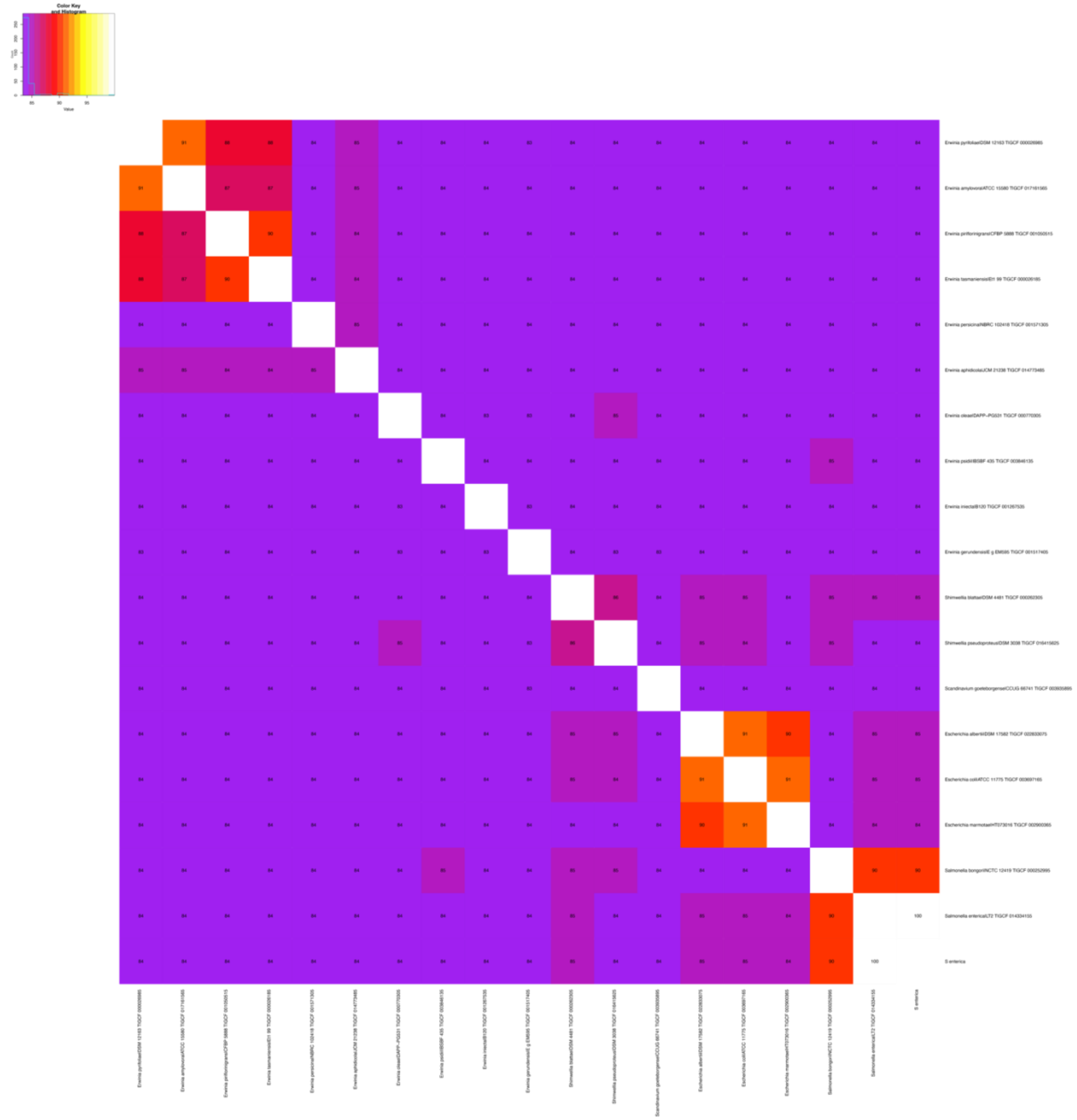

**Figure S8. Average nucleotide identity heatmap automatically produced by PHANTASM when using the whole genome sequence of *Salmonella enterica* LT2$^{T}$ as input.** The figure shows the raw output generated by PHANTASM without any modifications. Taxa are listed in the same order as is found in the tree in **Figure S6**. The distribution of values is shown in the top left corner. The input genome used was GCF_000006945.2.

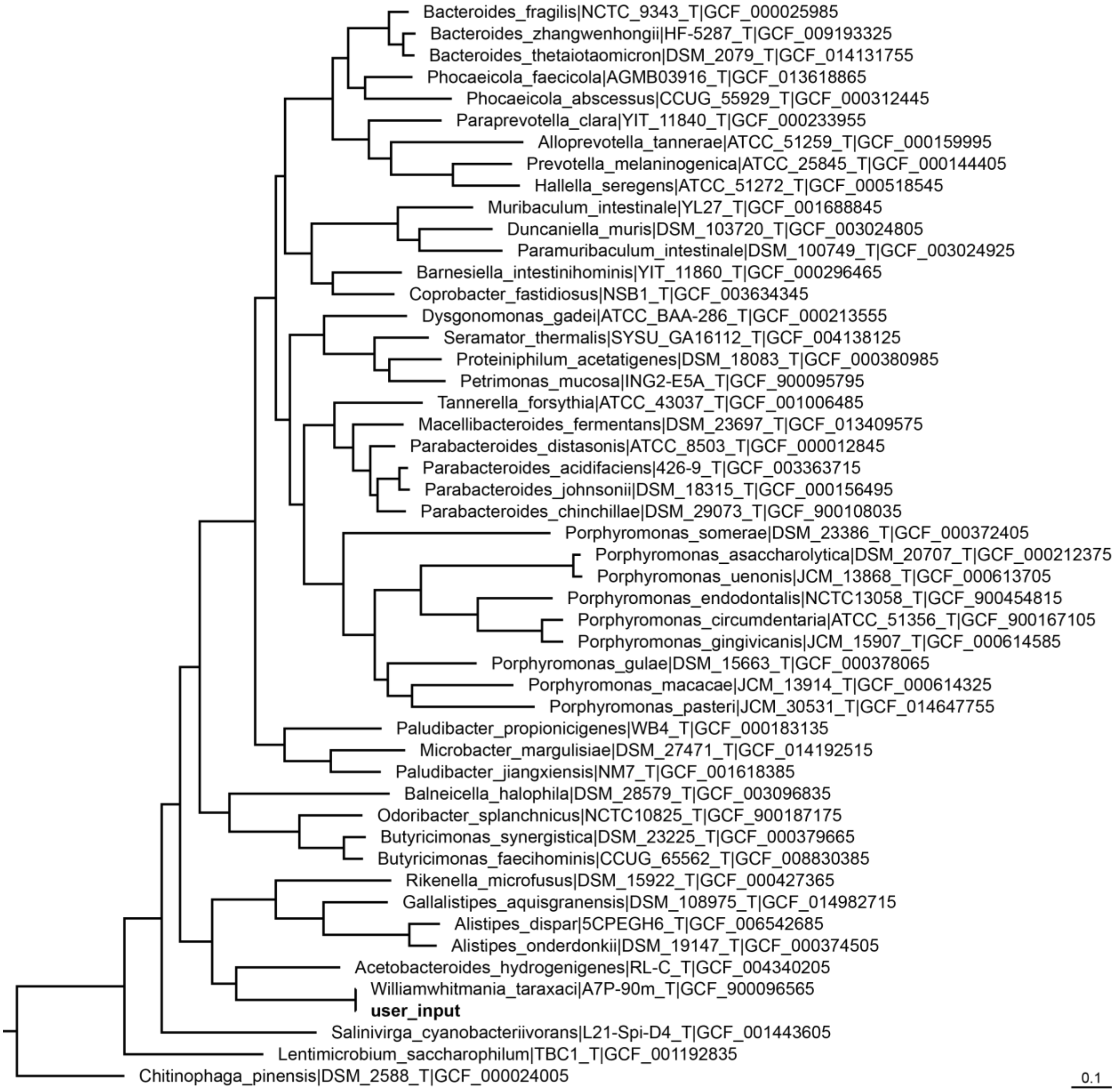


**Figure S9. Initial phylogeny constructed by PHANTASM when using the whole genome sequence of *Williamwhitmania taraxaci* A7P-90m$^{T}$ as input.** The tree was constructed from a concatenated alignment of the core genes as described in Materials and Methods. The input genome (GCF_900096565.1) is bolded. All other genomes were automatically identified by PHANTASM. The tree was rooted on *Chitinophaga pinensis* DSM 2588$^{T}$ (GCF_000024005). The scale bar indicates the number of substitutions per site. The tree was visualized with FigTree v1.4.4 (https://github.com/rambaut/figtree/releases) and is a graphical representation of the raw output of PHANTASM.

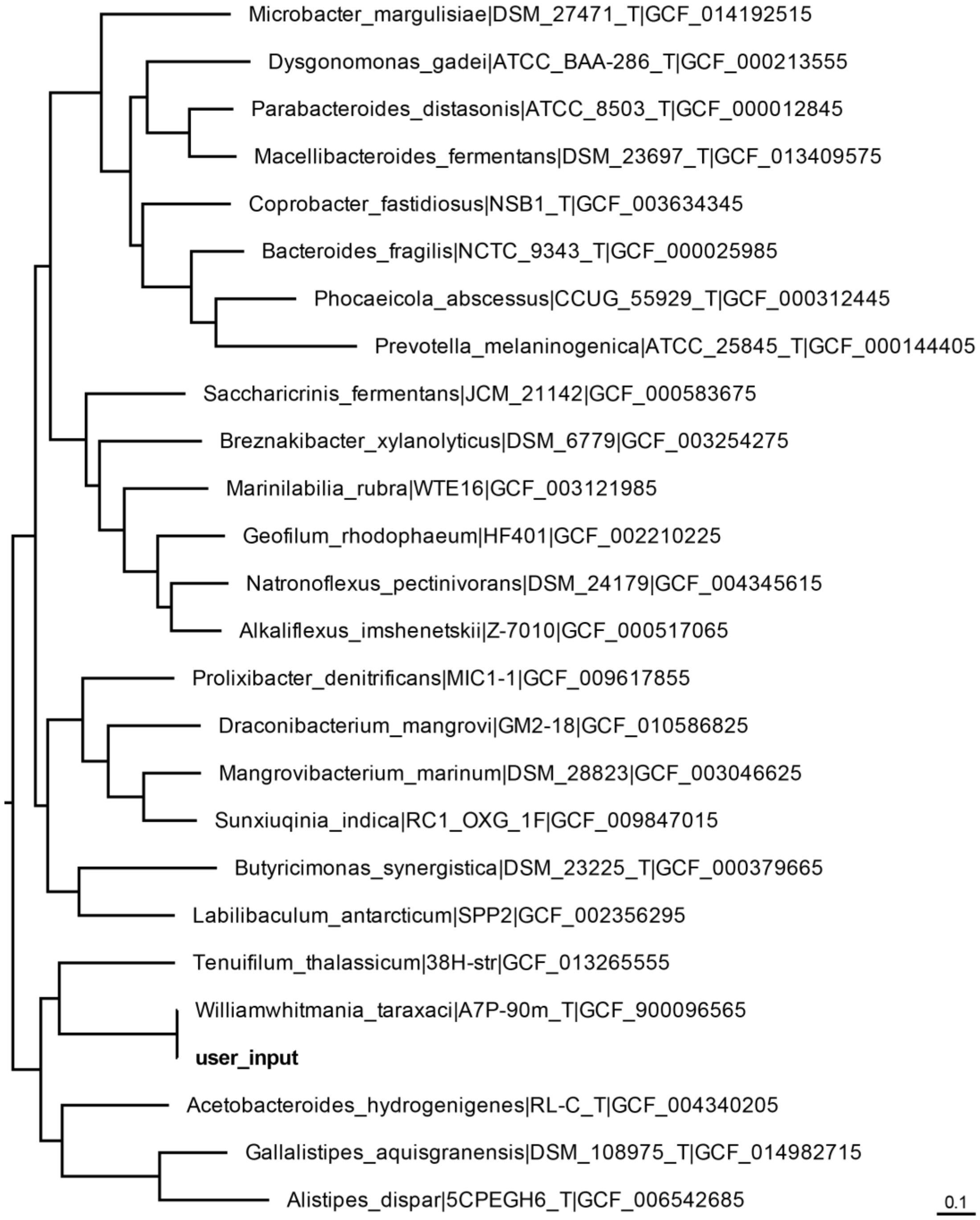


**Figure S10. Final phylogenomic reconstruction by PHANTASM when using the whole genome sequence of *Williamwhitmania taraxaci* A7P-90m$^{T}$ as input.** The tree was constructed from a concatenated alignment of the core genes as described in Materials and Methods. The input genome (GCF_900096565.1) is bolded. All other genomes were automatically identified by PHANTASM. The tree was rooted on *Chitinophaga pinensis* DSM 2588$^{T}$ (GCF_000024005), which was omitted. The scale bar indicates the number of substitutions per site. The tree was visualized with FigTree v1.4.4 (https://github.com/rambaut/figtree/releases) and is a graphical representation of the raw output of PHANTASM.

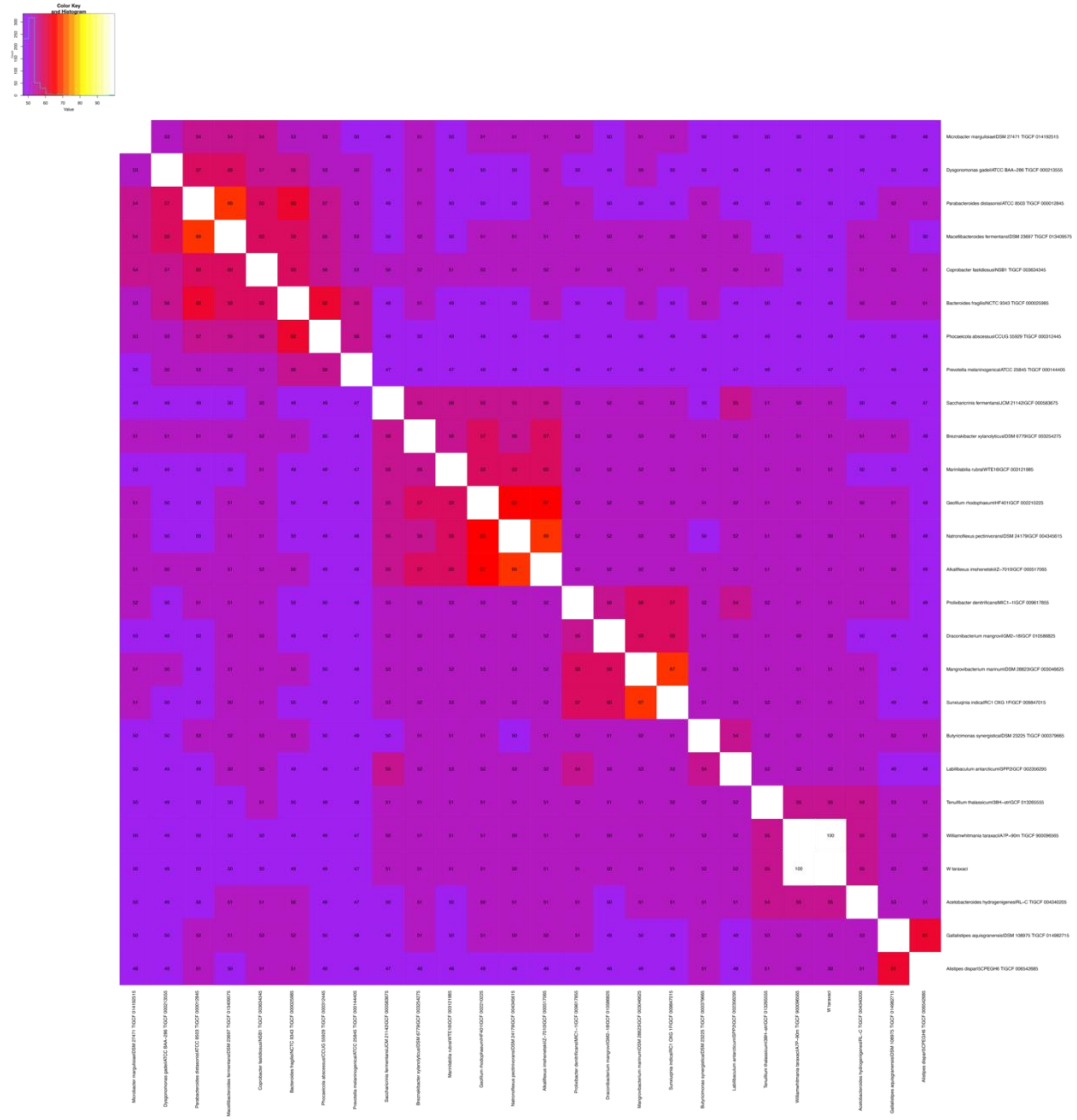

**Figure S11. Average amino acid identity heatmap automatically produced by PHANTASM when using the whole genome sequence of *Williamwhitmania taraxaci* A7P-90m$^{T}$ as input.** The figure shows the raw output generated by PHANTASM without any modifications. Taxa are listed in the same order as is found in the tree in **Figure S10**. The distribution of values is shown in the top left corner. The input genome used was GCF_900096565.1.

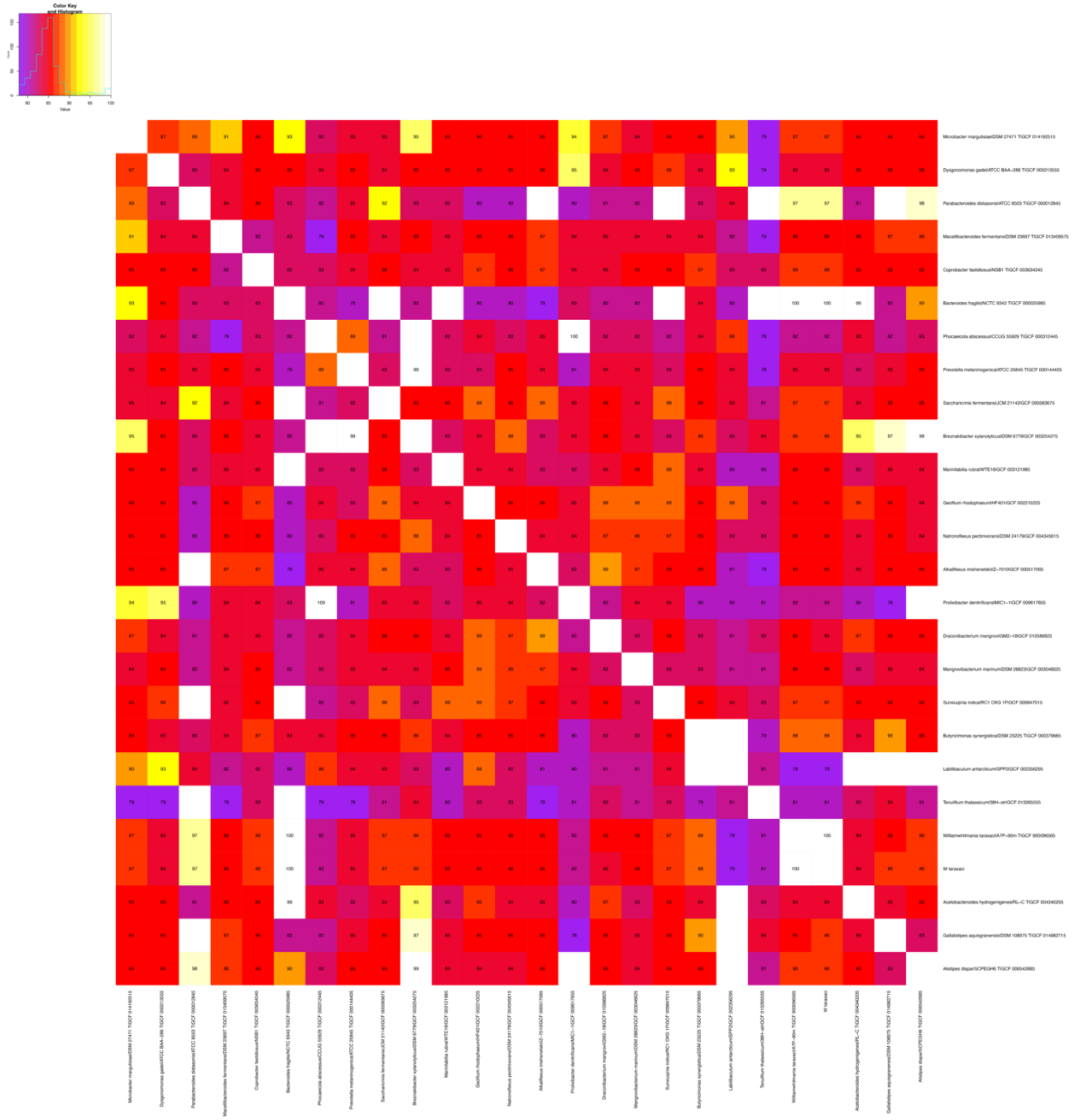

**Figure S12. Average nucleotide identity heatmap automatically produced by PHANTASM when using the whole genome sequence of *Williamwhitmania taraxaci* A7P-90m$^{T}$ as input.** The figure shows the raw output generated by PHANTASM without any modifications. Taxa are listed in the same order as is found in the tree in **Figure S10**. The distribution of values is shown in the top left corner. The input genome used was GCF_900096565.1.

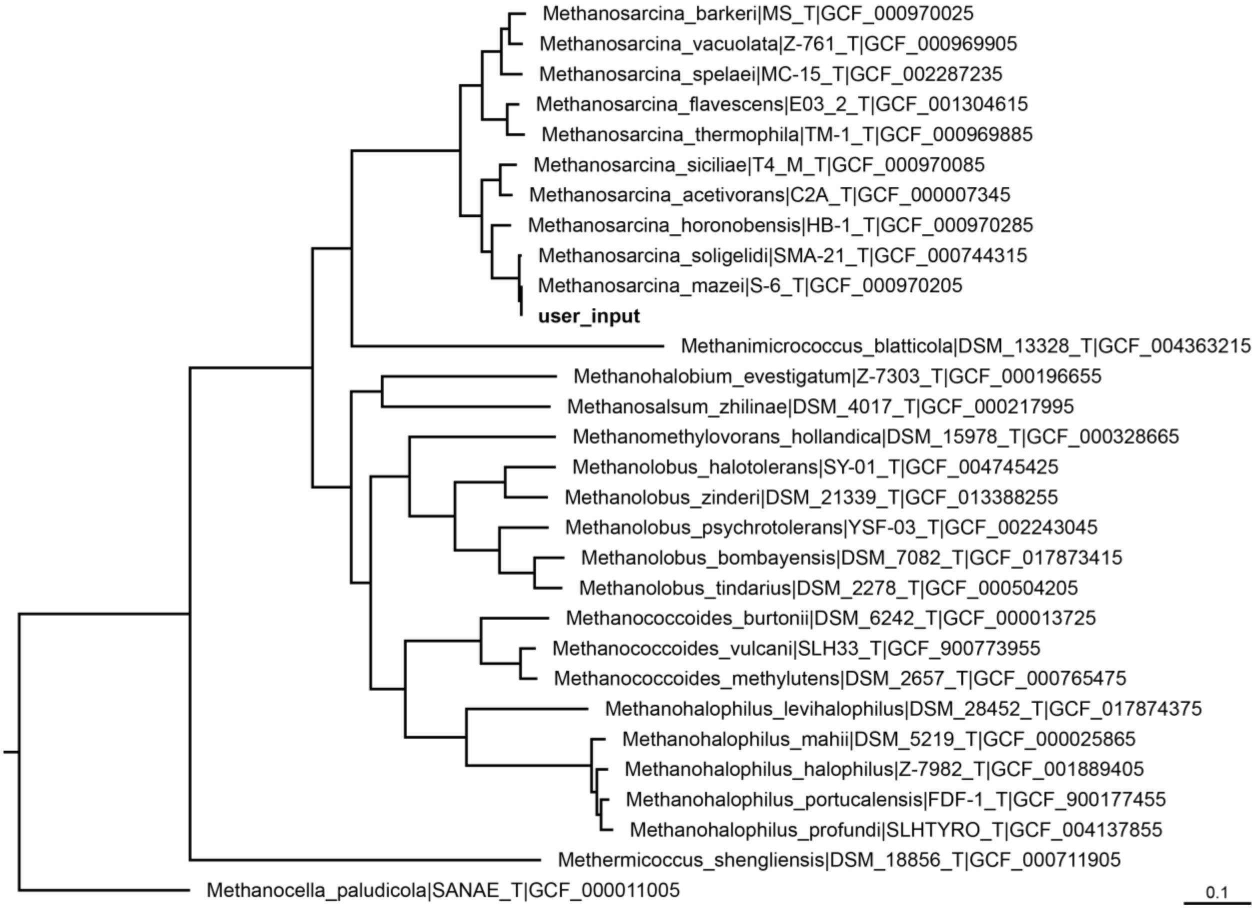


**Figure S13. Initial phylogeny constructed by PHANTASM when using the whole genome sequence of *Methanosarcina mazei* S-6$^{T}$ as input.** The tree was constructed from a concatenated alignment of the core genes as described in Materials and Methods. The input genome (GCF_000970205.1) is bolded. All other genomes were automatically identified by PHANTASM. The tree was rooted on *Methanocella paludicola* SANAE$^{T}$ (GCF_000011005). The scale bar indicates the number of substitutions per site. The tree was visualized with FigTree v1.4.4 (https://github.com/rambaut/figtree/releases) and is a graphical representation of the raw output of PHANTASM.

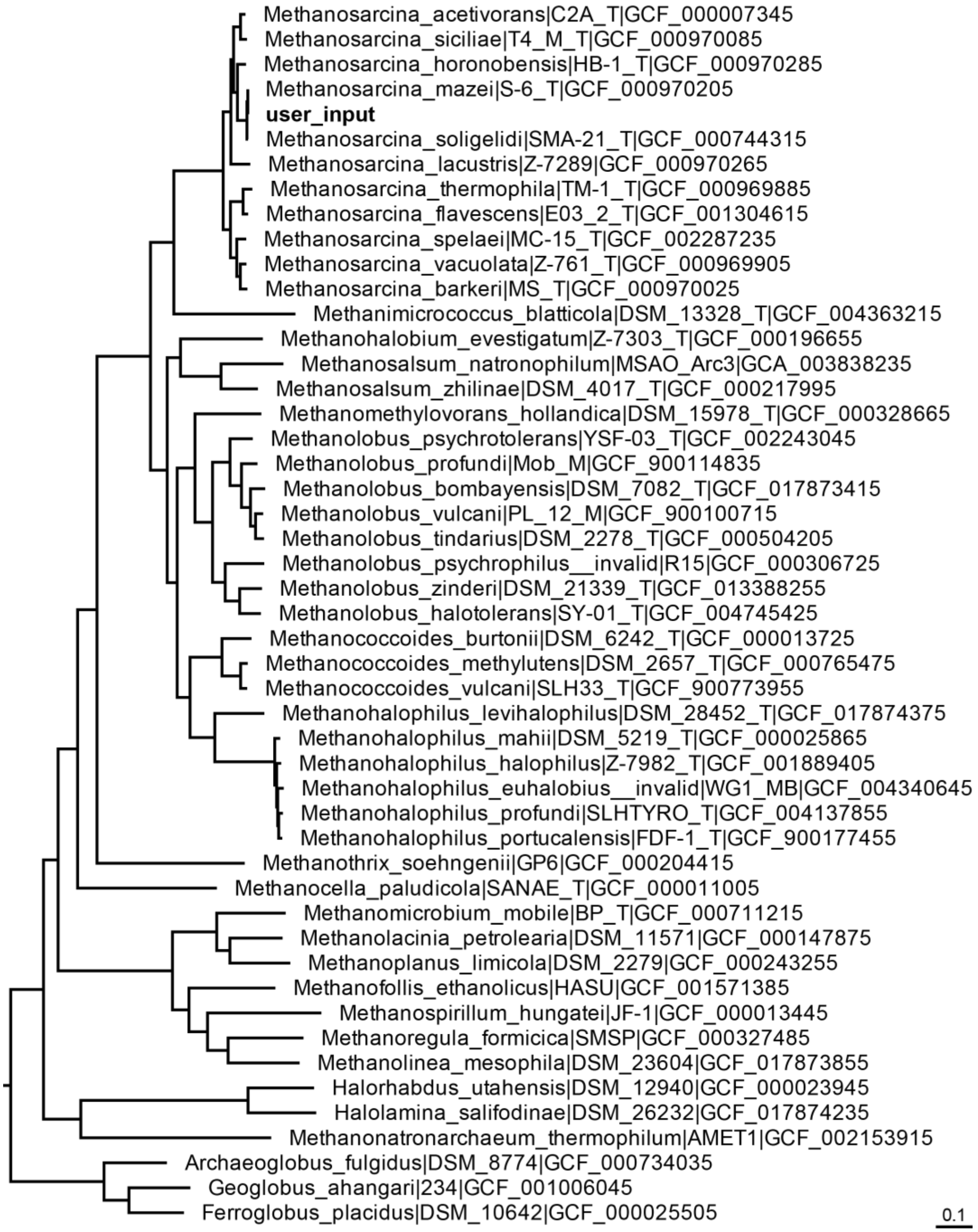


**Figure S14. Final phylogenomic reconstruction by PHANTASM when using the whole genome sequence of *Methanosarcina mazei* S-6$^{T}$ as input.** The tree was constructed from a concatenated alignment of the core genes as described in Materials and Methods. The input genome (GCF_000970205.1) is bolded. All other genomes were automatically identified by PHANTASM. The tree was rooted on *Nitrososphaera viennensis* EN76$^{T}$ (GCF_000698785), which was omitted. The scale bar indicates the number of substitutions per site. The tree was visualized with FigTree v1.4.4 (https://github.com/rambaut/figtree/releases) and is a graphical representation of the raw output of PHANTASM.

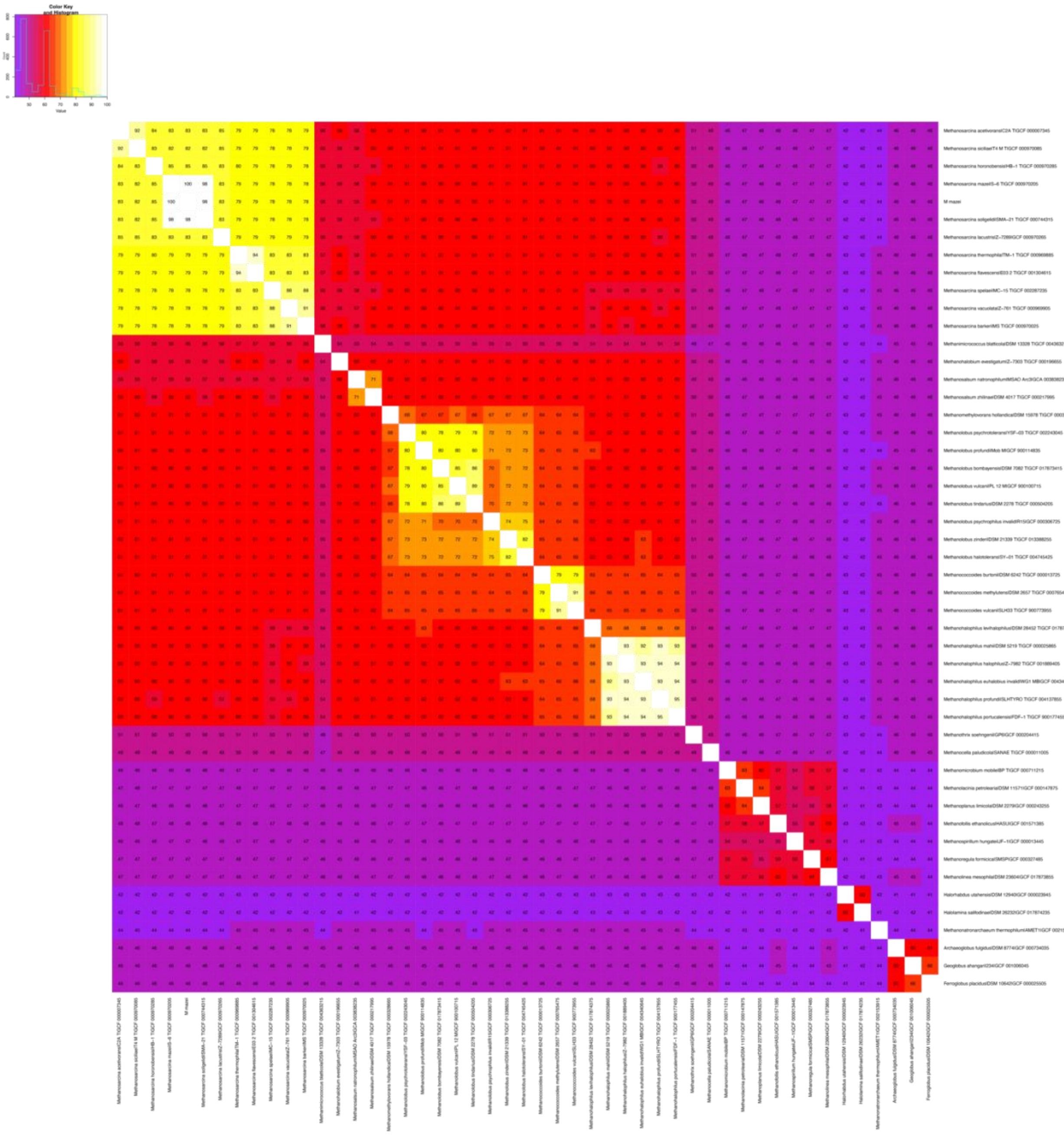

**Figure S15. Average amino acid identity heatmap automatically produced by PHANTASM when using the whole genome sequence of *Methanosarcina mazei* S-6$^{T}$ as input.** The figure shows the raw output generated by PHANTASM without any modifications. Taxa are listed in the same order as is found in the tree in **Figure S14**. The distribution of values is shown in the top left corner. The input genome used was GCF_000970205.1.

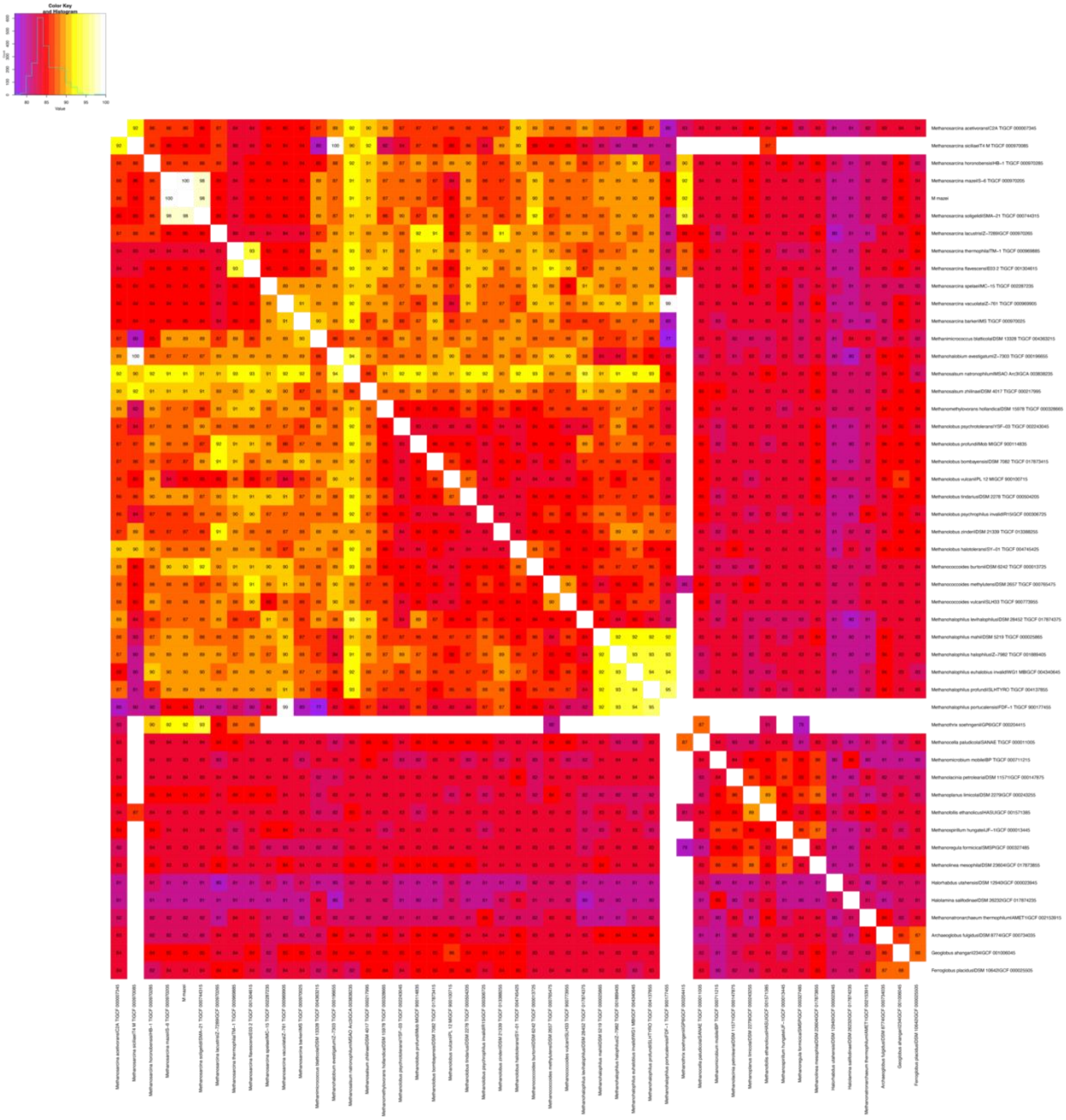

**Figure S16. Average nucleotide identity heatmap automatically produced by PHANTASM when using the whole genome sequence of *Methanosarcina mazei* S-6$^{T}$ as input.** The figure shows the raw output generated by PHANTASM without any modifications. Taxa are listed in the same order as is found in the tree in **Figure S14**. The distribution of values is shown in the top left corner. The input genome used was GCF_000970205.1.

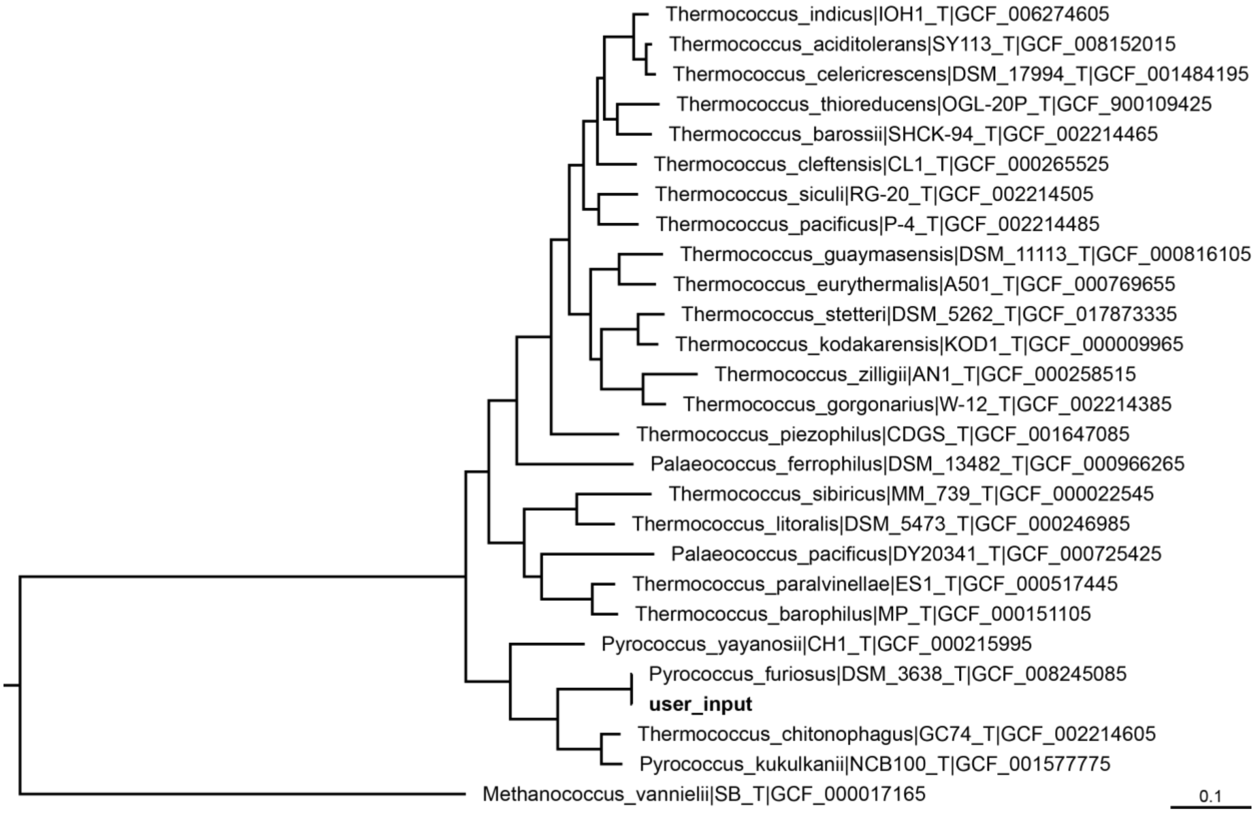


**Figure S17. Initial phylogeny constructed by PHANTASM when using the whole genome sequence of *Pyrococcus furiosus* DSM 3638$^T$ as input.** The tree was constructed from a concatenated alignment of the core genes as described in Materials and Methods. The input genome (GCF_008245085.1) is bolded. All other genomes were automatically identified by PHANTASM. The tree was rooted on *Methanococcus vannielii* SB$^T$ (GCF_000017165). The scale bar indicates the number of substitutions per site. The tree was visualized with FigTree v1.4.4 (https://github.com/rambaut/figtree/releases) and is a graphical representation of the raw output of PHANTASM.

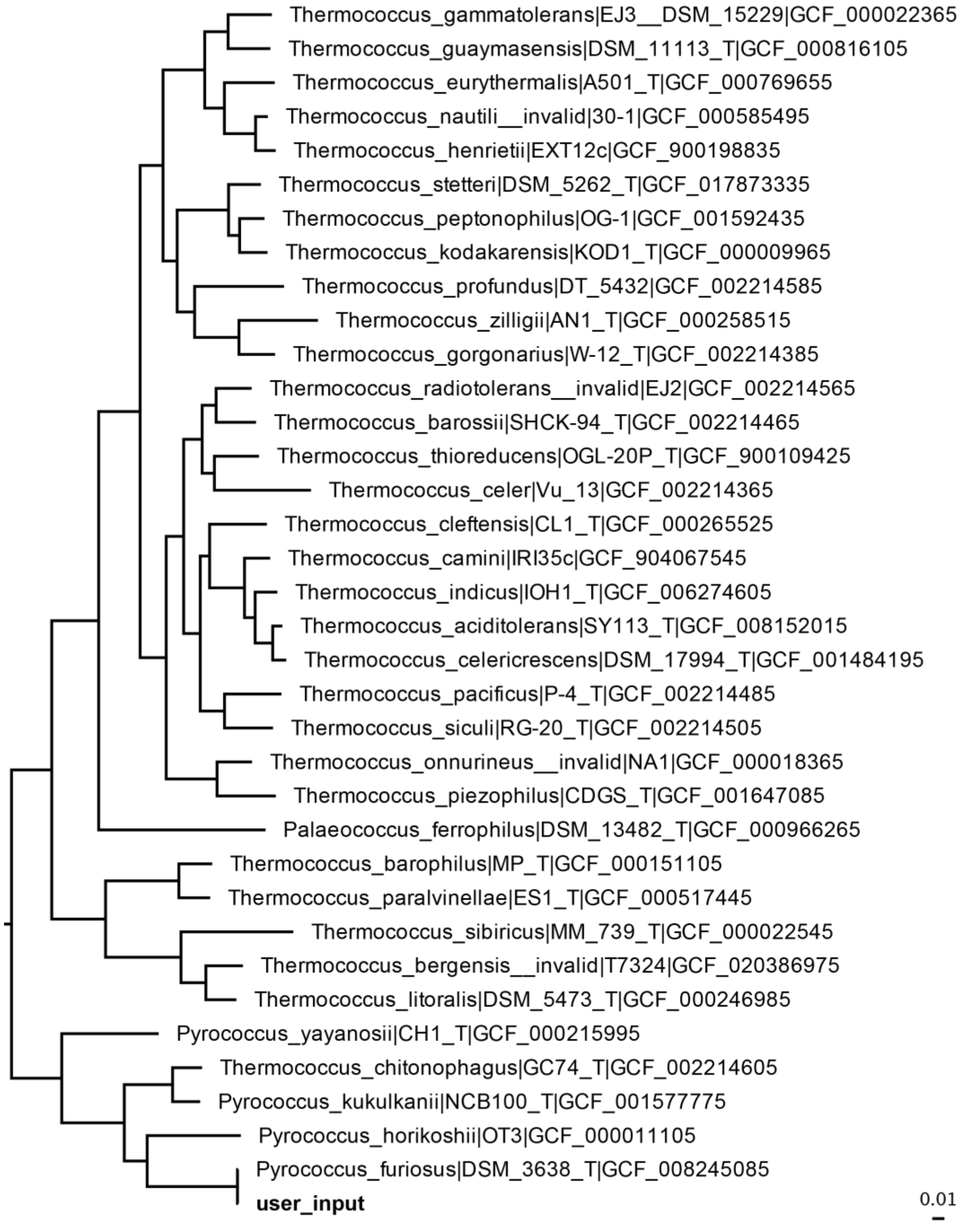


**Figure S18. Final phylogenomic reconstruction by PHANTASM when using the whole genome sequence of *Pyrococcus furiosus* DSM 3638$^{T}$ as input.** The tree was constructed from a concatenated alignment of the core genes as described in Materials and Methods. The input genome (GCF_008245085.1) is bolded. All other genomes were automatically identified by PHANTASM. The tree was rooted on *Nitrososphaera viennensis* EN76$^{T}$ (GCF_000698785), which was omitted. The scale bar indicates the number of substitutions per site. The tree was visualized with FigTree v1.4.4 (https://github.com/rambaut/figtree/releases) and is a graphical representation of the raw output of PHANTASM.

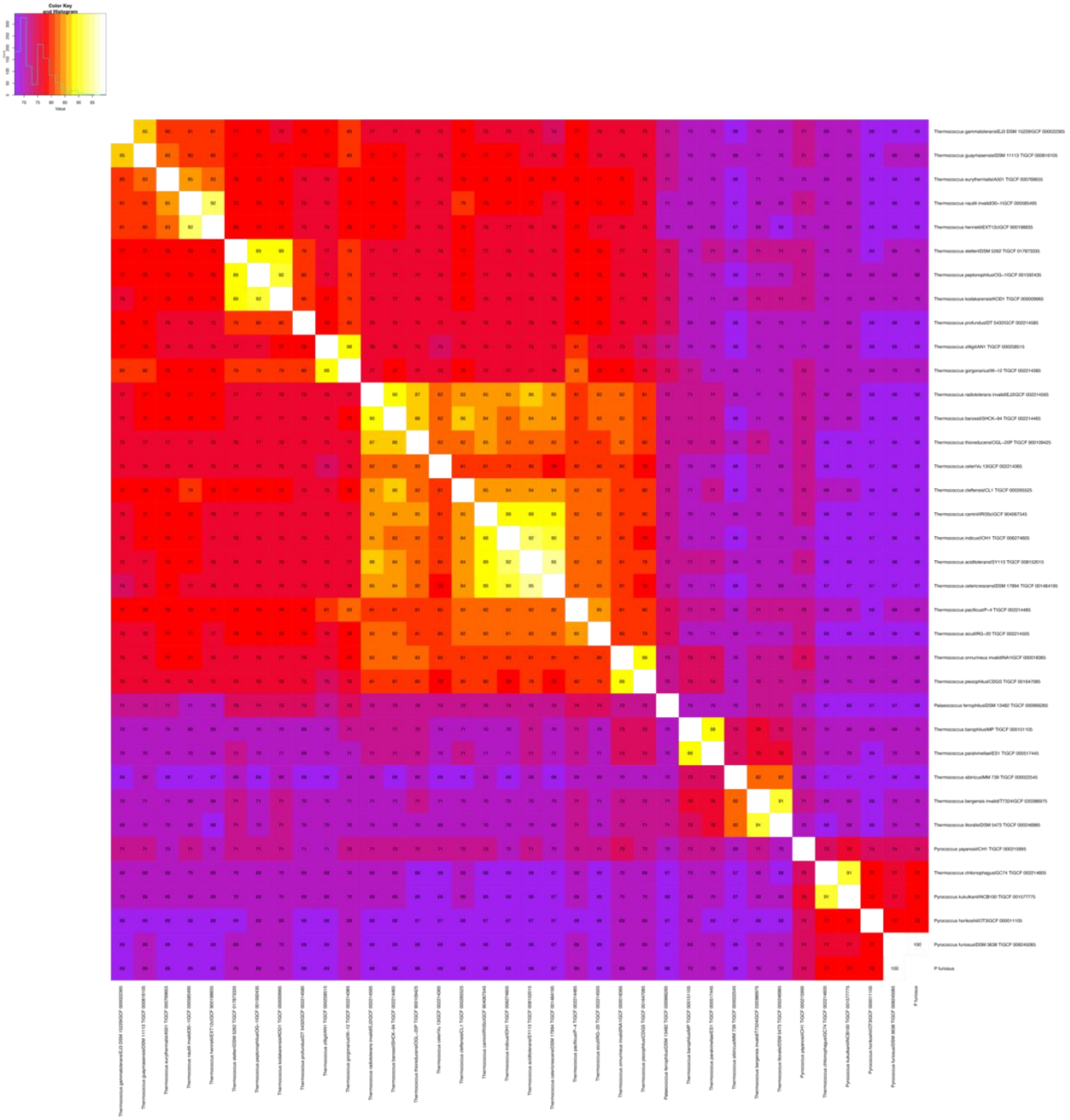

**Figure S19. Average amino acid identity heatmap automatically produced by PHANTASM when using the whole genome sequence of *Pyrococcus furiosus* DSM 3638$^{T}$ as input.** The figure shows the raw output generated by PHANTASM without any modifications. Taxa are listed in the same order as is found in the tree in **Figure S18**. The distribution of values is shown in the top left corner. The input genome used was GCF_008245085.1.

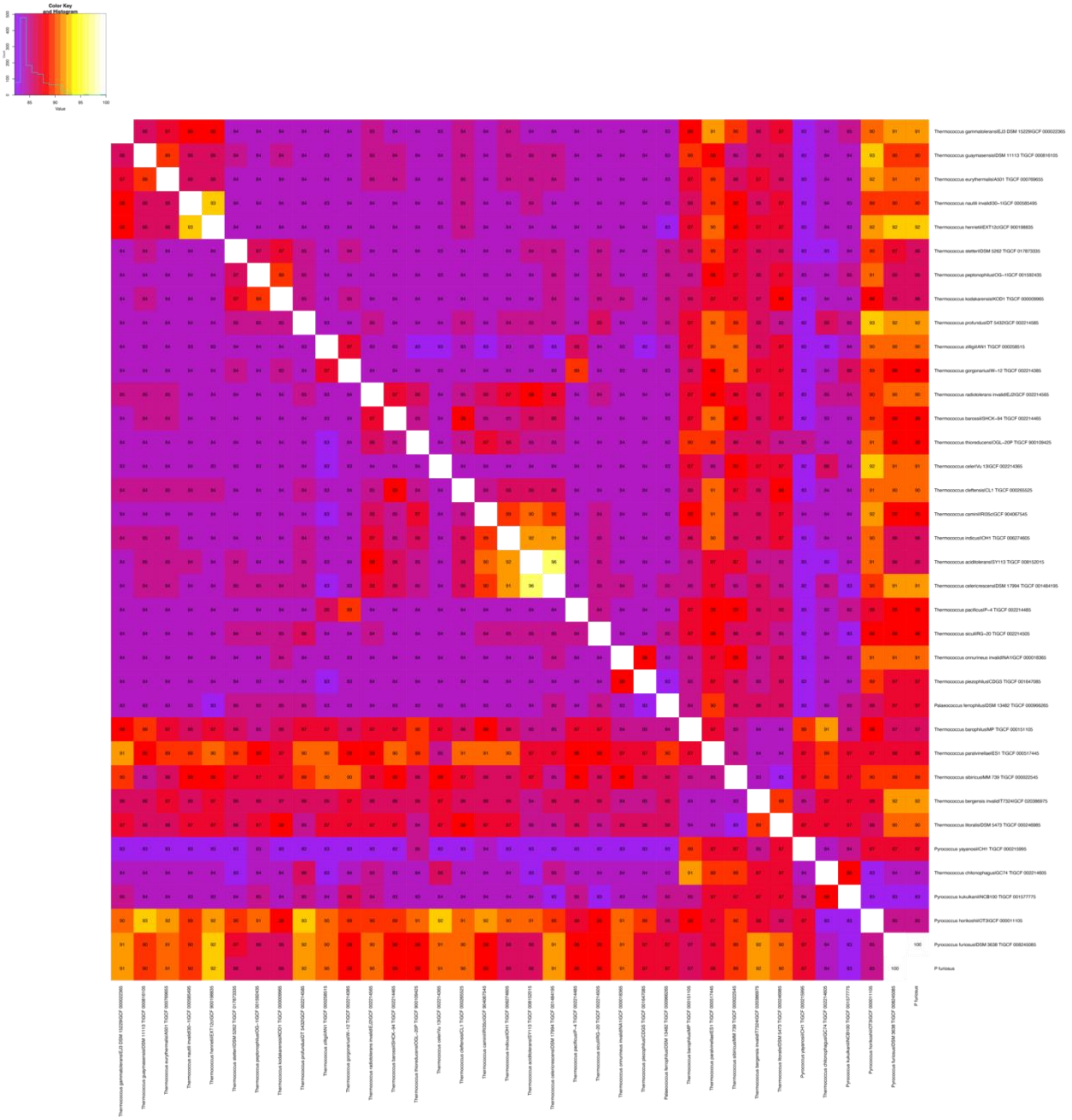

**Figure S20. Average nucleotide identity heatmap automatically produced by PHANTASM when using the whole genome sequence of *Pyrococcus furiosus* DSM 3638$^{T}$ as input.** The figure shows the raw output generated by PHANTASM without any modifications. Taxa are listed in the same order as is found in the tree in **Figure S18**. The distribution of values is shown in the top left corner. The input genome used was GCF_008245085.1.

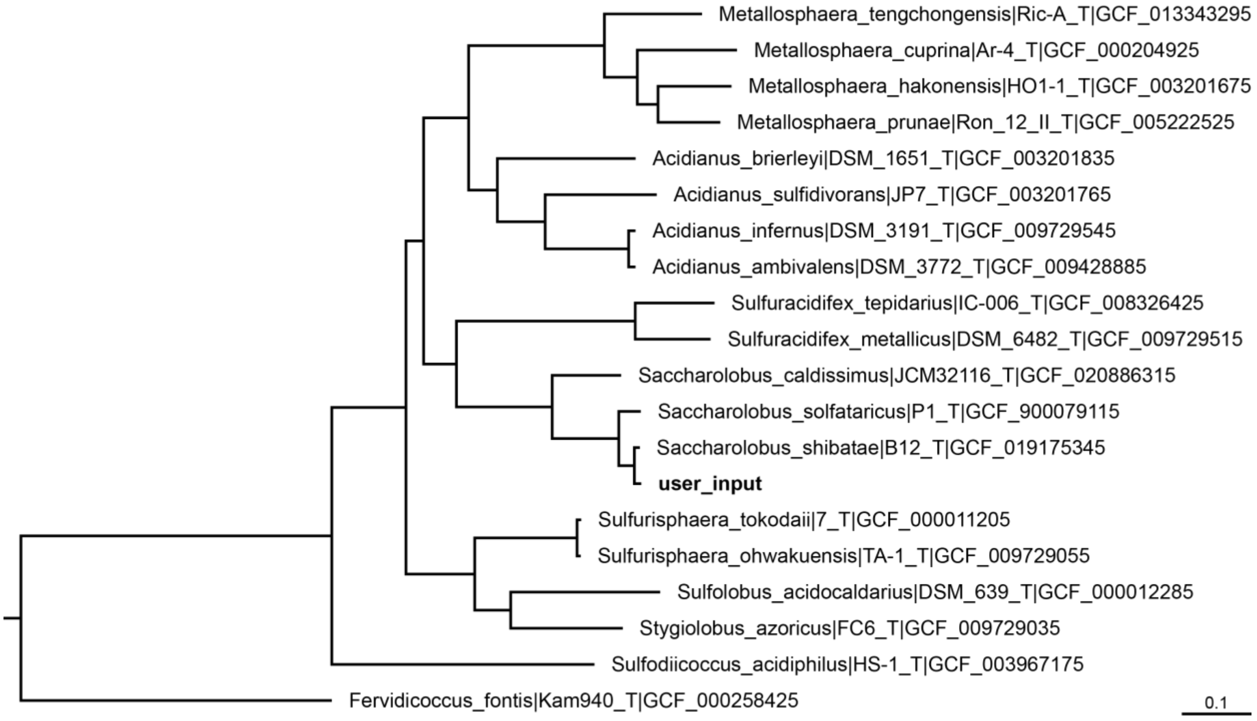


**Figure S21. Initial phylogeny constructed by PHANTASM when using the whole genome sequence of "*Sulfolobus islandicus*" Y.N.15.51 as input.** The tree was constructed from a concatenated alignment of the core genes as described in Materials and Methods. The input genome (GCF_000022485.1) is bolded. All other genomes were automatically identified by PHANTASM. The tree was rooted on *Fervidicoccus fontis* Kam940$^T$ (GCF_000258425). The scale bar indicates the number of substitutions per site. The tree was visualized with FigTree v1.4.4 (https://github.com/rambaut/figtree/releases) and is a graphical representation of the raw output of PHANTASM.

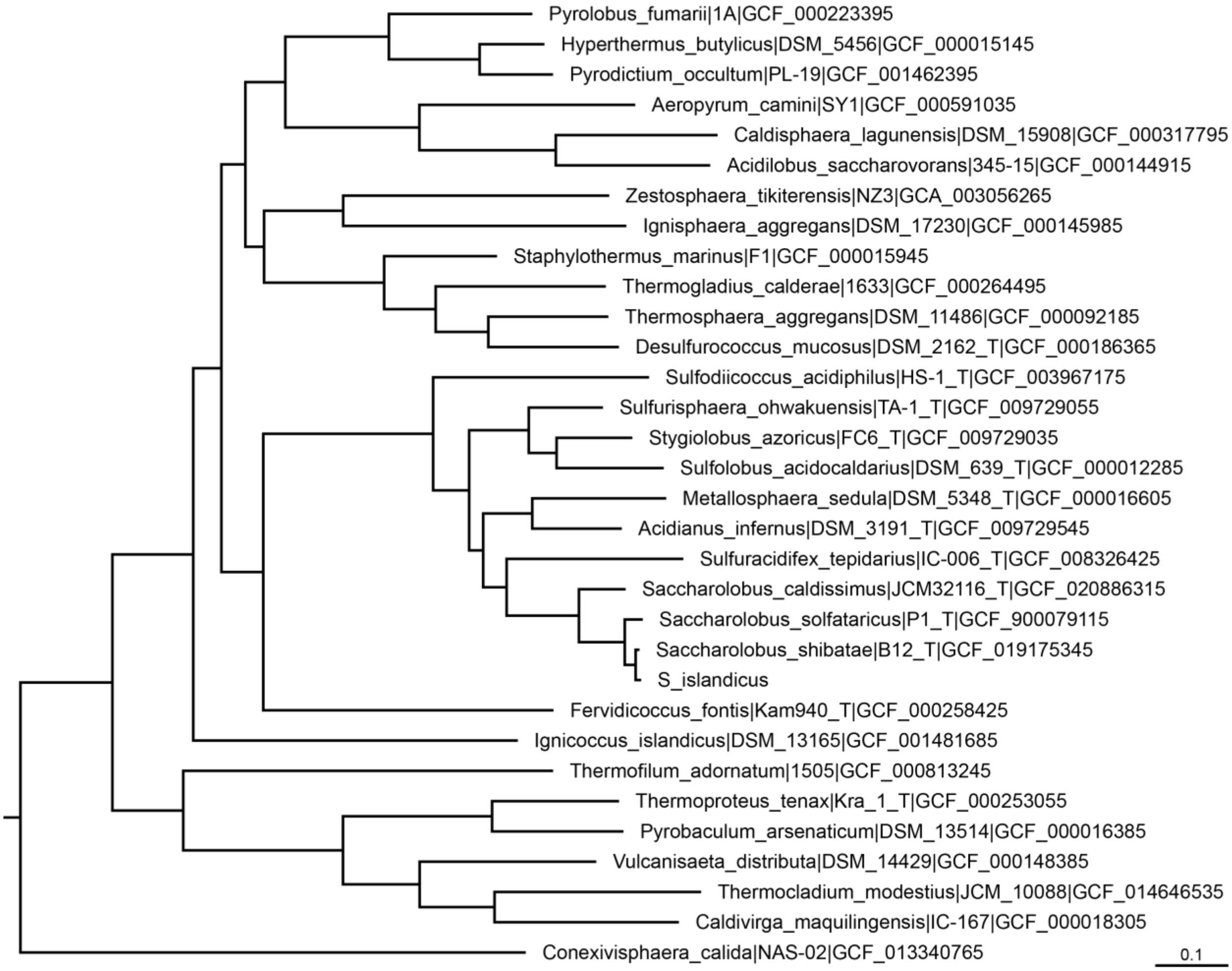


**Figure S22. Final phylogenomic reconstruction by PHANTASM when using the whole genome sequence of "*Sulfolobus islandicus*" Y.N.15.51 as input.** The tree was constructed from a concatenated alignment of the core genes as described in Materials and Methods. The input genome (GCF_000022485.1) is bolded. All other genomes were automatically identified by PHANTASM. The tree was rooted on *Methanobacterium formicicum* JCM 10132$^{T}$ (GCF_001316325), which was omitted. The scale bar indicates the number of substitutions per site. The tree was visualized with FigTree v1.4.4 (https://github.com/rambaut/figtree/releases) and is a graphical representation of the raw output of PHANTASM.

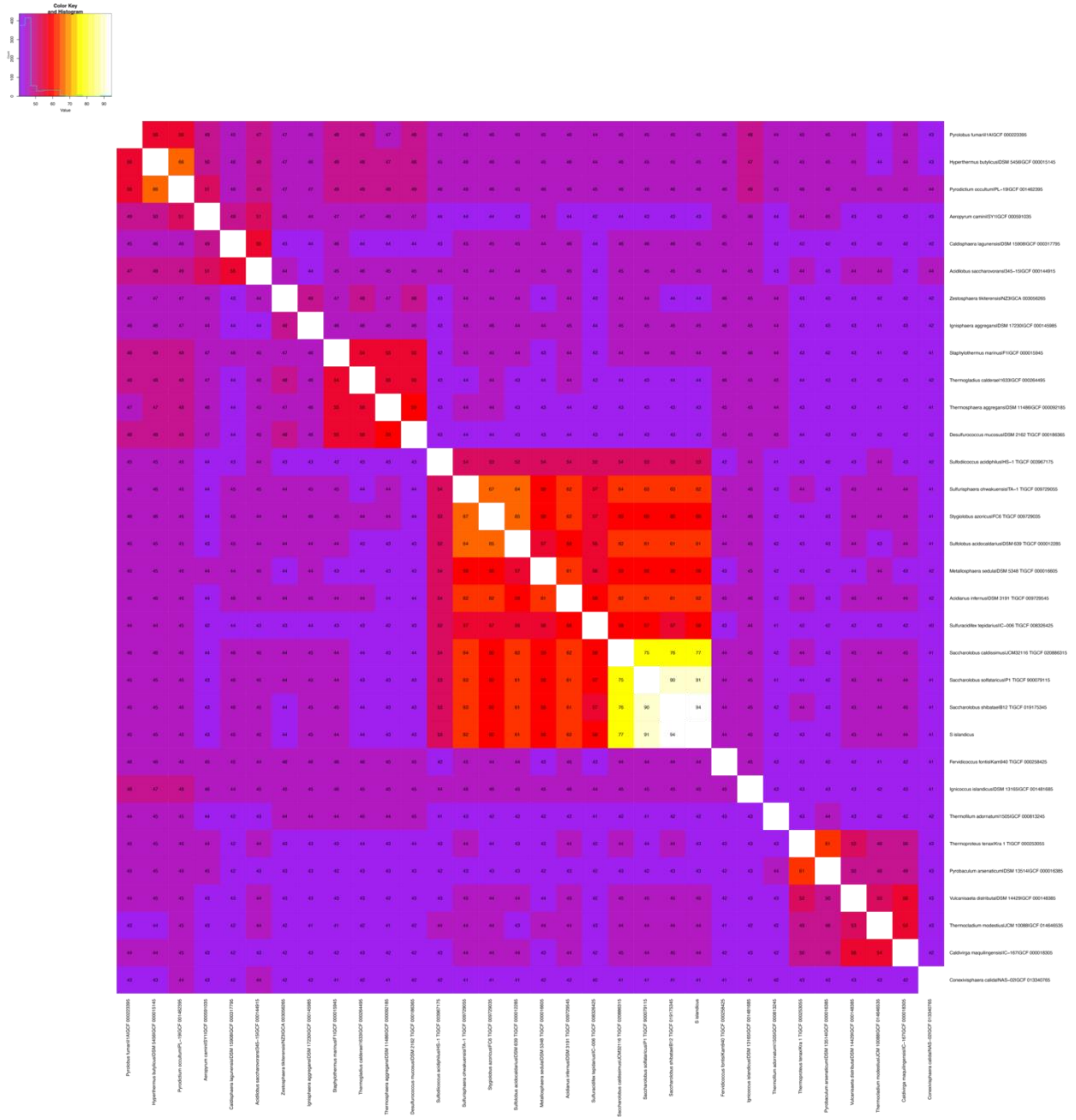

**Figure S23. Average amino acid identity heatmap automatically produced by PHANTASM when using the whole genome sequence of "*Sulfolobus islandicus*" Y.N.15.51 as input.** The figure shows the raw output generated by PHANTASM without any modifications. Taxa are listed in the same order as is found in the tree in **Figure S22**. The distribution of values is shown in the top left corner. The input genome used was GCF_000022485.1.

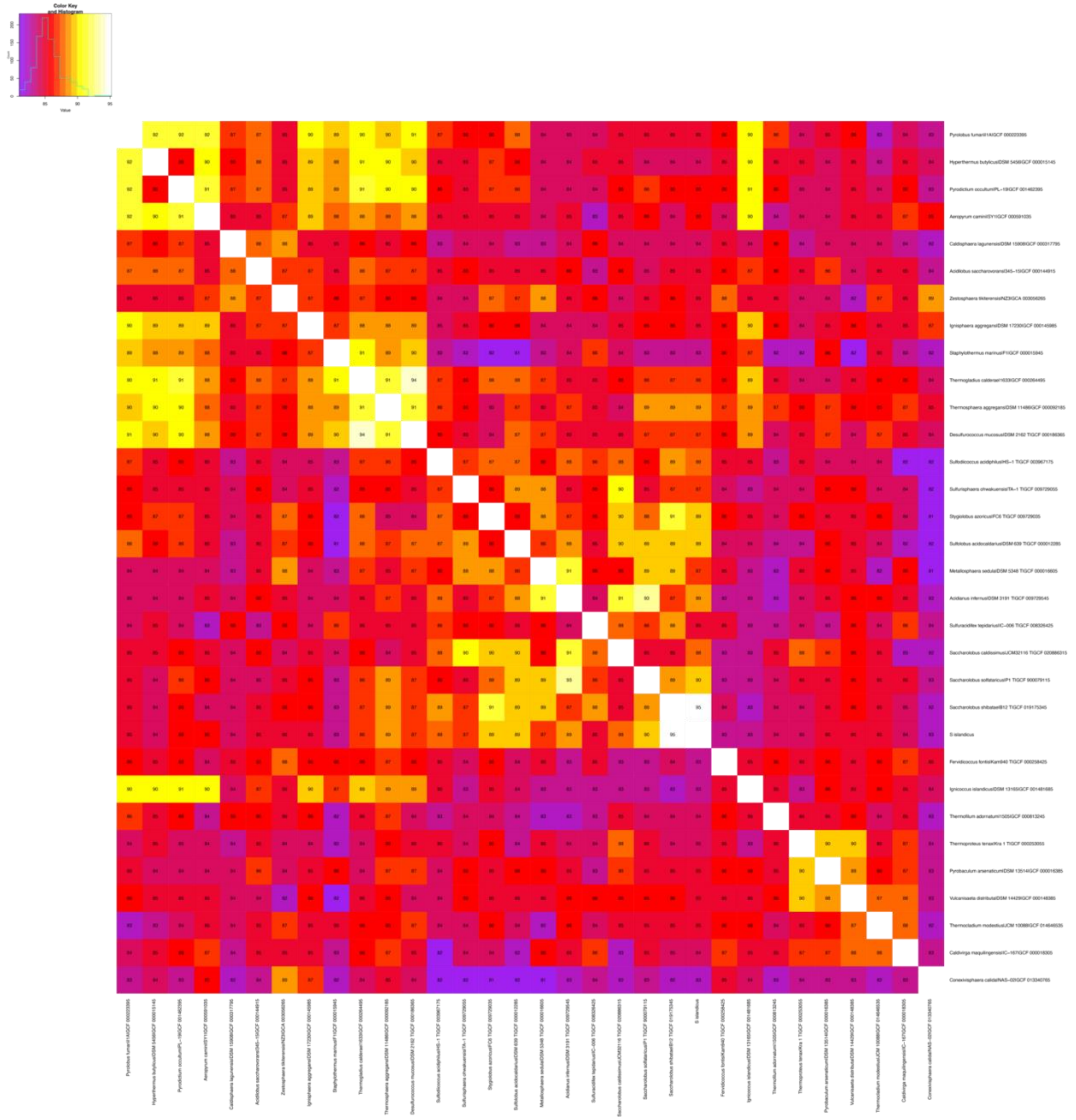

**Figure S24. Average nucleotide identity heatmap automatically produced by PHANTASM when using the whole genome sequence of "*Sulfolobus islandicus*" Y.N.15.51 as input.** The figure shows the raw output generated by PHANTASM without any modifications. Taxa are listed in the same order as is found in the tree in **Figure S22**. The distribution of values is shown in the top left corner. The input genome used was GCF_000022485.1.
