## Supplementary material for "Automating microbial taxonomy workflows with PHANTASM: PHylogenomic ANalyses for the TAxonomy and Systematics of Microbes": Table S2

**Table S2. Data from the List of Prokaryotic names with Standing in Nomenclature (LPSN) used by PHANTASM.** All websites were manually visited between January 5, 2023 and January 8, 2023.

| name | address |
|---|---|
| Anathece bachmannii | https://lpsn.dsmz.de/species/anathece-bachmannii |
| Anathece bacteriosa | https://lpsn.dsmz.de/species/anathece-bacteriosa |
| Anathece clathrata | https://lpsn.dsmz.de/species/anathece-clathrata |
| Anathece endophytica | https://lpsn.dsmz.de/species/anathece-endophytica |
| Anathece minutissima | https://lpsn.dsmz.de/species/anathece-minutissima |
| Anathece smithii | https://lpsn.dsmz.de/species/anathece-smithii |
| Anathece stratus | https://lpsn.dsmz.de/species/anathece-stratus |
| Aphanothece alascense | https://lpsn.dsmz.de/species/aphanothece-alascense |
| Aphanothece bachmannii | https://lpsn.dsmz.de/species/aphanothece-bachmannii |
| Aphanothece bacteriosa | https://lpsn.dsmz.de/species/aphanothece-bacteriosa |
| Aphanothece clathrata | https://lpsn.dsmz.de/species/aphanothece-clathrata |
| Aphanothece comasii | https://lpsn.dsmz.de/species/aphanothece-comasii |
| Aphanothece cylindracea | https://lpsn.dsmz.de/species/aphanothece-cylindracea |
| Aphanothece endophytica | https://lpsn.dsmz.de/species/aphanothece-endophytica |
| Aphanothece gelatinosa | https://lpsn.dsmz.de/species/aphanothece-gelatinosa |
| Aphanothece glebulenta | https://lpsn.dsmz.de/species/aphanothece-glebulenta |
| Aphanothece halophytica | https://lpsn.dsmz.de/species/aphanothece-halophytica |
| Aphanothece hegewaldii | https://lpsn.dsmz.de/species/aphanothece-hegewaldii |
| Aphanothece longior | https://lpsn.dsmz.de/species/aphanothece-longior |
| Aphanothece luteola | https://lpsn.dsmz.de/species/aphanothece-luteola |
| Aphanothece marina | https://lpsn.dsmz.de/species/aphanothece-marina |
| Aphanothece microscopica | https://lpsn.dsmz.de/species/aphanothece-microscopica |
| Aphanothece minutissima | https://lpsn.dsmz.de/species/aphanothece-minutissima |
| Aphanothece nidulans | https://lpsn.dsmz.de/species/aphanothece-nidulans |
| Aphanothece nostocopsis | https://lpsn.dsmz.de/species/aphanothece-nostocopsis |
| Aphanothece parallela | https://lpsn.dsmz.de/species/aphanothece-parallela |
| Aphanothece paralleliformis | https://lpsn.dsmz.de/species/aphanothece-paralleliformis |
| Aphanothece sacrum | https://lpsn.dsmz.de/species/aphanothece-sacrum |
| Aphanothece shiloi | https://lpsn.dsmz.de/species/aphanothece-shiloi |
| Aphanothece smithii | https://lpsn.dsmz.de/species/aphanothece-smithii |
| Aphanothece stratus | https://lpsn.dsmz.de/species/aphanothece-stratus |
| Aphanothece subsalina | https://lpsn.dsmz.de/species/aphanothece-subsalina |
| Aphanothece sulphurica | https://lpsn.dsmz.de/species/aphanothece-sulphurica |
| Caedimonas varicaedens | https://lpsn.dsmz.de/species/caedimonas-varicaedens |
| Chroococcus aurantiacus | https://lpsn.dsmz.de/species/chroococcus-aurantiacus |
| Chroococcus constrictus | https://lpsn.dsmz.de/species/chroococcus-constrictus |
| Chroococcus distans | https://lpsn.dsmz.de/species/chroococcus-distans |
| Chroococcus endophyticus | https://lpsn.dsmz.de/species/chroococcus-endophyticus |
| Chroococcus ercegovicii | https://lpsn.dsmz.de/species/chroococcus-ercegovicii |
| Chroococcus fuscoviolaceus | https://lpsn.dsmz.de/species/chroococcus-fuscoviolaceus |
| Chroococcus gomontii | https://lpsn.dsmz.de/species/chroococcus-gomontii |

| | |
|---|---|
| Chroococcus indicus | https://lpsn.dsmz.de/species/chroococcus-indicus |
| Chroococcus major | https://lpsn.dsmz.de/species/chroococcus-major |
| Chroococcus mediocris | https://lpsn.dsmz.de/species/chroococcus-mediocris |
| Chroococcus microscopicus | https://lpsn.dsmz.de/species/chroococcus-microscopicus |
| Chroococcus minor | https://lpsn.dsmz.de/species/chroococcus-minor |
| Chroococcus minutissimus | https://lpsn.dsmz.de/species/chroococcus-minutissimus |
| Chroococcus minutus | https://lpsn.dsmz.de/species/chroococcus-minutus |
| Chroococcus muralis | https://lpsn.dsmz.de/species/chroococcus-muralis |
| Chroococcus niger | https://lpsn.dsmz.de/species/chroococcus-niger |
| Chroococcus rufescens | https://lpsn.dsmz.de/species/chroococcus-rufescens |
| Chroococcus tenacoides | https://lpsn.dsmz.de/species/chroococcus-tenacoides |
| Chroococcus turgidus | https://lpsn.dsmz.de/species/chroococcus-turgidus |
| Chroococcus yellowstonensis | https://lpsn.dsmz.de/species/chroococcus-yellowstonensis |
| Cyanobacterium stanieri | https://lpsn.dsmz.de/species/cyanobacterium-stanieri |
| Cyanobium amethystinum | https://lpsn.dsmz.de/species/cyanobium-amethystinum |
| Cyanobium bacillare | https://lpsn.dsmz.de/species/cyanobium-bacillare |
| Cyanobium diatomicola | https://lpsn.dsmz.de/species/cyanobium-diatomicola |
| Cyanobium eximium | https://lpsn.dsmz.de/species/cyanobium-eximium |
| Cyanobium gaarderi | https://lpsn.dsmz.de/species/cyanobium-gaarderi |
| Cyanobium gracile | https://lpsn.dsmz.de/species/cyanobium-gracile |
| Cyanobium parvum | https://lpsn.dsmz.de/species/cyanobium-parvum |
| Cyanobium plancticum | https://lpsn.dsmz.de/species/cyanobium-plancticum |
| Cyanobium roseum | https://lpsn.dsmz.de/species/cyanobium-roseum |
| Cyanobium rubescens | https://lpsn.dsmz.de/species/cyanobium-rubescens |
| Cyanobium virgarosea | https://lpsn.dsmz.de/species/cyanobium-virgarosea |
| Cyanothece aeruginosa | https://lpsn.dsmz.de/species/cyanothece-aeruginosa |
| Cyanothece crassiuscula | https://lpsn.dsmz.de/species/cyanothece-crassiuscula |
| Cyanothece shiloi | https://lpsn.dsmz.de/species/cyanothece-shiloi |
| Cyanothece svehlovae | https://lpsn.dsmz.de/species/cyanothece-svehlovae |
| Epigloeosphaera filamentosa | https://lpsn.dsmz.de/species/epigloeosphaera-filamentosa |
| Epigloeosphaera glebulenta | https://lpsn.dsmz.de/species/epigloeosphaera-glebulenta |
| Geminocystis herdmanii | https://lpsn.dsmz.de/species/geminocystis-herdmanii |
| Geminocystis papuanica | https://lpsn.dsmz.de/species/geminocystis-papuanica |
| Leptolyngbya aeruginea | https://lpsn.dsmz.de/species/leptolyngbya-aeruginea |
| Leptolyngbya africana | https://lpsn.dsmz.de/species/leptolyngbya-africana |
| Leptolyngbya andina | https://lpsn.dsmz.de/species/leptolyngbya-andina |
| Leptolyngbya angusta | https://lpsn.dsmz.de/species/leptolyngbya-angusta |
| Leptolyngbya angustissima | https://lpsn.dsmz.de/species/leptolyngbya-angustissima |
| Leptolyngbya aspera | https://lpsn.dsmz.de/species/leptolyngbya-aspera |
| Leptolyngbya batrachosperma | https://lpsn.dsmz.de/species/leptolyngbya-batrachosperma |
| Leptolyngbya benthonica | https://lpsn.dsmz.de/species/leptolyngbya-benthonica |
| Leptolyngbya bigranulata | https://lpsn.dsmz.de/species/leptolyngbya-bigranulata |
| Leptolyngbya bijahensis | https://lpsn.dsmz.de/species/leptolyngbya-bijahensis |
| Leptolyngbya blennophila | https://lpsn.dsmz.de/species/leptolyngbya-blennophila |

| | |
|---|---|
| Leptolyngbya borchgrevinkii | https://lpsn.dsmz.de/species/leptolyngbya-borchgrevinkii |
| Leptolyngbya boryana | https://lpsn.dsmz.de/species/leptolyngbya-boryana |
| Leptolyngbya breviarticulata | https://lpsn.dsmz.de/species/leptolyngbya-breviarticulata |
| Leptolyngbya calothrichoides | https://lpsn.dsmz.de/species/leptolyngbya-calothrichoides |
| Leptolyngbya carnea | https://lpsn.dsmz.de/species/leptolyngbya-carnea |
| Leptolyngbya cartilaginea | https://lpsn.dsmz.de/species/leptolyngbya-cartilaginea |
| Leptolyngbya cataractarum | https://lpsn.dsmz.de/species/leptolyngbya-cataractarum |
| Leptolyngbya cebennensis | https://lpsn.dsmz.de/species/leptolyngbya-cebennensis |
| Leptolyngbya chlorina | https://lpsn.dsmz.de/species/leptolyngbya-chlorina |
| Leptolyngbya compacta | https://lpsn.dsmz.de/species/leptolyngbya-compacta |
| Leptolyngbya copelandii | https://lpsn.dsmz.de/species/leptolyngbya-copelandii |
| Leptolyngbya corticola | https://lpsn.dsmz.de/species/leptolyngbya-corticola |
| Leptolyngbya crassior | https://lpsn.dsmz.de/species/leptolyngbya-crassior |
| Leptolyngbya crispata | https://lpsn.dsmz.de/species/leptolyngbya-crispata |
| Leptolyngbya dangeardii | https://lpsn.dsmz.de/species/leptolyngbya-dangeardii |
| Leptolyngbya delicatula | https://lpsn.dsmz.de/species/leptolyngbya-delicatula |
| Leptolyngbya desmidiacearum | https://lpsn.dsmz.de/species/leptolyngbya-desmidiacearum |
| Leptolyngbya distincta | https://lpsn.dsmz.de/species/leptolyngbya-distincta |
| Leptolyngbya ectocarpi | https://lpsn.dsmz.de/species/leptolyngbya-ectocarpi |
| Leptolyngbya eliskae | https://lpsn.dsmz.de/species/leptolyngbya-eliskae |
| Leptolyngbya elongata | https://lpsn.dsmz.de/species/leptolyngbya-elongata |
| Leptolyngbya endolithica | https://lpsn.dsmz.de/species/leptolyngbya-endolithica |
| Leptolyngbya ercegovicii | https://lpsn.dsmz.de/species/leptolyngbya-ercegovicii |
| Leptolyngbya erebi | https://lpsn.dsmz.de/species/leptolyngbya-erebi |
| Leptolyngbya fallax | https://lpsn.dsmz.de/species/leptolyngbya-fallax |
| Leptolyngbya ferruginea | https://lpsn.dsmz.de/species/leptolyngbya-ferruginea |
| Leptolyngbya foveolarum | https://lpsn.dsmz.de/species/leptolyngbya-foveolarum |
| Leptolyngbya fragilis | https://lpsn.dsmz.de/species/leptolyngbya-fragilis |
| Leptolyngbya fritschiana | https://lpsn.dsmz.de/species/leptolyngbya-fritschiana |
| Leptolyngbya fritschii | https://lpsn.dsmz.de/species/leptolyngbya-fritschii |
| Leptolyngbya gelatinicola | https://lpsn.dsmz.de/species/leptolyngbya-gelatinicola |
| Leptolyngbya gelatinosa | https://lpsn.dsmz.de/species/leptolyngbya-gelatinosa |
| Leptolyngbya glacialis | https://lpsn.dsmz.de/species/leptolyngbya-glacialis |
| Leptolyngbya gloeophila | https://lpsn.dsmz.de/species/leptolyngbya-gloeophila |
| Leptolyngbya gracilis | https://lpsn.dsmz.de/species/leptolyngbya-gracilis |
| Leptolyngbya gracillima | https://lpsn.dsmz.de/species/leptolyngbya-gracillima |
| Leptolyngbya granulifera | https://lpsn.dsmz.de/species/leptolyngbya-granulifera |
| Leptolyngbya hansgirgiana | https://lpsn.dsmz.de/species/leptolyngbya-hansgirgiana |
| Leptolyngbya henningsii | https://lpsn.dsmz.de/species/leptolyngbya-henningsii |
| Leptolyngbya homogenea | https://lpsn.dsmz.de/species/leptolyngbya-homogenea |
| Leptolyngbya hormoides | https://lpsn.dsmz.de/species/leptolyngbya-hormoides |
| Leptolyngbya hypolimnetica | https://lpsn.dsmz.de/species/leptolyngbya-hypolimnetica |
| Leptolyngbya jadertina | https://lpsn.dsmz.de/species/leptolyngbya-jadertina |
| Leptolyngbya komarovii | https://lpsn.dsmz.de/species/leptolyngbya-komarovii |

| | |
|---|---|
| Leptolyngbya lagerheimii | https://lpsn.dsmz.de/species/leptolyngbya-lagerheimii |
| Leptolyngbya laminosa | https://lpsn.dsmz.de/species/leptolyngbya-laminosa |
| Leptolyngbya lapidea | https://lpsn.dsmz.de/species/leptolyngbya-lapidea |
| Leptolyngbya lignicola | https://lpsn.dsmz.de/species/leptolyngbya-lignicola |
| Leptolyngbya maior | https://lpsn.dsmz.de/species/leptolyngbya-maior |
| Leptolyngbya margaretheana | https://lpsn.dsmz.de/species/leptolyngbya-margaretheana |
| Leptolyngbya margaritata | https://lpsn.dsmz.de/species/leptolyngbya-margaritata |
| Leptolyngbya marina | https://lpsn.dsmz.de/species/leptolyngbya-marina |
| Leptolyngbya membraniporae | https://lpsn.dsmz.de/species/leptolyngbya-membraniporae |
| Leptolyngbya minor | https://lpsn.dsmz.de/species/leptolyngbya-minor |
| Leptolyngbya minuta | https://lpsn.dsmz.de/species/leptolyngbya-minuta |
| Leptolyngbya minutissima | https://lpsn.dsmz.de/species/leptolyngbya-minutissima |
| Leptolyngbya mucicola | https://lpsn.dsmz.de/species/leptolyngbya-mucicola |
| Leptolyngbya mucosa | https://lpsn.dsmz.de/species/leptolyngbya-mucosa |
| Leptolyngbya muralis | https://lpsn.dsmz.de/species/leptolyngbya-muralis |
| Leptolyngbya mycoidea | https://lpsn.dsmz.de/species/leptolyngbya-mycoidea |
| Leptolyngbya nana | https://lpsn.dsmz.de/species/leptolyngbya-nana |
| Leptolyngbya nigrescens | https://lpsn.dsmz.de/species/leptolyngbya-nigrescens |
| Leptolyngbya norvegica | https://lpsn.dsmz.de/species/leptolyngbya-norvegica |
| Leptolyngbya nostochoides | https://lpsn.dsmz.de/species/leptolyngbya-nostochoides |
| Leptolyngbya nostocorum | https://lpsn.dsmz.de/species/leptolyngbya-nostocorum |
| Leptolyngbya notata | https://lpsn.dsmz.de/species/leptolyngbya-notata |
| Leptolyngbya ochracea | https://lpsn.dsmz.de/species/leptolyngbya-ochracea |
| Leptolyngbya olivacea | https://lpsn.dsmz.de/species/leptolyngbya-olivacea |
| Leptolyngbya orientalis | https://lpsn.dsmz.de/species/leptolyngbya-orientalis |
| Leptolyngbya patinae | https://lpsn.dsmz.de/species/leptolyngbya-patinae |
| Leptolyngbya perelegans | https://lpsn.dsmz.de/species/leptolyngbya-perelegans |
| Leptolyngbya perforans | https://lpsn.dsmz.de/species/leptolyngbya-perforans |
| Leptolyngbya phormidioides | https://lpsn.dsmz.de/species/leptolyngbya-phormidioides |
| Leptolyngbya polymorpha | https://lpsn.dsmz.de/species/leptolyngbya-polymorpha |
| Leptolyngbya polysiphoniae | https://lpsn.dsmz.de/species/leptolyngbya-polysiphoniae |
| Leptolyngbya protospira | https://lpsn.dsmz.de/species/leptolyngbya-protospira |
| Leptolyngbya pseudotenuis | https://lpsn.dsmz.de/species/leptolyngbya-pseudotenuis |
| Leptolyngbya pseudovalderiana | https://lpsn.dsmz.de/species/leptolyngbya-pseudovalderiana |
| Leptolyngbya purpurascens | https://lpsn.dsmz.de/species/leptolyngbya-purpurascens |
| Leptolyngbya purpurea | https://lpsn.dsmz.de/species/leptolyngbya-purpurea |
| Leptolyngbya ramosa | https://lpsn.dsmz.de/species/leptolyngbya-ramosa |
| Leptolyngbya rivulariarum | https://lpsn.dsmz.de/species/leptolyngbya-rivulariarum |
| Leptolyngbya rubra | https://lpsn.dsmz.de/species/leptolyngbya-rubra |
| Leptolyngbya saxicola | https://lpsn.dsmz.de/species/leptolyngbya-saxicola |
| Leptolyngbya schmidlei | https://lpsn.dsmz.de/species/leptolyngbya-schmidlei |
| Leptolyngbya scottii | https://lpsn.dsmz.de/species/leptolyngbya-scottii |
| Leptolyngbya scytonematicola | https://lpsn.dsmz.de/species/leptolyngbya-scytonematicola |
| Leptolyngbya spiralis | https://lpsn.dsmz.de/species/leptolyngbya-spiralis |

| | |
|---|---|
| Leptolyngbya subcapitata | https://lpsn.dsmz.de/species/leptolyngbya-subcapitata |
| Leptolyngbya subcyanea | https://lpsn.dsmz.de/species/leptolyngbya-subcyanea |
| Leptolyngbya subsalina | https://lpsn.dsmz.de/species/leptolyngbya-subsalina |
| Leptolyngbya subterranea | https://lpsn.dsmz.de/species/leptolyngbya-subterranea |
| Leptolyngbya subtilissima | https://lpsn.dsmz.de/species/leptolyngbya-subtilissima |
| Leptolyngbya subtruncata | https://lpsn.dsmz.de/species/leptolyngbya-subtruncata |
| Leptolyngbya subuliformis | https://lpsn.dsmz.de/species/leptolyngbya-subuliformis |
| Leptolyngbya tenerrima | https://lpsn.dsmz.de/species/leptolyngbya-tenerrima |
| Leptolyngbya tenuis | https://lpsn.dsmz.de/species/leptolyngbya-tenuis |
| Leptolyngbya tenuissima | https://lpsn.dsmz.de/species/leptolyngbya-tenuissima |
| Leptolyngbya thermalis | https://lpsn.dsmz.de/species/leptolyngbya-thermalis |
| Leptolyngbya thermarum | https://lpsn.dsmz.de/species/leptolyngbya-thermarum |
| Leptolyngbya thermobia | https://lpsn.dsmz.de/species/leptolyngbya-thermobia |
| Leptolyngbya thermophila | https://lpsn.dsmz.de/species/leptolyngbya-thermophila |
| Leptolyngbya treleasei | https://lpsn.dsmz.de/species/leptolyngbya-treleasei |
| Leptolyngbya truncata | https://lpsn.dsmz.de/species/leptolyngbya-truncata |
| Leptolyngbya valderiana | https://lpsn.dsmz.de/species/leptolyngbya-valderiana |
| Leptolyngbya vandenberghenii | https://lpsn.dsmz.de/species/leptolyngbya-vandenberghenii |
| Leptolyngbya vesiculosa | https://lpsn.dsmz.de/species/leptolyngbya-vesiculosa |
| Leptolyngbya vincentii | https://lpsn.dsmz.de/species/leptolyngbya-vincentii |
| Leptolyngbya voronichiniana | https://lpsn.dsmz.de/species/leptolyngbya-voronichiniana |
| Leptolyngbya weedii | https://lpsn.dsmz.de/species/leptolyngbya-weedii |
| Leptolyngbya weissii | https://lpsn.dsmz.de/species/leptolyngbya-weissii |
| Leptolyngbya woronichinii | https://lpsn.dsmz.de/species/leptolyngbya-woronichinii |
| Leptolyngbya yellowstonensis | https://lpsn.dsmz.de/species/leptolyngbya-yellowstonensis |
| Oscillatoria additicia | https://lpsn.dsmz.de/species/oscillatoria-additicia |
| Oscillatoria anguiformis | https://lpsn.dsmz.de/species/oscillatoria-anguiformis |
| Oscillatoria angusta | https://lpsn.dsmz.de/species/oscillatoria-angusta |
| Oscillatoria anomala | https://lpsn.dsmz.de/species/oscillatoria-anomala |
| Oscillatoria bonnemaisonii | https://lpsn.dsmz.de/species/oscillatoria-bonnemaisonii |
| Oscillatoria conformis | https://lpsn.dsmz.de/species/oscillatoria-conformis |
| Oscillatoria cortiana | https://lpsn.dsmz.de/species/oscillatoria-cortiana |
| Oscillatoria crassa | https://lpsn.dsmz.de/species/oscillatoria-crassa |
| Oscillatoria depauperata | https://lpsn.dsmz.de/species/oscillatoria-depauperata |
| Oscillatoria difficilis | https://lpsn.dsmz.de/species/oscillatoria-difficilis |
| Oscillatoria euboeica | https://lpsn.dsmz.de/species/oscillatoria-euboeica |
| Oscillatoria funiformis | https://lpsn.dsmz.de/species/oscillatoria-funiformis |
| Oscillatoria hameloides | https://lpsn.dsmz.de/species/oscillatoria-hameloides |
| Oscillatoria indica | https://lpsn.dsmz.de/species/oscillatoria-indica |
| Oscillatoria lachneri | https://lpsn.dsmz.de/species/oscillatoria-lachneri |
| Oscillatoria leavittiae | https://lpsn.dsmz.de/species/oscillatoria-leavittiae |
| Oscillatoria levis | https://lpsn.dsmz.de/species/oscillatoria-levis |
| Oscillatoria limosa | https://lpsn.dsmz.de/species/oscillatoria-limosa |
| Oscillatoria margaritifera | https://lpsn.dsmz.de/species/oscillatoria-margaritifera |

| | |
|---|---|
| Oscillatoria obscura | https://lpsn.dsmz.de/species/oscillatoria-obscura |
| Oscillatoria princeps | https://lpsn.dsmz.de/species/oscillatoria-princeps |
| Oscillatoria rhamphoidea | https://lpsn.dsmz.de/species/oscillatoria-rhamphoidea |
| Oscillatoria rosea | https://lpsn.dsmz.de/species/oscillatoria-rosea |
| Oscillatoria sancta | https://lpsn.dsmz.de/species/oscillatoria-sancta |
| Oscillatoria subbrevis | https://lpsn.dsmz.de/species/oscillatoria-subbrevis |
| Oscillatoria subcortiana | https://lpsn.dsmz.de/species/oscillatoria-subcortiana |
| Oscillatoria tenuis | https://lpsn.dsmz.de/species/oscillatoria-tenuis |
| Oscillatoria trichoides | https://lpsn.dsmz.de/species/oscillatoria-trichoides |
| Oscillatoria variabilis | https://lpsn.dsmz.de/species/oscillatoria-variabilis |
| Pseudanabaena acicularis | https://lpsn.dsmz.de/species/pseudanabaena-acicularis |
| Pseudanabaena amphigranulata | https://lpsn.dsmz.de/species/pseudanabaena-amphigranulata |
| Pseudanabaena apiculatoflexuosa | https://lpsn.dsmz.de/species/pseudanabaena-apiculatoflexuosa |
| Pseudanabaena arcuata | https://lpsn.dsmz.de/species/pseudanabaena-arcuata |
| Pseudanabaena batrachospermorum | https://lpsn.dsmz.de/species/pseudanabaena-batrachospermorum |
| Pseudanabaena belizensis | https://lpsn.dsmz.de/species/pseudanabaena-belizensis |
| Pseudanabaena catenata | https://lpsn.dsmz.de/species/pseudanabaena-catenata |
| Pseudanabaena cinerea | https://lpsn.dsmz.de/species/pseudanabaena-cinerea |
| Pseudanabaena dictyothalla | https://lpsn.dsmz.de/species/pseudanabaena-dictyothalla |
| Pseudanabaena endophytica | https://lpsn.dsmz.de/species/pseudanabaena-endophytica |
| Pseudanabaena foetida | https://lpsn.dsmz.de/species/pseudanabaena-foetida |
| Pseudanabaena galeata | https://lpsn.dsmz.de/species/pseudanabaena-galeata |
| Pseudanabaena limnetica | https://lpsn.dsmz.de/species/pseudanabaena-limnetica |
| Pseudanabaena minima | https://lpsn.dsmz.de/species/pseudanabaena-minima |
| Pseudanabaena moniliformis | https://lpsn.dsmz.de/species/pseudanabaena-moniliformis |
| Pseudanabaena persicina | https://lpsn.dsmz.de/species/pseudanabaena-persicina |
| Pseudanabaena raphidoides | https://lpsn.dsmz.de/species/pseudanabaena-raphidoides |
| Pseudanabaena rosea | https://lpsn.dsmz.de/species/pseudanabaena-rosea |
| Pseudanabaena spelaea | https://lpsn.dsmz.de/species/pseudanabaena-spelaea |
| Pseudanabaena starmachii | https://lpsn.dsmz.de/species/pseudanabaena-starmachii |
| Pseudanabaena thermalis | https://lpsn.dsmz.de/species/pseudanabaena-thermalis |
| Pseudanabaena ulula | https://lpsn.dsmz.de/species/pseudanabaena-ulula |
| Pseudanabaena voronichinii | https://lpsn.dsmz.de/species/pseudanabaena-voronichinii |
| Pseudanabaena westiana | https://lpsn.dsmz.de/species/pseudanabaena-westiana |
| Pseudanabaena yagii | https://lpsn.dsmz.de/species/pseudanabaena-yagii |
| Spirulina aeruginea | https://lpsn.dsmz.de/species/spirulina-aeruginea |
| Spirulina caldaria | https://lpsn.dsmz.de/species/spirulina-caldaria |
| Spirulina condensata | https://lpsn.dsmz.de/species/spirulina-condensata |
| Spirulina laxa | https://lpsn.dsmz.de/species/spirulina-laxa |
| Spirulina laxissima | https://lpsn.dsmz.de/species/spirulina-laxissima |
| Spirulina magnifica | https://lpsn.dsmz.de/species/spirulina-magnifica |
| Spirulina major | https://lpsn.dsmz.de/species/spirulina-major |
| Spirulina platensis | https://lpsn.dsmz.de/species/spirulina-platensis |
| Spirulina subsalsa | https://lpsn.dsmz.de/species/spirulina-subsalsa |

| | |
|---|---|
| Synechococcus aeruginosus | https://lpsn.dsmz.de/species/synechococcus-aeruginosus |
| Synechococcus amethystinus | https://lpsn.dsmz.de/species/synechococcus-amethystinus |
| Synechococcus arcuatus | https://lpsn.dsmz.de/species/synechococcus-arcuatus |
| Synechococcus bacillaris | https://lpsn.dsmz.de/species/synechococcus-bacillaris |
| Synechococcus bigranulatus | https://lpsn.dsmz.de/species/synechococcus-bigranulatus |
| Synechococcus cedrorum | https://lpsn.dsmz.de/species/synechococcus-cedrorum |
| Synechococcus diatomicola | https://lpsn.dsmz.de/species/synechococcus-diatomicola |
| Synechococcus elongatus | https://lpsn.dsmz.de/species/synechococcus-elongatus |
| Synechococcus eximius | https://lpsn.dsmz.de/species/synechococcus-eximius |
| Synechococcus gaarderi | https://lpsn.dsmz.de/species/synechococcus-gaarderi |
| Synechococcus leopoliensis | https://lpsn.dsmz.de/species/synechococcus-leopoliensis |
| Synechococcus lividus | https://lpsn.dsmz.de/species/synechococcus-lividus |
| Synechococcus marinus | https://lpsn.dsmz.de/species/synechococcus-marinus |
| Synechococcus moorigangaii | https://lpsn.dsmz.de/species/synechococcus-moorigangaii |
| Synechococcus mundulus | https://lpsn.dsmz.de/species/synechococcus-mundulus |
| Synechococcus nidulans | https://lpsn.dsmz.de/species/synechococcus-nidulans |
| Synechococcus plancticus | https://lpsn.dsmz.de/species/synechococcus-plancticus |
| Synechococcus rhodobaktron | https://lpsn.dsmz.de/species/synechococcus-rhodobaktron |
| Synechococcus roseus | https://lpsn.dsmz.de/species/synechococcus-roseus |
| Synechococcus rubescens | https://lpsn.dsmz.de/species/synechococcus-rubescens |
| Synechococcus subsalsus | https://lpsn.dsmz.de/species/synechococcus-subsalsus |
| Synechococcus sulphuricus | https://lpsn.dsmz.de/species/synechococcus-sulphuricus |
| Synechococcus vescus | https://lpsn.dsmz.de/species/synechococcus-vescus |
| Synechococcus viridissimus | https://lpsn.dsmz.de/species/synechococcus-viridissimus |
| Synechococcus vulcanus | https://lpsn.dsmz.de/species/synechococcus-vulcanus |
| Thermostichus amphigranulatus | https://lpsn.dsmz.de/species/thermostichus-amphigranulatus |
| Thermostichus arcuatus | https://lpsn.dsmz.de/species/thermostichus-arcuatus |
| Thermostichus bigranulatus | https://lpsn.dsmz.de/species/thermostichus-bigranulatus |
| Thermostichus lividus | https://lpsn.dsmz.de/species/thermostichus-lividus |
| Thermostichus praelongus | https://lpsn.dsmz.de/species/thermostichus-praelongus |
| Thermostichus sublividus | https://lpsn.dsmz.de/species/thermostichus-sublividus |
| Thermostichus sulphuricus | https://lpsn.dsmz.de/species/thermostichus-sulphuricus |
| Thermostichus vescus | https://lpsn.dsmz.de/species/thermostichus-vescus |
| Thermostichus viridissimus | https://lpsn.dsmz.de/species/thermostichus-viridissimus |
| Thermostichus vulcanus | https://lpsn.dsmz.de/species/thermostichus-vulcanus |
| Abditibacterium | https://lpsn.dsmz.de/genus/abditibacterium |
| Abiotrophia | https://lpsn.dsmz.de/genus/abiotrophia |
| Absicoccus | https://lpsn.dsmz.de/genus/absicoccus |
| Absiella | https://lpsn.dsmz.de/genus/absiella |
| Abyssalbus | https://lpsn.dsmz.de/genus/abyssalbus |
| Abyssibacter | https://lpsn.dsmz.de/genus/abyssibacter |
| Abyssicoccus | https://lpsn.dsmz.de/genus/abyssicoccus |
| Abyssivirga | https://lpsn.dsmz.de/genus/abyssivirga |
| Acanthopleuribacter | https://lpsn.dsmz.de/genus/acanthopleuribacter |

| | |
|---|---|
| Acaricomes | https://lpsn.dsmz.de/genus/acaricomes |
| Acerihabitans | https://lpsn.dsmz.de/genus/acerihabitans |
| Acetanaerobacterium | https://lpsn.dsmz.de/genus/acetanaerobacterium |
| Acetatifactor | https://lpsn.dsmz.de/genus/acetatifactor |
| Acetilactobacillus | https://lpsn.dsmz.de/genus/acetilactobacillus |
| Acetitomaculum | https://lpsn.dsmz.de/genus/acetitomaculum |
| Acetivibrio | https://lpsn.dsmz.de/genus/acetivibrio |
| Acetoanaerobium | https://lpsn.dsmz.de/genus/acetoanaerobium |
| Acetobacter | https://lpsn.dsmz.de/genus/acetobacter |
| Acetobacterium | https://lpsn.dsmz.de/genus/acetobacterium |
| Acetobacteroides | https://lpsn.dsmz.de/genus/acetobacteroides |
| Acetofilamentum | https://lpsn.dsmz.de/genus/acetofilamentum |
| Acetogenium | https://lpsn.dsmz.de/genus/acetogenium |
| Acetohalobium | https://lpsn.dsmz.de/genus/acetohalobium |
| Acetomicrobium | https://lpsn.dsmz.de/genus/acetomicrobium |
| Acetonema | https://lpsn.dsmz.de/genus/acetonema |
| Acetothermus | https://lpsn.dsmz.de/genus/acetothermus |
| Acholeplasma | https://lpsn.dsmz.de/genus/acholeplasma |
| Achromatium | https://lpsn.dsmz.de/genus/achromatium |
| Achromobacter | https://lpsn.dsmz.de/genus/achromobacter |
| Acidaminobacter | https://lpsn.dsmz.de/genus/acidaminobacter |
| Acidaminococcus | https://lpsn.dsmz.de/genus/acidaminococcus |
| Acidianus | https://lpsn.dsmz.de/genus/acidianus |
| Acidibacter | https://lpsn.dsmz.de/genus/acidibacter |
| Acidibrevibacterium | https://lpsn.dsmz.de/genus/acidibrevibacterium |
| Acidicaldus | https://lpsn.dsmz.de/genus/acidicaldus |
| Acidicapsa | https://lpsn.dsmz.de/genus/acidicapsa |
| Acidiferrimicrobium | https://lpsn.dsmz.de/genus/acidiferrimicrobium |
| Acidiferrobacter | https://lpsn.dsmz.de/genus/acidiferrobacter |
| Acidihalobacter | https://lpsn.dsmz.de/genus/acidihalobacter |
| Acidilobus | https://lpsn.dsmz.de/genus/acidilobus |
| Acidiluteibacter | https://lpsn.dsmz.de/genus/acidiluteibacter |
| Acidimangrovimonas | https://lpsn.dsmz.de/genus/acidimangrovimonas |
| Acidimicrobium | https://lpsn.dsmz.de/genus/acidimicrobium |
| Acidiphilium | https://lpsn.dsmz.de/genus/acidiphilium |
| Acidipila | https://lpsn.dsmz.de/genus/acidipila |
| Acidiplasma | https://lpsn.dsmz.de/genus/acidiplasma |
| Acidipropionibacterium | https://lpsn.dsmz.de/genus/acidipropionibacterium |
| Acidisarcina | https://lpsn.dsmz.de/genus/acidisarcina |
| Acidisoma | https://lpsn.dsmz.de/genus/acidisoma |
| Acidisphaera | https://lpsn.dsmz.de/genus/acidisphaera |
| Aciditerrimonas | https://lpsn.dsmz.de/genus/aciditerrimonas |
| Acidithiobacillus | https://lpsn.dsmz.de/genus/acidithiobacillus |
| Acidithrix | https://lpsn.dsmz.de/genus/acidithrix |

| | |
|---|---|
| Acidobacterium | https://lpsn.dsmz.de/genus/acidobacterium |
| Acidocella | https://lpsn.dsmz.de/genus/acidocella |
| Acidomonas | https://lpsn.dsmz.de/genus/acidomonas |
| Acidothermus | https://lpsn.dsmz.de/genus/acidothermus |
| Acidovorax | https://lpsn.dsmz.de/genus/acidovorax |
| Acinetobacter | https://lpsn.dsmz.de/genus/acinetobacter |
| Acrocarpospora | https://lpsn.dsmz.de/genus/acrocarpospora |
| Actibacter | https://lpsn.dsmz.de/genus/actibacter |
| Actibacterium | https://lpsn.dsmz.de/genus/actibacterium |
| Actimicrobium | https://lpsn.dsmz.de/genus/actimicrobium |
| Actinacidiphila | https://lpsn.dsmz.de/genus/actinacidiphila |
| Actinaurispora | https://lpsn.dsmz.de/genus/actinaurispora |
| Actinoallomurus | https://lpsn.dsmz.de/genus/actinoallomurus |
| Actinoalloteichus | https://lpsn.dsmz.de/genus/actinoalloteichus |
| Actinobacillus | https://lpsn.dsmz.de/genus/actinobacillus |
| Actinobaculum | https://lpsn.dsmz.de/genus/actinobaculum |
| Actinobispora | https://lpsn.dsmz.de/genus/actinobispora |
| Actinocatenispora | https://lpsn.dsmz.de/genus/actinocatenispora |
| Actinocorallia | https://lpsn.dsmz.de/genus/actinocorallia |
| Actinocrinis | https://lpsn.dsmz.de/genus/actinocrinis |
| Actinocrispum | https://lpsn.dsmz.de/genus/actinocrispum |
| Actinokineospora | https://lpsn.dsmz.de/genus/actinokineospora |
| Actinomadura | https://lpsn.dsmz.de/genus/actinomadura |
| Actinomarinicola | https://lpsn.dsmz.de/genus/actinomarinicola |
| Actinomyces | https://lpsn.dsmz.de/genus/actinomyces |
| Actinomycetospora | https://lpsn.dsmz.de/genus/actinomycetospora |
| Actinophytocola | https://lpsn.dsmz.de/genus/actinophytocola |
| Actinoplanes | https://lpsn.dsmz.de/genus/actinoplanes |
| Actinopolymorpha | https://lpsn.dsmz.de/genus/actinopolymorpha |
| Actinopolyspora | https://lpsn.dsmz.de/genus/actinopolyspora |
| Actinopycnidium | https://lpsn.dsmz.de/genus/actinopycnidium |
| Actinorectispora | https://lpsn.dsmz.de/genus/actinorectispora |
| Actinorhabdospora | https://lpsn.dsmz.de/genus/actinorhabdospora |
| Actinorugispora | https://lpsn.dsmz.de/genus/actinorugispora |
| Actinospica | https://lpsn.dsmz.de/genus/actinospica |
| Actinosporangium | https://lpsn.dsmz.de/genus/actinosporangium |
| Actinosynnema | https://lpsn.dsmz.de/genus/actinosynnema |
| Actinotalea | https://lpsn.dsmz.de/genus/actinotalea |
| Actinotignum | https://lpsn.dsmz.de/genus/actinotignum |
| Actirhodobacter | https://lpsn.dsmz.de/genus/actirhodobacter |
| Acutalibacter | https://lpsn.dsmz.de/genus/acutalibacter |
| Acuticoccus | https://lpsn.dsmz.de/genus/acuticoccus |
| Adhaeretor | https://lpsn.dsmz.de/genus/adhaeretor |
| Adhaeribacter | https://lpsn.dsmz.de/genus/adhaeribacter |

| | |
|---|---|
| Adlercreutzia | https://lpsn.dsmz.de/genus/adlercreutzia |
| Advenella | https://lpsn.dsmz.de/genus/advenella |
| Aedoeadaptatus | https://lpsn.dsmz.de/genus/aedoeadaptatus |
| Aegicerativicinus | https://lpsn.dsmz.de/genus/aegicerativicinus |
| Aegyptianella | https://lpsn.dsmz.de/genus/aegyptianella |
| Aeoliella | https://lpsn.dsmz.de/genus/aeoliella |
| Aequoribacter | https://lpsn.dsmz.de/genus/aequoribacter |
| Aequorivita | https://lpsn.dsmz.de/genus/aequorivita |
| Aeribacillus | https://lpsn.dsmz.de/genus/aeribacillus |
| Aeriscardovia | https://lpsn.dsmz.de/genus/aeriscardovia |
| Aerococcus | https://lpsn.dsmz.de/genus/aerococcus |
| Aeromicrobium | https://lpsn.dsmz.de/genus/aeromicrobium |
| Aeromonas | https://lpsn.dsmz.de/genus/aeromonas |
| Aerophototrophica | https://lpsn.dsmz.de/genus/aerophototrophica |
| Aeropyrum | https://lpsn.dsmz.de/genus/aeropyrum |
| Aerosticca | https://lpsn.dsmz.de/genus/aerosticca |
| Aestuariibacter | https://lpsn.dsmz.de/genus/aestuariibacter |
| Aestuariibaculum | https://lpsn.dsmz.de/genus/aestuariibaculum |
| Aestuariibius | https://lpsn.dsmz.de/genus/aestuariibius |
| Aestuariicella | https://lpsn.dsmz.de/genus/aestuariicella |
| Aestuariicoccus | https://lpsn.dsmz.de/genus/aestuariicoccus |
| Aestuariicola | https://lpsn.dsmz.de/genus/aestuariicola |
| Aestuariihabitans | https://lpsn.dsmz.de/genus/aestuariihabitans |
| Aestuariimicrobium | https://lpsn.dsmz.de/genus/aestuariimicrobium |
| Aestuariimonas | https://lpsn.dsmz.de/genus/aestuariimonas |
| Aestuariirhabdus | https://lpsn.dsmz.de/genus/aestuariirhabdus |
| Aestuariisphingobium | https://lpsn.dsmz.de/genus/aestuariisphingobium |
| Aestuariispira | https://lpsn.dsmz.de/genus/aestuariispira |
| Aestuariivirga | https://lpsn.dsmz.de/genus/aestuariivirga |
| Aestuariivita | https://lpsn.dsmz.de/genus/aestuariivita |
| Aestuariivivens | https://lpsn.dsmz.de/genus/aestuariivivens |
| Aestuarium | https://lpsn.dsmz.de/genus/aestuarium |
| Aetherobacter | https://lpsn.dsmz.de/genus/aetherobacter |
| Afifella | https://lpsn.dsmz.de/genus/afifella |
| Afipia | https://lpsn.dsmz.de/genus/afipia |
| Agaribacter | https://lpsn.dsmz.de/genus/agaribacter |
| Agaribacterium | https://lpsn.dsmz.de/genus/agaribacterium |
| Agaricicola | https://lpsn.dsmz.de/genus/agaricicola |
| Agarilytica | https://lpsn.dsmz.de/genus/agarilytica |
| Agarivorans | https://lpsn.dsmz.de/genus/agarivorans |
| Agathobacter | https://lpsn.dsmz.de/genus/agathobacter |
| Agathobaculum | https://lpsn.dsmz.de/genus/agathobaculum |
| Aggregatibacter | https://lpsn.dsmz.de/genus/aggregatibacter |
| Aggregatilinea | https://lpsn.dsmz.de/genus/aggregatilinea |

| | |
|---|---|
| Aggregicoccus | https://lpsn.dsmz.de/genus/aggregicoccus |
| Agilicoccus | https://lpsn.dsmz.de/genus/agilicoccus |
| Agitococcus | https://lpsn.dsmz.de/genus/agitococcus |
| Agreia | https://lpsn.dsmz.de/genus/agreia |
| Agrilactobacillus | https://lpsn.dsmz.de/genus/agrilactobacillus |
| Agriterribacter | https://lpsn.dsmz.de/genus/agriterribacter |
| Agrobacterium | https://lpsn.dsmz.de/genus/agrobacterium |
| Agrococcus | https://lpsn.dsmz.de/genus/agrococcus |
| Agromonas | https://lpsn.dsmz.de/genus/agromonas |
| Agromyces | https://lpsn.dsmz.de/genus/agromyces |
| Ahniella | https://lpsn.dsmz.de/genus/ahniella |
| Ahrensia | https://lpsn.dsmz.de/genus/ahrensia |
| Aidingibacillus | https://lpsn.dsmz.de/genus/aidingibacillus |
| Aidingimonas | https://lpsn.dsmz.de/genus/aidingimonas |
| Akkermansia | https://lpsn.dsmz.de/genus/akkermansia |
| Albibacillus | https://lpsn.dsmz.de/genus/albibacillus |
| Albibacter | https://lpsn.dsmz.de/genus/albibacter |
| Albibacterium | https://lpsn.dsmz.de/genus/albibacterium |
| Albidiferax | https://lpsn.dsmz.de/genus/albidiferax |
| Albidoferax | https://lpsn.dsmz.de/genus/albidoferax |
| Albidovulum | https://lpsn.dsmz.de/genus/albidovulum |
| Albimonas | https://lpsn.dsmz.de/genus/albimonas |
| Albirhodobacter | https://lpsn.dsmz.de/genus/albirhodobacter |
| Alcaligenes | https://lpsn.dsmz.de/genus/alcaligenes |
| Alcalilimnicola | https://lpsn.dsmz.de/genus/alcalilimnicola |
| Alcanivorax | https://lpsn.dsmz.de/genus/alcanivorax |
| Aldersonia | https://lpsn.dsmz.de/genus/aldersonia |
| Algibacillus | https://lpsn.dsmz.de/genus/algibacillus |
| Algibacter | https://lpsn.dsmz.de/genus/algibacter |
| Algicoccus | https://lpsn.dsmz.de/genus/algicoccus |
| Algicola | https://lpsn.dsmz.de/genus/algicola |
| Algihabitans | https://lpsn.dsmz.de/genus/algihabitans |
| Algimonas | https://lpsn.dsmz.de/genus/algimonas |
| Alginatibacterium | https://lpsn.dsmz.de/genus/alginatibacterium |
| Algiphilus | https://lpsn.dsmz.de/genus/algiphilus |
| Algisphaera | https://lpsn.dsmz.de/genus/algisphaera |
| Algitalea | https://lpsn.dsmz.de/genus/algitalea |
| Algoriella | https://lpsn.dsmz.de/genus/algoriella |
| Algoriphagus | https://lpsn.dsmz.de/genus/algoriphagus |
| Aliagarivorans | https://lpsn.dsmz.de/genus/aliagarivorans |
| Aliarcobacter | https://lpsn.dsmz.de/genus/aliarcobacter |
| Alicycliphilus | https://lpsn.dsmz.de/genus/alicycliphilus |
| Alicyclobacillus | https://lpsn.dsmz.de/genus/alicyclobacillus |
| Alienimonas | https://lpsn.dsmz.de/genus/alienimonas |

| | |
|---|---|
| Aliibacillus | https://lpsn.dsmz.de/genus/aliibacillus |
| Aliicoccus | https://lpsn.dsmz.de/genus/aliicoccus |
| Aliidiomarina | https://lpsn.dsmz.de/genus/aliidiomarina |
| Aliidongia | https://lpsn.dsmz.de/genus/aliidongia |
| Aliifodinibius | https://lpsn.dsmz.de/genus/aliifodinibius |
| Aliiglaciecola | https://lpsn.dsmz.de/genus/aliiglaciecola |
| Aliikangiella | https://lpsn.dsmz.de/genus/aliikangiella |
| Aliiroseovarius | https://lpsn.dsmz.de/genus/aliiroseovarius |
| Aliiruegeria | https://lpsn.dsmz.de/genus/aliiruegeria |
| Aliishimia | https://lpsn.dsmz.de/genus/aliishimia |
| Aliivibrio | https://lpsn.dsmz.de/genus/aliivibrio |
| Alishewanella | https://lpsn.dsmz.de/genus/alishewanella |
| Alistipes | https://lpsn.dsmz.de/genus/alistipes |
| Alitiscatomonas | https://lpsn.dsmz.de/genus/alitiscatomonas |
| Alkalibacillus | https://lpsn.dsmz.de/genus/alkalibacillus |
| Alkalibacter | https://lpsn.dsmz.de/genus/alkalibacter |
| Alkalibacterium | https://lpsn.dsmz.de/genus/alkalibacterium |
| Alkalibaculum | https://lpsn.dsmz.de/genus/alkalibaculum |
| Alkalicaulis | https://lpsn.dsmz.de/genus/alkalicaulis |
| Alkalicella | https://lpsn.dsmz.de/genus/alkalicella |
| Alkalicoccobacillus | https://lpsn.dsmz.de/genus/alkalicoccobacillus |
| Alkalicoccus | https://lpsn.dsmz.de/genus/alkalicoccus |
| Alkaliflexus | https://lpsn.dsmz.de/genus/alkaliflexus |
| Alkalihalobacillus | https://lpsn.dsmz.de/genus/alkalihalobacillus |
| Alkalihalobacterium | https://lpsn.dsmz.de/genus/alkalihalobacterium |
| Alkalihalophilus | https://lpsn.dsmz.de/genus/alkalihalophilus |
| Alkalilactibacillus | https://lpsn.dsmz.de/genus/alkalilactibacillus |
| Alkalilacustris | https://lpsn.dsmz.de/genus/alkalilacustris |
| Alkalilimnicola | https://lpsn.dsmz.de/genus/alkalilimnicola |
| Alkalimarinus | https://lpsn.dsmz.de/genus/alkalimarinus |
| Alkalimicrobium | https://lpsn.dsmz.de/genus/alkalimicrobium |
| Alkalimonas | https://lpsn.dsmz.de/genus/alkalimonas |
| Alkaliphilus | https://lpsn.dsmz.de/genus/alkaliphilus |
| Alkalispirochaeta | https://lpsn.dsmz.de/genus/alkalispirochaeta |
| Alkalitalea | https://lpsn.dsmz.de/genus/alkalitalea |
| Alkanibacter | https://lpsn.dsmz.de/genus/alkanibacter |
| Alkanindiges | https://lpsn.dsmz.de/genus/alkanindiges |
| Allgaiera | https://lpsn.dsmz.de/genus/allgaiera |
| Allisonella | https://lpsn.dsmz.de/genus/allisonella |
| Alloacidobacterium | https://lpsn.dsmz.de/genus/alloacidobacterium |
| Alloactinosynnema | https://lpsn.dsmz.de/genus/alloactinosynnema |
| Allobacillus | https://lpsn.dsmz.de/genus/allobacillus |
| Allobaculum | https://lpsn.dsmz.de/genus/allobaculum |
| Allobranchiibius | https://lpsn.dsmz.de/genus/allobranchiibius |

| | |
|---|---|
| Allocatelliglobosispora | https://lpsn.dsmz.de/genus/allocatelliglobosispora |
| Allochromatium | https://lpsn.dsmz.de/genus/allochromatium |
| Allofrancisella | https://lpsn.dsmz.de/genus/allofrancisella |
| Allofustis | https://lpsn.dsmz.de/genus/allofustis |
| Allohahella | https://lpsn.dsmz.de/genus/allohahella |
| Allohumibacter | https://lpsn.dsmz.de/genus/allohumibacter |
| Alloiococcus | https://lpsn.dsmz.de/genus/alloiococcus |
| Allokutzneria | https://lpsn.dsmz.de/genus/allokutzneria |
| Allomonas | https://lpsn.dsmz.de/genus/allomonas |
| Allonocardiopsis | https://lpsn.dsmz.de/genus/allonocardiopsis |
| Allopontixanthobacter | https://lpsn.dsmz.de/genus/allopontixanthobacter |
| Alloprevotella | https://lpsn.dsmz.de/genus/alloprevotella |
| Allopseudarcicella | https://lpsn.dsmz.de/genus/allopseudarcicella |
| Allorhizobium | https://lpsn.dsmz.de/genus/allorhizobium |
| Allorhizocola | https://lpsn.dsmz.de/genus/allorhizocola |
| Allosaccharopolyspora | https://lpsn.dsmz.de/genus/allosaccharopolyspora |
| Allosalinactinospora | https://lpsn.dsmz.de/genus/allosalinactinospora |
| Alloscardovia | https://lpsn.dsmz.de/genus/alloscardovia |
| Allosediminivita | https://lpsn.dsmz.de/genus/allosediminivita |
| Allosphingosinicella | https://lpsn.dsmz.de/genus/allosphingosinicella |
| Allostreptomyces | https://lpsn.dsmz.de/genus/allostreptomyces |
| Alpinimonas | https://lpsn.dsmz.de/genus/alpinimonas |
| Alsobacter | https://lpsn.dsmz.de/genus/alsobacter |
| Alteracholeplasma | https://lpsn.dsmz.de/genus/alteracholeplasma |
| Alteraurantiacibacter | https://lpsn.dsmz.de/genus/alteraurantiacibacter |
| Altererythrobacter | https://lpsn.dsmz.de/genus/altererythrobacter |
| Alteribacillus | https://lpsn.dsmz.de/genus/alteribacillus |
| Alteribacter | https://lpsn.dsmz.de/genus/alteribacter |
| Altericroceibacterium | https://lpsn.dsmz.de/genus/altericroceibacterium |
| Alterinioella | https://lpsn.dsmz.de/genus/alterinioella |
| Alteripontixanthobacter | https://lpsn.dsmz.de/genus/alteripontixanthobacter |
| Alteriqipengyuania | https://lpsn.dsmz.de/genus/alteriqipengyuania |
| Alterococcus | https://lpsn.dsmz.de/genus/alterococcus |
| Alteromonas | https://lpsn.dsmz.de/genus/alteromonas |
| Alysiella | https://lpsn.dsmz.de/genus/alysiella |
| Amantichitinum | https://lpsn.dsmz.de/genus/amantichitinum |
| Amaricoccus | https://lpsn.dsmz.de/genus/amaricoccus |
| Amedibacillus | https://lpsn.dsmz.de/genus/amedibacillus |
| Amedibacterium | https://lpsn.dsmz.de/genus/amedibacterium |
| Ameyamaea | https://lpsn.dsmz.de/genus/ameyamaea |
| Aminicella | https://lpsn.dsmz.de/genus/aminicella |
| Aminiphilus | https://lpsn.dsmz.de/genus/aminiphilus |
| Aminipila | https://lpsn.dsmz.de/genus/aminipila |
| Aminivibrio | https://lpsn.dsmz.de/genus/aminivibrio |

| | |
|---|---|
| Aminobacter | https://lpsn.dsmz.de/genus/aminobacter |
| Aminobacterium | https://lpsn.dsmz.de/genus/aminobacterium |
| Aminocella | https://lpsn.dsmz.de/genus/aminocella |
| Aminomonas | https://lpsn.dsmz.de/genus/aminomonas |
| Ammonifex | https://lpsn.dsmz.de/genus/ammonifex |
| Ammoniibacillus | https://lpsn.dsmz.de/genus/ammoniibacillus |
| Ammoniphilus | https://lpsn.dsmz.de/genus/ammoniphilus |
| Amnibacterium | https://lpsn.dsmz.de/genus/amnibacterium |
| Amniculibacterium | https://lpsn.dsmz.de/genus/amniculibacterium |
| Amnimonas | https://lpsn.dsmz.de/genus/amnimonas |
| Amoebobacter | https://lpsn.dsmz.de/genus/amoebobacter |
| Amorphosporangium | https://lpsn.dsmz.de/genus/amorphosporangium |
| Amorphus | https://lpsn.dsmz.de/genus/amorphus |
| Amphibacillus | https://lpsn.dsmz.de/genus/amphibacillus |
| Amphiplicatus | https://lpsn.dsmz.de/genus/amphiplicatus |
| Amphritea | https://lpsn.dsmz.de/genus/amphritea |
| Ampullariella | https://lpsn.dsmz.de/genus/ampullariella |
| Ampullimonas | https://lpsn.dsmz.de/genus/ampullimonas |
| Amycolata | https://lpsn.dsmz.de/genus/amycolata |
| Amycolatopsis | https://lpsn.dsmz.de/genus/amycolatopsis |
| Amycolicicoccus | https://lpsn.dsmz.de/genus/amycolicicoccus |
| Amylibacter | https://lpsn.dsmz.de/genus/amylibacter |
| Amylolactobacillus | https://lpsn.dsmz.de/genus/amylolactobacillus |
| Anaeroarcus | https://lpsn.dsmz.de/genus/anaeroarcus |
| Anaerobacillus | https://lpsn.dsmz.de/genus/anaerobacillus |
| Anaerobacter | https://lpsn.dsmz.de/genus/anaerobacter |
| Anaerobacterium | https://lpsn.dsmz.de/genus/anaerobacterium |
| Anaerobaculum | https://lpsn.dsmz.de/genus/anaerobaculum |
| Anaerobiospirillum | https://lpsn.dsmz.de/genus/anaerobiospirillum |
| Anaerobium | https://lpsn.dsmz.de/genus/anaerobium |
| Anaerobranca | https://lpsn.dsmz.de/genus/anaerobranca |
| Anaerobutyricum | https://lpsn.dsmz.de/genus/anaerobutyricum |
| Anaerocella | https://lpsn.dsmz.de/genus/anaerocella |
| Anaerococcus | https://lpsn.dsmz.de/genus/anaerococcus |
| Anaerocolumna | https://lpsn.dsmz.de/genus/anaerocolumna |
| Anaerofilum | https://lpsn.dsmz.de/genus/anaerofilum |
| Anaerofustis | https://lpsn.dsmz.de/genus/anaerofustis |
| Anaeroglobus | https://lpsn.dsmz.de/genus/anaeroglobus |
| Anaerohalosphaera | https://lpsn.dsmz.de/genus/anaerohalosphaera |
| Anaerolinea | https://lpsn.dsmz.de/genus/anaerolinea |
| Anaeromicrobium | https://lpsn.dsmz.de/genus/anaeromicrobium |
| Anaeromonas | https://lpsn.dsmz.de/genus/anaeromonas |
| Anaeromusa | https://lpsn.dsmz.de/genus/anaeromusa |
| Anaeromyxobacter | https://lpsn.dsmz.de/genus/anaeromyxobacter |

| | |
|---|---|
| Anaeropeptidivorans | https://lpsn.dsmz.de/genus/anaeropeptidivorans |
| Anaerophaga | https://lpsn.dsmz.de/genus/anaerophaga |
| Anaeroplasma | https://lpsn.dsmz.de/genus/anaeroplasma |
| Anaerorhabdus | https://lpsn.dsmz.de/genus/anaerorhabdus |
| Anaerosacchariphilus | https://lpsn.dsmz.de/genus/anaerosacchariphilus |
| Anaerosalibacter | https://lpsn.dsmz.de/genus/anaerosalibacter |
| Anaerosinus | https://lpsn.dsmz.de/genus/anaerosinus |
| Anaerosolibacter | https://lpsn.dsmz.de/genus/anaerosolibacter |
| Anaerosphaera | https://lpsn.dsmz.de/genus/anaerosphaera |
| Anaerospora | https://lpsn.dsmz.de/genus/anaerospora |
| Anaerosporobacter | https://lpsn.dsmz.de/genus/anaerosporobacter |
| Anaerosporomusa | https://lpsn.dsmz.de/genus/anaerosporomusa |
| Anaerostipes | https://lpsn.dsmz.de/genus/anaerostipes |
| Anaerotaenia | https://lpsn.dsmz.de/genus/anaerotaenia |
| Anaerotalea | https://lpsn.dsmz.de/genus/anaerotalea |
| Anaerotignum | https://lpsn.dsmz.de/genus/anaerotignum |
| Anaerotruncus | https://lpsn.dsmz.de/genus/anaerotruncus |
| Anaerovibrio | https://lpsn.dsmz.de/genus/anaerovibrio |
| Anaerovirgula | https://lpsn.dsmz.de/genus/anaerovirgula |
| Anaerovorax | https://lpsn.dsmz.de/genus/anaerovorax |
| Anaplasma | https://lpsn.dsmz.de/genus/anaplasma |
| Anathece | https://lpsn.dsmz.de/genus/anathece |
| Anatilimnocola | https://lpsn.dsmz.de/genus/anatilimnocola |
| Ancalochloris | https://lpsn.dsmz.de/genus/ancalochloris |
| Ancalomicrobium | https://lpsn.dsmz.de/genus/ancalomicrobium |
| Ancylobacter | https://lpsn.dsmz.de/genus/ancylobacter |
| Ancylomarina | https://lpsn.dsmz.de/genus/ancylomarina |
| Anderseniella | https://lpsn.dsmz.de/genus/anderseniella |
| Andreesenia | https://lpsn.dsmz.de/genus/andreesenia |
| Andreprevotia | https://lpsn.dsmz.de/genus/andreprevotia |
| Aneurinibacillus | https://lpsn.dsmz.de/genus/aneurinibacillus |
| Angiococcus | https://lpsn.dsmz.de/genus/angiococcus |
| Angulomicrobium | https://lpsn.dsmz.de/genus/angulomicrobium |
| Angustibacter | https://lpsn.dsmz.de/genus/angustibacter |
| Annwoodia | https://lpsn.dsmz.de/genus/annwoodia |
| Anoxybacillus | https://lpsn.dsmz.de/genus/anoxybacillus |
| Anoxybacter | https://lpsn.dsmz.de/genus/anoxybacter |
| Anoxynatronum | https://lpsn.dsmz.de/genus/anoxynatronum |
| Anseongella | https://lpsn.dsmz.de/genus/anseongella |
| Antarcticibacterium | https://lpsn.dsmz.de/genus/antarcticibacterium |
| Antarcticimicrobium | https://lpsn.dsmz.de/genus/antarcticimicrobium |
| Antarcticimonas | https://lpsn.dsmz.de/genus/antarcticimonas |
| Antarctobacter | https://lpsn.dsmz.de/genus/antarctobacter |
| Anthropogastromicrobium | https://lpsn.dsmz.de/genus/anthropogastromicrobium |

| | |
|---|---|
| Antribacter | https://lpsn.dsmz.de/genus/antribacter |
| Antricoccus | https://lpsn.dsmz.de/genus/antricoccus |
| Antrihabitans | https://lpsn.dsmz.de/genus/antrihabitans |
| Aphanothece | https://lpsn.dsmz.de/genus/aphanothece |
| Apibacter | https://lpsn.dsmz.de/genus/apibacter |
| Apilactobacillus | https://lpsn.dsmz.de/genus/apilactobacillus |
| Aquabacter | https://lpsn.dsmz.de/genus/aquabacter |
| Aquabacterium | https://lpsn.dsmz.de/genus/aquabacterium |
| Aquamicrobium | https://lpsn.dsmz.de/genus/aquamicrobium |
| Aquariibacter | https://lpsn.dsmz.de/genus/aquariibacter |
| Aquaspirillum | https://lpsn.dsmz.de/genus/aquaspirillum |
| Aquaticitalea | https://lpsn.dsmz.de/genus/aquaticitalea |
| Aquella | https://lpsn.dsmz.de/genus/aquella |
| Aquibacillus | https://lpsn.dsmz.de/genus/aquibacillus |
| Aquibacter | https://lpsn.dsmz.de/genus/aquibacter |
| Aquibium | https://lpsn.dsmz.de/genus/aquibium |
| Aquicella | https://lpsn.dsmz.de/genus/aquicella |
| Aquicoccus | https://lpsn.dsmz.de/genus/aquicoccus |
| Aquidulcibacter | https://lpsn.dsmz.de/genus/aquidulcibacter |
| Aquifex | https://lpsn.dsmz.de/genus/aquifex |
| Aquiflexum | https://lpsn.dsmz.de/genus/aquiflexum |
| Aquihabitans | https://lpsn.dsmz.de/genus/aquihabitans |
| Aquiluna | https://lpsn.dsmz.de/genus/aquiluna |
| Aquimarina | https://lpsn.dsmz.de/genus/aquimarina |
| Aquimixticola | https://lpsn.dsmz.de/genus/aquimixticola |
| Aquimonas | https://lpsn.dsmz.de/genus/aquimonas |
| Aquincola | https://lpsn.dsmz.de/genus/aquincola |
| Aquipluma | https://lpsn.dsmz.de/genus/aquipluma |
| Aquipuribacter | https://lpsn.dsmz.de/genus/aquipuribacter |
| Aquirhabdus | https://lpsn.dsmz.de/genus/aquirhabdus |
| Aquirufa | https://lpsn.dsmz.de/genus/aquirufa |
| Aquisalibacillus | https://lpsn.dsmz.de/genus/aquisalibacillus |
| Aquisalimonas | https://lpsn.dsmz.de/genus/aquisalimonas |
| Aquisalinus | https://lpsn.dsmz.de/genus/aquisalinus |
| Aquisediminimonas | https://lpsn.dsmz.de/genus/aquisediminimonas |
| Aquisphaera | https://lpsn.dsmz.de/genus/aquisphaera |
| Aquitalea | https://lpsn.dsmz.de/genus/aquitalea |
| Arachidicoccus | https://lpsn.dsmz.de/genus/arachidicoccus |
| Arachnia | https://lpsn.dsmz.de/genus/arachnia |
| Arboricoccus | https://lpsn.dsmz.de/genus/arboricoccus |
| Arboriscoccus | https://lpsn.dsmz.de/genus/arboriscoccus |
| Arcanobacterium | https://lpsn.dsmz.de/genus/arcanobacterium |
| Archaeoglobus | https://lpsn.dsmz.de/genus/archaeoglobus |
| Archangium | https://lpsn.dsmz.de/genus/archangium |

| | |
|---|---|
| Arcicella | https://lpsn.dsmz.de/genus/arcicella |
| Arcobacter | https://lpsn.dsmz.de/genus/arcobacter |
| Arcticibacter | https://lpsn.dsmz.de/genus/arcticibacter |
| Arcticibacterium | https://lpsn.dsmz.de/genus/arcticibacterium |
| Arcticiflavibacter | https://lpsn.dsmz.de/genus/arcticiflavibacter |
| Ardenticatena | https://lpsn.dsmz.de/genus/ardenticatena |
| Arenibacillus | https://lpsn.dsmz.de/genus/arenibacillus |
| Arenibacter | https://lpsn.dsmz.de/genus/arenibacter |
| Arenibacterium | https://lpsn.dsmz.de/genus/arenibacterium |
| Arenibaculum | https://lpsn.dsmz.de/genus/arenibaculum |
| Arenicella | https://lpsn.dsmz.de/genus/arenicella |
| Arenimicrobium | https://lpsn.dsmz.de/genus/arenimicrobium |
| Arenimonas | https://lpsn.dsmz.de/genus/arenimonas |
| Arenitalea | https://lpsn.dsmz.de/genus/arenitalea |
| Arenivirga | https://lpsn.dsmz.de/genus/arenivirga |
| Arhodomonas | https://lpsn.dsmz.de/genus/arhodomonas |
| Aridibacter | https://lpsn.dsmz.de/genus/aridibacter |
| Armatimonas | https://lpsn.dsmz.de/genus/armatimonas |
| Aromatoleum | https://lpsn.dsmz.de/genus/aromatoleum |
| Arsenicibacter | https://lpsn.dsmz.de/genus/arsenicibacter |
| Arsenicicoccus | https://lpsn.dsmz.de/genus/arsenicicoccus |
| Arsenicitalea | https://lpsn.dsmz.de/genus/arsenicitalea |
| Arsenophonus | https://lpsn.dsmz.de/genus/arsenophonus |
| Arsukibacterium | https://lpsn.dsmz.de/genus/arsukibacterium |
| Arthrobacter | https://lpsn.dsmz.de/genus/arthrobacter |
| Arthrospiribacter | https://lpsn.dsmz.de/genus/arthrospiribacter |
| Arundinibacter | https://lpsn.dsmz.de/genus/arundinibacter |
| Arvibacter | https://lpsn.dsmz.de/genus/arvibacter |
| Asaccharobacter | https://lpsn.dsmz.de/genus/asaccharobacter |
| Asaccharospora | https://lpsn.dsmz.de/genus/asaccharospora |
| Asaia | https://lpsn.dsmz.de/genus/asaia |
| Asanoa | https://lpsn.dsmz.de/genus/asanoa |
| Ascidiaceihabitans | https://lpsn.dsmz.de/genus/ascidiaceihabitans |
| Ascidiimonas | https://lpsn.dsmz.de/genus/ascidiimonas |
| Asinibacterium | https://lpsn.dsmz.de/genus/asinibacterium |
| Asprobacter | https://lpsn.dsmz.de/genus/asprobacter |
| Aspromonas | https://lpsn.dsmz.de/genus/aspromonas |
| Asteroleplasma | https://lpsn.dsmz.de/genus/asteroleplasma |
| Asticcacaulis | https://lpsn.dsmz.de/genus/asticcacaulis |
| Athalassotoga | https://lpsn.dsmz.de/genus/athalassotoga |
| Atopobacter | https://lpsn.dsmz.de/genus/atopobacter |
| Atopobium | https://lpsn.dsmz.de/genus/atopobium |
| Atopococcus | https://lpsn.dsmz.de/genus/atopococcus |
| Atopomonas | https://lpsn.dsmz.de/genus/atopomonas |

| | |
|---|---|
| Atopostipes | https://lpsn.dsmz.de/genus/atopostipes |
| Atribacter | https://lpsn.dsmz.de/genus/atribacter |
| Aurantiacibacter | https://lpsn.dsmz.de/genus/aurantiacibacter |
| Aurantiacicella | https://lpsn.dsmz.de/genus/aurantiacicella |
| Aurantibacter | https://lpsn.dsmz.de/genus/aurantibacter |
| Aurantimicrobium | https://lpsn.dsmz.de/genus/aurantimicrobium |
| Aurantimonas | https://lpsn.dsmz.de/genus/aurantimonas |
| Aurantisolimonas | https://lpsn.dsmz.de/genus/aurantisolimonas |
| Aurantivirga | https://lpsn.dsmz.de/genus/aurantivirga |
| Auraticoccus | https://lpsn.dsmz.de/genus/auraticoccus |
| Aureibacillus | https://lpsn.dsmz.de/genus/aureibacillus |
| Aureibacter | https://lpsn.dsmz.de/genus/aureibacter |
| Aureibaculum | https://lpsn.dsmz.de/genus/aureibaculum |
| Aureicoccus | https://lpsn.dsmz.de/genus/aureicoccus |
| Aureimonas | https://lpsn.dsmz.de/genus/aureimonas |
| Aureisphaera | https://lpsn.dsmz.de/genus/aureisphaera |
| Aureispira | https://lpsn.dsmz.de/genus/aureispira |
| Aureitalea | https://lpsn.dsmz.de/genus/aureitalea |
| Aureivirga | https://lpsn.dsmz.de/genus/aureivirga |
| Aureliella | https://lpsn.dsmz.de/genus/aureliella |
| Aureobacterium | https://lpsn.dsmz.de/genus/aureobacterium |
| Auricoccus | https://lpsn.dsmz.de/genus/auricoccus |
| Auritidibacter | https://lpsn.dsmz.de/genus/auritidibacter |
| Austwickia | https://lpsn.dsmz.de/genus/austwickia |
| Avibacterium | https://lpsn.dsmz.de/genus/avibacterium |
| Azoarcus | https://lpsn.dsmz.de/genus/azoarcus |
| Azohydromonas | https://lpsn.dsmz.de/genus/azohydromonas |
| Azomonas | https://lpsn.dsmz.de/genus/azomonas |
| Azomonotrichon | https://lpsn.dsmz.de/genus/azomonotrichon |
| Azonexus | https://lpsn.dsmz.de/genus/azonexus |
| Azorhizobium | https://lpsn.dsmz.de/genus/azorhizobium |
| Azorhizophilus | https://lpsn.dsmz.de/genus/azorhizophilus |
| Azospira | https://lpsn.dsmz.de/genus/azospira |
| Azospirillum | https://lpsn.dsmz.de/genus/azospirillum |
| Azotobacter | https://lpsn.dsmz.de/genus/azotobacter |
| Azovibrio | https://lpsn.dsmz.de/genus/azovibrio |
| Bacillus | https://lpsn.dsmz.de/genus/bacillus |
| Bacteriolyticum | https://lpsn.dsmz.de/genus/bacteriolyticum |
| Bacterionema | https://lpsn.dsmz.de/genus/bacterionema |
| Bacterioplanes | https://lpsn.dsmz.de/genus/bacterioplanes |
| Bacterioplanoides | https://lpsn.dsmz.de/genus/bacterioplanoides |
| Bacteriovorax | https://lpsn.dsmz.de/genus/bacteriovorax |
| Bacteroides | https://lpsn.dsmz.de/genus/bacteroides |
| Bactoderma | https://lpsn.dsmz.de/genus/bactoderma |

| | |
|---|---|
| Baekduia | https://lpsn.dsmz.de/genus/baekduia |
| Baia | https://lpsn.dsmz.de/genus/baia |
| Baileyella | https://lpsn.dsmz.de/genus/baileyella |
| Bailinhaonella | https://lpsn.dsmz.de/genus/bailinhaonella |
| Balnearium | https://lpsn.dsmz.de/genus/balnearium |
| Balneatrix | https://lpsn.dsmz.de/genus/balneatrix |
| Balneicella | https://lpsn.dsmz.de/genus/balneicella |
| Balneimonas | https://lpsn.dsmz.de/genus/balneimonas |
| Balneola | https://lpsn.dsmz.de/genus/balneola |
| Balneomonas | https://lpsn.dsmz.de/genus/balneomonas |
| Barnesiella | https://lpsn.dsmz.de/genus/barnesiella |
| Barrientosiimonas | https://lpsn.dsmz.de/genus/barrientosiimonas |
| Bartonella | https://lpsn.dsmz.de/genus/bartonella |
| Basfia | https://lpsn.dsmz.de/genus/basfia |
| Basilea | https://lpsn.dsmz.de/genus/basilea |
| Bauldia | https://lpsn.dsmz.de/genus/bauldia |
| Bavariicoccus | https://lpsn.dsmz.de/genus/bavariicoccus |
| Bdellovibrio | https://lpsn.dsmz.de/genus/bdellovibrio |
| Beggiatoa | https://lpsn.dsmz.de/genus/beggiatoa |
| Beijerinckia | https://lpsn.dsmz.de/genus/beijerinckia |
| Belliella | https://lpsn.dsmz.de/genus/belliella |
| Bellilinea | https://lpsn.dsmz.de/genus/bellilinea |
| Belnapia | https://lpsn.dsmz.de/genus/belnapia |
| Beneckea | https://lpsn.dsmz.de/genus/beneckea |
| Bergeriella | https://lpsn.dsmz.de/genus/bergeriella |
| Bergeyella | https://lpsn.dsmz.de/genus/bergeyella |
| Bermanella | https://lpsn.dsmz.de/genus/bermanella |
| Bernardetia | https://lpsn.dsmz.de/genus/bernardetia |
| Berryella | https://lpsn.dsmz.de/genus/berryella |
| Beutenbergia | https://lpsn.dsmz.de/genus/beutenbergia |
| Bhargavaea | https://lpsn.dsmz.de/genus/bhargavaea |
| Bibersteinia | https://lpsn.dsmz.de/genus/bibersteinia |
| Bieblia | https://lpsn.dsmz.de/genus/bieblia |
| Bifidobacterium | https://lpsn.dsmz.de/genus/bifidobacterium |
| Bilifractor | https://lpsn.dsmz.de/genus/bilifractor |
| Bilophila | https://lpsn.dsmz.de/genus/bilophila |
| Biomaibacter | https://lpsn.dsmz.de/genus/biomaibacter |
| Biostraticola | https://lpsn.dsmz.de/genus/biostraticola |
| Bisgaardia | https://lpsn.dsmz.de/genus/bisgaardia |
| Bizionia | https://lpsn.dsmz.de/genus/bizionia |
| Blastobacter | https://lpsn.dsmz.de/genus/blastobacter |
| Blastocatella | https://lpsn.dsmz.de/genus/blastocatella |
| Blastochloris | https://lpsn.dsmz.de/genus/blastochloris |
| Blastococcus | https://lpsn.dsmz.de/genus/blastococcus |

| | |
|---|---|
| Blastomonas | https://lpsn.dsmz.de/genus/blastomonas |
| Blastopirellula | https://lpsn.dsmz.de/genus/blastopirellula |
| Blattabacterium | https://lpsn.dsmz.de/genus/blattabacterium |
| Blautia | https://lpsn.dsmz.de/genus/blautia |
| Bogoriella | https://lpsn.dsmz.de/genus/bogoriella |
| Bombella | https://lpsn.dsmz.de/genus/bombella |
| Bombilactobacillus | https://lpsn.dsmz.de/genus/bombilactobacillus |
| Bombiscardovia | https://lpsn.dsmz.de/genus/bombiscardovia |
| Bordetella | https://lpsn.dsmz.de/genus/bordetella |
| Borrelia | https://lpsn.dsmz.de/genus/borrelia |
| Borreliella | https://lpsn.dsmz.de/genus/borreliella |
| Bosea | https://lpsn.dsmz.de/genus/bosea |
| Boseongicola | https://lpsn.dsmz.de/genus/boseongicola |
| Botrimarina | https://lpsn.dsmz.de/genus/botrimarina |
| Botryobacter | https://lpsn.dsmz.de/genus/botryobacter |
| Boudabousia | https://lpsn.dsmz.de/genus/boudabousia |
| Bounagaea | https://lpsn.dsmz.de/genus/bounagaea |
| Bowdenia | https://lpsn.dsmz.de/genus/bowdenia |
| Bowdeniella | https://lpsn.dsmz.de/genus/bowdeniella |
| Bowmanella | https://lpsn.dsmz.de/genus/bowmanella |
| Brachybacterium | https://lpsn.dsmz.de/genus/brachybacterium |
| Brachymonas | https://lpsn.dsmz.de/genus/brachymonas |
| Brachyspira | https://lpsn.dsmz.de/genus/brachyspira |
| Brackiella | https://lpsn.dsmz.de/genus/brackiella |
| Bradymonas | https://lpsn.dsmz.de/genus/bradymonas |
| Bradyrhizobium | https://lpsn.dsmz.de/genus/bradyrhizobium |
| Branchiibius | https://lpsn.dsmz.de/genus/branchiibius |
| Branhamella | https://lpsn.dsmz.de/genus/branhamella |
| Brassicibacter | https://lpsn.dsmz.de/genus/brassicibacter |
| Bremerella | https://lpsn.dsmz.de/genus/bremerella |
| Brenneria | https://lpsn.dsmz.de/genus/brenneria |
| Breoghania | https://lpsn.dsmz.de/genus/breoghania |
| Brevibacillus | https://lpsn.dsmz.de/genus/brevibacillus |
| Brevibacterium | https://lpsn.dsmz.de/genus/brevibacterium |
| Brevifollis | https://lpsn.dsmz.de/genus/brevifollis |
| Brevilactibacter | https://lpsn.dsmz.de/genus/brevilactibacter |
| Brevinema | https://lpsn.dsmz.de/genus/brevinema |
| Brevirhabdus | https://lpsn.dsmz.de/genus/brevirhabdus |
| Brevitalea | https://lpsn.dsmz.de/genus/brevitalea |
| Brevundimonas | https://lpsn.dsmz.de/genus/brevundimonas |
| Breznakia | https://lpsn.dsmz.de/genus/breznakia |
| Breznakibacter | https://lpsn.dsmz.de/genus/breznakibacter |
| Breznakiella | https://lpsn.dsmz.de/genus/breznakiella |
| Brochothrix | https://lpsn.dsmz.de/genus/brochothrix |

| | |
|---|---|
| Brockia | https://lpsn.dsmz.de/genus/brockia |
| Brooklawnia | https://lpsn.dsmz.de/genus/brooklawnia |
| Brotolimicola | https://lpsn.dsmz.de/genus/brotolimicola |
| Brotonthovivens | https://lpsn.dsmz.de/genus/brotonthovivens |
| Brucella | https://lpsn.dsmz.de/genus/brucella |
| Brumimicrobium | https://lpsn.dsmz.de/genus/brumimicrobium |
| Bryantella | https://lpsn.dsmz.de/genus/bryantella |
| Bryobacter | https://lpsn.dsmz.de/genus/bryobacter |
| Bryocella | https://lpsn.dsmz.de/genus/bryocella |
| Buchananella | https://lpsn.dsmz.de/genus/buchananella |
| Buchnera | https://lpsn.dsmz.de/genus/buchnera |
| Budvicia | https://lpsn.dsmz.de/genus/budvicia-1 |
| Budvicia | https://lpsn.dsmz.de/genus/budvicia |
| Bulleidia | https://lpsn.dsmz.de/genus/bulleidia |
| Bullifex | https://lpsn.dsmz.de/genus/bullifex |
| Burkholderia | https://lpsn.dsmz.de/genus/burkholderia |
| Buttiauxella | https://lpsn.dsmz.de/genus/buttiauxella |
| Butyratibacter | https://lpsn.dsmz.de/genus/butyratibacter |
| Butyribacter | https://lpsn.dsmz.de/genus/butyribacter |
| Butyricicoccus | https://lpsn.dsmz.de/genus/butyricicoccus |
| Butyricimonas | https://lpsn.dsmz.de/genus/butyricimonas |
| Butyrivibrio | https://lpsn.dsmz.de/genus/butyrivibrio |
| Byssovorax | https://lpsn.dsmz.de/genus/byssovorax |
| Bythopirellula | https://lpsn.dsmz.de/genus/bythopirellula |
| Caballeronia | https://lpsn.dsmz.de/genus/caballeronia |
| Caecibacterium | https://lpsn.dsmz.de/genus/caecibacterium |
| Caedibacter | https://lpsn.dsmz.de/genus/caedibacter |
| Caedimonas | https://lpsn.dsmz.de/genus/caedimonas |
| Caenibacillus | https://lpsn.dsmz.de/genus/caenibacillus |
| Caenibacterium | https://lpsn.dsmz.de/genus/caenibacterium |
| Caenibius | https://lpsn.dsmz.de/genus/caenibius |
| Caenimicrobium | https://lpsn.dsmz.de/genus/caenimicrobium |
| Caenimonas | https://lpsn.dsmz.de/genus/caenimonas |
| Caenispirillum | https://lpsn.dsmz.de/genus/caenispirillum |
| Calculibacillus | https://lpsn.dsmz.de/genus/calculibacillus |
| Caldalkalibacillus | https://lpsn.dsmz.de/genus/caldalkalibacillus |
| Caldanaerobacter | https://lpsn.dsmz.de/genus/caldanaerobacter |
| Caldanaerobius | https://lpsn.dsmz.de/genus/caldanaerobius |
| Caldanaerovirga | https://lpsn.dsmz.de/genus/caldanaerovirga |
| Calderihabitans | https://lpsn.dsmz.de/genus/calderihabitans |
| Calderobacterium | https://lpsn.dsmz.de/genus/calderobacterium |
| Caldibacillus | https://lpsn.dsmz.de/genus/caldibacillus |
| Caldicellulosiruptor | https://lpsn.dsmz.de/genus/caldicellulosiruptor |
| Caldichromatium | https://lpsn.dsmz.de/genus/caldichromatium |

| | |
|---|---|
| Caldicoprobacter | https://lpsn.dsmz.de/genus/caldicoprobacter |
| Caldilinea | https://lpsn.dsmz.de/genus/caldilinea |
| Caldimicrobium | https://lpsn.dsmz.de/genus/caldimicrobium |
| Caldimonas | https://lpsn.dsmz.de/genus/caldimonas |
| Caldisalinibacter | https://lpsn.dsmz.de/genus/caldisalinibacter |
| Caldisericum | https://lpsn.dsmz.de/genus/caldisericum |
| Caldisphaera | https://lpsn.dsmz.de/genus/caldisphaera |
| Calditerricola | https://lpsn.dsmz.de/genus/calditerricola |
| Calditerrivibrio | https://lpsn.dsmz.de/genus/calditerrivibrio |
| Caldithrix | https://lpsn.dsmz.de/genus/caldithrix |
| Caldivirga | https://lpsn.dsmz.de/genus/caldivirga |
| Caldovatus | https://lpsn.dsmz.de/genus/caldovatus |
| Calidifontibacillus | https://lpsn.dsmz.de/genus/calidifontibacillus |
| Calidifontibacter | https://lpsn.dsmz.de/genus/calidifontibacter |
| Calidifontimicrobium | https://lpsn.dsmz.de/genus/calidifontimicrobium |
| Calidithermus | https://lpsn.dsmz.de/genus/calidithermus |
| Caloramator | https://lpsn.dsmz.de/genus/caloramator |
| Caloranaerobacter | https://lpsn.dsmz.de/genus/caloranaerobacter |
| Caloribacterium | https://lpsn.dsmz.de/genus/caloribacterium |
| Calorimonas | https://lpsn.dsmz.de/genus/calorimonas |
| Calorithrix | https://lpsn.dsmz.de/genus/calorithrix |
| Calycomorphotria | https://lpsn.dsmz.de/genus/calycomorphotria |
| Calymmatobacterium | https://lpsn.dsmz.de/genus/calymmatobacterium |
| Camelimonas | https://lpsn.dsmz.de/genus/camelimonas |
| Camelliibacillus | https://lpsn.dsmz.de/genus/camelliibacillus |
| Caminibacter | https://lpsn.dsmz.de/genus/caminibacter |
| Caminicella | https://lpsn.dsmz.de/genus/caminicella |
| Campylobacter | https://lpsn.dsmz.de/genus/campylobacter |
| Candidimonas | https://lpsn.dsmz.de/genus/candidimonas |
| Canibacter | https://lpsn.dsmz.de/genus/canibacter |
| Canicola | https://lpsn.dsmz.de/genus/canicola |
| Capillibacterium | https://lpsn.dsmz.de/genus/capillibacterium |
| Capillimicrobium | https://lpsn.dsmz.de/genus/capillimicrobium |
| Capnocytophaga | https://lpsn.dsmz.de/genus/capnocytophaga |
| Caproicibacter | https://lpsn.dsmz.de/genus/caproicibacter |
| Caproicibacterium | https://lpsn.dsmz.de/genus/caproicibacterium |
| Caproiciproducens | https://lpsn.dsmz.de/genus/caproiciproducens |
| Capsularis | https://lpsn.dsmz.de/genus/capsularis |
| Capsulimonas | https://lpsn.dsmz.de/genus/capsulimonas |
| Carbonactinospora | https://lpsn.dsmz.de/genus/carbonactinospora |
| Carbophilus | https://lpsn.dsmz.de/genus/carbophilus |
| Carboxydibrachium | https://lpsn.dsmz.de/genus/carboxydibrachium |
| Carboxydobrachium | https://lpsn.dsmz.de/genus/carboxydobrachium |
| Carboxydocella | https://lpsn.dsmz.de/genus/carboxydocella |

| | |
|---|---|
| Carboxydothermus | https://lpsn.dsmz.de/genus/carboxydothermus |
| Carboxylicivirga | https://lpsn.dsmz.de/genus/carboxylicivirga |
| Cardiobacterium | https://lpsn.dsmz.de/genus/cardiobacterium |
| Carideicomes | https://lpsn.dsmz.de/genus/carideicomes |
| Carnimonas | https://lpsn.dsmz.de/genus/carnimonas |
| Carnobacterium | https://lpsn.dsmz.de/genus/carnobacterium |
| Caryophanon | https://lpsn.dsmz.de/genus/caryophanon |
| Caseobacter | https://lpsn.dsmz.de/genus/caseobacter |
| Casimicrobium | https://lpsn.dsmz.de/genus/casimicrobium |
| Castellaniella | https://lpsn.dsmz.de/genus/castellaniella |
| Catabacter | https://lpsn.dsmz.de/genus/catabacter |
| Catalinimonas | https://lpsn.dsmz.de/genus/catalinimonas |
| Catellatospora | https://lpsn.dsmz.de/genus/catellatospora |
| Catellibacterium | https://lpsn.dsmz.de/genus/catellibacterium |
| Catellicoccus | https://lpsn.dsmz.de/genus/catellicoccus |
| Catelliglobosispora | https://lpsn.dsmz.de/genus/catelliglobosispora |
| Catenibacillus | https://lpsn.dsmz.de/genus/catenibacillus |
| Catenibacterium | https://lpsn.dsmz.de/genus/catenibacterium |
| Catenisphaera | https://lpsn.dsmz.de/genus/catenisphaera |
| Catenococcus | https://lpsn.dsmz.de/genus/catenococcus |
| Catenovulum | https://lpsn.dsmz.de/genus/catenovulum |
| Catenulispora | https://lpsn.dsmz.de/genus/catenulispora |
| Catenuloplanes | https://lpsn.dsmz.de/genus/catenuloplanes |
| Catonella | https://lpsn.dsmz.de/genus/catonella |
| Caulifigura | https://lpsn.dsmz.de/genus/caulifigura |
| Caulobacter | https://lpsn.dsmz.de/genus/caulobacter |
| Cavicella | https://lpsn.dsmz.de/genus/cavicella |
| Caviibacter | https://lpsn.dsmz.de/genus/caviibacter |
| Caviibacterium | https://lpsn.dsmz.de/genus/caviibacterium |
| Cecembia | https://lpsn.dsmz.de/genus/cecembia |
| Cedecea | https://lpsn.dsmz.de/genus/cedecea |
| Celeribacter | https://lpsn.dsmz.de/genus/celeribacter |
| Celerinatantimonas | https://lpsn.dsmz.de/genus/celerinatantimonas |
| Cellulomonas | https://lpsn.dsmz.de/genus/cellulomonas |
| Cellulophaga | https://lpsn.dsmz.de/genus/cellulophaga |
| Cellulosibacter | https://lpsn.dsmz.de/genus/cellulosibacter |
| Cellulosilyticum | https://lpsn.dsmz.de/genus/cellulosilyticum |
| Cellulosimicrobium | https://lpsn.dsmz.de/genus/cellulosimicrobium |
| Cellvibrio | https://lpsn.dsmz.de/genus/cellvibrio |
| Centipeda | https://lpsn.dsmz.de/genus/centipeda |
| Cerasibacillus | https://lpsn.dsmz.de/genus/cerasibacillus |
| Cerasicoccus | https://lpsn.dsmz.de/genus/cerasicoccus |
| Cereibacter | https://lpsn.dsmz.de/genus/cereibacter |
| Cesiribacter | https://lpsn.dsmz.de/genus/cesiribacter |

| | |
|---|---|
| Cetia | https://lpsn.dsmz.de/genus/cetia |
| Cetobacterium | https://lpsn.dsmz.de/genus/cetobacterium |
| Chachezhania | https://lpsn.dsmz.de/genus/chachezhania |
| Chainia | https://lpsn.dsmz.de/genus/chainia |
| Chakrabartia | https://lpsn.dsmz.de/genus/chakrabartia |
| Changchengzhania | https://lpsn.dsmz.de/genus/changchengzhania |
| Chania | https://lpsn.dsmz.de/genus/chania |
| Chelativorans | https://lpsn.dsmz.de/genus/chelativorans |
| Chelatobacter | https://lpsn.dsmz.de/genus/chelatobacter |
| Chelatococcus | https://lpsn.dsmz.de/genus/chelatococcus |
| Chelonobacter | https://lpsn.dsmz.de/genus/chelonobacter |
| Chenggangzhangella | https://lpsn.dsmz.de/genus/chenggangzhangella |
| Chengkuizengella | https://lpsn.dsmz.de/genus/chengkuizengella |
| Chiayiivirga | https://lpsn.dsmz.de/genus/chiayiivirga |
| Chimaereicella | https://lpsn.dsmz.de/genus/chimaereicella |
| Chimaeribacter | https://lpsn.dsmz.de/genus/chimaeribacter |
| Chioneia | https://lpsn.dsmz.de/genus/chioneia |
| Chishuiella | https://lpsn.dsmz.de/genus/chishuiella |
| Chitinasiproducens | https://lpsn.dsmz.de/genus/chitinasiproducens |
| Chitinibacter | https://lpsn.dsmz.de/genus/chitinibacter |
| Chitinilyticum | https://lpsn.dsmz.de/genus/chitinilyticum |
| Chitinimonas | https://lpsn.dsmz.de/genus/chitinimonas |
| Chitiniphilus | https://lpsn.dsmz.de/genus/chitiniphilus |
| Chitinispirillum | https://lpsn.dsmz.de/genus/chitinispirillum |
| Chitinivibrio | https://lpsn.dsmz.de/genus/chitinivibrio |
| Chitinivorax | https://lpsn.dsmz.de/genus/chitinivorax |
| Chitinophaga | https://lpsn.dsmz.de/genus/chitinophaga |
| Chlamydia | https://lpsn.dsmz.de/genus/chlamydia |
| Chlamydophila | https://lpsn.dsmz.de/genus/chlamydophila |
| Chlorobaculum | https://lpsn.dsmz.de/genus/chlorobaculum |
| Chlorobium | https://lpsn.dsmz.de/genus/chlorobium |
| Chloroflexus | https://lpsn.dsmz.de/genus/chloroflexus |
| Chloroherpeton | https://lpsn.dsmz.de/genus/chloroherpeton |
| Chloronema | https://lpsn.dsmz.de/genus/chloronema |
| Chondromyces | https://lpsn.dsmz.de/genus/chondromyces |
| Christensenella | https://lpsn.dsmz.de/genus/christensenella |
| Chromatium | https://lpsn.dsmz.de/genus/chromatium |
| Chromatocurvus | https://lpsn.dsmz.de/genus/chromatocurvus |
| Chromobacterium | https://lpsn.dsmz.de/genus/chromobacterium |
| Chromocurvus | https://lpsn.dsmz.de/genus/chromocurvus |
| Chromohalobacter | https://lpsn.dsmz.de/genus/chromohalobacter |
| Chroococcus | https://lpsn.dsmz.de/genus/chroococcus |
| Chryseobacterium | https://lpsn.dsmz.de/genus/chryseobacterium |
| Chryseoglobus | https://lpsn.dsmz.de/genus/chryseoglobus |

| | |
|---|---|
| Chryseolinea | https://lpsn.dsmz.de/genus/chryseolinea |
| Chryseomicrobium | https://lpsn.dsmz.de/genus/chryseomicrobium |
| Chryseomonas | https://lpsn.dsmz.de/genus/chryseomonas |
| Chryseotalea | https://lpsn.dsmz.de/genus/chryseotalea |
| Chrysiogenes | https://lpsn.dsmz.de/genus/chrysiogenes |
| Chthonobacter | https://lpsn.dsmz.de/genus/chthonobacter |
| Chthonomonas | https://lpsn.dsmz.de/genus/chthonomonas |
| Chujaibacter | https://lpsn.dsmz.de/genus/chujaibacter |
| Chungangia | https://lpsn.dsmz.de/genus/chungangia |
| Ciceribacter | https://lpsn.dsmz.de/genus/ciceribacter |
| Citreicella | https://lpsn.dsmz.de/genus/citreicella |
| Citreicoccus | https://lpsn.dsmz.de/genus/citreicoccus |
| Citreimonas | https://lpsn.dsmz.de/genus/citreimonas |
| Citreitalea | https://lpsn.dsmz.de/genus/citreitalea |
| Citricoccus | https://lpsn.dsmz.de/genus/citricoccus |
| Citrifermentans | https://lpsn.dsmz.de/genus/citrifermentans |
| Citrobacter | https://lpsn.dsmz.de/genus/citrobacter |
| Citroniella | https://lpsn.dsmz.de/genus/citroniella |
| Clavibacter | https://lpsn.dsmz.de/genus/clavibacter |
| Clevelandina | https://lpsn.dsmz.de/genus/clevelandina |
| Cloacibacillus | https://lpsn.dsmz.de/genus/cloacibacillus |
| Cloacibacterium | https://lpsn.dsmz.de/genus/cloacibacterium |
| Clostridiisalibacter | https://lpsn.dsmz.de/genus/clostridiisalibacter |
| Clostridioides | https://lpsn.dsmz.de/genus/clostridioides |
| Clostridium | https://lpsn.dsmz.de/genus/clostridium |
| Cnuella | https://lpsn.dsmz.de/genus/cnuella |
| Cnuibacter | https://lpsn.dsmz.de/genus/cnuibacter |
| Cobetia | https://lpsn.dsmz.de/genus/cobetia |
| Cochleicola | https://lpsn.dsmz.de/genus/cochleicola |
| Cochlodiniinecator | https://lpsn.dsmz.de/genus/cochlodiniinecator |
| Cocleimonas | https://lpsn.dsmz.de/genus/cocleimonas |
| Coenonia | https://lpsn.dsmz.de/genus/coenonia |
| Coetzeea | https://lpsn.dsmz.de/genus/coetzeea |
| Cognatazoarcus | https://lpsn.dsmz.de/genus/cognatazoarcus |
| Cognaticolwellia | https://lpsn.dsmz.de/genus/cognaticolwellia |
| Cognatishimia | https://lpsn.dsmz.de/genus/cognatishimia |
| Cognatiyoonia | https://lpsn.dsmz.de/genus/cognatiyoonia |
| Cohaesibacter | https://lpsn.dsmz.de/genus/cohaesibacter |
| Cohnella | https://lpsn.dsmz.de/genus/cohnella |
| Collimonas | https://lpsn.dsmz.de/genus/collimonas |
| Collinsella | https://lpsn.dsmz.de/genus/collinsella |
| Colwellia | https://lpsn.dsmz.de/genus/colwellia |
| Comamonas | https://lpsn.dsmz.de/genus/comamonas |
| Commensalibacter | https://lpsn.dsmz.de/genus/commensalibacter |

| | |
|---|---|
| Companilactobacillus | https://lpsn.dsmz.de/genus/companilactobacillus |
| Compostibacillus | https://lpsn.dsmz.de/genus/compostibacillus |
| Compostibacter | https://lpsn.dsmz.de/genus/compostibacter |
| Compostimonas | https://lpsn.dsmz.de/genus/compostimonas |
| Conchiformibium | https://lpsn.dsmz.de/genus/conchiformibium |
| Conchiformibius | https://lpsn.dsmz.de/genus/conchiformibius |
| Conexibacter | https://lpsn.dsmz.de/genus/conexibacter |
| Conexivisphaera | https://lpsn.dsmz.de/genus/conexivisphaera |
| Confluentibacter | https://lpsn.dsmz.de/genus/confluentibacter |
| Confluentimicrobium | https://lpsn.dsmz.de/genus/confluentimicrobium |
| Conglomeromonas | https://lpsn.dsmz.de/genus/conglomeromonas |
| Congregibacter | https://lpsn.dsmz.de/genus/congregibacter |
| Conservatibacter | https://lpsn.dsmz.de/genus/conservatibacter |
| Consotaella | https://lpsn.dsmz.de/genus/consotaella |
| Constrictibacter | https://lpsn.dsmz.de/genus/constrictibacter |
| Convivina | https://lpsn.dsmz.de/genus/convivina |
| Conyzicola | https://lpsn.dsmz.de/genus/conyzicola |
| Copranaerobaculum | https://lpsn.dsmz.de/genus/copranaerobaculum |
| Coprobacillus | https://lpsn.dsmz.de/genus/coprobacillus |
| Coprobacter | https://lpsn.dsmz.de/genus/coprobacter |
| Coprococcus | https://lpsn.dsmz.de/genus/coprococcus |
| Coprothermobacter | https://lpsn.dsmz.de/genus/coprothermobacter |
| Coraliihabitans | https://lpsn.dsmz.de/genus/coraliihabitans |
| Coraliomargarita | https://lpsn.dsmz.de/genus/coraliomargarita |
| Corallibacter | https://lpsn.dsmz.de/genus/corallibacter |
| Corallincola | https://lpsn.dsmz.de/genus/corallincola |
| Corallococcus | https://lpsn.dsmz.de/genus/corallococcus |
| Coralloluteibacterium | https://lpsn.dsmz.de/genus/coralloluteibacterium |
| Corallomonas | https://lpsn.dsmz.de/genus/corallomonas |
| Coriobacterium | https://lpsn.dsmz.de/genus/coriobacterium |
| Corticibacter | https://lpsn.dsmz.de/genus/corticibacter |
| Corticibacterium | https://lpsn.dsmz.de/genus/corticibacterium |
| Corticicoccus | https://lpsn.dsmz.de/genus/corticicoccus |
| Corticimicrobacter | https://lpsn.dsmz.de/genus/corticimicrobacter |
| Corynebacterium | https://lpsn.dsmz.de/genus/corynebacterium |
| Cosenzaea | https://lpsn.dsmz.de/genus/cosenzaea |
| Costertonia | https://lpsn.dsmz.de/genus/costertonia |
| Couchioplanes | https://lpsn.dsmz.de/genus/couchioplanes |
| Cowdria | https://lpsn.dsmz.de/genus/cowdria |
| Coxiella | https://lpsn.dsmz.de/genus/coxiella |
| Crabtreella | https://lpsn.dsmz.de/genus/crabtreella |
| Crassaminicella | https://lpsn.dsmz.de/genus/crassaminicella |
| Crateriforma | https://lpsn.dsmz.de/genus/crateriforma |
| Craterilacuibacter | https://lpsn.dsmz.de/genus/craterilacuibacter |

| | |
|---|---|
| Craurococcus | https://lpsn.dsmz.de/genus/craurococcus |
| Crenalkalicoccus | https://lpsn.dsmz.de/genus/crenalkalicoccus |
| Crenobacter | https://lpsn.dsmz.de/genus/crenobacter |
| Crenotalea | https://lpsn.dsmz.de/genus/crenotalea |
| Crenothrix | https://lpsn.dsmz.de/genus/crenothrix |
| Cribrihabitans | https://lpsn.dsmz.de/genus/cribrihabitans |
| Cricetibacter | https://lpsn.dsmz.de/genus/cricetibacter |
| Criibacterium | https://lpsn.dsmz.de/genus/criibacterium |
| Cristispira | https://lpsn.dsmz.de/genus/cristispira |
| Croceibacter | https://lpsn.dsmz.de/genus/croceibacter |
| Croceibacterium | https://lpsn.dsmz.de/genus/croceibacterium |
| Croceicoccus | https://lpsn.dsmz.de/genus/croceicoccus |
| Croceifilum | https://lpsn.dsmz.de/genus/croceifilum |
| Croceimicrobium | https://lpsn.dsmz.de/genus/croceimicrobium |
| Croceitalea | https://lpsn.dsmz.de/genus/croceitalea |
| Croceivirga | https://lpsn.dsmz.de/genus/croceivirga |
| Crocinitomix | https://lpsn.dsmz.de/genus/crocinitomix |
| Cronobacter | https://lpsn.dsmz.de/genus/cronobacter |
| Crossiella | https://lpsn.dsmz.de/genus/crossiella |
| Cruoricaptor | https://lpsn.dsmz.de/genus/cruoricaptor |
| Cryobacterium | https://lpsn.dsmz.de/genus/cryobacterium |
| Cryomorpha | https://lpsn.dsmz.de/genus/cryomorpha |
| Cryptanaerobacter | https://lpsn.dsmz.de/genus/cryptanaerobacter |
| Cryptobacterium | https://lpsn.dsmz.de/genus/cryptobacterium |
| Cryptosporangium | https://lpsn.dsmz.de/genus/cryptosporangium |
| Cucumibacter | https://lpsn.dsmz.de/genus/cucumibacter |
| Culicoidibacter | https://lpsn.dsmz.de/genus/culicoidibacter |
| Cumulibacter | https://lpsn.dsmz.de/genus/cumulibacter |
| Cuneatibacter | https://lpsn.dsmz.de/genus/cuneatibacter |
| Cuniculiplasma | https://lpsn.dsmz.de/genus/cuniculiplasma |
| Cupidesulfovibrio | https://lpsn.dsmz.de/genus/cupidesulfovibrio |
| Cupriavidus | https://lpsn.dsmz.de/genus/cupriavidus |
| Curtobacterium | https://lpsn.dsmz.de/genus/curtobacterium |
| Curvibacter | https://lpsn.dsmz.de/genus/curvibacter |
| Curvivirga | https://lpsn.dsmz.de/genus/curvivirga |
| Cutibacterium | https://lpsn.dsmz.de/genus/cutibacterium |
| Cyanobacterium | https://lpsn.dsmz.de/genus/cyanobacterium |
| Cyanobium | https://lpsn.dsmz.de/genus/cyanobium |
| Cyanothece | https://lpsn.dsmz.de/genus/cyanothece |
| Cyclobacterium | https://lpsn.dsmz.de/genus/cyclobacterium |
| Cycloclasticus | https://lpsn.dsmz.de/genus/cycloclasticus |
| Cypionkella | https://lpsn.dsmz.de/genus/cypionkella |
| Cysteiniphilum | https://lpsn.dsmz.de/genus/cysteiniphilum |
| Cystobacter | https://lpsn.dsmz.de/genus/cystobacter |

| | |
|---|---|
| Cytobacillus | https://lpsn.dsmz.de/genus/cytobacillus |
| Cytophaga | https://lpsn.dsmz.de/genus/cytophaga |
| Dactylosporangium | https://lpsn.dsmz.de/genus/dactylosporangium |
| Daeguia | https://lpsn.dsmz.de/genus/daeguia |
| Daejeonella | https://lpsn.dsmz.de/genus/daejeonella |
| Daejeonia | https://lpsn.dsmz.de/genus/daejeonia |
| Dankookia | https://lpsn.dsmz.de/genus/dankookia |
| Dasania | https://lpsn.dsmz.de/genus/dasania |
| Dechloromonas | https://lpsn.dsmz.de/genus/dechloromonas |
| Dechlorosoma | https://lpsn.dsmz.de/genus/dechlorosoma |
| Deefgea | https://lpsn.dsmz.de/genus/deefgea |
| Deferribacter | https://lpsn.dsmz.de/genus/deferribacter |
| Deferrisoma | https://lpsn.dsmz.de/genus/deferrisoma |
| Defluvibacter | https://lpsn.dsmz.de/genus/defluvibacter |
| Defluvicoccus | https://lpsn.dsmz.de/genus/defluvicoccus |
| Defluviicoccus | https://lpsn.dsmz.de/genus/defluviicoccus |
| Defluviimonas | https://lpsn.dsmz.de/genus/defluviimonas |
| Defluviitalea | https://lpsn.dsmz.de/genus/defluviitalea |
| Defluviitoga | https://lpsn.dsmz.de/genus/defluviitoga |
| Dehalobacter | https://lpsn.dsmz.de/genus/dehalobacter |
| Dehalococcoides | https://lpsn.dsmz.de/genus/dehalococcoides |
| Dehalogenimonas | https://lpsn.dsmz.de/genus/dehalogenimonas |
| Dehalospirillum | https://lpsn.dsmz.de/genus/dehalospirillum |
| Deinobacter | https://lpsn.dsmz.de/genus/deinobacter |
| Deinobacterium | https://lpsn.dsmz.de/genus/deinobacterium |
| Deinococcus | https://lpsn.dsmz.de/genus/deinococcus |
| Deleya | https://lpsn.dsmz.de/genus/deleya |
| Delftia | https://lpsn.dsmz.de/genus/delftia |
| Dellaglioa | https://lpsn.dsmz.de/genus/dellaglioa |
| Demequina | https://lpsn.dsmz.de/genus/demequina |
| Demetria | https://lpsn.dsmz.de/genus/demetria |
| Deminuibacter | https://lpsn.dsmz.de/genus/deminuibacter |
| Dendrosporobacter | https://lpsn.dsmz.de/genus/dendrosporobacter |
| Denitratisoma | https://lpsn.dsmz.de/genus/denitratisoma |
| Denitrificimonas | https://lpsn.dsmz.de/genus/denitrificimonas |
| Denitrobacterium | https://lpsn.dsmz.de/genus/denitrobacterium |
| Denitrobaculum | https://lpsn.dsmz.de/genus/denitrobaculum |
| Denitrovibrio | https://lpsn.dsmz.de/genus/denitrovibrio |
| Dermabacter | https://lpsn.dsmz.de/genus/dermabacter |
| Dermacoccus | https://lpsn.dsmz.de/genus/dermacoccus |
| Dermatophilus | https://lpsn.dsmz.de/genus/dermatophilus |
| Derxia | https://lpsn.dsmz.de/genus/derxia |
| Desemzia | https://lpsn.dsmz.de/genus/desemzia |
| Desertactinospora | https://lpsn.dsmz.de/genus/desertactinospora |

| | |
|---|---|
| Desertiactinospora | https://lpsn.dsmz.de/genus/desertiactinospora |
| Desertibacillus | https://lpsn.dsmz.de/genus/desertibacillus |
| Desertibacter | https://lpsn.dsmz.de/genus/desertibacter |
| Deserticella | https://lpsn.dsmz.de/genus/deserticella |
| Desertihabitans | https://lpsn.dsmz.de/genus/desertihabitans |
| Desertimonas | https://lpsn.dsmz.de/genus/desertimonas |
| Desmospora | https://lpsn.dsmz.de/genus/desmospora |
| Desulfacinum | https://lpsn.dsmz.de/genus/desulfacinum |
| Desulfallas | https://lpsn.dsmz.de/genus/desulfallas |
| Desulfarculus | https://lpsn.dsmz.de/genus/desulfarculus |
| Desulfatibacillum | https://lpsn.dsmz.de/genus/desulfatibacillum |
| Desulfatiferula | https://lpsn.dsmz.de/genus/desulfatiferula |
| Desulfatiglans | https://lpsn.dsmz.de/genus/desulfatiglans |
| Desulfatirhabdium | https://lpsn.dsmz.de/genus/desulfatirhabdium |
| Desulfatitalea | https://lpsn.dsmz.de/genus/desulfatitalea |
| Desulfitibacter | https://lpsn.dsmz.de/genus/desulfitibacter |
| Desulfitispora | https://lpsn.dsmz.de/genus/desulfitispora |
| Desulfitobacterium | https://lpsn.dsmz.de/genus/desulfitobacterium |
| Desulfobacca | https://lpsn.dsmz.de/genus/desulfobacca |
| Desulfobacter | https://lpsn.dsmz.de/genus/desulfobacter |
| Desulfobacterium | https://lpsn.dsmz.de/genus/desulfobacterium |
| Desulfobacula | https://lpsn.dsmz.de/genus/desulfobacula |
| Desulfobaculum | https://lpsn.dsmz.de/genus/desulfobaculum |
| Desulfobotulus | https://lpsn.dsmz.de/genus/desulfobotulus |
| Desulfobulbus | https://lpsn.dsmz.de/genus/desulfobulbus |
| Desulfocapsa | https://lpsn.dsmz.de/genus/desulfocapsa |
| Desulfocarbo | https://lpsn.dsmz.de/genus/desulfocarbo |
| Desulfocastanea | https://lpsn.dsmz.de/genus/desulfocastanea |
| Desulfocella | https://lpsn.dsmz.de/genus/desulfocella |
| Desulfocicer | https://lpsn.dsmz.de/genus/desulfocicer |
| Desulfococcus | https://lpsn.dsmz.de/genus/desulfococcus |
| Desulfoconvexum | https://lpsn.dsmz.de/genus/desulfoconvexum |
| Desulfocucumis | https://lpsn.dsmz.de/genus/desulfocucumis |
| Desulfocurvibacter | https://lpsn.dsmz.de/genus/desulfocurvibacter |
| Desulfocurvus | https://lpsn.dsmz.de/genus/desulfocurvus |
| Desulfofaba | https://lpsn.dsmz.de/genus/desulfofaba |
| Desulfofalx | https://lpsn.dsmz.de/genus/desulfofalx |
| Desulfofarcimen | https://lpsn.dsmz.de/genus/desulfofarcimen |
| Desulfoferrobacter | https://lpsn.dsmz.de/genus/desulfoferrobacter |
| Desulfofrigus | https://lpsn.dsmz.de/genus/desulfofrigus |
| Desulfofundulus | https://lpsn.dsmz.de/genus/desulfofundulus |
| Desulfofustis | https://lpsn.dsmz.de/genus/desulfofustis |
| Desulfoglaeba | https://lpsn.dsmz.de/genus/desulfoglaeba |
| Desulfogranum | https://lpsn.dsmz.de/genus/desulfogranum |

| | |
|---|---|
| Desulfohalobium | https://lpsn.dsmz.de/genus/desulfohalobium |
| Desulfohalotomaculum | https://lpsn.dsmz.de/genus/desulfohalotomaculum |
| Desulfohalovibrio | https://lpsn.dsmz.de/genus/desulfohalovibrio |
| Desulfolucanica | https://lpsn.dsmz.de/genus/desulfolucanica |
| Desulfolucianica | https://lpsn.dsmz.de/genus/desulfolucianica |
| Desulfoluna | https://lpsn.dsmz.de/genus/desulfoluna |
| Desulfolutivibrio | https://lpsn.dsmz.de/genus/desulfolutivibrio |
| Desulfomarina | https://lpsn.dsmz.de/genus/desulfomarina |
| Desulfomicrobium | https://lpsn.dsmz.de/genus/desulfomicrobium |
| Desulfomonas | https://lpsn.dsmz.de/genus/desulfomonas |
| Desulfomonile | https://lpsn.dsmz.de/genus/desulfomonile |
| Desulfomusa | https://lpsn.dsmz.de/genus/desulfomusa |
| Desulfonatronobacter | https://lpsn.dsmz.de/genus/desulfonatronobacter |
| Desulfonatronospira | https://lpsn.dsmz.de/genus/desulfonatronospira |
| Desulfonatronovibrio | https://lpsn.dsmz.de/genus/desulfonatronovibrio |
| Desulfonatronum | https://lpsn.dsmz.de/genus/desulfonatronum |
| Desulfonauticus | https://lpsn.dsmz.de/genus/desulfonauticus |
| Desulfonema | https://lpsn.dsmz.de/genus/desulfonema |
| Desulfonispora | https://lpsn.dsmz.de/genus/desulfonispora |
| Desulfopila | https://lpsn.dsmz.de/genus/desulfopila |
| Desulfoplanes | https://lpsn.dsmz.de/genus/desulfoplanes |
| Desulfoprunum | https://lpsn.dsmz.de/genus/desulfoprunum |
| Desulforadius | https://lpsn.dsmz.de/genus/desulforadius |
| Desulforamulus | https://lpsn.dsmz.de/genus/desulforamulus |
| Desulforapulum | https://lpsn.dsmz.de/genus/desulforapulum |
| Desulforegula | https://lpsn.dsmz.de/genus/desulforegula |
| Desulforhabdus | https://lpsn.dsmz.de/genus/desulforhabdus |
| Desulforhopalus | https://lpsn.dsmz.de/genus/desulforhopalus |
| Desulfosalsimonas | https://lpsn.dsmz.de/genus/desulfosalsimonas |
| Desulfosarcina | https://lpsn.dsmz.de/genus/desulfosarcina |
| Desulfoscipio | https://lpsn.dsmz.de/genus/desulfoscipio |
| Desulfosoma | https://lpsn.dsmz.de/genus/desulfosoma |
| Desulfospira | https://lpsn.dsmz.de/genus/desulfospira |
| Desulfosporosinus | https://lpsn.dsmz.de/genus/desulfosporosinus |
| Desulfosudis | https://lpsn.dsmz.de/genus/desulfosudis |
| Desulfotalea | https://lpsn.dsmz.de/genus/desulfotalea |
| Desulfothermobacter | https://lpsn.dsmz.de/genus/desulfothermobacter |
| Desulfothermus | https://lpsn.dsmz.de/genus/desulfothermus |
| Desulfotignum | https://lpsn.dsmz.de/genus/desulfotignum |
| Desulfotomaculum | https://lpsn.dsmz.de/genus/desulfotomaculum |
| Desulfotruncus | https://lpsn.dsmz.de/genus/desulfotruncus |
| Desulfovectis | https://lpsn.dsmz.de/genus/desulfovectis |
| Desulfovermiculus | https://lpsn.dsmz.de/genus/desulfovermiculus |
| Desulfovibrio | https://lpsn.dsmz.de/genus/desulfovibrio |

| | |
|---|---|
| Desulfovirga | https://lpsn.dsmz.de/genus/desulfovirga |
| Desulfovirgula | https://lpsn.dsmz.de/genus/desulfovirgula |
| Desulfurella | https://lpsn.dsmz.de/genus/desulfurella |
| Desulfuribacillus | https://lpsn.dsmz.de/genus/desulfuribacillus |
| Desulfurispira | https://lpsn.dsmz.de/genus/desulfurispira |
| Desulfurispirillum | https://lpsn.dsmz.de/genus/desulfurispirillum |
| Desulfurispora | https://lpsn.dsmz.de/genus/desulfurispora |
| Desulfurivibrio | https://lpsn.dsmz.de/genus/desulfurivibrio |
| Desulfurobacterium | https://lpsn.dsmz.de/genus/desulfurobacterium |
| Desulfurococcus | https://lpsn.dsmz.de/genus/desulfurococcus |
| Desulfurolobus | https://lpsn.dsmz.de/genus/desulfurolobus |
| Desulfuromonas | https://lpsn.dsmz.de/genus/desulfuromonas |
| Desulfuromusa | https://lpsn.dsmz.de/genus/desulfuromusa |
| Dethiobacter | https://lpsn.dsmz.de/genus/dethiobacter |
| Dethiosulfatarculus | https://lpsn.dsmz.de/genus/dethiosulfatarculus |
| Dethiosulfatibacter | https://lpsn.dsmz.de/genus/dethiosulfatibacter |
| Dethiosulfovibrio | https://lpsn.dsmz.de/genus/dethiosulfovibrio |
| Devosia | https://lpsn.dsmz.de/genus/devosia |
| Devriesea | https://lpsn.dsmz.de/genus/devriesea |
| Dialister | https://lpsn.dsmz.de/genus/dialister |
| Diaminobutyricibacter | https://lpsn.dsmz.de/genus/diaminobutyricibacter |
| Diaminobutyricimonas | https://lpsn.dsmz.de/genus/diaminobutyricimonas |
| Diaphorobacter | https://lpsn.dsmz.de/genus/diaphorobacter |
| Dichelobacter | https://lpsn.dsmz.de/genus/dichelobacter |
| Dichotomicrobium | https://lpsn.dsmz.de/genus/dichotomicrobium |
| Dickeya | https://lpsn.dsmz.de/genus/dickeya |
| Dictyobacter | https://lpsn.dsmz.de/genus/dictyobacter |
| Dictyoglomus | https://lpsn.dsmz.de/genus/dictyoglomus |
| Dielma | https://lpsn.dsmz.de/genus/dielma |
| Dietzia | https://lpsn.dsmz.de/genus/dietzia |
| Dinghuibacter | https://lpsn.dsmz.de/genus/dinghuibacter |
| Dinoroseobacter | https://lpsn.dsmz.de/genus/dinoroseobacter |
| Diplocalyx | https://lpsn.dsmz.de/genus/diplocalyx |
| Diplocloster | https://lpsn.dsmz.de/genus/diplocloster |
| Diplorickettsia | https://lpsn.dsmz.de/genus/diplorickettsia |
| Dissulfuribacter | https://lpsn.dsmz.de/genus/dissulfuribacter |
| Dissulfurimicrobium | https://lpsn.dsmz.de/genus/dissulfurimicrobium |
| Dissulfurirhabdus | https://lpsn.dsmz.de/genus/dissulfurirhabdus |
| Dissulfurispira | https://lpsn.dsmz.de/genus/dissulfurispira |
| Dokdonella | https://lpsn.dsmz.de/genus/dokdonella |
| Dokdonia | https://lpsn.dsmz.de/genus/dokdonia |
| Dolosicoccus | https://lpsn.dsmz.de/genus/dolosicoccus |
| Dolosigranulum | https://lpsn.dsmz.de/genus/dolosigranulum |
| Domibacillus | https://lpsn.dsmz.de/genus/domibacillus |

| | |
|---|---|
| Donghaeana | https://lpsn.dsmz.de/genus/donghaeana |
| Donghicola | https://lpsn.dsmz.de/genus/donghicola |
| Dongia | https://lpsn.dsmz.de/genus/dongia |
| Dongshaea | https://lpsn.dsmz.de/genus/dongshaea |
| Dorea | https://lpsn.dsmz.de/genus/dorea |
| Draconibacterium | https://lpsn.dsmz.de/genus/draconibacterium |
| Dubosiella | https://lpsn.dsmz.de/genus/dubosiella |
| Duganella | https://lpsn.dsmz.de/genus/duganella |
| Duncaniella | https://lpsn.dsmz.de/genus/duncaniella |
| Dyadobacter | https://lpsn.dsmz.de/genus/dyadobacter |
| Dyella | https://lpsn.dsmz.de/genus/dyella |
| Dysgonomonas | https://lpsn.dsmz.de/genus/dysgonomonas |
| Dysosmobacter | https://lpsn.dsmz.de/genus/dysosmobacter |
| Echinicola | https://lpsn.dsmz.de/genus/echinicola |
| Echinimonas | https://lpsn.dsmz.de/genus/echinimonas |
| Ectobacillus | https://lpsn.dsmz.de/genus/ectobacillus |
| Ectothiorhodosinus | https://lpsn.dsmz.de/genus/ectothiorhodosinus |
| Ectothiorhodospira | https://lpsn.dsmz.de/genus/ectothiorhodospira |
| Edaphobacter | https://lpsn.dsmz.de/genus/edaphobacter |
| Edaphobaculum | https://lpsn.dsmz.de/genus/edaphobaculum |
| Edaphocola | https://lpsn.dsmz.de/genus/edaphocola |
| Edaphorhabdus | https://lpsn.dsmz.de/genus/edaphorhabdus |
| Edwardsiella | https://lpsn.dsmz.de/genus/edwardsiella |
| Effluviibacter | https://lpsn.dsmz.de/genus/effluviibacter |
| Effusibacillus | https://lpsn.dsmz.de/genus/effusibacillus |
| Eggerthella | https://lpsn.dsmz.de/genus/eggerthella-1 |
| Eggerthella | https://lpsn.dsmz.de/genus/eggerthella |
| Eggerthia | https://lpsn.dsmz.de/genus/eggerthia |
| Egibacter | https://lpsn.dsmz.de/genus/egibacter |
| Egicoccus | https://lpsn.dsmz.de/genus/egicoccus |
| Ehrlichia | https://lpsn.dsmz.de/genus/ehrlichia |
| Eikenella | https://lpsn.dsmz.de/genus/eikenella |
| Eilatimonas | https://lpsn.dsmz.de/genus/eilatimonas |
| Eionea | https://lpsn.dsmz.de/genus/eionea |
| Eisenbergiella | https://lpsn.dsmz.de/genus/eisenbergiella |
| Eisenibacter | https://lpsn.dsmz.de/genus/eisenibacter |
| Ekhidna | https://lpsn.dsmz.de/genus/ekhidna |
| Elioraea | https://lpsn.dsmz.de/genus/elioraea |
| Elizabethkingia | https://lpsn.dsmz.de/genus/elizabethkingia |
| Elkelangia | https://lpsn.dsmz.de/genus/elkelangia |
| Ellagibacter | https://lpsn.dsmz.de/genus/ellagibacter |
| Elstera | https://lpsn.dsmz.de/genus/elstera |
| Elusimicrobium | https://lpsn.dsmz.de/genus/elusimicrobium |
| Elytrosporangium | https://lpsn.dsmz.de/genus/elytrosporangium |

| | |
|---|---|
| Embleya | https://lpsn.dsmz.de/genus/embleya |
| Emcibacter | https://lpsn.dsmz.de/genus/emcibacter |
| Empedobacter | https://lpsn.dsmz.de/genus/empedobacter |
| Emticicia | https://lpsn.dsmz.de/genus/emticicia |
| Endobacter | https://lpsn.dsmz.de/genus/endobacter |
| Endobacterium | https://lpsn.dsmz.de/genus/endobacterium |
| Endomicrobium | https://lpsn.dsmz.de/genus/endomicrobium |
| Endothiovibrio | https://lpsn.dsmz.de/genus/endothiovibrio |
| Endozoicomonas | https://lpsn.dsmz.de/genus/endozoicomonas |
| Enemella | https://lpsn.dsmz.de/genus/enemella |
| Enhydrobacter | https://lpsn.dsmz.de/genus/enhydrobacter |
| Enhygromyxa | https://lpsn.dsmz.de/genus/enhygromyxa |
| Enorma | https://lpsn.dsmz.de/genus/enorma |
| Ensifer | https://lpsn.dsmz.de/genus/ensifer |
| Enteractinococcus | https://lpsn.dsmz.de/genus/enteractinococcus |
| Enterobacillus | https://lpsn.dsmz.de/genus/enterobacillus |
| Enterobacter | https://lpsn.dsmz.de/genus/enterobacter |
| Enterocloster | https://lpsn.dsmz.de/genus/enterocloster |
| Enterococcus | https://lpsn.dsmz.de/genus/enterococcus |
| Enterorhabdus | https://lpsn.dsmz.de/genus/enterorhabdus |
| Enteroscipio | https://lpsn.dsmz.de/genus/enteroscipio |
| Enterovibrio | https://lpsn.dsmz.de/genus/enterovibrio |
| Enterovirga | https://lpsn.dsmz.de/genus/enterovirga |
| Entomobacter | https://lpsn.dsmz.de/genus/entomobacter |
| Entomohabitans | https://lpsn.dsmz.de/genus/entomohabitans |
| Entomomonas | https://lpsn.dsmz.de/genus/entomomonas |
| Entomoplasma | https://lpsn.dsmz.de/genus/entomoplasma |
| Eoetvoesia | https://lpsn.dsmz.de/genus/eoetvoesia |
| Eperythrozoon | https://lpsn.dsmz.de/genus/eperythrozoon |
| Ephemeroptericola | https://lpsn.dsmz.de/genus/ephemeroptericola |
| Epibacterium | https://lpsn.dsmz.de/genus/epibacterium |
| Epidermidibacterium | https://lpsn.dsmz.de/genus/epidermidibacterium |
| Epigloeosphaera | https://lpsn.dsmz.de/genus/epigloeosphaera |
| Epilithonimonas | https://lpsn.dsmz.de/genus/epilithonimonas |
| Ercella | https://lpsn.dsmz.de/genus/ercella |
| Ereboglobus | https://lpsn.dsmz.de/genus/ereboglobus |
| Eremococcus | https://lpsn.dsmz.de/genus/eremococcus |
| Erwinia | https://lpsn.dsmz.de/genus/erwinia |
| Erysipelothrix | https://lpsn.dsmz.de/genus/erysipelothrix |
| Erythrobacter | https://lpsn.dsmz.de/genus/erythrobacter |
| Erythromicrobium | https://lpsn.dsmz.de/genus/erythromicrobium |
| Erythromonas | https://lpsn.dsmz.de/genus/erythromonas |
| Escherichia | https://lpsn.dsmz.de/genus/escherichia |
| Ethanoligenens | https://lpsn.dsmz.de/genus/ethanoligenens |

| | |
|---|---|
| Eubacterium | https://lpsn.dsmz.de/genus/eubacterium |
| Eudoraea | https://lpsn.dsmz.de/genus/eudoraea |
| Euzebya | https://lpsn.dsmz.de/genus/euzebya |
| Euzebyella | https://lpsn.dsmz.de/genus/euzebyella |
| Evansella | https://lpsn.dsmz.de/genus/evansella |
| Ewingella | https://lpsn.dsmz.de/genus/ewingella |
| Excellospora | https://lpsn.dsmz.de/genus/excellospora |
| Exiguobacterium | https://lpsn.dsmz.de/genus/exiguobacterium |
| Exilibacterium | https://lpsn.dsmz.de/genus/exilibacterium |
| Exilispira | https://lpsn.dsmz.de/genus/exilispira |
| Extensimonas | https://lpsn.dsmz.de/genus/extensimonas |
| Extibacter | https://lpsn.dsmz.de/genus/extibacter |
| Ezakiella | https://lpsn.dsmz.de/genus/ezakiella |
| Fabibacter | https://lpsn.dsmz.de/genus/fabibacter |
| Fabivirga | https://lpsn.dsmz.de/genus/fabivirga |
| Facilibium | https://lpsn.dsmz.de/genus/facilibium |
| Facklamia | https://lpsn.dsmz.de/genus/facklamia |
| Faecalibacillus | https://lpsn.dsmz.de/genus/faecalibacillus |
| Faecalibacter | https://lpsn.dsmz.de/genus/faecalibacter |
| Faecalibacterium | https://lpsn.dsmz.de/genus/faecalibacterium |
| Faecalibaculum | https://lpsn.dsmz.de/genus/faecalibaculum |
| Faecalicatena | https://lpsn.dsmz.de/genus/faecalicatena |
| Faecalicoccus | https://lpsn.dsmz.de/genus/faecalicoccus |
| Faecalimonas | https://lpsn.dsmz.de/genus/faecalimonas |
| Faecalitalea | https://lpsn.dsmz.de/genus/faecalitalea |
| Faenia | https://lpsn.dsmz.de/genus/faenia |
| Falcatimonas | https://lpsn.dsmz.de/genus/falcatimonas |
| Falcivibrio | https://lpsn.dsmz.de/genus/falcivibrio |
| Falsarthrobacter | https://lpsn.dsmz.de/genus/falsarthrobacter |
| Falseniella | https://lpsn.dsmz.de/genus/falseniella |
| Falsibacillus | https://lpsn.dsmz.de/genus/falsibacillus |
| Falsihalocynthiibacter | https://lpsn.dsmz.de/genus/falsihalocynthiibacter |
| Falsiporphyromonas | https://lpsn.dsmz.de/genus/falsiporphyromonas |
| Falsirhodobacter | https://lpsn.dsmz.de/genus/falsirhodobacter |
| Falsiroseomonas | https://lpsn.dsmz.de/genus/falsiroseomonas |
| Falsiruegeria | https://lpsn.dsmz.de/genus/falsiruegeria |
| Falsochrobactrum | https://lpsn.dsmz.de/genus/falsochrobactrum |
| Fangia | https://lpsn.dsmz.de/genus/fangia |
| Fannyhessea | https://lpsn.dsmz.de/genus/fannyhessea |
| Fastidiosibacter | https://lpsn.dsmz.de/genus/fastidiosibacter |
| Fastidiosipila | https://lpsn.dsmz.de/genus/fastidiosipila |
| Faucicola | https://lpsn.dsmz.de/genus/faucicola |
| Fenollaria | https://lpsn.dsmz.de/genus/fenollaria |
| Ferdinandcohnia | https://lpsn.dsmz.de/genus/ferdinandcohnia |

| | |
|---|---|
| Fermentibacillus | https://lpsn.dsmz.de/genus/fermentibacillus |
| Fermentimonas | https://lpsn.dsmz.de/genus/fermentimonas |
| Ferribacterium | https://lpsn.dsmz.de/genus/ferribacterium |
| Ferrigenium | https://lpsn.dsmz.de/genus/ferrigenium |
| Ferrimicrobium | https://lpsn.dsmz.de/genus/ferrimicrobium |
| Ferrimonas | https://lpsn.dsmz.de/genus/ferrimonas |
| Ferriphaselus | https://lpsn.dsmz.de/genus/ferriphaselus |
| Ferrithrix | https://lpsn.dsmz.de/genus/ferrithrix |
| Ferroglobus | https://lpsn.dsmz.de/genus/ferroglobus |
| Ferroplasma | https://lpsn.dsmz.de/genus/ferroplasma |
| Ferrovibrio | https://lpsn.dsmz.de/genus/ferrovibrio |
| Ferruginibacter | https://lpsn.dsmz.de/genus/ferruginibacter |
| Ferruginivarius | https://lpsn.dsmz.de/genus/ferruginivarius |
| Fertoeibacter | https://lpsn.dsmz.de/genus/fertoeibacter |
| Fervidicella | https://lpsn.dsmz.de/genus/fervidicella |
| Fervidicoccus | https://lpsn.dsmz.de/genus/fervidicoccus |
| Fervidicola | https://lpsn.dsmz.de/genus/fervidicola |
| Fervidobacterium | https://lpsn.dsmz.de/genus/fervidobacterium |
| Fibrella | https://lpsn.dsmz.de/genus/fibrella |
| Fibrisoma | https://lpsn.dsmz.de/genus/fibrisoma |
| Fibrobacter | https://lpsn.dsmz.de/genus/fibrobacter |
| Fictibacillus | https://lpsn.dsmz.de/genus/fictibacillus |
| Filibacter | https://lpsn.dsmz.de/genus/filibacter |
| Filifactor | https://lpsn.dsmz.de/genus/filifactor |
| Filimonas | https://lpsn.dsmz.de/genus/filimonas |
| Filobacillus | https://lpsn.dsmz.de/genus/filobacillus |
| Filobacterium | https://lpsn.dsmz.de/genus/filobacterium |
| Filomicrobium | https://lpsn.dsmz.de/genus/filomicrobium |
| Fimbriiglobus | https://lpsn.dsmz.de/genus/fimbriiglobus |
| Fimbriimonas | https://lpsn.dsmz.de/genus/fimbriimonas |
| Finegoldia | https://lpsn.dsmz.de/genus/finegoldia |
| Flagellatimonas | https://lpsn.dsmz.de/genus/flagellatimonas |
| Flagellimonas | https://lpsn.dsmz.de/genus/flagellimonas |
| Flammeovirga | https://lpsn.dsmz.de/genus/flammeovirga |
| Flaviaestuariibacter | https://lpsn.dsmz.de/genus/flaviaestuariibacter |
| Flaviaesturariibacter | https://lpsn.dsmz.de/genus/flaviaesturariibacter |
| Flavicella | https://lpsn.dsmz.de/genus/flavicella |
| Flaviflagellibacter | https://lpsn.dsmz.de/genus/flaviflagellibacter |
| Flaviflexus | https://lpsn.dsmz.de/genus/flaviflexus |
| Flavihalobacter | https://lpsn.dsmz.de/genus/flavihalobacter |
| Flavihumibacter | https://lpsn.dsmz.de/genus/flavihumibacter |
| Flavilitoribacter | https://lpsn.dsmz.de/genus/flavilitoribacter |
| Flavimaricola | https://lpsn.dsmz.de/genus/flavimaricola |
| Flavimarina | https://lpsn.dsmz.de/genus/flavimarina |

| | |
|---|---|
| Flavimobilis | https://lpsn.dsmz.de/genus/flavimobilis |
| Flavimonas | https://lpsn.dsmz.de/genus/flavimonas |
| Flavipsychrobacter | https://lpsn.dsmz.de/genus/flavipsychrobacter |
| Flaviramulus | https://lpsn.dsmz.de/genus/flaviramulus |
| Flavirhabdus | https://lpsn.dsmz.de/genus/flavirhabdus |
| Flavisolibacter | https://lpsn.dsmz.de/genus/flavisolibacter |
| Flavitalea | https://lpsn.dsmz.de/genus/flavitalea |
| Flavivirga | https://lpsn.dsmz.de/genus/flavivirga |
| Flavobacterium | https://lpsn.dsmz.de/genus/flavobacterium |
| Flavonifractor | https://lpsn.dsmz.de/genus/flavonifractor |
| Flectobacillus | https://lpsn.dsmz.de/genus/flectobacillus |
| Flexibacter | https://lpsn.dsmz.de/genus/flexibacter |
| Flexilinea | https://lpsn.dsmz.de/genus/flexilinea |
| Flexistipes | https://lpsn.dsmz.de/genus/flexistipes |
| Flexithrix | https://lpsn.dsmz.de/genus/flexithrix |
| Flexivirga | https://lpsn.dsmz.de/genus/flexivirga |
| Flindersiella | https://lpsn.dsmz.de/genus/flindersiella |
| Flintibacter | https://lpsn.dsmz.de/genus/flintibacter |
| Floccifex | https://lpsn.dsmz.de/genus/floccifex |
| Flocculibacter | https://lpsn.dsmz.de/genus/flocculibacter |
| Floricoccus | https://lpsn.dsmz.de/genus/floricoccus |
| Fluoribacter | https://lpsn.dsmz.de/genus/fluoribacter |
| Fluviibacter | https://lpsn.dsmz.de/genus/fluviibacter |
| Fluviibacterium | https://lpsn.dsmz.de/genus/fluviibacterium |
| Fluviicoccus | https://lpsn.dsmz.de/genus/fluviicoccus |
| Fluviicola | https://lpsn.dsmz.de/genus/fluviicola |
| Fluviimonas | https://lpsn.dsmz.de/genus/fluviimonas |
| Fluviispira | https://lpsn.dsmz.de/genus/fluviispira |
| Fodinibacter | https://lpsn.dsmz.de/genus/fodinibacter |
| Fodinibius | https://lpsn.dsmz.de/genus/fodinibius |
| Fodinicola | https://lpsn.dsmz.de/genus/fodinicola |
| Fodinicurvata | https://lpsn.dsmz.de/genus/fodinicurvata |
| Fontibacillus | https://lpsn.dsmz.de/genus/fontibacillus |
| Fontibacter | https://lpsn.dsmz.de/genus/fontibacter |
| Fonticella | https://lpsn.dsmz.de/genus/fonticella |
| Fontimonas | https://lpsn.dsmz.de/genus/fontimonas |
| Formivibrio | https://lpsn.dsmz.de/genus/formivibrio |
| Formosa | https://lpsn.dsmz.de/genus/formosa |
| Formosimonas | https://lpsn.dsmz.de/genus/formosimonas |
| Fournierella | https://lpsn.dsmz.de/genus/fournierella |
| Francisella | https://lpsn.dsmz.de/genus/francisella |
| Franconibacter | https://lpsn.dsmz.de/genus/franconibacter |
| Frankia | https://lpsn.dsmz.de/genus/frankia |
| Franklinella | https://lpsn.dsmz.de/genus/franklinella |

| | |
|---|---|
| Frateuria | https://lpsn.dsmz.de/genus/frateuria |
| Frederiksenia | https://lpsn.dsmz.de/genus/frederiksenia |
| Fredinandcohnia | https://lpsn.dsmz.de/genus/fredinandcohnia |
| Fretibacter | https://lpsn.dsmz.de/genus/fretibacter |
| Fretibacterium | https://lpsn.dsmz.de/genus/fretibacterium |
| Friedmanniella | https://lpsn.dsmz.de/genus/friedmanniella |
| Frigidibacter | https://lpsn.dsmz.de/genus/frigidibacter |
| Frigoribacterium | https://lpsn.dsmz.de/genus/frigoribacterium |
| Frigoriflavimonas | https://lpsn.dsmz.de/genus/frigoriflavimonas |
| Frigoriglobus | https://lpsn.dsmz.de/genus/frigoriglobus |
| Frischella | https://lpsn.dsmz.de/genus/frischella |
| Frisingicoccus | https://lpsn.dsmz.de/genus/frisingicoccus |
| Frondibacter | https://lpsn.dsmz.de/genus/frondibacter |
| Frondicola | https://lpsn.dsmz.de/genus/frondicola |
| Frondihabitans | https://lpsn.dsmz.de/genus/frondihabitans |
| Fructilactobacillus | https://lpsn.dsmz.de/genus/fructilactobacillus |
| Fructobacillus | https://lpsn.dsmz.de/genus/fructobacillus |
| Fuchsiella | https://lpsn.dsmz.de/genus/fuchsiella |
| Fudania | https://lpsn.dsmz.de/genus/fudania |
| Fuerstiella | https://lpsn.dsmz.de/genus/fuerstiella |
| Fulvibacter | https://lpsn.dsmz.de/genus/fulvibacter |
| Fulvimarina | https://lpsn.dsmz.de/genus/fulvimarina |
| Fulvimonas | https://lpsn.dsmz.de/genus/fulvimonas |
| Fulvitalea | https://lpsn.dsmz.de/genus/fulvitalea |
| Fulvivirga | https://lpsn.dsmz.de/genus/fulvivirga |
| Fumia | https://lpsn.dsmz.de/genus/fumia |
| Fundibacter | https://lpsn.dsmz.de/genus/fundibacter |
| Fundicoccus | https://lpsn.dsmz.de/genus/fundicoccus |
| Fundidesulfovibrio | https://lpsn.dsmz.de/genus/fundidesulfovibrio |
| Furfurilactobacillus | https://lpsn.dsmz.de/genus/furfurilactobacillus |
| Fuscovulum | https://lpsn.dsmz.de/genus/fuscovulum |
| Fusibacter | https://lpsn.dsmz.de/genus/fusibacter |
| Fusicatenibacter | https://lpsn.dsmz.de/genus/fusicatenibacter |
| Fusobacterium | https://lpsn.dsmz.de/genus/fusobacterium |
| Gabonibacter | https://lpsn.dsmz.de/genus/gabonibacter |
| Gaetbulibacter | https://lpsn.dsmz.de/genus/gaetbulibacter |
| Gaetbulicola | https://lpsn.dsmz.de/genus/gaetbulicola |
| Gaetbulimicrobium | https://lpsn.dsmz.de/genus/gaetbulimicrobium |
| Gaiella | https://lpsn.dsmz.de/genus/gaiella |
| Galactobacter | https://lpsn.dsmz.de/genus/galactobacter |
| Galbibacter | https://lpsn.dsmz.de/genus/galbibacter |
| Galbitalea | https://lpsn.dsmz.de/genus/galbitalea |
| Galenea | https://lpsn.dsmz.de/genus/galenea |
| Gallaecimonas | https://lpsn.dsmz.de/genus/gallaecimonas |

| | |
|---|---|
| Gallalistipes | https://lpsn.dsmz.de/genus/gallalistipes |
| Gallibacter | https://lpsn.dsmz.de/genus/gallibacter |
| Gallibacterium | https://lpsn.dsmz.de/genus/gallibacterium |
| Gallicola | https://lpsn.dsmz.de/genus/gallicola |
| Gallintestinimicrobium | https://lpsn.dsmz.de/genus/gallintestinimicrobium |
| Gallionella | https://lpsn.dsmz.de/genus/gallionella |
| Galliscardovia | https://lpsn.dsmz.de/genus/galliscardovia |
| Gandjariella | https://lpsn.dsmz.de/genus/gandjariella |
| Gangjinia | https://lpsn.dsmz.de/genus/gangjinia |
| Garciella | https://lpsn.dsmz.de/genus/garciella |
| Gardnerella | https://lpsn.dsmz.de/genus/gardnerella |
| Garicola | https://lpsn.dsmz.de/genus/garicola |
| Garritya | https://lpsn.dsmz.de/genus/garritya |
| Gehongia | https://lpsn.dsmz.de/genus/gehongia |
| Gelatiniphilus | https://lpsn.dsmz.de/genus/gelatiniphilus |
| Gelidibacter | https://lpsn.dsmz.de/genus/gelidibacter |
| Gellertiella | https://lpsn.dsmz.de/genus/gellertiella |
| Gelria | https://lpsn.dsmz.de/genus/gelria |
| Gemella | https://lpsn.dsmz.de/genus/gemella |
| Geminicoccus | https://lpsn.dsmz.de/genus/geminicoccus |
| Geminisphaera | https://lpsn.dsmz.de/genus/geminisphaera |
| Geminocystis | https://lpsn.dsmz.de/genus/geminocystis |
| Gemmata | https://lpsn.dsmz.de/genus/gemmata |
| Gemmatimonas | https://lpsn.dsmz.de/genus/gemmatimonas |
| Gemmiger | https://lpsn.dsmz.de/genus/gemmiger |
| Gemmobacter | https://lpsn.dsmz.de/genus/gemmobacter |
| Geoalkalibacter | https://lpsn.dsmz.de/genus/geoalkalibacter |
| Geoanaerobacter | https://lpsn.dsmz.de/genus/geoanaerobacter |
| Geobacillus | https://lpsn.dsmz.de/genus/geobacillus |
| Geobacter | https://lpsn.dsmz.de/genus/geobacter |
| Geodermatophilus | https://lpsn.dsmz.de/genus/geodermatophilus |
| Geofilum | https://lpsn.dsmz.de/genus/geofilum |
| Geoglobus | https://lpsn.dsmz.de/genus/geoglobus |
| Geojedonia | https://lpsn.dsmz.de/genus/geojedonia |
| Geomesophilobacter | https://lpsn.dsmz.de/genus/geomesophilobacter |
| Geomicrobium | https://lpsn.dsmz.de/genus/geomicrobium |
| Geomobilimonas | https://lpsn.dsmz.de/genus/geomobilimonas |
| Geomonas | https://lpsn.dsmz.de/genus/geomonas |
| Geopsychrobacter | https://lpsn.dsmz.de/genus/geopsychrobacter |
| Georgenia | https://lpsn.dsmz.de/genus/georgenia |
| Georgfuchsia | https://lpsn.dsmz.de/genus/georgfuchsia |
| Georhizobium | https://lpsn.dsmz.de/genus/georhizobium |
| Geosporobacter | https://lpsn.dsmz.de/genus/geosporobacter |
| Geotalea | https://lpsn.dsmz.de/genus/geotalea |

| | |
|---|---|
| Geothermobacter | https://lpsn.dsmz.de/genus/geothermobacter |
| Geothermomicrobium | https://lpsn.dsmz.de/genus/geothermomicrobium |
| Geothrix | https://lpsn.dsmz.de/genus/geothrix |
| Geotoga | https://lpsn.dsmz.de/genus/geotoga |
| Geovibrio | https://lpsn.dsmz.de/genus/geovibrio |
| Gibbsiella | https://lpsn.dsmz.de/genus/gibbsiella |
| Giesbergeria | https://lpsn.dsmz.de/genus/giesbergeria |
| Gilliamella | https://lpsn.dsmz.de/genus/gilliamella |
| Gillisia | https://lpsn.dsmz.de/genus/gillisia |
| Gilvibacter | https://lpsn.dsmz.de/genus/gilvibacter |
| Gilvimarinus | https://lpsn.dsmz.de/genus/gilvimarinus |
| Gimesia | https://lpsn.dsmz.de/genus/gimesia |
| Glaciecola | https://lpsn.dsmz.de/genus/glaciecola |
| Glacieibacterium | https://lpsn.dsmz.de/genus/glacieibacterium |
| Glaciibacter | https://lpsn.dsmz.de/genus/glaciibacter |
| Glaciihabitans | https://lpsn.dsmz.de/genus/glaciihabitans |
| Glaciimonas | https://lpsn.dsmz.de/genus/glaciimonas |
| Glaesserella | https://lpsn.dsmz.de/genus/glaesserella |
| Gleimia | https://lpsn.dsmz.de/genus/gleimia |
| Globicatella | https://lpsn.dsmz.de/genus/globicatella |
| Gluconacetobacter | https://lpsn.dsmz.de/genus/gluconacetobacter |
| Gluconoacetobacter | https://lpsn.dsmz.de/genus/gluconoacetobacter |
| Gluconobacter | https://lpsn.dsmz.de/genus/gluconobacter |
| Glutamicibacter | https://lpsn.dsmz.de/genus/glutamicibacter |
| Glycocaulis | https://lpsn.dsmz.de/genus/glycocaulis |
| Glycomyces | https://lpsn.dsmz.de/genus/glycomyces |
| Goodfellowia | https://lpsn.dsmz.de/genus/goodfellowia |
| Goodfellowiella | https://lpsn.dsmz.de/genus/goodfellowiella |
| Gordona | https://lpsn.dsmz.de/genus/gordona |
| Gordonia | https://lpsn.dsmz.de/genus/gordonia |
| Gordonibacter | https://lpsn.dsmz.de/genus/gordonibacter |
| Gorillibacterium | https://lpsn.dsmz.de/genus/gorillibacterium |
| Gottfriedia | https://lpsn.dsmz.de/genus/gottfriedia |
| Gottschalkia | https://lpsn.dsmz.de/genus/gottschalkia |
| Gracilibacillus | https://lpsn.dsmz.de/genus/gracilibacillus |
| Gracilibacter | https://lpsn.dsmz.de/genus/gracilibacter |
| Gracilimonas | https://lpsn.dsmz.de/genus/gracilimonas |
| Gracilinema | https://lpsn.dsmz.de/genus/gracilinema |
| Grahamella | https://lpsn.dsmz.de/genus/grahamella |
| Gramella | https://lpsn.dsmz.de/genus/gramella |
| Granulibacter | https://lpsn.dsmz.de/genus/granulibacter |
| Granulicatella | https://lpsn.dsmz.de/genus/granulicatella |
| Granulicella | https://lpsn.dsmz.de/genus/granulicella |
| Granulicoccus | https://lpsn.dsmz.de/genus/granulicoccus |

| | |
|---|---|
| Granulimonas | https://lpsn.dsmz.de/genus/granulimonas |
| Granulosicoccus | https://lpsn.dsmz.de/genus/granulosicoccus |
| Grimontia | https://lpsn.dsmz.de/genus/grimontia |
| Gryllotalpicola | https://lpsn.dsmz.de/genus/gryllotalpicola |
| Gudongella | https://lpsn.dsmz.de/genus/gudongella |
| Guggenheimella | https://lpsn.dsmz.de/genus/guggenheimella |
| Gulbenkiania | https://lpsn.dsmz.de/genus/gulbenkiania |
| Gulosibacter | https://lpsn.dsmz.de/genus/gulosibacter |
| Guopingia | https://lpsn.dsmz.de/genus/guopingia |
| Guyparkeria | https://lpsn.dsmz.de/genus/guyparkeria |
| Gymnodinialimonas | https://lpsn.dsmz.de/genus/gymnodinialimonas |
| Gynuella | https://lpsn.dsmz.de/genus/gynuella |
| Gynurincola | https://lpsn.dsmz.de/genus/gynurincola |
| Haematobacter | https://lpsn.dsmz.de/genus/haematobacter |
| Haematomicrobium | https://lpsn.dsmz.de/genus/haematomicrobium |
| Haematospirillum | https://lpsn.dsmz.de/genus/haematospirillum |
| Haemobartonella | https://lpsn.dsmz.de/genus/haemobartonella |
| Haemophilus | https://lpsn.dsmz.de/genus/haemophilus |
| Hafnia | https://lpsn.dsmz.de/genus/hafnia |
| Hahella | https://lpsn.dsmz.de/genus/hahella |
| Hahyoungchilella | https://lpsn.dsmz.de/genus/hahyoungchilella |
| Haladaptatus | https://lpsn.dsmz.de/genus/haladaptatus |
| Halalkaliarchaeum | https://lpsn.dsmz.de/genus/halalkaliarchaeum |
| Halalkalibacillus | https://lpsn.dsmz.de/genus/halalkalibacillus |
| Halalkalibacter | https://lpsn.dsmz.de/genus/halalkalibacter |
| Halalkalibacterium | https://lpsn.dsmz.de/genus/halalkalibacterium |
| Halalkalibacterium | https://lpsn.dsmz.de/genus/halalkalibacterium-1 |
| Halalkalibaculum | https://lpsn.dsmz.de/genus/halalkalibaculum |
| Halalkalicoccus | https://lpsn.dsmz.de/genus/halalkalicoccus |
| Halalkalirubrum | https://lpsn.dsmz.de/genus/halalkalirubrum |
| Halanaeroarchaeum | https://lpsn.dsmz.de/genus/halanaeroarchaeum |
| Halanaerobacter | https://lpsn.dsmz.de/genus/halanaerobacter |
| Halanaerobaculum | https://lpsn.dsmz.de/genus/halanaerobaculum |
| Halanaerobium | https://lpsn.dsmz.de/genus/halanaerobium |
| Halapricum | https://lpsn.dsmz.de/genus/halapricum |
| Halarchaeum | https://lpsn.dsmz.de/genus/halarchaeum |
| Halarcobacter | https://lpsn.dsmz.de/genus/halarcobacter |
| Halarsenatibacter | https://lpsn.dsmz.de/genus/halarsenatibacter |
| Halegenticoccus | https://lpsn.dsmz.de/genus/halegenticoccus |
| Haliangium | https://lpsn.dsmz.de/genus/haliangium |
| Haliea | https://lpsn.dsmz.de/genus/haliea |
| Halioglobus | https://lpsn.dsmz.de/genus/halioglobus |
| Halioxenophilus | https://lpsn.dsmz.de/genus/halioxenophilus |
| Haliscomenobacter | https://lpsn.dsmz.de/genus/haliscomenobacter |

| | |
|---|---|
| Hallella | https://lpsn.dsmz.de/genus/hallella |
| Hallerella | https://lpsn.dsmz.de/genus/hallerella |
| Haloactinobacterium | https://lpsn.dsmz.de/genus/haloactinobacterium |
| Haloactinomyces | https://lpsn.dsmz.de/genus/haloactinomyces |
| Haloactinopolyspora | https://lpsn.dsmz.de/genus/haloactinopolyspora |
| Haloactinospora | https://lpsn.dsmz.de/genus/haloactinospora |
| Haloanaerobacter | https://lpsn.dsmz.de/genus/haloanaerobacter |
| Haloanaerobium | https://lpsn.dsmz.de/genus/haloanaerobium |
| Haloarchaeobius | https://lpsn.dsmz.de/genus/haloarchaeobius |
| Haloarcobacter | https://lpsn.dsmz.de/genus/haloarcobacter |
| Haloarcula | https://lpsn.dsmz.de/genus/haloarcula |
| Halobacillus | https://lpsn.dsmz.de/genus/halobacillus |
| Halobacteriovorax | https://lpsn.dsmz.de/genus/halobacteriovorax |
| Halobacterium | https://lpsn.dsmz.de/genus/halobacterium |
| Halobacteroides | https://lpsn.dsmz.de/genus/halobacteroides |
| Halobaculum | https://lpsn.dsmz.de/genus/halobaculum |
| Halobellus | https://lpsn.dsmz.de/genus/halobellus |
| Halobiforma | https://lpsn.dsmz.de/genus/halobiforma |
| Halobium | https://lpsn.dsmz.de/genus/halobium |
| Halocalculus | https://lpsn.dsmz.de/genus/halocalculus |
| Halocatena | https://lpsn.dsmz.de/genus/halocatena |
| Halocella | https://lpsn.dsmz.de/genus/halocella |
| Halochromatium | https://lpsn.dsmz.de/genus/halochromatium |
| Halococcoides | https://lpsn.dsmz.de/genus/halococcoides |
| Halococcus | https://lpsn.dsmz.de/genus/halococcus |
| Halocynthiibacter | https://lpsn.dsmz.de/genus/halocynthiibacter |
| Halodesulfovibrio | https://lpsn.dsmz.de/genus/halodesulfovibrio |
| Halodesulfurarchaeum | https://lpsn.dsmz.de/genus/halodesulfurarchaeum |
| Halodurantibacterium | https://lpsn.dsmz.de/genus/halodurantibacterium |
| Haloechinothrix | https://lpsn.dsmz.de/genus/haloechinothrix |
| Haloferax | https://lpsn.dsmz.de/genus/haloferax |
| Haloferula | https://lpsn.dsmz.de/genus/haloferula |
| Halofilum | https://lpsn.dsmz.de/genus/halofilum |
| Haloflavibacter | https://lpsn.dsmz.de/genus/haloflavibacter |
| Halogeometricum | https://lpsn.dsmz.de/genus/halogeometricum |
| Haloglomus | https://lpsn.dsmz.de/genus/haloglomus |
| Haloglycomyces | https://lpsn.dsmz.de/genus/haloglycomyces |
| Halogranum | https://lpsn.dsmz.de/genus/halogranum |
| Halohasta | https://lpsn.dsmz.de/genus/halohasta |
| Haloimpatiens | https://lpsn.dsmz.de/genus/haloimpatiens |
| Haloincola | https://lpsn.dsmz.de/genus/haloincola |
| Halolactibacillus | https://lpsn.dsmz.de/genus/halolactibacillus |
| Halolamina | https://lpsn.dsmz.de/genus/halolamina |
| Halomarina | https://lpsn.dsmz.de/genus/halomarina |

| | |
|---|---|
| Halomethanococcus | https://lpsn.dsmz.de/genus/halomethanococcus |
| Halomicroarcula | https://lpsn.dsmz.de/genus/halomicroarcula |
| Halomicrobium | https://lpsn.dsmz.de/genus/halomicrobium |
| Halomicrococcus | https://lpsn.dsmz.de/genus/halomicrococcus |
| Halomonas | https://lpsn.dsmz.de/genus/halomonas |
| Halonatronum | https://lpsn.dsmz.de/genus/halonatronum |
| Halonotius | https://lpsn.dsmz.de/genus/halonotius |
| Haloparvum | https://lpsn.dsmz.de/genus/haloparvum |
| Halopelagius | https://lpsn.dsmz.de/genus/halopelagius |
| Halopenitus | https://lpsn.dsmz.de/genus/halopenitus |
| Halopeptonella | https://lpsn.dsmz.de/genus/halopeptonella |
| Halopiger | https://lpsn.dsmz.de/genus/halopiger |
| Haloplanus | https://lpsn.dsmz.de/genus/haloplanus |
| Haloplasma | https://lpsn.dsmz.de/genus/haloplasma |
| Halopolyspora | https://lpsn.dsmz.de/genus/halopolyspora |
| Haloprofundus | https://lpsn.dsmz.de/genus/haloprofundus |
| Halopseudomonas | https://lpsn.dsmz.de/genus/halopseudomonas |
| Haloquadratum | https://lpsn.dsmz.de/genus/haloquadratum |
| Halorhabdus | https://lpsn.dsmz.de/genus/halorhabdus |
| Halorhodospira | https://lpsn.dsmz.de/genus/halorhodospira |
| Halorientalis | https://lpsn.dsmz.de/genus/halorientalis |
| Halorubellus | https://lpsn.dsmz.de/genus/halorubellus |
| Halorubrobacterium | https://lpsn.dsmz.de/genus/halorubrobacterium |
| Halorubrum | https://lpsn.dsmz.de/genus/halorubrum |
| Halorussus | https://lpsn.dsmz.de/genus/halorussus |
| Halosaccharopolyspora | https://lpsn.dsmz.de/genus/halosaccharopolyspora |
| Halosarcina | https://lpsn.dsmz.de/genus/halosarcina |
| Halosegnis | https://lpsn.dsmz.de/genus/halosegnis |
| Halosiccatus | https://lpsn.dsmz.de/genus/halosiccatus |
| Halosimplex | https://lpsn.dsmz.de/genus/halosimplex |
| Halosolutus | https://lpsn.dsmz.de/genus/halosolutus |
| Halospina | https://lpsn.dsmz.de/genus/halospina |
| Halospirulina | https://lpsn.dsmz.de/genus/halospirulina |
| Halostagnicola | https://lpsn.dsmz.de/genus/halostagnicola |
| Halostella | https://lpsn.dsmz.de/genus/halostella |
| Halotalea | https://lpsn.dsmz.de/genus/halotalea |
| Haloterrigena | https://lpsn.dsmz.de/genus/haloterrigena |
| Halothermothrix | https://lpsn.dsmz.de/genus/halothermothrix |
| Halothiobacillus | https://lpsn.dsmz.de/genus/halothiobacillus |
| Halovarius | https://lpsn.dsmz.de/genus/halovarius |
| Halovenus | https://lpsn.dsmz.de/genus/halovenus |
| Halovibrio | https://lpsn.dsmz.de/genus/halovibrio |
| Halovivax | https://lpsn.dsmz.de/genus/halovivax |
| Halovulum | https://lpsn.dsmz.de/genus/halovulum |

| | |
|---|---|
| Halpernia | https://lpsn.dsmz.de/genus/halpernia |
| Hamadaea | https://lpsn.dsmz.de/genus/hamadaea |
| Hansschlegelia | https://lpsn.dsmz.de/genus/hansschlegelia |
| Hanstruepera | https://lpsn.dsmz.de/genus/hanstruepera |
| Haoranjiania | https://lpsn.dsmz.de/genus/haoranjiania |
| Haploplasma | https://lpsn.dsmz.de/genus/haploplasma |
| Harenicola | https://lpsn.dsmz.de/genus/harenicola |
| Harryflintia | https://lpsn.dsmz.de/genus/harryflintia |
| Hartmannibacter | https://lpsn.dsmz.de/genus/hartmannibacter |
| Hasllibacter | https://lpsn.dsmz.de/genus/hasllibacter |
| Hathewaya | https://lpsn.dsmz.de/genus/hathewaya |
| Hazenella | https://lpsn.dsmz.de/genus/hazenella |
| Helcobacillus | https://lpsn.dsmz.de/genus/helcobacillus |
| Helcococcus | https://lpsn.dsmz.de/genus/helcococcus |
| Helicobacter | https://lpsn.dsmz.de/genus/helicobacter |
| Heliimonas | https://lpsn.dsmz.de/genus/heliimonas |
| Heliobacillus | https://lpsn.dsmz.de/genus/heliobacillus |
| Heliobacterium | https://lpsn.dsmz.de/genus/heliobacterium |
| Heliomarina | https://lpsn.dsmz.de/genus/heliomarina |
| Heliomicrobium | https://lpsn.dsmz.de/genus/heliomicrobium |
| Heliophilum | https://lpsn.dsmz.de/genus/heliophilum |
| Heliorestis | https://lpsn.dsmz.de/genus/heliorestis |
| Heliothrix | https://lpsn.dsmz.de/genus/heliothrix |
| Hellea | https://lpsn.dsmz.de/genus/hellea |
| Helmutkoenigia | https://lpsn.dsmz.de/genus/helmutkoenigia |
| Hemophilus | https://lpsn.dsmz.de/genus/hemophilus |
| Henriciella | https://lpsn.dsmz.de/genus/henriciella |
| Hephaestia | https://lpsn.dsmz.de/genus/hephaestia |
| Herbaspirillum | https://lpsn.dsmz.de/genus/herbaspirillum |
| Herbiconiux | https://lpsn.dsmz.de/genus/herbiconiux |
| Herbidospora | https://lpsn.dsmz.de/genus/herbidospora |
| Herbihabitans | https://lpsn.dsmz.de/genus/herbihabitans |
| Herbinix | https://lpsn.dsmz.de/genus/herbinix |
| Herbivorax | https://lpsn.dsmz.de/genus/herbivorax |
| Herminiimonas | https://lpsn.dsmz.de/genus/herminiimonas |
| Herpetosiphon | https://lpsn.dsmz.de/genus/herpetosiphon |
| Hespellia | https://lpsn.dsmz.de/genus/hespellia |
| Heyndrickxia | https://lpsn.dsmz.de/genus/heyndrickxia |
| Hippea | https://lpsn.dsmz.de/genus/hippea |
| Hirschia | https://lpsn.dsmz.de/genus/hirschia |
| Histidinibacterium | https://lpsn.dsmz.de/genus/histidinibacterium |
| Histophilus | https://lpsn.dsmz.de/genus/histophilus |
| Hoeflea | https://lpsn.dsmz.de/genus/hoeflea |
| Holdemanella | https://lpsn.dsmz.de/genus/holdemanella |

| | |
|---|---|
| Holdemania | https://lpsn.dsmz.de/genus/holdemania |
| Hollandina | https://lpsn.dsmz.de/genus/hollandina |
| Holophaga | https://lpsn.dsmz.de/genus/holophaga |
| Holospora | https://lpsn.dsmz.de/genus/holospora-1 |
| Holospora | https://lpsn.dsmz.de/genus/holospora |
| Holzapfelia | https://lpsn.dsmz.de/genus/holzapfelia |
| Hominimerdicola | https://lpsn.dsmz.de/genus/hominimerdicola |
| Hoministercoradaptatus | https://lpsn.dsmz.de/genus/hoministercoradaptatus |
| Homoserinibacter | https://lpsn.dsmz.de/genus/homoserinibacter |
| Homoserinimonas | https://lpsn.dsmz.de/genus/homoserinimonas |
| Hongia | https://lpsn.dsmz.de/genus/hongia |
| Hongiella | https://lpsn.dsmz.de/genus/hongiella |
| Hongsoonwoonella | https://lpsn.dsmz.de/genus/hongsoonwoonella |
| Hoppeia | https://lpsn.dsmz.de/genus/hoppeia |
| Hornefia | https://lpsn.dsmz.de/genus/hornefia |
| Howardella | https://lpsn.dsmz.de/genus/howardella |
| Hoyosella | https://lpsn.dsmz.de/genus/hoyosella |
| Huaishuia | https://lpsn.dsmz.de/genus/huaishuia |
| Huakuichenia | https://lpsn.dsmz.de/genus/huakuichenia |
| Huanghella | https://lpsn.dsmz.de/genus/huanghella |
| Huaxiibacter | https://lpsn.dsmz.de/genus/huaxiibacter |
| Hugenholtzia | https://lpsn.dsmz.de/genus/hugenholtzia |
| Huintestinicola | https://lpsn.dsmz.de/genus/huintestinicola |
| Humibacillus | https://lpsn.dsmz.de/genus/humibacillus |
| Humibacter | https://lpsn.dsmz.de/genus/humibacter |
| Humicoccus | https://lpsn.dsmz.de/genus/humicoccus |
| Humidesulfovibrio | https://lpsn.dsmz.de/genus/humidesulfovibrio |
| Humihabitans | https://lpsn.dsmz.de/genus/humihabitans |
| Humitalea | https://lpsn.dsmz.de/genus/humitalea |
| Hungateiclostridium | https://lpsn.dsmz.de/genus/hungateiclostridium |
| Hungatella | https://lpsn.dsmz.de/genus/hungatella |
| Hutsoniella | https://lpsn.dsmz.de/genus/hutsoniella |
| Hwangdonia | https://lpsn.dsmz.de/genus/hwangdonia |
| Hwanghaeella | https://lpsn.dsmz.de/genus/hwanghaeella |
| Hwanghaeicola | https://lpsn.dsmz.de/genus/hwanghaeicola |
| Hyalangium | https://lpsn.dsmz.de/genus/hyalangium |
| Hydrobacter | https://lpsn.dsmz.de/genus/hydrobacter |
| Hydrocarboniclastica | https://lpsn.dsmz.de/genus/hydrocarboniclastica |
| Hydrocarboniphaga | https://lpsn.dsmz.de/genus/hydrocarboniphaga |
| Hydrogenibacillus | https://lpsn.dsmz.de/genus/hydrogenibacillus |
| Hydrogeniiclostidium | https://lpsn.dsmz.de/genus/hydrogeniiclostidium |
| Hydrogeniiclostridium | https://lpsn.dsmz.de/genus/hydrogeniiclostridium |
| Hydrogenimonas | https://lpsn.dsmz.de/genus/hydrogenimonas |
| Hydrogenispora | https://lpsn.dsmz.de/genus/hydrogenispora |

| | |
|---|---|
| Hydrogenivirga | https://lpsn.dsmz.de/genus/hydrogenivirga |
| Hydrogenoanaerobacterium | https://lpsn.dsmz.de/genus/hydrogenoanaerobacterium |
| Hydrogenobacter | https://lpsn.dsmz.de/genus/hydrogenobacter |
| Hydrogenobaculum | https://lpsn.dsmz.de/genus/hydrogenobaculum |
| Hydrogenophaga | https://lpsn.dsmz.de/genus/hydrogenophaga |
| Hydrogenophilus | https://lpsn.dsmz.de/genus/hydrogenophilus |
| Hydrogenothermus | https://lpsn.dsmz.de/genus/hydrogenothermus |
| Hydrogenovibrio | https://lpsn.dsmz.de/genus/hydrogenovibrio |
| Hydromonas | https://lpsn.dsmz.de/genus/hydromonas |
| Hydrotalea | https://lpsn.dsmz.de/genus/hydrotalea |
| Hylemonella | https://lpsn.dsmz.de/genus/hylemonella |
| Hymenobacter | https://lpsn.dsmz.de/genus/hymenobacter |
| Hypericibacter | https://lpsn.dsmz.de/genus/hypericibacter |
| Hyperthermus | https://lpsn.dsmz.de/genus/hyperthermus |
| Hyphobacterium | https://lpsn.dsmz.de/genus/hyphobacterium |
| Hyphococcus | https://lpsn.dsmz.de/genus/hyphococcus |
| Hyphomicrobium | https://lpsn.dsmz.de/genus/hyphomicrobium |
| Hyphomonas | https://lpsn.dsmz.de/genus/hyphomonas |
| Hypnocyclicus | https://lpsn.dsmz.de/genus/hypnocyclicus |
| Hyunsoonleella | https://lpsn.dsmz.de/genus/hyunsoonleella |
| Iamia | https://lpsn.dsmz.de/genus/iamia |
| Ichthyenterobacterium | https://lpsn.dsmz.de/genus/ichthyenterobacterium |
| Ichthyobacterium | https://lpsn.dsmz.de/genus/ichthyobacterium |
| Ideonella | https://lpsn.dsmz.de/genus/ideonella |
| Idiomarina | https://lpsn.dsmz.de/genus/idiomarina |
| Ignatzschineria | https://lpsn.dsmz.de/genus/ignatzschineria |
| Ignavibacterium | https://lpsn.dsmz.de/genus/ignavibacterium |
| Ignavigranum | https://lpsn.dsmz.de/genus/ignavigranum |
| Ignicoccus | https://lpsn.dsmz.de/genus/ignicoccus |
| Ignisphaera | https://lpsn.dsmz.de/genus/ignisphaera |
| Ileibacterium | https://lpsn.dsmz.de/genus/ileibacterium |
| Ilumatobacter | https://lpsn.dsmz.de/genus/ilumatobacter |
| Ilyobacter | https://lpsn.dsmz.de/genus/ilyobacter |
| Ilyomonas | https://lpsn.dsmz.de/genus/ilyomonas |
| Imhoffiella | https://lpsn.dsmz.de/genus/imhoffiella |
| Immundisolibacter | https://lpsn.dsmz.de/genus/immundisolibacter |
| Imperialibacter | https://lpsn.dsmz.de/genus/imperialibacter |
| Imtechella | https://lpsn.dsmz.de/genus/imtechella |
| Inconstantimicrobium | https://lpsn.dsmz.de/genus/inconstantimicrobium |
| Indibacter | https://lpsn.dsmz.de/genus/indibacter |
| Indiicoccus | https://lpsn.dsmz.de/genus/indiicoccus |
| Indioceanicola | https://lpsn.dsmz.de/genus/indioceanicola |
| Inhella | https://lpsn.dsmz.de/genus/inhella |
| Inmirania | https://lpsn.dsmz.de/genus/inmirania |

| | |
|---|---|
| Inquilinus | https://lpsn.dsmz.de/genus/inquilinus |
| Insectihabitans | https://lpsn.dsmz.de/genus/insectihabitans |
| Insolitispirillum | https://lpsn.dsmz.de/genus/insolitispirillum |
| Intestinibacter | https://lpsn.dsmz.de/genus/intestinibacter |
| Intestinibaculum | https://lpsn.dsmz.de/genus/intestinibaculum |
| Intestinimonas | https://lpsn.dsmz.de/genus/intestinimonas |
| Intestinirhabdus | https://lpsn.dsmz.de/genus/intestinirhabdus |
| Intrasporangium | https://lpsn.dsmz.de/genus/intrasporangium |
| Iodidimonas | https://lpsn.dsmz.de/genus/iodidimonas |
| Iodobacter | https://lpsn.dsmz.de/genus/iodobacter |
| Irregularibacter | https://lpsn.dsmz.de/genus/irregularibacter |
| Isachenkonia | https://lpsn.dsmz.de/genus/isachenkonia |
| Isobaculum | https://lpsn.dsmz.de/genus/isobaculum |
| Isochromatium | https://lpsn.dsmz.de/genus/isochromatium |
| Isoptericola | https://lpsn.dsmz.de/genus/isoptericola |
| Isosphaera | https://lpsn.dsmz.de/genus/isosphaera |
| Izhakiella | https://lpsn.dsmz.de/genus/izhakiella |
| Jahnella | https://lpsn.dsmz.de/genus/jahnella |
| Jahnia | https://lpsn.dsmz.de/genus/jahnia |
| Janibacter | https://lpsn.dsmz.de/genus/janibacter |
| Jannaschia | https://lpsn.dsmz.de/genus/jannaschia |
| Janthinobacterium | https://lpsn.dsmz.de/genus/janthinobacterium |
| Jatrophihabitans | https://lpsn.dsmz.de/genus/jatrophihabitans |
| Jejudonia | https://lpsn.dsmz.de/genus/jejudonia |
| Jejuia | https://lpsn.dsmz.de/genus/jejuia |
| Jeongeupia | https://lpsn.dsmz.de/genus/jeongeupia |
| Jeotgalibaca | https://lpsn.dsmz.de/genus/jeotgalibaca |
| Jeotgalibacillus | https://lpsn.dsmz.de/genus/jeotgalibacillus |
| Jeotgalicoccus | https://lpsn.dsmz.de/genus/jeotgalicoccus |
| Jezberella | https://lpsn.dsmz.de/genus/jezberella |
| Jhaorihella | https://lpsn.dsmz.de/genus/jhaorihella |
| Jiangella | https://lpsn.dsmz.de/genus/jiangella |
| Jiella | https://lpsn.dsmz.de/genus/jiella |
| Jinshanibacter | https://lpsn.dsmz.de/genus/jinshanibacter |
| Jishengella | https://lpsn.dsmz.de/genus/jishengella |
| Jiulongibacter | https://lpsn.dsmz.de/genus/jiulongibacter |
| Johnsonella | https://lpsn.dsmz.de/genus/johnsonella |
| Jonesia | https://lpsn.dsmz.de/genus/jonesia |
| Jongsikchunia | https://lpsn.dsmz.de/genus/jongsikchunia |
| Jonquetella | https://lpsn.dsmz.de/genus/jonquetella |
| Joostella | https://lpsn.dsmz.de/genus/joostella |
| Jutongia | https://lpsn.dsmz.de/genus/jutongia |
| Kaistella | https://lpsn.dsmz.de/genus/kaistella |
| Kaistia | https://lpsn.dsmz.de/genus/kaistia |

| | |
|---|---|
| Kalamiella | https://lpsn.dsmz.de/genus/kalamiella |
| Kallipyga | https://lpsn.dsmz.de/genus/kallipyga |
| Kallotenue | https://lpsn.dsmz.de/genus/kallotenue |
| Kandeliimicrobium | https://lpsn.dsmz.de/genus/kandeliimicrobium |
| Kandleria | https://lpsn.dsmz.de/genus/kandleria |
| Kangiella | https://lpsn.dsmz.de/genus/kangiella |
| Kangsaoukella | https://lpsn.dsmz.de/genus/kangsaoukella |
| Keratinibaculum | https://lpsn.dsmz.de/genus/keratinibaculum |
| Kerstersia | https://lpsn.dsmz.de/genus/kerstersia |
| Ketobacter | https://lpsn.dsmz.de/genus/ketobacter |
| Kibdelosporangium | https://lpsn.dsmz.de/genus/kibdelosporangium |
| Kiloniella | https://lpsn.dsmz.de/genus/kiloniella |
| Kineobactrum | https://lpsn.dsmz.de/genus/kineobactrum |
| Kineococcus | https://lpsn.dsmz.de/genus/kineococcus |
| Kineosphaera | https://lpsn.dsmz.de/genus/kineosphaera |
| Kineosporia | https://lpsn.dsmz.de/genus/kineosporia |
| Kineothrix | https://lpsn.dsmz.de/genus/kineothrix |
| Kingella | https://lpsn.dsmz.de/genus/kingella |
| Kinneretia | https://lpsn.dsmz.de/genus/kinneretia |
| Kiritimatiella | https://lpsn.dsmz.de/genus/kiritimatiella |
| Kistimonas | https://lpsn.dsmz.de/genus/kistimonas |
| Kitasatoa | https://lpsn.dsmz.de/genus/kitasatoa |
| Kitasatospora | https://lpsn.dsmz.de/genus/kitasatospora |
| Kitasatosporia | https://lpsn.dsmz.de/genus/kitasatosporia |
| Klebsiella | https://lpsn.dsmz.de/genus/klebsiella |
| Klenkia | https://lpsn.dsmz.de/genus/klenkia |
| Klugiella | https://lpsn.dsmz.de/genus/klugiella |
| Kluyvera | https://lpsn.dsmz.de/genus/kluyvera |
| Knoellia | https://lpsn.dsmz.de/genus/knoellia |
| Kocuria | https://lpsn.dsmz.de/genus/kocuria |
| Kofleria | https://lpsn.dsmz.de/genus/kofleria |
| Koleobacter | https://lpsn.dsmz.de/genus/koleobacter |
| Komagataeibacter | https://lpsn.dsmz.de/genus/komagataeibacter |
| Kordia | https://lpsn.dsmz.de/genus/kordia |
| Kordiimonas | https://lpsn.dsmz.de/genus/kordiimonas |
| Koreibacter | https://lpsn.dsmz.de/genus/koreibacter |
| Kosakonia | https://lpsn.dsmz.de/genus/kosakonia |
| Koserella | https://lpsn.dsmz.de/genus/koserella |
| Kosmotoga | https://lpsn.dsmz.de/genus/kosmotoga |
| Kozakia | https://lpsn.dsmz.de/genus/kozakia |
| Krasilnikovia | https://lpsn.dsmz.de/genus/krasilnikovia |
| Krasilnikoviella | https://lpsn.dsmz.de/genus/krasilnikoviella |
| Kribbella | https://lpsn.dsmz.de/genus/kribbella |
| Kribbia | https://lpsn.dsmz.de/genus/kribbia |

| | |
|---|---|
| Kriegella | https://lpsn.dsmz.de/genus/kriegella |
| Krokinobacter | https://lpsn.dsmz.de/genus/krokinobacter |
| Kroppenstedtia | https://lpsn.dsmz.de/genus/kroppenstedtia |
| Ktedobacter | https://lpsn.dsmz.de/genus/ktedobacter |
| Ktedonobacter | https://lpsn.dsmz.de/genus/ktedonobacter |
| Ktedonospora | https://lpsn.dsmz.de/genus/ktedonospora |
| Ktedonosporobacter | https://lpsn.dsmz.de/genus/ktedonosporobacter |
| Kurthia | https://lpsn.dsmz.de/genus/kurthia |
| Kushneria | https://lpsn.dsmz.de/genus/kushneria |
| Kutzneria | https://lpsn.dsmz.de/genus/kutzneria |
| Kyrpidia | https://lpsn.dsmz.de/genus/kyrpidia |
| Kytococcus | https://lpsn.dsmz.de/genus/kytococcus |
| Labedaea | https://lpsn.dsmz.de/genus/labedaea |
| Labedella | https://lpsn.dsmz.de/genus/labedella |
| Labilibacter | https://lpsn.dsmz.de/genus/labilibacter |
| Labilibaculum | https://lpsn.dsmz.de/genus/labilibaculum |
| Labilithrix | https://lpsn.dsmz.de/genus/labilithrix |
| Labrenzia | https://lpsn.dsmz.de/genus/labrenzia |
| Labrys | https://lpsn.dsmz.de/genus/labrys |
| Laceyella | https://lpsn.dsmz.de/genus/laceyella |
| Lachnoanaerobaculum | https://lpsn.dsmz.de/genus/lachnoanaerobaculum |
| Lachnobacterium | https://lpsn.dsmz.de/genus/lachnobacterium |
| Lachnospira | https://lpsn.dsmz.de/genus/lachnospira |
| Lachnotalea | https://lpsn.dsmz.de/genus/lachnotalea |
| Lacibacter | https://lpsn.dsmz.de/genus/lacibacter |
| Lacibacterium | https://lpsn.dsmz.de/genus/lacibacterium |
| Lacihabitans | https://lpsn.dsmz.de/genus/lacihabitans |
| Lacimicrobium | https://lpsn.dsmz.de/genus/lacimicrobium |
| Lacimonas | https://lpsn.dsmz.de/genus/lacimonas |
| Lacinutrix | https://lpsn.dsmz.de/genus/lacinutrix |
| Lacipirellula | https://lpsn.dsmz.de/genus/lacipirellula |
| Lacisediminihabitans | https://lpsn.dsmz.de/genus/lacisediminihabitans |
| Lacisediminimonas | https://lpsn.dsmz.de/genus/lacisediminimonas |
| Lacrimispora | https://lpsn.dsmz.de/genus/lacrimispora |
| Lacticaseibacillus | https://lpsn.dsmz.de/genus/lacticaseibacillus |
| Lacticigenium | https://lpsn.dsmz.de/genus/lacticigenium |
| Lactiplantibacillus | https://lpsn.dsmz.de/genus/lactiplantibacillus |
| Lactivibrio | https://lpsn.dsmz.de/genus/lactivibrio |
| Lactobacillus | https://lpsn.dsmz.de/genus/lactobacillus |
| Lactococcus | https://lpsn.dsmz.de/genus/lactococcus |
| Lactonifactor | https://lpsn.dsmz.de/genus/lactonifactor |
| Lactosphaera | https://lpsn.dsmz.de/genus/lactosphaera |
| Lactovum | https://lpsn.dsmz.de/genus/lactovum |
| Laedolimicola | https://lpsn.dsmz.de/genus/laedolimicola |

| | |
|---|---|
| Lamprobacter | https://lpsn.dsmz.de/genus/lamprobacter |
| Lamprocystis | https://lpsn.dsmz.de/genus/lamprocystis |
| Lampropedia | https://lpsn.dsmz.de/genus/lampropedia |
| Lancefieldella | https://lpsn.dsmz.de/genus/lancefieldella |
| Lapidilactobacillus | https://lpsn.dsmz.de/genus/lapidilactobacillus |
| Lapillicoccus | https://lpsn.dsmz.de/genus/lapillicoccus |
| Laribacter | https://lpsn.dsmz.de/genus/laribacter |
| Larkinella | https://lpsn.dsmz.de/genus/larkinella |
| Larsenimonas | https://lpsn.dsmz.de/genus/larsenimonas |
| Latilactobacillus | https://lpsn.dsmz.de/genus/latilactobacillus |
| Lautropia | https://lpsn.dsmz.de/genus/lautropia |
| Lawsonella | https://lpsn.dsmz.de/genus/lawsonella |
| Lawsonia | https://lpsn.dsmz.de/genus/lawsonia |
| Lawsonibacter | https://lpsn.dsmz.de/genus/lawsonibacter |
| Leadbettera | https://lpsn.dsmz.de/genus/leadbettera |
| Leadbetterella | https://lpsn.dsmz.de/genus/leadbetterella |
| Lebetimonas | https://lpsn.dsmz.de/genus/lebetimonas |
| Lechevalieria | https://lpsn.dsmz.de/genus/lechevalieria |
| Leclercia | https://lpsn.dsmz.de/genus/leclercia |
| Lederbergia | https://lpsn.dsmz.de/genus/lederbergia |
| Leeia | https://lpsn.dsmz.de/genus/leeia |
| Leekyejoonella | https://lpsn.dsmz.de/genus/leekyejoonella |
| Leeuwenhoekiella | https://lpsn.dsmz.de/genus/leeuwenhoekiella |
| Legionella | https://lpsn.dsmz.de/genus/legionella |
| Leifsonia | https://lpsn.dsmz.de/genus/leifsonia |
| Leisingera | https://lpsn.dsmz.de/genus/leisingera |
| Lelliottia | https://lpsn.dsmz.de/genus/lelliottia |
| Leminorella | https://lpsn.dsmz.de/genus/leminorella |
| Lentibacillus | https://lpsn.dsmz.de/genus/lentibacillus |
| Lentibacter | https://lpsn.dsmz.de/genus/lentibacter |
| Lentihominibacter | https://lpsn.dsmz.de/genus/lentihominibacter |
| Lentilactobacillus | https://lpsn.dsmz.de/genus/lentilactobacillus |
| Lentilitoribacter | https://lpsn.dsmz.de/genus/lentilitoribacter |
| Lentimicrobium | https://lpsn.dsmz.de/genus/lentimicrobium |
| Lentisphaera | https://lpsn.dsmz.de/genus/lentisphaera |
| Lentzea | https://lpsn.dsmz.de/genus/lentzea |
| Leptobacterium | https://lpsn.dsmz.de/genus/leptobacterium |
| Leptogranulimonas | https://lpsn.dsmz.de/genus/leptogranulimonas |
| Leptolinea | https://lpsn.dsmz.de/genus/leptolinea |
| Leptolyngbya | https://lpsn.dsmz.de/genus/leptolyngbya |
| Leptonema | https://lpsn.dsmz.de/genus/leptonema |
| Leptospira | https://lpsn.dsmz.de/genus/leptospira |
| Leptospirillum | https://lpsn.dsmz.de/genus/leptospirillum |
| Leptothrix | https://lpsn.dsmz.de/genus/leptothrix |

| | |
|---|---|
| Leptotrichia | https://lpsn.dsmz.de/genus/leptotrichia |
| Leucobacter | https://lpsn.dsmz.de/genus/leucobacter |
| Leuconostoc | https://lpsn.dsmz.de/genus/leuconostoc |
| Leucothrix | https://lpsn.dsmz.de/genus/leucothrix |
| Levilactobacillus | https://lpsn.dsmz.de/genus/levilactobacillus |
| Levilinea | https://lpsn.dsmz.de/genus/levilinea |
| Levinea | https://lpsn.dsmz.de/genus/levinea |
| Lewinella | https://lpsn.dsmz.de/genus/lewinella |
| Libanicoccus | https://lpsn.dsmz.de/genus/libanicoccus |
| Liberibacter | https://lpsn.dsmz.de/genus/liberibacter |
| Lichenibacterium | https://lpsn.dsmz.de/genus/lichenibacterium |
| Lichenicoccus | https://lpsn.dsmz.de/genus/lichenicoccus |
| Lichenicola | https://lpsn.dsmz.de/genus/lichenicola |
| Lichenihabitans | https://lpsn.dsmz.de/genus/lichenihabitans |
| Ligaoa | https://lpsn.dsmz.de/genus/ligaoa |
| Ligilactobacillus | https://lpsn.dsmz.de/genus/ligilactobacillus |
| Lignipirellula | https://lpsn.dsmz.de/genus/lignipirellula |
| Lihuaxuella | https://lpsn.dsmz.de/genus/lihuaxuella |
| Limibacillus | https://lpsn.dsmz.de/genus/limibacillus |
| Limibacter | https://lpsn.dsmz.de/genus/limibacter |
| Limibaculum | https://lpsn.dsmz.de/genus/limibaculum |
| Limihaloglobus | https://lpsn.dsmz.de/genus/limihaloglobus |
| Limimaricola | https://lpsn.dsmz.de/genus/limimaricola |
| Limimonas | https://lpsn.dsmz.de/genus/limimonas |
| Limisalsivibrio | https://lpsn.dsmz.de/genus/limisalsivibrio |
| Limisphaera | https://lpsn.dsmz.de/genus/limisphaera |
| Limnobacter | https://lpsn.dsmz.de/genus/limnobacter |
| Limnobaculum | https://lpsn.dsmz.de/genus/limnobaculum |
| Limnochorda | https://lpsn.dsmz.de/genus/limnochorda |
| Limnoglobus | https://lpsn.dsmz.de/genus/limnoglobus |
| Limnohabitans | https://lpsn.dsmz.de/genus/limnohabitans |
| Limnovirga | https://lpsn.dsmz.de/genus/limnovirga |
| Limoniibacter | https://lpsn.dsmz.de/genus/limoniibacter |
| Limosilactobacillus | https://lpsn.dsmz.de/genus/limosilactobacillus |
| Lipingzhangella | https://lpsn.dsmz.de/genus/lipingzhangella |
| Liquorilactobacillus | https://lpsn.dsmz.de/genus/liquorilactobacillus |
| Lishizhenia | https://lpsn.dsmz.de/genus/lishizhenia |
| Listeria | https://lpsn.dsmz.de/genus/listeria |
| Listonella | https://lpsn.dsmz.de/genus/listonella |
| Litchfieldia | https://lpsn.dsmz.de/genus/litchfieldia |
| Litoreibacter | https://lpsn.dsmz.de/genus/litoreibacter |
| Litoribacillus | https://lpsn.dsmz.de/genus/litoribacillus |
| Litoribacter | https://lpsn.dsmz.de/genus/litoribacter |
| Litoribacterium | https://lpsn.dsmz.de/genus/litoribacterium |

| | |
|---|---|
| Litoribaculum | https://lpsn.dsmz.de/genus/litoribaculum |
| Litoribrevibacter | https://lpsn.dsmz.de/genus/litoribrevibacter |
| Litoricola | https://lpsn.dsmz.de/genus/litoricola |
| Litorihabitans | https://lpsn.dsmz.de/genus/litorihabitans |
| Litorilinea | https://lpsn.dsmz.de/genus/litorilinea |
| Litorilituus | https://lpsn.dsmz.de/genus/litorilituus |
| Litorimicrobium | https://lpsn.dsmz.de/genus/litorimicrobium |
| Litorimonas | https://lpsn.dsmz.de/genus/litorimonas |
| Litorisediminicola | https://lpsn.dsmz.de/genus/litorisediminicola |
| Litorisediminivivens | https://lpsn.dsmz.de/genus/litorisediminivivens |
| Litorivita | https://lpsn.dsmz.de/genus/litorivita |
| Litorivivens | https://lpsn.dsmz.de/genus/litorivivens |
| Loigolactobacillus | https://lpsn.dsmz.de/genus/loigolactobacillus |
| Loktanella | https://lpsn.dsmz.de/genus/loktanella |
| Lonepinella | https://lpsn.dsmz.de/genus/lonepinella |
| Longibacter | https://lpsn.dsmz.de/genus/longibacter |
| Longibaculum | https://lpsn.dsmz.de/genus/longibaculum |
| Longicatena | https://lpsn.dsmz.de/genus/longicatena |
| Longilinea | https://lpsn.dsmz.de/genus/longilinea |
| Longimicrobium | https://lpsn.dsmz.de/genus/longimicrobium |
| Longimonas | https://lpsn.dsmz.de/genus/longimonas |
| Longimycelium | https://lpsn.dsmz.de/genus/longimycelium |
| Longirhabdus | https://lpsn.dsmz.de/genus/longirhabdus |
| Longispora | https://lpsn.dsmz.de/genus/longispora |
| Longivirga | https://lpsn.dsmz.de/genus/longivirga |
| Lonsdalea | https://lpsn.dsmz.de/genus/lonsdalea |
| Lottiidibacillus | https://lpsn.dsmz.de/genus/lottiidibacillus |
| Lucibacterium | https://lpsn.dsmz.de/genus/lucibacterium |
| Lucifera | https://lpsn.dsmz.de/genus/lucifera |
| Luedemannella | https://lpsn.dsmz.de/genus/luedemannella |
| Lujinxingia | https://lpsn.dsmz.de/genus/lujinxingia |
| Luminiphilus | https://lpsn.dsmz.de/genus/luminiphilus |
| Lunatibacter | https://lpsn.dsmz.de/genus/lunatibacter |
| Lunatimonas | https://lpsn.dsmz.de/genus/lunatimonas |
| Luoshenia | https://lpsn.dsmz.de/genus/luoshenia |
| Lutaonella | https://lpsn.dsmz.de/genus/lutaonella |
| Luteibacter | https://lpsn.dsmz.de/genus/luteibacter |
| Luteibaculum | https://lpsn.dsmz.de/genus/luteibaculum |
| Luteimicrobium | https://lpsn.dsmz.de/genus/luteimicrobium |
| Luteimonas | https://lpsn.dsmz.de/genus/luteimonas |
| Luteipulveratus | https://lpsn.dsmz.de/genus/luteipulveratus |
| Luteirhabdus | https://lpsn.dsmz.de/genus/luteirhabdus |
| Luteitalea | https://lpsn.dsmz.de/genus/luteitalea |
| Luteithermobacter | https://lpsn.dsmz.de/genus/luteithermobacter |

| | |
|---|---|
| Luteivirga | https://lpsn.dsmz.de/genus/luteivirga |
| Luteococcus | https://lpsn.dsmz.de/genus/luteococcus |
| Luteolibacter | https://lpsn.dsmz.de/genus/luteolibacter |
| Luteovulum | https://lpsn.dsmz.de/genus/luteovulum |
| Lutibacter | https://lpsn.dsmz.de/genus/lutibacter |
| Lutibaculum | https://lpsn.dsmz.de/genus/lutibaculum |
| Lutimaribacter | https://lpsn.dsmz.de/genus/lutimaribacter |
| Lutimonas | https://lpsn.dsmz.de/genus/lutimonas |
| Lutispora | https://lpsn.dsmz.de/genus/lutispora |
| Lysinibacillus | https://lpsn.dsmz.de/genus/lysinibacillus |
| Lysinibacter | https://lpsn.dsmz.de/genus/lysinibacter |
| Lysinimicrobium | https://lpsn.dsmz.de/genus/lysinimicrobium |
| Lysinimonas | https://lpsn.dsmz.de/genus/lysinimonas |
| Lysobacter | https://lpsn.dsmz.de/genus/lysobacter |
| Lyticum | https://lpsn.dsmz.de/genus/lyticum |
| Mabikibacter | https://lpsn.dsmz.de/genus/mabikibacter |
| Macellibacteroides | https://lpsn.dsmz.de/genus/macellibacteroides |
| Macrococcus | https://lpsn.dsmz.de/genus/macrococcus |
| Macromonas | https://lpsn.dsmz.de/genus/macromonas |
| Mageeibacillus | https://lpsn.dsmz.de/genus/mageeibacillus |
| Magnetococcus | https://lpsn.dsmz.de/genus/magnetococcus |
| Magnetospira | https://lpsn.dsmz.de/genus/magnetospira |
| Magnetospirillum | https://lpsn.dsmz.de/genus/magnetospirillum |
| Magnetovibrio | https://lpsn.dsmz.de/genus/magnetovibrio |
| Mahella | https://lpsn.dsmz.de/genus/mahella |
| Maioricimonas | https://lpsn.dsmz.de/genus/maioricimonas |
| Malaciobacter | https://lpsn.dsmz.de/genus/malaciobacter |
| Malacobacter | https://lpsn.dsmz.de/genus/malacobacter |
| Malacoplasma | https://lpsn.dsmz.de/genus/malacoplasma |
| Maledivibacter | https://lpsn.dsmz.de/genus/maledivibacter |
| Malikia | https://lpsn.dsmz.de/genus/malikia |
| Maliponia | https://lpsn.dsmz.de/genus/maliponia |
| Malonomonas | https://lpsn.dsmz.de/genus/malonomonas |
| Mameliella | https://lpsn.dsmz.de/genus/mameliella |
| Mammaliicoccus | https://lpsn.dsmz.de/genus/mammaliicoccus |
| Mangrovactinospora | https://lpsn.dsmz.de/genus/mangrovactinospora |
| Mangrovibacter | https://lpsn.dsmz.de/genus/mangrovibacter |
| Mangrovibacterium | https://lpsn.dsmz.de/genus/mangrovibacterium |
| Mangrovicella | https://lpsn.dsmz.de/genus/mangrovicella |
| Mangrovicoccus | https://lpsn.dsmz.de/genus/mangrovicoccus |
| Mangroviflexus | https://lpsn.dsmz.de/genus/mangroviflexus |
| Mangrovihabitans | https://lpsn.dsmz.de/genus/mangrovihabitans |
| Mangrovimonas | https://lpsn.dsmz.de/genus/mangrovimonas |
| Mangrovitalea | https://lpsn.dsmz.de/genus/mangrovitalea |

| | |
|---|---|
| Mangrovivirga | https://lpsn.dsmz.de/genus/mangrovivirga |
| Mannheimia | https://lpsn.dsmz.de/genus/mannheimia |
| Margalitia | https://lpsn.dsmz.de/genus/margalitia |
| Maribacter | https://lpsn.dsmz.de/genus/maribacter |
| Maribaculum | https://lpsn.dsmz.de/genus/maribaculum |
| Maribellus | https://lpsn.dsmz.de/genus/maribellus |
| Maribius | https://lpsn.dsmz.de/genus/maribius |
| Maribrevibacterium | https://lpsn.dsmz.de/genus/maribrevibacterium |
| Maricaulis | https://lpsn.dsmz.de/genus/maricaulis |
| Marichromatium | https://lpsn.dsmz.de/genus/marichromatium |
| Maricurvus | https://lpsn.dsmz.de/genus/maricurvus |
| Maridesulfovibrio | https://lpsn.dsmz.de/genus/maridesulfovibrio |
| Marihabitans | https://lpsn.dsmz.de/genus/marihabitans |
| Marimicrobium | https://lpsn.dsmz.de/genus/marimicrobium |
| Marimonas | https://lpsn.dsmz.de/genus/marimonas |
| Marinactinospora | https://lpsn.dsmz.de/genus/marinactinospora |
| Marinagarivorans | https://lpsn.dsmz.de/genus/marinagarivorans |
| Marinibacillus | https://lpsn.dsmz.de/genus/marinibacillus |
| Marinibacterium | https://lpsn.dsmz.de/genus/marinibacterium |
| Marinibactrum | https://lpsn.dsmz.de/genus/marinibactrum |
| Marinibaculum | https://lpsn.dsmz.de/genus/marinibaculum |
| Mariniblastus | https://lpsn.dsmz.de/genus/mariniblastus |
| Marinicauda | https://lpsn.dsmz.de/genus/marinicauda |
| Marinicaulis | https://lpsn.dsmz.de/genus/marinicaulis |
| Marinicella | https://lpsn.dsmz.de/genus/marinicella |
| Marinicola | https://lpsn.dsmz.de/genus/marinicola |
| Marinicrinis | https://lpsn.dsmz.de/genus/marinicrinis |
| Marinifilum | https://lpsn.dsmz.de/genus/marinifilum |
| Mariniflexile | https://lpsn.dsmz.de/genus/mariniflexile |
| Marinilabilia | https://lpsn.dsmz.de/genus/marinilabilia |
| Marinilactibacillus | https://lpsn.dsmz.de/genus/marinilactibacillus |
| Mariniluteicoccus | https://lpsn.dsmz.de/genus/mariniluteicoccus |
| Marinimicrobium | https://lpsn.dsmz.de/genus/marinimicrobium |
| Marininema | https://lpsn.dsmz.de/genus/marininema |
| Mariniphaga | https://lpsn.dsmz.de/genus/mariniphaga |
| Mariniplasma | https://lpsn.dsmz.de/genus/mariniplasma |
| Mariniradius | https://lpsn.dsmz.de/genus/mariniradius |
| Marinirhabdus | https://lpsn.dsmz.de/genus/marinirhabdus |
| Marinisporobacter | https://lpsn.dsmz.de/genus/marinisporobacter |
| Marinisubtilis | https://lpsn.dsmz.de/genus/marinisubtilis |
| Marinitenerispora | https://lpsn.dsmz.de/genus/marinitenerispora |
| Marinithermofilum | https://lpsn.dsmz.de/genus/marinithermofilum |
| Marinithermus | https://lpsn.dsmz.de/genus/marinithermus |
| Marinitoga | https://lpsn.dsmz.de/genus/marinitoga |

| | |
|---|---|
| Marinivirga | https://lpsn.dsmz.de/genus/marinivirga |
| Marinobacter | https://lpsn.dsmz.de/genus/marinobacter |
| Marinobacterium | https://lpsn.dsmz.de/genus/marinobacterium |
| Marinococcus | https://lpsn.dsmz.de/genus/marinococcus |
| Marinomonas | https://lpsn.dsmz.de/genus/marinomonas |
| Marinoscillum | https://lpsn.dsmz.de/genus/marinoscillum |
| Marinospirillum | https://lpsn.dsmz.de/genus/marinospirillum |
| Marinovum | https://lpsn.dsmz.de/genus/marinovum |
| Mariprofundus | https://lpsn.dsmz.de/genus/mariprofundus |
| Maripseudobacter | https://lpsn.dsmz.de/genus/maripseudobacter |
| Marisediminicola | https://lpsn.dsmz.de/genus/marisediminicola |
| Marisediminitalea | https://lpsn.dsmz.de/genus/marisediminitalea |
| Marispirillum | https://lpsn.dsmz.de/genus/marispirillum |
| Marispirochaeta | https://lpsn.dsmz.de/genus/marispirochaeta |
| Maritalea | https://lpsn.dsmz.de/genus/maritalea |
| Maritimibacter | https://lpsn.dsmz.de/genus/maritimibacter |
| Maritimimonas | https://lpsn.dsmz.de/genus/maritimimonas |
| Marivibrio | https://lpsn.dsmz.de/genus/marivibrio |
| Marivirga | https://lpsn.dsmz.de/genus/marivirga |
| Marivita | https://lpsn.dsmz.de/genus/marivita |
| Marivivens | https://lpsn.dsmz.de/genus/marivivens |
| Marixanthomonas | https://lpsn.dsmz.de/genus/marixanthomonas |
| Marmoricola | https://lpsn.dsmz.de/genus/marmoricola |
| Marortus | https://lpsn.dsmz.de/genus/marortus |
| Martelella | https://lpsn.dsmz.de/genus/martelella |
| Marvinbryantia | https://lpsn.dsmz.de/genus/marvinbryantia |
| Massilia | https://lpsn.dsmz.de/genus/massilia |
| Massiliimalia | https://lpsn.dsmz.de/genus/massiliimalia |
| Mechercharimyces | https://lpsn.dsmz.de/genus/mechercharimyces |
| Mediterraneibacter | https://lpsn.dsmz.de/genus/mediterraneibacter |
| Megalodesulfovibrio | https://lpsn.dsmz.de/genus/megalodesulfovibrio |
| Megamonas | https://lpsn.dsmz.de/genus/megamonas |
| Meganema | https://lpsn.dsmz.de/genus/meganema |
| Megasphaera | https://lpsn.dsmz.de/genus/megasphaera |
| Meinhardsimonia | https://lpsn.dsmz.de/genus/meinhardsimonia |
| Meiothermus | https://lpsn.dsmz.de/genus/meiothermus |
| Melaminivora | https://lpsn.dsmz.de/genus/melaminivora |
| Melghiribacillus | https://lpsn.dsmz.de/genus/melghiribacillus |
| Melghirimyces | https://lpsn.dsmz.de/genus/melghirimyces |
| Melioribacter | https://lpsn.dsmz.de/genus/melioribacter |
| Melisococcus | https://lpsn.dsmz.de/genus/melisococcus |
| Melissococcus | https://lpsn.dsmz.de/genus/melissococcus |
| Melitea | https://lpsn.dsmz.de/genus/melitea |
| Melittangium | https://lpsn.dsmz.de/genus/melittangium |

| | |
|---|---|
| Membranicola | https://lpsn.dsmz.de/genus/membranicola |
| Membranihabitans | https://lpsn.dsmz.de/genus/membranihabitans |
| Meniscus | https://lpsn.dsmz.de/genus/meniscus |
| Merdimonas | https://lpsn.dsmz.de/genus/merdimonas |
| Meridianimaribacter | https://lpsn.dsmz.de/genus/meridianimaribacter |
| Meridianimarinicoccus | https://lpsn.dsmz.de/genus/meridianimarinicoccus |
| Mesoaciditoga | https://lpsn.dsmz.de/genus/mesoaciditoga |
| Mesobacillus | https://lpsn.dsmz.de/genus/mesobacillus |
| Mesobacterium | https://lpsn.dsmz.de/genus/mesobacterium |
| Mesobaculum | https://lpsn.dsmz.de/genus/mesobaculum |
| Mesocricetibacter | https://lpsn.dsmz.de/genus/mesocricetibacter |
| Mesoflavibacter | https://lpsn.dsmz.de/genus/mesoflavibacter |
| Mesohalobacter | https://lpsn.dsmz.de/genus/mesohalobacter |
| Mesomycoplasma | https://lpsn.dsmz.de/genus/mesomycoplasma |
| Mesonia | https://lpsn.dsmz.de/genus/mesonia |
| Mesophilobacter | https://lpsn.dsmz.de/genus/mesophilobacter |
| Mesoplasma | https://lpsn.dsmz.de/genus/mesoplasma |
| Mesorhizobium | https://lpsn.dsmz.de/genus/mesorhizobium |
| Mesosutterella | https://lpsn.dsmz.de/genus/mesosutterella |
| Mesotoga | https://lpsn.dsmz.de/genus/mesotoga |
| Metabacillus | https://lpsn.dsmz.de/genus/metabacillus |
| Metakosakonia | https://lpsn.dsmz.de/genus/metakosakonia |
| Metallibacterium | https://lpsn.dsmz.de/genus/metallibacterium |
| Metallosphaera | https://lpsn.dsmz.de/genus/metallosphaera |
| Metalysinibacillus | https://lpsn.dsmz.de/genus/metalysinibacillus |
| Metamycoplasma | https://lpsn.dsmz.de/genus/metamycoplasma |
| Metaplanococcus | https://lpsn.dsmz.de/genus/metaplanococcus |
| Metascardovia | https://lpsn.dsmz.de/genus/metascardovia |
| Metasolibacillus | https://lpsn.dsmz.de/genus/metasolibacillus |
| Methanimicrococcus | https://lpsn.dsmz.de/genus/methanimicrococcus |
| Methanobacterium | https://lpsn.dsmz.de/genus/methanobacterium |
| Methanobrevibacter | https://lpsn.dsmz.de/genus/methanobrevibacter |
| Methanocalculus | https://lpsn.dsmz.de/genus/methanocalculus |
| Methanocaldococcus | https://lpsn.dsmz.de/genus/methanocaldococcus |
| Methanocella | https://lpsn.dsmz.de/genus/methanocella |
| Methanococcoides | https://lpsn.dsmz.de/genus/methanococcoides |
| Methanococcus | https://lpsn.dsmz.de/genus/methanococcus |
| Methanocorpusculum | https://lpsn.dsmz.de/genus/methanocorpusculum |
| Methanoculleus | https://lpsn.dsmz.de/genus/methanoculleus |
| Methanofervidicoccus | https://lpsn.dsmz.de/genus/methanofervidicoccus |
| Methanofollis | https://lpsn.dsmz.de/genus/methanofollis |
| Methanogenium | https://lpsn.dsmz.de/genus/methanogenium |
| Methanohalobium | https://lpsn.dsmz.de/genus/methanohalobium |
| Methanohalophilus | https://lpsn.dsmz.de/genus/methanohalophilus |

| | |
|---|---|
| Methanolacinia | https://lpsn.dsmz.de/genus/methanolacinia |
| Methanolinea | https://lpsn.dsmz.de/genus/methanolinea |
| Methanolobus | https://lpsn.dsmz.de/genus/methanolobus |
| Methanomassiliicoccus | https://lpsn.dsmz.de/genus/methanomassiliicoccus |
| Methanomethylovorans | https://lpsn.dsmz.de/genus/methanomethylovorans |
| Methanomicrobium | https://lpsn.dsmz.de/genus/methanomicrobium |
| Methanomicrococcus | https://lpsn.dsmz.de/genus/methanomicrococcus |
| Methanonatronarchaeum | https://lpsn.dsmz.de/genus/methanonatronarchaeum |
| Methanoplanus | https://lpsn.dsmz.de/genus/methanoplanus |
| Methanopyrus | https://lpsn.dsmz.de/genus/methanopyrus |
| Methanoregula | https://lpsn.dsmz.de/genus/methanoregula |
| Methanosaeta | https://lpsn.dsmz.de/genus/methanosaeta |
| Methanosalsum | https://lpsn.dsmz.de/genus/methanosalsum |
| Methanosarcina | https://lpsn.dsmz.de/genus/methanosarcina |
| Methanosphaera | https://lpsn.dsmz.de/genus/methanosphaera |
| Methanosphaerula | https://lpsn.dsmz.de/genus/methanosphaerula |
| Methanospirillum | https://lpsn.dsmz.de/genus/methanospirillum |
| Methanothermobacter | https://lpsn.dsmz.de/genus/methanothermobacter |
| Methanothermococcus | https://lpsn.dsmz.de/genus/methanothermococcus |
| Methanothermus | https://lpsn.dsmz.de/genus/methanothermus |
| Methanothrix | https://lpsn.dsmz.de/genus/methanothrix |
| Methanotorris | https://lpsn.dsmz.de/genus/methanotorris |
| Methermicoccus | https://lpsn.dsmz.de/genus/methermicoccus |
| Methylarcula | https://lpsn.dsmz.de/genus/methylarcula |
| Methylibium | https://lpsn.dsmz.de/genus/methylibium |
| Methylicorpusculum | https://lpsn.dsmz.de/genus/methylicorpusculum |
| Methylobacillus | https://lpsn.dsmz.de/genus/methylobacillus |
| Methylobacter | https://lpsn.dsmz.de/genus/methylobacter |
| Methylobacterium | https://lpsn.dsmz.de/genus/methylobacterium |
| Methylobrevis | https://lpsn.dsmz.de/genus/methylobrevis |
| Methylocaldum | https://lpsn.dsmz.de/genus/methylocaldum |
| Methylocapsa | https://lpsn.dsmz.de/genus/methylocapsa |
| Methyloceanibacter | https://lpsn.dsmz.de/genus/methyloceanibacter |
| Methylocella | https://lpsn.dsmz.de/genus/methylocella |
| Methylococcus | https://lpsn.dsmz.de/genus/methylococcus |
| Methylocucumis | https://lpsn.dsmz.de/genus/methylocucumis |
| Methylocystis | https://lpsn.dsmz.de/genus/methylocystis |
| Methyloferula | https://lpsn.dsmz.de/genus/methyloferula |
| Methylogaea | https://lpsn.dsmz.de/genus/methylogaea |
| Methyloglobulus | https://lpsn.dsmz.de/genus/methyloglobulus |
| Methylohalobius | https://lpsn.dsmz.de/genus/methylohalobius |
| Methylohalomonas | https://lpsn.dsmz.de/genus/methylohalomonas |
| Methyloligella | https://lpsn.dsmz.de/genus/methyloligella |
| Methylomagnum | https://lpsn.dsmz.de/genus/methylomagnum |

| | |
|---|---|
| Methylomarinovum | https://lpsn.dsmz.de/genus/methylomarinovum |
| Methylomarinum | https://lpsn.dsmz.de/genus/methylomarinum |
| Methylomicrobium | https://lpsn.dsmz.de/genus/methylomicrobium |
| Methylomonas | https://lpsn.dsmz.de/genus/methylomonas |
| Methylomusa | https://lpsn.dsmz.de/genus/methylomusa |
| Methylonatrum | https://lpsn.dsmz.de/genus/methylonatrum |
| Methyloparacoccus | https://lpsn.dsmz.de/genus/methyloparacoccus |
| Methylophaga | https://lpsn.dsmz.de/genus/methylophaga |
| Methylophilus | https://lpsn.dsmz.de/genus/methylophilus |
| Methylopila | https://lpsn.dsmz.de/genus/methylopila |
| Methyloprofundus | https://lpsn.dsmz.de/genus/methyloprofundus |
| Methyloradius | https://lpsn.dsmz.de/genus/methyloradius |
| Methylorhabdus | https://lpsn.dsmz.de/genus/methylorhabdus |
| Methylorosula | https://lpsn.dsmz.de/genus/methylorosula |
| Methylorubrum | https://lpsn.dsmz.de/genus/methylorubrum |
| Methylosarcina | https://lpsn.dsmz.de/genus/methylosarcina |
| Methylosinus | https://lpsn.dsmz.de/genus/methylosinus |
| Methylosoma | https://lpsn.dsmz.de/genus/methylosoma |
| Methylosphaera | https://lpsn.dsmz.de/genus/methylosphaera |
| Methylotenera | https://lpsn.dsmz.de/genus/methylotenera |
| Methyloterricola | https://lpsn.dsmz.de/genus/methyloterricola |
| Methyloterrigena | https://lpsn.dsmz.de/genus/methyloterrigena |
| Methylothermus | https://lpsn.dsmz.de/genus/methylothermus |
| Methylotuvimicrobium | https://lpsn.dsmz.de/genus/methylotuvimicrobium |
| Methyloversatilis | https://lpsn.dsmz.de/genus/methyloversatilis |
| Methylovirgula | https://lpsn.dsmz.de/genus/methylovirgula |
| Methylovorus | https://lpsn.dsmz.de/genus/methylovorus |
| Methylovulum | https://lpsn.dsmz.de/genus/methylovulum |
| Micavibrio | https://lpsn.dsmz.de/genus/micavibrio |
| Microaerobacter | https://lpsn.dsmz.de/genus/microaerobacter |
| Microbacter | https://lpsn.dsmz.de/genus/microbacter |
| Microbacterium | https://lpsn.dsmz.de/genus/microbacterium |
| Microbaculum | https://lpsn.dsmz.de/genus/microbaculum |
| Microbispora | https://lpsn.dsmz.de/genus/microbispora |
| Microbulbifer | https://lpsn.dsmz.de/genus/microbulbifer |
| Microcella | https://lpsn.dsmz.de/genus/microcella |
| Micrococcoides | https://lpsn.dsmz.de/genus/micrococcoides |
| Micrococcus | https://lpsn.dsmz.de/genus/micrococcus |
| Microcyclus | https://lpsn.dsmz.de/genus/microcyclus |
| Microellobosporia | https://lpsn.dsmz.de/genus/microellobosporia |
| Microlunatus | https://lpsn.dsmz.de/genus/microlunatus |
| Micromonas | https://lpsn.dsmz.de/genus/micromonas |
| Micromonospora | https://lpsn.dsmz.de/genus/micromonospora |
| Micropepsis | https://lpsn.dsmz.de/genus/micropepsis |

| | |
|---|---|
| Micropolyspora | https://lpsn.dsmz.de/genus/micropolyspora |
| Micropruina | https://lpsn.dsmz.de/genus/micropruina |
| Microscilla | https://lpsn.dsmz.de/genus/microscilla |
| Microsphaera | https://lpsn.dsmz.de/genus/microsphaera |
| Microterricola | https://lpsn.dsmz.de/genus/microterricola |
| Microtetraspora | https://lpsn.dsmz.de/genus/microtetraspora |
| Microvenator | https://lpsn.dsmz.de/genus/microvenator |
| Microvirga | https://lpsn.dsmz.de/genus/microvirga |
| Microvirgula | https://lpsn.dsmz.de/genus/microvirgula |
| Millisia | https://lpsn.dsmz.de/genus/millisia |
| Miltoncostaea | https://lpsn.dsmz.de/genus/miltoncostaea |
| Minicystis | https://lpsn.dsmz.de/genus/minicystis |
| Miniimonas | https://lpsn.dsmz.de/genus/miniimonas |
| Miniphocaeibacter | https://lpsn.dsmz.de/genus/miniphocaeibacter |
| Miniphocibacter | https://lpsn.dsmz.de/genus/miniphocibacter-1 |
| Minwuia | https://lpsn.dsmz.de/genus/minwuia |
| Mitsuaria | https://lpsn.dsmz.de/genus/mitsuaria |
| Mitsuokella | https://lpsn.dsmz.de/genus/mitsuokella |
| Mixta | https://lpsn.dsmz.de/genus/mixta |
| Mizugakiibacter | https://lpsn.dsmz.de/genus/mizugakiibacter |
| Mobilicoccus | https://lpsn.dsmz.de/genus/mobilicoccus |
| Mobilisporobacter | https://lpsn.dsmz.de/genus/mobilisporobacter |
| Mobilitalea | https://lpsn.dsmz.de/genus/mobilitalea |
| Mobiluncus | https://lpsn.dsmz.de/genus/mobiluncus |
| Modestobacter | https://lpsn.dsmz.de/genus/modestobacter |
| Modicisalibacter | https://lpsn.dsmz.de/genus/modicisalibacter |
| Moellerella | https://lpsn.dsmz.de/genus/moellerella |
| Mogibacterium | https://lpsn.dsmz.de/genus/mogibacterium |
| Moheibacter | https://lpsn.dsmz.de/genus/moheibacter |
| Monaibacterium | https://lpsn.dsmz.de/genus/monaibacterium |
| Monashia | https://lpsn.dsmz.de/genus/monashia |
| Mongoliibacter | https://lpsn.dsmz.de/genus/mongoliibacter |
| Mongoliicoccus | https://lpsn.dsmz.de/genus/mongoliicoccus |
| Mongoliimonas | https://lpsn.dsmz.de/genus/mongoliimonas |
| Mongoliitalea | https://lpsn.dsmz.de/genus/mongoliitalea |
| Monoglobus | https://lpsn.dsmz.de/genus/monoglobus |
| Mooreia | https://lpsn.dsmz.de/genus/mooreia |
| Moorella | https://lpsn.dsmz.de/genus/moorella |
| Moraxella | https://lpsn.dsmz.de/genus/moraxella |
| Morganella | https://lpsn.dsmz.de/genus/morganella |
| Moritella | https://lpsn.dsmz.de/genus/moritella |
| Morococcus | https://lpsn.dsmz.de/genus/morococcus |
| Moryella | https://lpsn.dsmz.de/genus/moryella |
| Motilibacter | https://lpsn.dsmz.de/genus/motilibacter |

| | |
|---|---|
| Motilimonas | https://lpsn.dsmz.de/genus/motilimonas |
| Motiliproteus | https://lpsn.dsmz.de/genus/motiliproteus |
| Mucibacter | https://lpsn.dsmz.de/genus/mucibacter |
| Mucilaginibacter | https://lpsn.dsmz.de/genus/mucilaginibacter |
| Mucinivorans | https://lpsn.dsmz.de/genus/mucinivorans |
| Mucisphaera | https://lpsn.dsmz.de/genus/mucisphaera |
| Mucispirillum | https://lpsn.dsmz.de/genus/mucispirillum |
| Mumia | https://lpsn.dsmz.de/genus/mumia |
| Murdochiella | https://lpsn.dsmz.de/genus/murdochiella |
| Muribacter | https://lpsn.dsmz.de/genus/muribacter |
| Muribaculum | https://lpsn.dsmz.de/genus/muribaculum |
| Muricauda | https://lpsn.dsmz.de/genus/muricauda |
| Muricoccus | https://lpsn.dsmz.de/genus/muricoccus |
| Muricomes | https://lpsn.dsmz.de/genus/muricomes |
| Muricoprocola | https://lpsn.dsmz.de/genus/muricoprocola |
| Muriicola | https://lpsn.dsmz.de/genus/muriicola-1 |
| Muriicola | https://lpsn.dsmz.de/genus/muriicola |
| Muriiphilus | https://lpsn.dsmz.de/genus/muriiphilus |
| Murimonas | https://lpsn.dsmz.de/genus/murimonas |
| Murinocardiopsis | https://lpsn.dsmz.de/genus/murinocardiopsis |
| Muriventricola | https://lpsn.dsmz.de/genus/muriventricola |
| Musicola | https://lpsn.dsmz.de/genus/musicola |
| Myceligenerans | https://lpsn.dsmz.de/genus/myceligenerans |
| Mycetocola | https://lpsn.dsmz.de/genus/mycetocola |
| Mycetohabitans | https://lpsn.dsmz.de/genus/mycetohabitans |
| Mycoavidus | https://lpsn.dsmz.de/genus/mycoavidus |
| Mycobacterium | https://lpsn.dsmz.de/genus/mycobacterium |
| Mycobacteroides | https://lpsn.dsmz.de/genus/mycobacteroides |
| Mycolicibacillus | https://lpsn.dsmz.de/genus/mycolicibacillus |
| Mycolicibacter | https://lpsn.dsmz.de/genus/mycolicibacter |
| Mycolicibacterium | https://lpsn.dsmz.de/genus/mycolicibacterium |
| Mycoplana | https://lpsn.dsmz.de/genus/mycoplana |
| Mycoplasma | https://lpsn.dsmz.de/genus/mycoplasma |
| Mycoplasmoides | https://lpsn.dsmz.de/genus/mycoplasmoides |
| Mycoplasmopsis | https://lpsn.dsmz.de/genus/mycoplasmopsis |
| Myroides | https://lpsn.dsmz.de/genus/myroides |
| Myxococcus | https://lpsn.dsmz.de/genus/myxococcus |
| Mzabimyces | https://lpsn.dsmz.de/genus/mzabimyces |
| Naasia | https://lpsn.dsmz.de/genus/naasia |
| Nafulsella | https://lpsn.dsmz.de/genus/nafulsella |
| Nakamurella | https://lpsn.dsmz.de/genus/nakamurella |
| Namhaeicola | https://lpsn.dsmz.de/genus/namhaeicola |
| Nanchangia | https://lpsn.dsmz.de/genus/nanchangia |
| Nannocystis | https://lpsn.dsmz.de/genus/nannocystis |

| | |
|---|---|
| Nanobdella | https://lpsn.dsmz.de/genus/nanobdella |
| Natranaerobaculum | https://lpsn.dsmz.de/genus/natranaerobaculum |
| Natranaerobius | https://lpsn.dsmz.de/genus/natranaerobius |
| Natranaerofaba | https://lpsn.dsmz.de/genus/natranaerofaba |
| Natranaerovirga | https://lpsn.dsmz.de/genus/natranaerovirga |
| Natrarchaeobaculum | https://lpsn.dsmz.de/genus/natrarchaeobaculum |
| Natrarchaeobius | https://lpsn.dsmz.de/genus/natrarchaeobius |
| Natrialba | https://lpsn.dsmz.de/genus/natrialba |
| Natribacillus | https://lpsn.dsmz.de/genus/natribacillus |
| Natribaculum | https://lpsn.dsmz.de/genus/natribaculum |
| Natrinema | https://lpsn.dsmz.de/genus/natrinema |
| Natroniella | https://lpsn.dsmz.de/genus/natroniella |
| Natronincola | https://lpsn.dsmz.de/genus/natronincola |
| Natronoarchaeum | https://lpsn.dsmz.de/genus/natronoarchaeum |
| Natronobacillus | https://lpsn.dsmz.de/genus/natronobacillus |
| Natronobacterium | https://lpsn.dsmz.de/genus/natronobacterium |
| Natronobiforma | https://lpsn.dsmz.de/genus/natronobiforma |
| Natronocella | https://lpsn.dsmz.de/genus/natronocella |
| Natronococcus | https://lpsn.dsmz.de/genus/natronococcus |
| Natronoflexus | https://lpsn.dsmz.de/genus/natronoflexus |
| Natronoglycomyces | https://lpsn.dsmz.de/genus/natronoglycomyces |
| Natronoincola | https://lpsn.dsmz.de/genus/natronoincola |
| Natronolimnobius | https://lpsn.dsmz.de/genus/natronolimnobius |
| Natronolimnohabitans | https://lpsn.dsmz.de/genus/natronolimnohabitans |
| Natronomonas | https://lpsn.dsmz.de/genus/natronomonas |
| Natronorubrum | https://lpsn.dsmz.de/genus/natronorubrum |
| Natronospira | https://lpsn.dsmz.de/genus/natronospira |
| Natronospirillum | https://lpsn.dsmz.de/genus/natronospirillum |
| Natronotalea | https://lpsn.dsmz.de/genus/natronotalea |
| Natronovirga | https://lpsn.dsmz.de/genus/natronovirga |
| Naumannella | https://lpsn.dsmz.de/genus/naumannella |
| Nautella | https://lpsn.dsmz.de/genus/nautella |
| Nautilia | https://lpsn.dsmz.de/genus/nautilia |
| Naxibacter | https://lpsn.dsmz.de/genus/naxibacter |
| Necropsobacter | https://lpsn.dsmz.de/genus/necropsobacter |
| Negadavirga | https://lpsn.dsmz.de/genus/negadavirga |
| Negativibacillus | https://lpsn.dsmz.de/genus/negativibacillus |
| Negativicoccus | https://lpsn.dsmz.de/genus/negativicoccus |
| Neiella | https://lpsn.dsmz.de/genus/neiella |
| Neisseria | https://lpsn.dsmz.de/genus/neisseria |
| Nemorella | https://lpsn.dsmz.de/genus/nemorella |
| Nemorincola | https://lpsn.dsmz.de/genus/nemorincola |
| Neoasaia | https://lpsn.dsmz.de/genus/neoasaia |
| Neobacillus | https://lpsn.dsmz.de/genus/neobacillus |

| | |
|---|---|
| Neokomagataea | https://lpsn.dsmz.de/genus/neokomagataea |
| Neolewinella | https://lpsn.dsmz.de/genus/neolewinella |
| Neomegalonema | https://lpsn.dsmz.de/genus/neomegalonema |
| Neomicrococcus | https://lpsn.dsmz.de/genus/neomicrococcus |
| Neopseudomonas | https://lpsn.dsmz.de/genus/neopseudomonas |
| Neorhizobium | https://lpsn.dsmz.de/genus/neorhizobium |
| Neorickettsia | https://lpsn.dsmz.de/genus/neorickettsia |
| Neoroseomonas | https://lpsn.dsmz.de/genus/neoroseomonas |
| Neoscardovia | https://lpsn.dsmz.de/genus/neoscardovia |
| Neotabrizicola | https://lpsn.dsmz.de/genus/neotabrizicola |
| Neptunicella | https://lpsn.dsmz.de/genus/neptunicella |
| Neptunicoccus | https://lpsn.dsmz.de/genus/neptunicoccus |
| Neptuniibacter | https://lpsn.dsmz.de/genus/neptuniibacter |
| Neptunitalea | https://lpsn.dsmz.de/genus/neptunitalea |
| Neptunomonas | https://lpsn.dsmz.de/genus/neptunomonas |
| Nereida | https://lpsn.dsmz.de/genus/nereida |
| Nesiotobacter | https://lpsn.dsmz.de/genus/nesiotobacter |
| Nesterenkonia | https://lpsn.dsmz.de/genus/nesterenkonia |
| Nevskia | https://lpsn.dsmz.de/genus/nevskia |
| Nguyenibacter | https://lpsn.dsmz.de/genus/nguyenibacter |
| Niabella | https://lpsn.dsmz.de/genus/niabella |
| Niallia | https://lpsn.dsmz.de/genus/niallia |
| Niastella | https://lpsn.dsmz.de/genus/niastella |
| Nibrella | https://lpsn.dsmz.de/genus/nibrella |
| Nibribacter | https://lpsn.dsmz.de/genus/nibribacter |
| Nibricoccus | https://lpsn.dsmz.de/genus/nibricoccus |
| Nicoletella | https://lpsn.dsmz.de/genus/nicoletella |
| Nicolia | https://lpsn.dsmz.de/genus/nicolia |
| Ningiella | https://lpsn.dsmz.de/genus/ningiella |
| Nioella | https://lpsn.dsmz.de/genus/nioella |
| Nisaea | https://lpsn.dsmz.de/genus/nisaea |
| Nitratidesulfovibrio | https://lpsn.dsmz.de/genus/nitratidesulfovibrio |
| Nitratifractor | https://lpsn.dsmz.de/genus/nitratifractor |
| Nitratireductor | https://lpsn.dsmz.de/genus/nitratireductor |
| Nitratiruptor | https://lpsn.dsmz.de/genus/nitratiruptor |
| Nitriliruptor | https://lpsn.dsmz.de/genus/nitriliruptor |
| Nitrincola | https://lpsn.dsmz.de/genus/nitrincola |
| Nitritalea | https://lpsn.dsmz.de/genus/nitritalea |
| Nitrobacter | https://lpsn.dsmz.de/genus/nitrobacter |
| Nitrococcus | https://lpsn.dsmz.de/genus/nitrococcus |
| Nitrogeniibacter | https://lpsn.dsmz.de/genus/nitrogeniibacter |
| Nitrolancea | https://lpsn.dsmz.de/genus/nitrolancea |
| Nitrosarchaeum | https://lpsn.dsmz.de/genus/nitrosarchaeum |
| Nitrosococcus | https://lpsn.dsmz.de/genus/nitrosococcus |

| | |
|---|---|
| Nitrosolobus | https://lpsn.dsmz.de/genus/nitrosolobus |
| Nitrosomonas | https://lpsn.dsmz.de/genus/nitrosomonas |
| Nitrosophilus | https://lpsn.dsmz.de/genus/nitrosophilus |
| Nitrosopumilus | https://lpsn.dsmz.de/genus/nitrosopumilus |
| Nitrososphaera | https://lpsn.dsmz.de/genus/nitrososphaera |
| Nitrosospira | https://lpsn.dsmz.de/genus/nitrosospira |
| Nitrospina | https://lpsn.dsmz.de/genus/nitrospina |
| Nitrospira | https://lpsn.dsmz.de/genus/nitrospira |
| Nitrospirillum | https://lpsn.dsmz.de/genus/nitrospirillum |
| Niveibacterium | https://lpsn.dsmz.de/genus/niveibacterium |
| Niveispirillum | https://lpsn.dsmz.de/genus/niveispirillum |
| Niveitalea | https://lpsn.dsmz.de/genus/niveitalea |
| Nocardia | https://lpsn.dsmz.de/genus/nocardia |
| Nocardioides | https://lpsn.dsmz.de/genus/nocardioides |
| Nocardiopsis | https://lpsn.dsmz.de/genus/nocardiopsis |
| Nonlabens | https://lpsn.dsmz.de/genus/nonlabens |
| Nonomuraea | https://lpsn.dsmz.de/genus/nonomuraea |
| Nonomuria | https://lpsn.dsmz.de/genus/nonomuria |
| Nosocomiicoccus | https://lpsn.dsmz.de/genus/nosocomiicoccus |
| Notoacmeibacter | https://lpsn.dsmz.de/genus/notoacmeibacter |
| Novacetimonas | https://lpsn.dsmz.de/genus/novacetimonas |
| Novibacillus | https://lpsn.dsmz.de/genus/novibacillus |
| Noviherbaspirillum | https://lpsn.dsmz.de/genus/noviherbaspirillum |
| Novimethylophilus | https://lpsn.dsmz.de/genus/novimethylophilus |
| Novipirellula | https://lpsn.dsmz.de/genus/novipirellula |
| Novispirillum | https://lpsn.dsmz.de/genus/novispirillum |
| Novisyntrophococcus | https://lpsn.dsmz.de/genus/novisyntrophococcus |
| Novosphingobium | https://lpsn.dsmz.de/genus/novosphingobium |
| Novosphingopyxis | https://lpsn.dsmz.de/genus/novosphingopyxis |
| Nubsella | https://lpsn.dsmz.de/genus/nubsella |
| Nucisporomicrobium | https://lpsn.dsmz.de/genus/nucisporomicrobium |
| Obesumbacterium | https://lpsn.dsmz.de/genus/obesumbacterium |
| Oblitimonas | https://lpsn.dsmz.de/genus/oblitimonas |
| Occallatibacter | https://lpsn.dsmz.de/genus/occallatibacter |
| Occidentia | https://lpsn.dsmz.de/genus/occidentia |
| Occultella | https://lpsn.dsmz.de/genus/occultella |
| Oceanibacterium | https://lpsn.dsmz.de/genus/oceanibacterium |
| Oceanibaculum | https://lpsn.dsmz.de/genus/oceanibaculum |
| Oceanibium | https://lpsn.dsmz.de/genus/oceanibium |
| Oceanibulbus | https://lpsn.dsmz.de/genus/oceanibulbus |
| Oceanicaulis | https://lpsn.dsmz.de/genus/oceanicaulis |
| Oceanicella | https://lpsn.dsmz.de/genus/oceanicella |
| Oceanicola | https://lpsn.dsmz.de/genus/oceanicola |
| Oceanidesulfovibrio | https://lpsn.dsmz.de/genus/oceanidesulfovibrio |

| | |
|---|---|
| Oceaniferula | https://lpsn.dsmz.de/genus/oceaniferula |
| Oceaniglobus | https://lpsn.dsmz.de/genus/oceaniglobus |
| Oceanihabitans | https://lpsn.dsmz.de/genus/oceanihabitans |
| Oceanimonas | https://lpsn.dsmz.de/genus/oceanimonas |
| Oceanipulchritudo | https://lpsn.dsmz.de/genus/oceanipulchritudo |
| Oceaniradius | https://lpsn.dsmz.de/genus/oceaniradius |
| Oceanirhabdus | https://lpsn.dsmz.de/genus/oceanirhabdus |
| Oceaniserpentilla | https://lpsn.dsmz.de/genus/oceaniserpentilla |
| Oceanisphaera | https://lpsn.dsmz.de/genus/oceanisphaera |
| Oceanispirochaeta | https://lpsn.dsmz.de/genus/oceanispirochaeta |
| Oceanitalea | https://lpsn.dsmz.de/genus/oceanitalea |
| Oceanithermus | https://lpsn.dsmz.de/genus/oceanithermus |
| Oceanivirga | https://lpsn.dsmz.de/genus/oceanivirga |
| Oceanobacillus | https://lpsn.dsmz.de/genus/oceanobacillus |
| Oceanobacter | https://lpsn.dsmz.de/genus/oceanobacter |
| Oceanococcus | https://lpsn.dsmz.de/genus/oceanococcus |
| Oceanomonas | https://lpsn.dsmz.de/genus/oceanomonas |
| Oceanospirillum | https://lpsn.dsmz.de/genus/oceanospirillum |
| Oceanotoga | https://lpsn.dsmz.de/genus/oceanotoga |
| Ochrobactrum | https://lpsn.dsmz.de/genus/ochrobactrum |
| Octadecabacter | https://lpsn.dsmz.de/genus/octadecabacter |
| Odoribacter | https://lpsn.dsmz.de/genus/odoribacter |
| Oecophyllibacter | https://lpsn.dsmz.de/genus/oecophyllibacter |
| Oenococcus | https://lpsn.dsmz.de/genus/oenococcus |
| Oerskovia | https://lpsn.dsmz.de/genus/oerskovia |
| Oharaeibacter | https://lpsn.dsmz.de/genus/oharaeibacter |
| Ohtaekwangia | https://lpsn.dsmz.de/genus/ohtaekwangia |
| Okibacterium | https://lpsn.dsmz.de/genus/okibacterium |
| Oleiagrimonas | https://lpsn.dsmz.de/genus/oleiagrimonas |
| Oleibacter | https://lpsn.dsmz.de/genus/oleibacter |
| Oleidesulfovibrio | https://lpsn.dsmz.de/genus/oleidesulfovibrio |
| Oleiharenicola | https://lpsn.dsmz.de/genus/oleiharenicola |
| Oleiliquidispirillum | https://lpsn.dsmz.de/genus/oleiliquidispirillum |
| Oleiphilus | https://lpsn.dsmz.de/genus/oleiphilus |
| Oleisolibacter | https://lpsn.dsmz.de/genus/oleisolibacter |
| Oleispira | https://lpsn.dsmz.de/genus/oleispira |
| Oligella | https://lpsn.dsmz.de/genus/oligella |
| Oligoflexus | https://lpsn.dsmz.de/genus/oligoflexus |
| Oligosphaera | https://lpsn.dsmz.de/genus/oligosphaera |
| Oligotropha | https://lpsn.dsmz.de/genus/oligotropha |
| Oliverpabstia | https://lpsn.dsmz.de/genus/oliverpabstia |
| Olivibacter | https://lpsn.dsmz.de/genus/olivibacter |
| Olleya | https://lpsn.dsmz.de/genus/olleya |
| Olsenella | https://lpsn.dsmz.de/genus/olsenella |

| | |
|---|---|
| Opitutus | https://lpsn.dsmz.de/genus/opitutus |
| Orbus | https://lpsn.dsmz.de/genus/orbus |
| Orenia | https://lpsn.dsmz.de/genus/orenia |
| Oribacterium | https://lpsn.dsmz.de/genus/oribacterium |
| Oribaculum | https://lpsn.dsmz.de/genus/oribaculum |
| Oricola | https://lpsn.dsmz.de/genus/oricola |
| Orientia | https://lpsn.dsmz.de/genus/orientia |
| Ornatilinea | https://lpsn.dsmz.de/genus/ornatilinea |
| Ornithinibacillus | https://lpsn.dsmz.de/genus/ornithinibacillus |
| Ornithinibacter | https://lpsn.dsmz.de/genus/ornithinibacter |
| Ornithinicoccus | https://lpsn.dsmz.de/genus/ornithinicoccus |
| Ornithinimicrobium | https://lpsn.dsmz.de/genus/ornithinimicrobium |
| Ornithobacterium | https://lpsn.dsmz.de/genus/ornithobacterium |
| Orrella | https://lpsn.dsmz.de/genus/orrella |
| Oryzihumus | https://lpsn.dsmz.de/genus/oryzihumus |
| Oryzisolibacter | https://lpsn.dsmz.de/genus/oryzisolibacter |
| Oryzobacter | https://lpsn.dsmz.de/genus/oryzobacter |
| Oryzomicrobium | https://lpsn.dsmz.de/genus/oryzomicrobium |
| Oryzomonas | https://lpsn.dsmz.de/genus/oryzomonas |
| Oscillatoria | https://lpsn.dsmz.de/genus/oscillatoria |
| Oscillibacter | https://lpsn.dsmz.de/genus/oscillibacter |
| Oscillochloris | https://lpsn.dsmz.de/genus/oscillochloris |
| Oscillospira | https://lpsn.dsmz.de/genus/oscillospira |
| Ostreibacterium | https://lpsn.dsmz.de/genus/ostreibacterium |
| Otariodibacter | https://lpsn.dsmz.de/genus/otariodibacter |
| Ottowia | https://lpsn.dsmz.de/genus/ottowia |
| Owenweeksia | https://lpsn.dsmz.de/genus/owenweeksia |
| Oxalicibacterium | https://lpsn.dsmz.de/genus/oxalicibacterium |
| Oxalobacter | https://lpsn.dsmz.de/genus/oxalobacter |
| Oxalophagus | https://lpsn.dsmz.de/genus/oxalophagus |
| Oxobacter | https://lpsn.dsmz.de/genus/oxobacter |
| Pacificamonas | https://lpsn.dsmz.de/genus/pacificamonas |
| Pacificibacter | https://lpsn.dsmz.de/genus/pacificibacter |
| Pacificimonas | https://lpsn.dsmz.de/genus/pacificimonas |
| Pacificitalea | https://lpsn.dsmz.de/genus/pacificitalea |
| Paenactinomyces | https://lpsn.dsmz.de/genus/paenactinomyces |
| Paenalcaligenes | https://lpsn.dsmz.de/genus/paenalcaligenes |
| Paenalkalicoccus | https://lpsn.dsmz.de/genus/paenalkalicoccus |
| Paenarthrobacter | https://lpsn.dsmz.de/genus/paenarthrobacter |
| Paenibacillus | https://lpsn.dsmz.de/genus/paenibacillus |
| Paeniclostridium | https://lpsn.dsmz.de/genus/paeniclostridium |
| Paeniglutamicibacter | https://lpsn.dsmz.de/genus/paeniglutamicibacter |
| Paenihalocynthiibacter | https://lpsn.dsmz.de/genus/paenihalocynthiibacter |
| Paenimaribius | https://lpsn.dsmz.de/genus/paenimaribius |

| | |
|---|---|
| Paenirhodobacter | https://lpsn.dsmz.de/genus/paenirhodobacter |
| Paeniroseomonas | https://lpsn.dsmz.de/genus/paeniroseomonas |
| Paenisporosarcina | https://lpsn.dsmz.de/genus/paenisporosarcina |
| Paenochrobactrum | https://lpsn.dsmz.de/genus/paenochrobactrum |
| Palaeococcus | https://lpsn.dsmz.de/genus/palaeococcus |
| Palleronia | https://lpsn.dsmz.de/genus/palleronia |
| Paludibacter | https://lpsn.dsmz.de/genus/paludibacter |
| Paludibacterium | https://lpsn.dsmz.de/genus/paludibacterium |
| Paludibaculum | https://lpsn.dsmz.de/genus/paludibaculum |
| Paludicola | https://lpsn.dsmz.de/genus/paludicola |
| Paludifilum | https://lpsn.dsmz.de/genus/paludifilum |
| Paludirhabdus | https://lpsn.dsmz.de/genus/paludirhabdus |
| Paludisphaera | https://lpsn.dsmz.de/genus/paludisphaera |
| Panacagrimonas | https://lpsn.dsmz.de/genus/panacagrimonas |
| Panacibacter | https://lpsn.dsmz.de/genus/panacibacter |
| Pandoraea | https://lpsn.dsmz.de/genus/pandoraea |
| Pannonibacter | https://lpsn.dsmz.de/genus/pannonibacter |
| Pantoea | https://lpsn.dsmz.de/genus/pantoea |
| Papillibacter | https://lpsn.dsmz.de/genus/papillibacter |
| Parabacteroides | https://lpsn.dsmz.de/genus/parabacteroides |
| Parablastomonas | https://lpsn.dsmz.de/genus/parablastomonas |
| Paraburkholderia | https://lpsn.dsmz.de/genus/paraburkholderia |
| Paracandidimonas | https://lpsn.dsmz.de/genus/paracandidimonas |
| Parachlamydia | https://lpsn.dsmz.de/genus/parachlamydia |
| Paracholeplasma | https://lpsn.dsmz.de/genus/paracholeplasma |
| Parachryseolinea | https://lpsn.dsmz.de/genus/parachryseolinea |
| Paracidobacterium | https://lpsn.dsmz.de/genus/paracidobacterium |
| Paraclostridium | https://lpsn.dsmz.de/genus/paraclostridium |
| Paracnuella | https://lpsn.dsmz.de/genus/paracnuella |
| Paracoccus | https://lpsn.dsmz.de/genus/paracoccus |
| Paraconexibacter | https://lpsn.dsmz.de/genus/paraconexibacter |
| Paracraurococcus | https://lpsn.dsmz.de/genus/paracraurococcus |
| Paradesulfitobacterium | https://lpsn.dsmz.de/genus/paradesulfitobacterium |
| Paradesulfovibrio | https://lpsn.dsmz.de/genus/paradesulfovibrio |
| Paradevosia | https://lpsn.dsmz.de/genus/paradevosia |
| Paradonghicola | https://lpsn.dsmz.de/genus/paradonghicola |
| Paraeggerthella | https://lpsn.dsmz.de/genus/paraeggerthella |
| Parafannyhessea | https://lpsn.dsmz.de/genus/parafannyhessea |
| Paraferrimonas | https://lpsn.dsmz.de/genus/paraferrimonas |
| Parafilimonas | https://lpsn.dsmz.de/genus/parafilimonas |
| Paraflavitalea | https://lpsn.dsmz.de/genus/paraflavitalea |
| Parafrigoribacterium | https://lpsn.dsmz.de/genus/parafrigoribacterium |
| Parageobacillus | https://lpsn.dsmz.de/genus/parageobacillus |
| Paraglaciecola | https://lpsn.dsmz.de/genus/paraglaciecola |

| | |
|---|---|
| Paraglycomyces | https://lpsn.dsmz.de/genus/paraglycomyces |
| Parahaliea | https://lpsn.dsmz.de/genus/parahaliea |
| Paraherbaspirillum | https://lpsn.dsmz.de/genus/paraherbaspirillum |
| Paralactobacillus | https://lpsn.dsmz.de/genus/paralactobacillus |
| Paralcaligenes | https://lpsn.dsmz.de/genus/paralcaligenes |
| Paraliobacillus | https://lpsn.dsmz.de/genus/paraliobacillus |
| Paralkalibacillus | https://lpsn.dsmz.de/genus/paralkalibacillus |
| Paralysiella | https://lpsn.dsmz.de/genus/paralysiella |
| Paramaledivibacter | https://lpsn.dsmz.de/genus/paramaledivibacter |
| Paramesonia | https://lpsn.dsmz.de/genus/paramesonia |
| Paramoritella | https://lpsn.dsmz.de/genus/paramoritella |
| Paramuribaculum | https://lpsn.dsmz.de/genus/paramuribaculum |
| Paraneptunicella | https://lpsn.dsmz.de/genus/paraneptunicella |
| Paraoceanicella | https://lpsn.dsmz.de/genus/paraoceanicella |
| Paraoerskovia | https://lpsn.dsmz.de/genus/paraoerskovia |
| Parapedobacter | https://lpsn.dsmz.de/genus/parapedobacter |
| Paraperlucidibaca | https://lpsn.dsmz.de/genus/paraperlucidibaca |
| Paraphaeobacter | https://lpsn.dsmz.de/genus/paraphaeobacter |
| Paraphotobacterium | https://lpsn.dsmz.de/genus/paraphotobacterium |
| Parapontixanthobacter | https://lpsn.dsmz.de/genus/parapontixanthobacter |
| Paraprevotella | https://lpsn.dsmz.de/genus/paraprevotella |
| Parapseudoflavitalea | https://lpsn.dsmz.de/genus/parapseudoflavitalea |
| Parapseudomonas | https://lpsn.dsmz.de/genus/parapseudomonas |
| Parapusillimonas | https://lpsn.dsmz.de/genus/parapusillimonas |
| Pararcticibacter | https://lpsn.dsmz.de/genus/pararcticibacter |
| Pararhizobium | https://lpsn.dsmz.de/genus/pararhizobium |
| Pararhodobacter | https://lpsn.dsmz.de/genus/pararhodobacter |
| Pararhodonellum | https://lpsn.dsmz.de/genus/pararhodonellum |
| Pararhodospirillum | https://lpsn.dsmz.de/genus/pararhodospirillum |
| Pararobbsia | https://lpsn.dsmz.de/genus/pararobbsia |
| Pararoseburia | https://lpsn.dsmz.de/genus/pararoseburia |
| Pararoseomonas | https://lpsn.dsmz.de/genus/pararoseomonas |
| Parascardovia | https://lpsn.dsmz.de/genus/parascardovia |
| Parasedimentitalea | https://lpsn.dsmz.de/genus/parasedimentitalea |
| Parasediminibacterium | https://lpsn.dsmz.de/genus/parasediminibacterium |
| Parasegetibacter | https://lpsn.dsmz.de/genus/parasegetibacter |
| Parashewanella | https://lpsn.dsmz.de/genus/parashewanella |
| Parasphaerochaeta | https://lpsn.dsmz.de/genus/parasphaerochaeta |
| Parasphingopyxis | https://lpsn.dsmz.de/genus/parasphingopyxis |
| Parasphingorhabdus | https://lpsn.dsmz.de/genus/parasphingorhabdus |
| Parasporobacterium | https://lpsn.dsmz.de/genus/parasporobacterium |
| Parasulfuritortus | https://lpsn.dsmz.de/genus/parasulfuritortus |
| Parasutterella | https://lpsn.dsmz.de/genus/parasutterella |
| Paratractidigestivibacter | https://lpsn.dsmz.de/genus/paratractidigestivibacter |

| | |
|---|---|
| Paraurantiacibacter | https://lpsn.dsmz.de/genus/paraurantiacibacter |
| Parazoarcus | https://lpsn.dsmz.de/genus/parazoarcus |
| Paremcibacter | https://lpsn.dsmz.de/genus/paremcibacter |
| Parendozoicomonas | https://lpsn.dsmz.de/genus/parendozoicomonas |
| Parenemella | https://lpsn.dsmz.de/genus/parenemella |
| Parerythrobacter | https://lpsn.dsmz.de/genus/parerythrobacter |
| Pareuzebyella | https://lpsn.dsmz.de/genus/pareuzebyella |
| Paroceanicella | https://lpsn.dsmz.de/genus/paroceanicella |
| Parolsenella | https://lpsn.dsmz.de/genus/parolsenella |
| Parvibacter | https://lpsn.dsmz.de/genus/parvibacter |
| Parvibaculum | https://lpsn.dsmz.de/genus/parvibaculum |
| Parvibium | https://lpsn.dsmz.de/genus/parvibium |
| Parvicella | https://lpsn.dsmz.de/genus/parvicella |
| Parvimonas | https://lpsn.dsmz.de/genus/parvimonas |
| Parviterribacter | https://lpsn.dsmz.de/genus/parviterribacter |
| Parvularcula | https://lpsn.dsmz.de/genus/parvularcula |
| Pasteurella | https://lpsn.dsmz.de/genus/pasteurella |
| Pasteuria | https://lpsn.dsmz.de/genus/pasteuria |
| Patiriisocius | https://lpsn.dsmz.de/genus/patiriisocius |
| Patulibacter | https://lpsn.dsmz.de/genus/patulibacter |
| Paucibacter | https://lpsn.dsmz.de/genus/paucibacter |
| Paucidesulfovibrio | https://lpsn.dsmz.de/genus/paucidesulfovibrio |
| Paucihalobacter | https://lpsn.dsmz.de/genus/paucihalobacter |
| Paucilactobacillus | https://lpsn.dsmz.de/genus/paucilactobacillus |
| Paucimonas | https://lpsn.dsmz.de/genus/paucimonas |
| Paucisalibacillus | https://lpsn.dsmz.de/genus/paucisalibacillus |
| Pauljensenia | https://lpsn.dsmz.de/genus/pauljensenia |
| Pectinatus | https://lpsn.dsmz.de/genus/pectinatus |
| Pectobacterium | https://lpsn.dsmz.de/genus/pectobacterium |
| Pediococcus | https://lpsn.dsmz.de/genus/pediococcus |
| Pedobacter | https://lpsn.dsmz.de/genus/pedobacter |
| Pedococcus | https://lpsn.dsmz.de/genus/pedococcus |
| Pedomicrobium | https://lpsn.dsmz.de/genus/pedomicrobium |
| Pedomonas | https://lpsn.dsmz.de/genus/pedomonas |
| Pelagerythrobacter | https://lpsn.dsmz.de/genus/pelagerythrobacter |
| Pelagibaca | https://lpsn.dsmz.de/genus/pelagibaca |
| Pelagibacillus | https://lpsn.dsmz.de/genus/pelagibacillus |
| Pelagibacterium | https://lpsn.dsmz.de/genus/pelagibacterium |
| Pelagibaculum | https://lpsn.dsmz.de/genus/pelagibaculum |
| Pelagibius | https://lpsn.dsmz.de/genus/pelagibius |
| Pelagicoccus | https://lpsn.dsmz.de/genus/pelagicoccus |
| Pelagicola | https://lpsn.dsmz.de/genus/pelagicola |
| Pelagihabitans | https://lpsn.dsmz.de/genus/pelagihabitans |
| Pelagimonas | https://lpsn.dsmz.de/genus/pelagimonas |

| | |
|---|---|
| Pelagirhabdus | https://lpsn.dsmz.de/genus/pelagirhabdus |
| Pelagivirga | https://lpsn.dsmz.de/genus/pelagivirga |
| Pelagovum | https://lpsn.dsmz.de/genus/pelagovum |
| Pelczaria | https://lpsn.dsmz.de/genus/pelczaria |
| Pelistega | https://lpsn.dsmz.de/genus/pelistega |
| Pelobacter | https://lpsn.dsmz.de/genus/pelobacter |
| Pelobium | https://lpsn.dsmz.de/genus/pelobium |
| Pelodictyon | https://lpsn.dsmz.de/genus/pelodictyon |
| Pelolinea | https://lpsn.dsmz.de/genus/pelolinea |
| Pelomicrobium | https://lpsn.dsmz.de/genus/pelomicrobium |
| Pelomonas | https://lpsn.dsmz.de/genus/pelomonas |
| Pelorhabdus | https://lpsn.dsmz.de/genus/pelorhabdus |
| Pelosinus | https://lpsn.dsmz.de/genus/pelosinus |
| Pelospora | https://lpsn.dsmz.de/genus/pelospora |
| Pelotalea | https://lpsn.dsmz.de/genus/pelotalea |
| Pelotomaculum | https://lpsn.dsmz.de/genus/pelotomaculum |
| Penaeicola | https://lpsn.dsmz.de/genus/penaeicola |
| Pengzhenrongella | https://lpsn.dsmz.de/genus/pengzhenrongella |
| Peptacetobacter | https://lpsn.dsmz.de/genus/peptacetobacter |
| Peptidiphaga | https://lpsn.dsmz.de/genus/peptidiphaga |
| Peptoanaerobacter | https://lpsn.dsmz.de/genus/peptoanaerobacter |
| Peptoclostridium | https://lpsn.dsmz.de/genus/peptoclostridium |
| Peptococcus | https://lpsn.dsmz.de/genus/peptococcus |
| Peptoniphilus | https://lpsn.dsmz.de/genus/peptoniphilus |
| Peptostreptococcus | https://lpsn.dsmz.de/genus/peptostreptococcus |
| Peredibacter | https://lpsn.dsmz.de/genus/peredibacter |
| Perexilibacter | https://lpsn.dsmz.de/genus/perexilibacter |
| Peribacillus | https://lpsn.dsmz.de/genus/peribacillus |
| Periweissella | https://lpsn.dsmz.de/genus/periweissella |
| Perlabentimonas | https://lpsn.dsmz.de/genus/perlabentimonas |
| Perlucidibaca | https://lpsn.dsmz.de/genus/perlucidibaca |
| Permianibacter | https://lpsn.dsmz.de/genus/permianibacter |
| Persephonella | https://lpsn.dsmz.de/genus/persephonella |
| Persicimonas | https://lpsn.dsmz.de/genus/persicimonas |
| Persicirhabdus | https://lpsn.dsmz.de/genus/persicirhabdus |
| Persicitalea | https://lpsn.dsmz.de/genus/persicitalea |
| Persicivirga | https://lpsn.dsmz.de/genus/persicivirga |
| Persicobacter | https://lpsn.dsmz.de/genus/persicobacter |
| Perspicuibacter | https://lpsn.dsmz.de/genus/perspicuibacter |
| Peterkaempfera | https://lpsn.dsmz.de/genus/peterkaempfera |
| Peteryoungia | https://lpsn.dsmz.de/genus/peteryoungia |
| Petrimonas | https://lpsn.dsmz.de/genus/petrimonas |
| Petrobacter | https://lpsn.dsmz.de/genus/petrobacter |
| Petrocella | https://lpsn.dsmz.de/genus/petrocella |

| | |
|---|---|
| Petroclostridium | https://lpsn.dsmz.de/genus/petroclostridium |
| Petrothermobacter | https://lpsn.dsmz.de/genus/petrothermobacter |
| Petrotoga | https://lpsn.dsmz.de/genus/petrotoga |
| Pfennigia | https://lpsn.dsmz.de/genus/pfennigia |
| Phaeacidiphilus | https://lpsn.dsmz.de/genus/phaeacidiphilus |
| Phaeobacter | https://lpsn.dsmz.de/genus/phaeobacter |
| Phaeobacterium | https://lpsn.dsmz.de/genus/phaeobacterium |
| Phaeochromatium | https://lpsn.dsmz.de/genus/phaeochromatium |
| Phaeocystidibacter | https://lpsn.dsmz.de/genus/phaeocystidibacter |
| Phaeodactylibacter | https://lpsn.dsmz.de/genus/phaeodactylibacter |
| Phaeospirillum | https://lpsn.dsmz.de/genus/phaeospirillum |
| Phaeovibrio | https://lpsn.dsmz.de/genus/phaeovibrio |
| Phaeovulum | https://lpsn.dsmz.de/genus/phaeovulum |
| Phascolarctobacterium | https://lpsn.dsmz.de/genus/phascolarctobacterium |
| Phaselicystis | https://lpsn.dsmz.de/genus/phaselicystis |
| Phaseolibacter | https://lpsn.dsmz.de/genus/phaseolibacter |
| Phenylobacterium | https://lpsn.dsmz.de/genus/phenylobacterium |
| Phnomibacter | https://lpsn.dsmz.de/genus/phnomibacter |
| Phocaeicola | https://lpsn.dsmz.de/genus/phocaeicola |
| Phocoenobacter | https://lpsn.dsmz.de/genus/phocoenobacter |
| Phorcysia | https://lpsn.dsmz.de/genus/phorcysia |
| Phosphitispora | https://lpsn.dsmz.de/genus/phosphitispora |
| Photobacterium | https://lpsn.dsmz.de/genus/photobacterium |
| Photorhabdus | https://lpsn.dsmz.de/genus/photorhabdus |
| Phragmitibacter | https://lpsn.dsmz.de/genus/phragmitibacter |
| Phreatobacter | https://lpsn.dsmz.de/genus/phreatobacter |
| Phycicoccus | https://lpsn.dsmz.de/genus/phycicoccus |
| Phycicola | https://lpsn.dsmz.de/genus/phycicola |
| Phycisphaera | https://lpsn.dsmz.de/genus/phycisphaera |
| Phycocomes | https://lpsn.dsmz.de/genus/phycocomes |
| Phyllobacterium | https://lpsn.dsmz.de/genus/phyllobacterium |
| Phytoactinopolyspora | https://lpsn.dsmz.de/genus/phytoactinopolyspora |
| Phytobacter | https://lpsn.dsmz.de/genus/phytobacter |
| Phytohabitans | https://lpsn.dsmz.de/genus/phytohabitans |
| Phytomonospora | https://lpsn.dsmz.de/genus/phytomonospora |
| Pibocella | https://lpsn.dsmz.de/genus/pibocella |
| Picrophilus | https://lpsn.dsmz.de/genus/picrophilus |
| Pigmentiphaga | https://lpsn.dsmz.de/genus/pigmentiphaga |
| Pikeienuella | https://lpsn.dsmz.de/genus/pikeienuella |
| Pilibacter | https://lpsn.dsmz.de/genus/pilibacter |
| Pilimelia | https://lpsn.dsmz.de/genus/pilimelia |
| Pillotina | https://lpsn.dsmz.de/genus/pillotina |
| Pimelobacter | https://lpsn.dsmz.de/genus/pimelobacter |
| Pinibacter | https://lpsn.dsmz.de/genus/pinibacter |

| | |
|---|---|
| Pinisolibacter | https://lpsn.dsmz.de/genus/pinisolibacter |
| Pirella | https://lpsn.dsmz.de/genus/pirella |
| Pirellula | https://lpsn.dsmz.de/genus/pirellula |
| Pirellulimonas | https://lpsn.dsmz.de/genus/pirellulimonas |
| Piscibacillus | https://lpsn.dsmz.de/genus/piscibacillus |
| Piscicoccus | https://lpsn.dsmz.de/genus/piscicoccus |
| Pisciglobus | https://lpsn.dsmz.de/genus/pisciglobus |
| Piscinibacter | https://lpsn.dsmz.de/genus/piscinibacter |
| Piscinibacterium | https://lpsn.dsmz.de/genus/piscinibacterium |
| Piscirickettsia | https://lpsn.dsmz.de/genus/piscirickettsia |
| Pistricoccus | https://lpsn.dsmz.de/genus/pistricoccus |
| Planctobacterium | https://lpsn.dsmz.de/genus/planctobacterium |
| Planctomicrobium | https://lpsn.dsmz.de/genus/planctomicrobium |
| Planctomonas | https://lpsn.dsmz.de/genus/planctomonas |
| Planctomyces | https://lpsn.dsmz.de/genus/planctomyces |
| Planctopirus | https://lpsn.dsmz.de/genus/planctopirus |
| Planifilum | https://lpsn.dsmz.de/genus/planifilum |
| Planktomarina | https://lpsn.dsmz.de/genus/planktomarina |
| Planktosalinus | https://lpsn.dsmz.de/genus/planktosalinus |
| Planktotalea | https://lpsn.dsmz.de/genus/planktotalea |
| Planobacterium | https://lpsn.dsmz.de/genus/planobacterium |
| Planobispora | https://lpsn.dsmz.de/genus/planobispora |
| Planococcus | https://lpsn.dsmz.de/genus/planococcus |
| Planomicrobium | https://lpsn.dsmz.de/genus/planomicrobium |
| Planomonospora | https://lpsn.dsmz.de/genus/planomonospora |
| Planopolyspora | https://lpsn.dsmz.de/genus/planopolyspora |
| Planosporangium | https://lpsn.dsmz.de/genus/planosporangium |
| Planotetraspora | https://lpsn.dsmz.de/genus/planotetraspora |
| Plantactinospora | https://lpsn.dsmz.de/genus/plantactinospora |
| Plantibacter | https://lpsn.dsmz.de/genus/plantibacter |
| Plasticicumulans | https://lpsn.dsmz.de/genus/plasticicumulans |
| Plastoroseomonas | https://lpsn.dsmz.de/genus/plastoroseomonas |
| Pleionea | https://lpsn.dsmz.de/genus/pleionea |
| Pleisomonas | https://lpsn.dsmz.de/genus/pleisomonas |
| Pleomorphobacterium | https://lpsn.dsmz.de/genus/pleomorphobacterium |
| Pleomorphochaeta | https://lpsn.dsmz.de/genus/pleomorphochaeta |
| Pleomorphomonas | https://lpsn.dsmz.de/genus/pleomorphomonas |
| Pleomorphovibrio | https://lpsn.dsmz.de/genus/pleomorphovibrio |
| Plesiocystis | https://lpsn.dsmz.de/genus/plesiocystis |
| Plesiomonas | https://lpsn.dsmz.de/genus/plesiomonas |
| Pluralibacter | https://lpsn.dsmz.de/genus/pluralibacter |
| Pokkaliibacter | https://lpsn.dsmz.de/genus/pokkaliibacter |
| Polaribacter | https://lpsn.dsmz.de/genus/polaribacter |
| Polaromonas | https://lpsn.dsmz.de/genus/polaromonas |

| | |
|---|---|
| Polyangium | https://lpsn.dsmz.de/genus/polyangium |
| Polycladomyces | https://lpsn.dsmz.de/genus/polycladomyces |
| Polycyclovorans | https://lpsn.dsmz.de/genus/polycyclovorans |
| Polygonibacillus | https://lpsn.dsmz.de/genus/polygonibacillus |
| Polymorphobacter | https://lpsn.dsmz.de/genus/polymorphobacter |
| Polymorphospora | https://lpsn.dsmz.de/genus/polymorphospora |
| Polynucleobacter | https://lpsn.dsmz.de/genus/polynucleobacter |
| Polystyrenella | https://lpsn.dsmz.de/genus/polystyrenella |
| Pontibaca | https://lpsn.dsmz.de/genus/pontibaca |
| Pontibacillus | https://lpsn.dsmz.de/genus/pontibacillus |
| Pontibacter | https://lpsn.dsmz.de/genus/pontibacter |
| Pontibacterium | https://lpsn.dsmz.de/genus/pontibacterium |
| Pontibrevibacter | https://lpsn.dsmz.de/genus/pontibrevibacter |
| Ponticaulis | https://lpsn.dsmz.de/genus/ponticaulis |
| Ponticoccus | https://lpsn.dsmz.de/genus/ponticoccus |
| Pontiella | https://lpsn.dsmz.de/genus/pontiella |
| Pontimicrobium | https://lpsn.dsmz.de/genus/pontimicrobium |
| Pontimonas | https://lpsn.dsmz.de/genus/pontimonas |
| Pontirhabdus | https://lpsn.dsmz.de/genus/pontirhabdus |
| Pontivivens | https://lpsn.dsmz.de/genus/pontivivens |
| Pontixanthobacter | https://lpsn.dsmz.de/genus/pontixanthobacter |
| Populibacterium | https://lpsn.dsmz.de/genus/populibacterium |
| Porcincola | https://lpsn.dsmz.de/genus/porcincola |
| Poriferisphaera | https://lpsn.dsmz.de/genus/poriferisphaera |
| Poritiphilus | https://lpsn.dsmz.de/genus/poritiphilus |
| Porphyrobacter | https://lpsn.dsmz.de/genus/porphyrobacter |
| Porphyromonas | https://lpsn.dsmz.de/genus/porphyromonas |
| Portibacter | https://lpsn.dsmz.de/genus/portibacter |
| Porticoccus | https://lpsn.dsmz.de/genus/porticoccus |
| Poseidonibacter | https://lpsn.dsmz.de/genus/poseidonibacter |
| Poseidonocella | https://lpsn.dsmz.de/genus/poseidonocella |
| Posidoniimonas | https://lpsn.dsmz.de/genus/posidoniimonas |
| Postechiella | https://lpsn.dsmz.de/genus/postechiella |
| Povalibacter | https://lpsn.dsmz.de/genus/povalibacter |
| Pradoshia | https://lpsn.dsmz.de/genus/pradoshia |
| Pragia | https://lpsn.dsmz.de/genus/pragia |
| Prauserella | https://lpsn.dsmz.de/genus/prauserella |
| Prescottella | https://lpsn.dsmz.de/genus/prescottella |
| Prevotella | https://lpsn.dsmz.de/genus/prevotella |
| Pricia | https://lpsn.dsmz.de/genus/pricia |
| Priestia | https://lpsn.dsmz.de/genus/priestia |
| Primorskyibacter | https://lpsn.dsmz.de/genus/primorskyibacter |
| Prochloron | https://lpsn.dsmz.de/genus/prochloron |
| Prochlorothrix | https://lpsn.dsmz.de/genus/prochlorothrix |

| | |
|---|---|
| Profundibacter | https://lpsn.dsmz.de/genus/profundibacter |
| Profundibacterium | https://lpsn.dsmz.de/genus/profundibacterium |
| Prolinoborus | https://lpsn.dsmz.de/genus/prolinoborus |
| Prolixibacter | https://lpsn.dsmz.de/genus/prolixibacter |
| Promicromonospora | https://lpsn.dsmz.de/genus/promicromonospora |
| Propionibacter | https://lpsn.dsmz.de/genus/propionibacter |
| Propionibacterium | https://lpsn.dsmz.de/genus/propionibacterium |
| Propionicicella | https://lpsn.dsmz.de/genus/propionicicella |
| Propioniciclava | https://lpsn.dsmz.de/genus/propioniciclava |
| Propionicimonas | https://lpsn.dsmz.de/genus/propionicimonas |
| Propioniferax | https://lpsn.dsmz.de/genus/propioniferax |
| Propionigenium | https://lpsn.dsmz.de/genus/propionigenium |
| Propionimicrobium | https://lpsn.dsmz.de/genus/propionimicrobium |
| Propionispira | https://lpsn.dsmz.de/genus/propionispira |
| Propionispora | https://lpsn.dsmz.de/genus/propionispora |
| Propionivibrio | https://lpsn.dsmz.de/genus/propionivibrio |
| Prosthecobacter | https://lpsn.dsmz.de/genus/prosthecobacter |
| Prosthecochloris | https://lpsn.dsmz.de/genus/prosthecochloris |
| Prosthecomicrobium | https://lpsn.dsmz.de/genus/prosthecomicrobium |
| Protaetiibacter | https://lpsn.dsmz.de/genus/protaetiibacter |
| Proteiniborus | https://lpsn.dsmz.de/genus/proteiniborus |
| Proteiniclasticum | https://lpsn.dsmz.de/genus/proteiniclasticum |
| Proteiniphilum | https://lpsn.dsmz.de/genus/proteiniphilum |
| Proteinivorax | https://lpsn.dsmz.de/genus/proteinivorax |
| Proteocatella | https://lpsn.dsmz.de/genus/proteocatella |
| Proteus | https://lpsn.dsmz.de/genus/proteus |
| Protomonas | https://lpsn.dsmz.de/genus/protomonas |
| Providencia | https://lpsn.dsmz.de/genus/providencia |
| Pseudacidobacterium | https://lpsn.dsmz.de/genus/pseudacidobacterium |
| Pseudacidovorax | https://lpsn.dsmz.de/genus/pseudacidovorax |
| Pseudactinotalea | https://lpsn.dsmz.de/genus/pseudactinotalea |
| Pseudaeromonas | https://lpsn.dsmz.de/genus/pseudaeromonas |
| Pseudaestuariivita | https://lpsn.dsmz.de/genus/pseudaestuariivita |
| Pseudahrensia | https://lpsn.dsmz.de/genus/pseudahrensia |
| Pseudalgibacter | https://lpsn.dsmz.de/genus/pseudalgibacter |
| Pseudalkalibacillus | https://lpsn.dsmz.de/genus/pseudalkalibacillus |
| Pseudaminobacter | https://lpsn.dsmz.de/genus/pseudaminobacter |
| Pseudanabaena | https://lpsn.dsmz.de/genus/pseudanabaena |
| Pseudaquidulcibacter | https://lpsn.dsmz.de/genus/pseudaquidulcibacter |
| Pseudarcicella | https://lpsn.dsmz.de/genus/pseudarcicella |
| Pseudarcobacter | https://lpsn.dsmz.de/genus/pseudarcobacter |
| Pseudarthrobacter | https://lpsn.dsmz.de/genus/pseudarthrobacter |
| Pseudazoarcus | https://lpsn.dsmz.de/genus/pseudazoarcus |
| Pseudemcibacter | https://lpsn.dsmz.de/genus/pseudemcibacter |

| | |
|---|---|
| Pseudenhygromyxa | https://lpsn.dsmz.de/genus/pseudenhygromyxa |
| Pseudescherichia | https://lpsn.dsmz.de/genus/pseudescherichia |
| Pseudidiomarina | https://lpsn.dsmz.de/genus/pseudidiomarina |
| Pseudoalteromonas | https://lpsn.dsmz.de/genus/pseudoalteromonas |
| Pseudoamycolata | https://lpsn.dsmz.de/genus/pseudoamycolata |
| Pseudoarcobacter | https://lpsn.dsmz.de/genus/pseudoarcobacter |
| Pseudobacter | https://lpsn.dsmz.de/genus/pseudobacter |
| Pseudobacteriovorax | https://lpsn.dsmz.de/genus/pseudobacteriovorax |
| Pseudobacteroides | https://lpsn.dsmz.de/genus/pseudobacteroides |
| Pseudobdellovibrio | https://lpsn.dsmz.de/genus/pseudobdellovibrio |
| Pseudobizionia | https://lpsn.dsmz.de/genus/pseudobizionia |
| Pseudobowmanella | https://lpsn.dsmz.de/genus/pseudobowmanella |
| Pseudobutyrivibrio | https://lpsn.dsmz.de/genus/pseudobutyrivibrio |
| Pseudobythopirellula | https://lpsn.dsmz.de/genus/pseudobythopirellula |
| Pseudocaedibacter | https://lpsn.dsmz.de/genus/pseudocaedibacter |
| Pseudochelatococcus | https://lpsn.dsmz.de/genus/pseudochelatococcus |
| Pseudochrobactrum | https://lpsn.dsmz.de/genus/pseudochrobactrum |
| Pseudochryseolinea | https://lpsn.dsmz.de/genus/pseudochryseolinea |
| Pseudocitrobacter | https://lpsn.dsmz.de/genus/pseudocitrobacter |
| Pseudoclavibacter | https://lpsn.dsmz.de/genus/pseudoclavibacter |
| Pseudocolwellia | https://lpsn.dsmz.de/genus/pseudocolwellia |
| Pseudodesulfovibrio | https://lpsn.dsmz.de/genus/pseudodesulfovibrio |
| Pseudodonghicola | https://lpsn.dsmz.de/genus/pseudodonghicola |
| Pseudoduganella | https://lpsn.dsmz.de/genus/pseudoduganella |
| Pseudoflavitalea | https://lpsn.dsmz.de/genus/pseudoflavitalea |
| Pseudoflavonifractor | https://lpsn.dsmz.de/genus/pseudoflavonifractor |
| Pseudofrancisella | https://lpsn.dsmz.de/genus/pseudofrancisella |
| Pseudofulvibacter | https://lpsn.dsmz.de/genus/pseudofulvibacter |
| Pseudofulvimonas | https://lpsn.dsmz.de/genus/pseudofulvimonas |
| Pseudogemmobacter | https://lpsn.dsmz.de/genus/pseudogemmobacter |
| Pseudoglutamicibacter | https://lpsn.dsmz.de/genus/pseudoglutamicibacter |
| Pseudogracilibacillus | https://lpsn.dsmz.de/genus/pseudogracilibacillus |
| Pseudogulbenkiania | https://lpsn.dsmz.de/genus/pseudogulbenkiania |
| Pseudohaliea | https://lpsn.dsmz.de/genus/pseudohaliea |
| Pseudohalioglobus | https://lpsn.dsmz.de/genus/pseudohalioglobus |
| Pseudohalocynthiibacter | https://lpsn.dsmz.de/genus/pseudohalocynthiibacter |
| Pseudohoeflea | https://lpsn.dsmz.de/genus/pseudohoeflea |
| Pseudohongiella | https://lpsn.dsmz.de/genus/pseudohongiella |
| Pseudokineococcus | https://lpsn.dsmz.de/genus/pseudokineococcus |
| Pseudolabrys | https://lpsn.dsmz.de/genus/pseudolabrys |
| Pseudoleptotrichia | https://lpsn.dsmz.de/genus/pseudoleptotrichia |
| Pseudolysinimonas | https://lpsn.dsmz.de/genus/pseudolysinimonas |
| Pseudolysobacter | https://lpsn.dsmz.de/genus/pseudolysobacter |
| Pseudomaribius | https://lpsn.dsmz.de/genus/pseudomaribius |

| | |
|---|---|
| Pseudomaricurvus | https://lpsn.dsmz.de/genus/pseudomaricurvus |
| Pseudomarimonas | https://lpsn.dsmz.de/genus/pseudomarimonas |
| Pseudomethylobacillus | https://lpsn.dsmz.de/genus/pseudomethylobacillus |
| Pseudomonas | https://lpsn.dsmz.de/genus/pseudomonas |
| Pseudoneobacillus | https://lpsn.dsmz.de/genus/pseudoneobacillus |
| Pseudonocardia | https://lpsn.dsmz.de/genus/pseudonocardia |
| Pseudooceanicola | https://lpsn.dsmz.de/genus/pseudooceanicola |
| Pseudooctadecabacter | https://lpsn.dsmz.de/genus/pseudooctadecabacter |
| Pseudopedobacter | https://lpsn.dsmz.de/genus/pseudopedobacter |
| Pseudopelagicola | https://lpsn.dsmz.de/genus/pseudopelagicola |
| Pseudophaeobacter | https://lpsn.dsmz.de/genus/pseudophaeobacter |
| Pseudoponticoccus | https://lpsn.dsmz.de/genus/pseudoponticoccus |
| Pseudopontixanthobacter | https://lpsn.dsmz.de/genus/pseudopontixanthobacter-1 |
| Pseudopontixanthobacter | https://lpsn.dsmz.de/genus/pseudopontixanthobacter |
| Pseudoprevotella | https://lpsn.dsmz.de/genus/pseudoprevotella |
| Pseudoprimorskyibacter | https://lpsn.dsmz.de/genus/pseudoprimorskyibacter |
| Pseudopropionibacterium | https://lpsn.dsmz.de/genus/pseudopropionibacterium |
| Pseudopuniceibacterium | https://lpsn.dsmz.de/genus/pseudopuniceibacterium |
| Pseudoramibacter | https://lpsn.dsmz.de/genus/pseudoramibacter |
| Pseudorhizobium | https://lpsn.dsmz.de/genus/pseudorhizobium |
| Pseudorhodobacter | https://lpsn.dsmz.de/genus/pseudorhodobacter |
| Pseudorhodoferax | https://lpsn.dsmz.de/genus/pseudorhodoferax |
| Pseudorhodoplanes | https://lpsn.dsmz.de/genus/pseudorhodoplanes |
| Pseudorivibacter | https://lpsn.dsmz.de/genus/pseudorivibacter |
| Pseudoroseicyclus | https://lpsn.dsmz.de/genus/pseudoroseicyclus |
| Pseudoroseomonas | https://lpsn.dsmz.de/genus/pseudoroseomonas |
| Pseudoroseovarius | https://lpsn.dsmz.de/genus/pseudoroseovarius |
| Pseudoruegeria | https://lpsn.dsmz.de/genus/pseudoruegeria |
| Pseudoscardovia | https://lpsn.dsmz.de/genus/pseudoscardovia |
| Pseudoseohaeicola | https://lpsn.dsmz.de/genus/pseudoseohaeicola |
| Pseudosphingobacterium | https://lpsn.dsmz.de/genus/pseudosphingobacterium |
| Pseudospirillum | https://lpsn.dsmz.de/genus/pseudospirillum |
| Pseudosporangium | https://lpsn.dsmz.de/genus/pseudosporangium |
| Pseudostreptobacillus | https://lpsn.dsmz.de/genus/pseudostreptobacillus |
| Pseudosulfitobacter | https://lpsn.dsmz.de/genus/pseudosulfitobacter |
| Pseudotabrizicola | https://lpsn.dsmz.de/genus/pseudotabrizicola |
| Pseudotenacibaculum | https://lpsn.dsmz.de/genus/pseudotenacibaculum |
| Pseudoteredinibacter | https://lpsn.dsmz.de/genus/pseudoteredinibacter |
| Pseudothauera | https://lpsn.dsmz.de/genus/pseudothauera |
| Pseudothermotoga | https://lpsn.dsmz.de/genus/pseudothermotoga |
| Pseudothioclava | https://lpsn.dsmz.de/genus/pseudothioclava |
| Pseudovibrio | https://lpsn.dsmz.de/genus/pseudovibrio |
| Pseudoxanthobacter | https://lpsn.dsmz.de/genus/pseudoxanthobacter |
| Pseudoxanthomonas | https://lpsn.dsmz.de/genus/pseudoxanthomonas |

| | |
|---|---|
| Pseudozobellia | https://lpsn.dsmz.de/genus/pseudozobellia |
| Psittacicella | https://lpsn.dsmz.de/genus/psittacicella |
| Psychrilyobacter | https://lpsn.dsmz.de/genus/psychrilyobacter |
| Psychrobacillus | https://lpsn.dsmz.de/genus/psychrobacillus |
| Psychrobacter | https://lpsn.dsmz.de/genus/psychrobacter |
| Psychrobium | https://lpsn.dsmz.de/genus/psychrobium |
| Psychroflexus | https://lpsn.dsmz.de/genus/psychroflexus |
| Psychroglaciecola | https://lpsn.dsmz.de/genus/psychroglaciecola |
| Psychromarinibacter | https://lpsn.dsmz.de/genus/psychromarinibacter |
| Psychromicrobium | https://lpsn.dsmz.de/genus/psychromicrobium |
| Psychromonas | https://lpsn.dsmz.de/genus/psychromonas |
| Psychroserpens | https://lpsn.dsmz.de/genus/psychroserpens |
| Psychrosphaera | https://lpsn.dsmz.de/genus/psychrosphaera |
| Puerhibacterium | https://lpsn.dsmz.de/genus/puerhibacterium |
| Pueribacillus | https://lpsn.dsmz.de/genus/pueribacillus |
| Puia | https://lpsn.dsmz.de/genus/puia |
| Pukyongia | https://lpsn.dsmz.de/genus/pukyongia |
| Pukyongiella | https://lpsn.dsmz.de/genus/pukyongiella |
| Pullulanibacillus | https://lpsn.dsmz.de/genus/pullulanibacillus |
| Pulveribacter | https://lpsn.dsmz.de/genus/pulveribacter |
| Puniceibacterium | https://lpsn.dsmz.de/genus/puniceibacterium |
| Puniceicoccus | https://lpsn.dsmz.de/genus/puniceicoccus |
| Pusillibacter | https://lpsn.dsmz.de/genus/pusillibacter |
| Pusillimonas | https://lpsn.dsmz.de/genus/pusillimonas |
| Pusillimonas | https://lpsn.dsmz.de/genus/pusillimonas-1 |
| Pustulibacterium | https://lpsn.dsmz.de/genus/pustulibacterium |
| Puteibacter | https://lpsn.dsmz.de/genus/puteibacter |
| Putridiphycobacter | https://lpsn.dsmz.de/genus/putridiphycobacter |
| Puzihella | https://lpsn.dsmz.de/genus/puzihella |
| Pyramidobacter | https://lpsn.dsmz.de/genus/pyramidobacter |
| Pyrinomonas | https://lpsn.dsmz.de/genus/pyrinomonas |
| Pyrobaculum | https://lpsn.dsmz.de/genus/pyrobaculum |
| Pyrococcus | https://lpsn.dsmz.de/genus/pyrococcus |
| Pyrodictium | https://lpsn.dsmz.de/genus/pyrodictium |
| Pyrolobus | https://lpsn.dsmz.de/genus/pyrolobus |
| Pyruvatibacter | https://lpsn.dsmz.de/genus/pyruvatibacter |
| Pyxicoccus | https://lpsn.dsmz.de/genus/pyxicoccus |
| Pyxidicoccus | https://lpsn.dsmz.de/genus/pyxidicoccus |
| Qaidamihabitans | https://lpsn.dsmz.de/genus/qaidamihabitans |
| Qingshengfania | https://lpsn.dsmz.de/genus/qingshengfania |
| Qingshengfaniella | https://lpsn.dsmz.de/genus/qingshengfaniella |
| Qipengyuania | https://lpsn.dsmz.de/genus/qipengyuania |
| Quadricoccus | https://lpsn.dsmz.de/genus/quadricoccus |
| Quadrisphaera | https://lpsn.dsmz.de/genus/quadrisphaera |

| | |
|---|---|
| Quatrionicoccus | https://lpsn.dsmz.de/genus/quatrionicoccus |
| Quinella | https://lpsn.dsmz.de/genus/quinella |
| Quisquiliibacterium | https://lpsn.dsmz.de/genus/quisquiliibacterium |
| Racemicystis | https://lpsn.dsmz.de/genus/racemicystis |
| Radiobacillus | https://lpsn.dsmz.de/genus/radiobacillus |
| Rahnella | https://lpsn.dsmz.de/genus/rahnella |
| Raineya | https://lpsn.dsmz.de/genus/raineya |
| Raineyella | https://lpsn.dsmz.de/genus/raineyella |
| Ralstonia | https://lpsn.dsmz.de/genus/ralstonia |
| Ramlibacter | https://lpsn.dsmz.de/genus/ramlibacter |
| Raoultella | https://lpsn.dsmz.de/genus/raoultella |
| Raoultibacter | https://lpsn.dsmz.de/genus/raoultibacter |
| Rapidithrix | https://lpsn.dsmz.de/genus/rapidithrix |
| Rariglobus | https://lpsn.dsmz.de/genus/rariglobus |
| Rarimicrobium | https://lpsn.dsmz.de/genus/rarimicrobium |
| Rarobacter | https://lpsn.dsmz.de/genus/rarobacter |
| Rathayibacter | https://lpsn.dsmz.de/genus/rathayibacter |
| Ravibacter | https://lpsn.dsmz.de/genus/ravibacter |
| Rectinema | https://lpsn.dsmz.de/genus/rectinema |
| Rehaibacterium | https://lpsn.dsmz.de/genus/rehaibacterium |
| Reichenbachia | https://lpsn.dsmz.de/genus/reichenbachia |
| Reichenbachiella | https://lpsn.dsmz.de/genus/reichenbachiella |
| Reinekea | https://lpsn.dsmz.de/genus/reinekea |
| Renibacterium | https://lpsn.dsmz.de/genus/renibacterium |
| Reticulibacter | https://lpsn.dsmz.de/genus/reticulibacter |
| Reyranella | https://lpsn.dsmz.de/genus/reyranella |
| Rhabdaerophilum | https://lpsn.dsmz.de/genus/rhabdaerophilum |
| Rhabdanaerobium | https://lpsn.dsmz.de/genus/rhabdanaerobium |
| Rhabdobacter | https://lpsn.dsmz.de/genus/rhabdobacter |
| Rhabdochromatium | https://lpsn.dsmz.de/genus/rhabdochromatium |
| Rhabdothermincola | https://lpsn.dsmz.de/genus/rhabdothermincola |
| Rhabdothermus | https://lpsn.dsmz.de/genus/rhabdothermus |
| Rheinheimera | https://lpsn.dsmz.de/genus/rheinheimera |
| Rhizobacter | https://lpsn.dsmz.de/genus/rhizobacter |
| Rhizobium | https://lpsn.dsmz.de/genus/rhizobium |
| Rhizocola | https://lpsn.dsmz.de/genus/rhizocola |
| Rhizomicrobium | https://lpsn.dsmz.de/genus/rhizomicrobium |
| Rhizomonas | https://lpsn.dsmz.de/genus/rhizomonas |
| Rhizorhabdus | https://lpsn.dsmz.de/genus/rhizorhabdus |
| Rhizorhapis | https://lpsn.dsmz.de/genus/rhizorhapis |
| Rhodanobacter | https://lpsn.dsmz.de/genus/rhodanobacter |
| Rhodobaca | https://lpsn.dsmz.de/genus/rhodobaca |
| Rhodobacter | https://lpsn.dsmz.de/genus/rhodobacter |
| Rhodobaculum | https://lpsn.dsmz.de/genus/rhodobaculum |

| | |
|---|---|
| Rhodobium | https://lpsn.dsmz.de/genus/rhodobium |
| Rhodoblastus | https://lpsn.dsmz.de/genus/rhodoblastus |
| Rhodocaloribacter | https://lpsn.dsmz.de/genus/rhodocaloribacter |
| Rhodocista | https://lpsn.dsmz.de/genus/rhodocista |
| Rhodococcus | https://lpsn.dsmz.de/genus/rhodococcus |
| Rhodocyclus | https://lpsn.dsmz.de/genus/rhodocyclus |
| Rhodocytophaga | https://lpsn.dsmz.de/genus/rhodocytophaga |
| Rhodoferax | https://lpsn.dsmz.de/genus/rhodoferax |
| Rhodoglobus | https://lpsn.dsmz.de/genus/rhodoglobus |
| Rhodohalobacter | https://lpsn.dsmz.de/genus/rhodohalobacter |
| Rhodoligotrophos | https://lpsn.dsmz.de/genus/rhodoligotrophos |
| Rhodoluna | https://lpsn.dsmz.de/genus/rhodoluna |
| Rhodomicrobium | https://lpsn.dsmz.de/genus/rhodomicrobium |
| Rhodonellum | https://lpsn.dsmz.de/genus/rhodonellum |
| Rhodophyticola | https://lpsn.dsmz.de/genus/rhodophyticola |
| Rhodopila | https://lpsn.dsmz.de/genus/rhodopila |
| Rhodopirellula | https://lpsn.dsmz.de/genus/rhodopirellula |
| Rhodoplanes | https://lpsn.dsmz.de/genus/rhodoplanes |
| Rhodopseudomonas | https://lpsn.dsmz.de/genus/rhodopseudomonas |
| Rhodosalinus | https://lpsn.dsmz.de/genus/rhodosalinus |
| Rhodospira | https://lpsn.dsmz.de/genus/rhodospira |
| Rhodospirillum | https://lpsn.dsmz.de/genus/rhodospirillum |
| Rhodothalassium | https://lpsn.dsmz.de/genus/rhodothalassium |
| Rhodothermus | https://lpsn.dsmz.de/genus/rhodothermus |
| Rhodovarius | https://lpsn.dsmz.de/genus/rhodovarius |
| Rhodovastum | https://lpsn.dsmz.de/genus/rhodovastum |
| Rhodovibrio | https://lpsn.dsmz.de/genus/rhodovibrio |
| Rhodovulum | https://lpsn.dsmz.de/genus/rhodovulum |
| Rickettsia | https://lpsn.dsmz.de/genus/rickettsia |
| Rickettsiella | https://lpsn.dsmz.de/genus/rickettsiella |
| Riemerella | https://lpsn.dsmz.de/genus/riemerella |
| Rikenella | https://lpsn.dsmz.de/genus/rikenella |
| Risungbinella | https://lpsn.dsmz.de/genus/risungbinella |
| Rivibacter | https://lpsn.dsmz.de/genus/rivibacter |
| Rivicola | https://lpsn.dsmz.de/genus/rivicola |
| Robbsia | https://lpsn.dsmz.de/genus/robbsia |
| Robertkochia | https://lpsn.dsmz.de/genus/robertkochia |
| Robertmurraya | https://lpsn.dsmz.de/genus/robertmurraya |
| Robiginitalea | https://lpsn.dsmz.de/genus/robiginitalea |
| Robiginitomaculum | https://lpsn.dsmz.de/genus/robiginitomaculum |
| Robinsoniella | https://lpsn.dsmz.de/genus/robinsoniella |
| Rochalimaea | https://lpsn.dsmz.de/genus/rochalimaea |
| Rodentibacter | https://lpsn.dsmz.de/genus/rodentibacter |
| Romboutsia | https://lpsn.dsmz.de/genus/romboutsia |

| | |
|---|---|
| Romeriopsis | https://lpsn.dsmz.de/genus/romeriopsis |
| Roseateles | https://lpsn.dsmz.de/genus/roseateles |
| Roseburia | https://lpsn.dsmz.de/genus/roseburia |
| Roseiarcus | https://lpsn.dsmz.de/genus/roseiarcus |
| Roseibaca | https://lpsn.dsmz.de/genus/roseibaca |
| Roseibacillus | https://lpsn.dsmz.de/genus/roseibacillus |
| Roseibacterium | https://lpsn.dsmz.de/genus/roseibacterium |
| Roseibium | https://lpsn.dsmz.de/genus/roseibium |
| Roseicella | https://lpsn.dsmz.de/genus/roseicella |
| Roseicitreum | https://lpsn.dsmz.de/genus/roseicitreum |
| Roseicyclus | https://lpsn.dsmz.de/genus/roseicyclus |
| Roseiflexus | https://lpsn.dsmz.de/genus/roseiflexus |
| Roseimarinus | https://lpsn.dsmz.de/genus/roseimarinus |
| Roseimaritima | https://lpsn.dsmz.de/genus/roseimaritima |
| Roseimicrobium | https://lpsn.dsmz.de/genus/roseimicrobium |
| Roseinatronobacter | https://lpsn.dsmz.de/genus/roseinatronobacter |
| Roseisalinus | https://lpsn.dsmz.de/genus/roseisalinus |
| Roseisolibacter | https://lpsn.dsmz.de/genus/roseisolibacter |
| Roseitalea | https://lpsn.dsmz.de/genus/roseitalea |
| Roseithermus | https://lpsn.dsmz.de/genus/roseithermus |
| Roseivirga | https://lpsn.dsmz.de/genus/roseivirga |
| Roseivivax | https://lpsn.dsmz.de/genus/roseivivax |
| Rosenbergiella | https://lpsn.dsmz.de/genus/rosenbergiella |
| Roseobacter | https://lpsn.dsmz.de/genus/roseobacter |
| Roseococcus | https://lpsn.dsmz.de/genus/roseococcus |
| Roseomonas | https://lpsn.dsmz.de/genus/roseomonas |
| Roseospira | https://lpsn.dsmz.de/genus/roseospira |
| Roseovarius | https://lpsn.dsmz.de/genus/roseovarius |
| Rosistilla | https://lpsn.dsmz.de/genus/rosistilla |
| Rossellomorea | https://lpsn.dsmz.de/genus/rossellomorea |
| Rothia | https://lpsn.dsmz.de/genus/rothia |
| Rouxiella | https://lpsn.dsmz.de/genus/rouxiella |
| Ruania | https://lpsn.dsmz.de/genus/ruania |
| Rubellimicrobium | https://lpsn.dsmz.de/genus/rubellimicrobium |
| Rubidibacter | https://lpsn.dsmz.de/genus/rubidibacter |
| Rubinisphaera | https://lpsn.dsmz.de/genus/rubinisphaera |
| Rubneribacter | https://lpsn.dsmz.de/genus/rubneribacter |
| Rubribacterium | https://lpsn.dsmz.de/genus/rubribacterium |
| Rubricella | https://lpsn.dsmz.de/genus/rubricella |
| Rubricoccus | https://lpsn.dsmz.de/genus/rubricoccus |
| Rubrimonas | https://lpsn.dsmz.de/genus/rubrimonas |
| Rubripirellula | https://lpsn.dsmz.de/genus/rubripirellula |
| Rubritalea | https://lpsn.dsmz.de/genus/rubritalea |
| Rubritepida | https://lpsn.dsmz.de/genus/rubritepida |

| | |
|---|---|
| Rubrivirga | https://lpsn.dsmz.de/genus/rubrivirga |
| Rubrivivax | https://lpsn.dsmz.de/genus/rubrivivax |
| Rubrobacter | https://lpsn.dsmz.de/genus/rubrobacter |
| Ructibacterium | https://lpsn.dsmz.de/genus/ructibacterium |
| Rudaea | https://lpsn.dsmz.de/genus/rudaea |
| Rudaeicoccus | https://lpsn.dsmz.de/genus/rudaeicoccus |
| Rudaibacter | https://lpsn.dsmz.de/genus/rudaibacter |
| Rudanella | https://lpsn.dsmz.de/genus/rudanella |
| Ruegeria | https://lpsn.dsmz.de/genus/ruegeria |
| Rufibacter | https://lpsn.dsmz.de/genus/rufibacter |
| Ruficoccus | https://lpsn.dsmz.de/genus/ruficoccus |
| Rugamonas | https://lpsn.dsmz.de/genus/rugamonas |
| Rugosibacter | https://lpsn.dsmz.de/genus/rugosibacter |
| Rugosimonospora | https://lpsn.dsmz.de/genus/rugosimonospora |
| Ruminiclostridium | https://lpsn.dsmz.de/genus/ruminiclostridium |
| Ruminobacter | https://lpsn.dsmz.de/genus/ruminobacter |
| Ruminococcoides | https://lpsn.dsmz.de/genus/ruminococcoides |
| Ruminococcus | https://lpsn.dsmz.de/genus/ruminococcus |
| Rummeliibacillus | https://lpsn.dsmz.de/genus/rummeliibacillus |
| Runella | https://lpsn.dsmz.de/genus/runella |
| Ruoffia | https://lpsn.dsmz.de/genus/ruoffia |
| Rurimicrobium | https://lpsn.dsmz.de/genus/rurimicrobium |
| Ruthenibacterium | https://lpsn.dsmz.de/genus/ruthenibacterium |
| Sabulibacter | https://lpsn.dsmz.de/genus/sabulibacter |
| Sabulilitoribacter | https://lpsn.dsmz.de/genus/sabulilitoribacter |
| Saccharedens | https://lpsn.dsmz.de/genus/saccharedens |
| Saccharibacillus | https://lpsn.dsmz.de/genus/saccharibacillus |
| Saccharibacter | https://lpsn.dsmz.de/genus/saccharibacter |
| Saccharicrinis | https://lpsn.dsmz.de/genus/saccharicrinis |
| Saccharobacter | https://lpsn.dsmz.de/genus/saccharobacter |
| Saccharobesus | https://lpsn.dsmz.de/genus/saccharobesus |
| Saccharococcus | https://lpsn.dsmz.de/genus/saccharococcus |
| Saccharofermentans | https://lpsn.dsmz.de/genus/saccharofermentans |
| Saccharolobus | https://lpsn.dsmz.de/genus/saccharolobus |
| Saccharomonospora | https://lpsn.dsmz.de/genus/saccharomonospora |
| Saccharophagus | https://lpsn.dsmz.de/genus/saccharophagus |
| Saccharopolyspora | https://lpsn.dsmz.de/genus/saccharopolyspora |
| Saccharospirillum | https://lpsn.dsmz.de/genus/saccharospirillum |
| Saccharothrix | https://lpsn.dsmz.de/genus/saccharothrix |
| Saezia | https://lpsn.dsmz.de/genus/saezia |
| Sagittula | https://lpsn.dsmz.de/genus/sagittula |
| Salana | https://lpsn.dsmz.de/genus/salana |
| Salaquimonas | https://lpsn.dsmz.de/genus/salaquimonas |
| Salarchaeum | https://lpsn.dsmz.de/genus/salarchaeum |

| | |
|---|---|
| Salegentibacter | https://lpsn.dsmz.de/genus/salegentibacter |
| Salibacillus | https://lpsn.dsmz.de/genus/salibacillus |
| Salibacter | https://lpsn.dsmz.de/genus/salibacter |
| Salibacterium | https://lpsn.dsmz.de/genus/salibacterium |
| Salibaculum | https://lpsn.dsmz.de/genus/salibaculum |
| Salicibibacter | https://lpsn.dsmz.de/genus/salicibibacter |
| Salicola | https://lpsn.dsmz.de/genus/salicola |
| Salifodinibacter | https://lpsn.dsmz.de/genus/salifodinibacter |
| Salilacibacter | https://lpsn.dsmz.de/genus/salilacibacter |
| Salimesophilobacter | https://lpsn.dsmz.de/genus/salimesophilobacter |
| Salimicrobium | https://lpsn.dsmz.de/genus/salimicrobium |
| Salinactinospora | https://lpsn.dsmz.de/genus/salinactinospora |
| Salinadaptatus | https://lpsn.dsmz.de/genus/salinadaptatus |
| Salinarchaeum | https://lpsn.dsmz.de/genus/salinarchaeum |
| Salinarimonas | https://lpsn.dsmz.de/genus/salinarimonas |
| Salinarubrum | https://lpsn.dsmz.de/genus/salinarubrum |
| Salinibacillus | https://lpsn.dsmz.de/genus/salinibacillus |
| Salinibacter | https://lpsn.dsmz.de/genus/salinibacter |
| Salinibacterium | https://lpsn.dsmz.de/genus/salinibacterium |
| Salinibaculum | https://lpsn.dsmz.de/genus/salinibaculum |
| Salinibius | https://lpsn.dsmz.de/genus/salinibius |
| Salinicoccus | https://lpsn.dsmz.de/genus/salinicoccus |
| Salinicola | https://lpsn.dsmz.de/genus/salinicola |
| Salinifilum | https://lpsn.dsmz.de/genus/salinifilum |
| Salinigranum | https://lpsn.dsmz.de/genus/salinigranum |
| Salinihabitans | https://lpsn.dsmz.de/genus/salinihabitans |
| Salinimicrobium | https://lpsn.dsmz.de/genus/salinimicrobium |
| Salinimonas | https://lpsn.dsmz.de/genus/salinimonas |
| Salininema | https://lpsn.dsmz.de/genus/salininema |
| Saliniradius | https://lpsn.dsmz.de/genus/saliniradius |
| Saliniramus | https://lpsn.dsmz.de/genus/saliniramus |
| Salinirepens | https://lpsn.dsmz.de/genus/salinirepens |
| Salinirubellus | https://lpsn.dsmz.de/genus/salinirubellus |
| Salinirubrum | https://lpsn.dsmz.de/genus/salinirubrum |
| Salinirussus | https://lpsn.dsmz.de/genus/salinirussus |
| Salinisphaera | https://lpsn.dsmz.de/genus/salinisphaera |
| Salinispira | https://lpsn.dsmz.de/genus/salinispira |
| Salinispirillum | https://lpsn.dsmz.de/genus/salinispirillum |
| Salinispora | https://lpsn.dsmz.de/genus/salinispora |
| Salinithrix | https://lpsn.dsmz.de/genus/salinithrix |
| Salinivenus | https://lpsn.dsmz.de/genus/salinivenus |
| Salinivibrio | https://lpsn.dsmz.de/genus/salinivibrio |
| Salinivirga | https://lpsn.dsmz.de/genus/salinivirga-1 |
| Salinivirga | https://lpsn.dsmz.de/genus/salinivirga |

| | |
|---|---|
| Salinovum | https://lpsn.dsmz.de/genus/salinovum |
| Salipaludibacillus | https://lpsn.dsmz.de/genus/salipaludibacillus |
| Saliphagus | https://lpsn.dsmz.de/genus/saliphagus |
| Salipiger | https://lpsn.dsmz.de/genus/salipiger |
| Salirhabdus | https://lpsn.dsmz.de/genus/salirhabdus |
| Salisaeta | https://lpsn.dsmz.de/genus/salisaeta |
| Salisediminibacterium | https://lpsn.dsmz.de/genus/salisediminibacterium |
| Saliterribacillus | https://lpsn.dsmz.de/genus/saliterribacillus |
| Salmonella | https://lpsn.dsmz.de/genus/salmonella |
| Salsipaludibacter | https://lpsn.dsmz.de/genus/salsipaludibacter |
| Salsuginibacillus | https://lpsn.dsmz.de/genus/salsuginibacillus |
| Salsuginimonas | https://lpsn.dsmz.de/genus/salsuginimonas |
| Samsonia | https://lpsn.dsmz.de/genus/samsonia |
| Sandaracinobacteroides | https://lpsn.dsmz.de/genus/sandaracinobacteroides |
| Sandaracinomonas | https://lpsn.dsmz.de/genus/sandaracinomonas |
| Sandaracinus | https://lpsn.dsmz.de/genus/sandaracinus |
| Sandarakinorhabdus | https://lpsn.dsmz.de/genus/sandarakinorhabdus |
| Sandarakinotalea | https://lpsn.dsmz.de/genus/sandarakinotalea |
| Sangeribacter | https://lpsn.dsmz.de/genus/sangeribacter |
| Sanguibacter | https://lpsn.dsmz.de/genus/sanguibacter |
| Sansalvadorimonas | https://lpsn.dsmz.de/genus/sansalvadorimonas |
| Saonia | https://lpsn.dsmz.de/genus/saonia |
| Sapientia | https://lpsn.dsmz.de/genus/sapientia |
| Saprospira | https://lpsn.dsmz.de/genus/saprospira |
| Sarcina | https://lpsn.dsmz.de/genus/sarcina |
| Sarcobium | https://lpsn.dsmz.de/genus/sarcobium |
| Savagea | https://lpsn.dsmz.de/genus/savagea |
| Saxeibacter | https://lpsn.dsmz.de/genus/saxeibacter |
| Scandinavium | https://lpsn.dsmz.de/genus/scandinavium |
| Scardovia | https://lpsn.dsmz.de/genus/scardovia |
| Schaalia | https://lpsn.dsmz.de/genus/schaalia |
| Schaedlerella | https://lpsn.dsmz.de/genus/schaedlerella |
| Schineria | https://lpsn.dsmz.de/genus/schineria |
| Schinkia | https://lpsn.dsmz.de/genus/schinkia |
| Schlegelella | https://lpsn.dsmz.de/genus/schlegelella |
| Schleiferia | https://lpsn.dsmz.de/genus/schleiferia |
| Schleiferilactobacillus | https://lpsn.dsmz.de/genus/schleiferilactobacillus |
| Schlesneria | https://lpsn.dsmz.de/genus/schlesneria |
| Schnuerera | https://lpsn.dsmz.de/genus/schnuerera |
| Schumannella | https://lpsn.dsmz.de/genus/schumannella |
| Schwartzia | https://lpsn.dsmz.de/genus/schwartzia |
| Sciscionella | https://lpsn.dsmz.de/genus/sciscionella |
| Scopulibacillus | https://lpsn.dsmz.de/genus/scopulibacillus |
| Scrofimicrobium | https://lpsn.dsmz.de/genus/scrofimicrobium |

| | |
|---|---|
| Sebaldella | https://lpsn.dsmz.de/genus/sebaldella |
| Secundilactobacillus | https://lpsn.dsmz.de/genus/secundilactobacillus |
| Sedimentibacter | https://lpsn.dsmz.de/genus/sedimentibacter |
| Sedimenticola | https://lpsn.dsmz.de/genus/sedimenticola |
| Sedimentisphaera | https://lpsn.dsmz.de/genus/sedimentisphaera |
| Sedimentitalea | https://lpsn.dsmz.de/genus/sedimentitalea |
| Sediminibacillus | https://lpsn.dsmz.de/genus/sediminibacillus |
| Sediminibacter | https://lpsn.dsmz.de/genus/sediminibacter |
| Sediminibacterium | https://lpsn.dsmz.de/genus/sediminibacterium |
| Sediminicola | https://lpsn.dsmz.de/genus/sediminicola |
| Sediminihabitans | https://lpsn.dsmz.de/genus/sediminihabitans |
| Sediminihaliea | https://lpsn.dsmz.de/genus/sediminihaliea |
| Sediminimonas | https://lpsn.dsmz.de/genus/sediminimonas |
| Sediminispirochaeta | https://lpsn.dsmz.de/genus/sediminispirochaeta |
| Sediminitomix | https://lpsn.dsmz.de/genus/sediminitomix |
| Sediminivirga | https://lpsn.dsmz.de/genus/sediminivirga |
| Segetibacter | https://lpsn.dsmz.de/genus/segetibacter |
| Segeticoccus | https://lpsn.dsmz.de/genus/segeticoccus |
| Segniliparus | https://lpsn.dsmz.de/genus/segniliparus |
| Segnochrobactrum | https://lpsn.dsmz.de/genus/segnochrobactrum |
| Seinonella | https://lpsn.dsmz.de/genus/seinonella |
| Sejongia | https://lpsn.dsmz.de/genus/sejongia |
| Seleniibacterium | https://lpsn.dsmz.de/genus/seleniibacterium |
| Seleniivibrio | https://lpsn.dsmz.de/genus/seleniivibrio |
| Selenomonas | https://lpsn.dsmz.de/genus/selenomonas |
| Seliberia | https://lpsn.dsmz.de/genus/seliberia |
| Sellimonas | https://lpsn.dsmz.de/genus/sellimonas |
| Seminibacterium | https://lpsn.dsmz.de/genus/seminibacterium |
| Senegalemassilia | https://lpsn.dsmz.de/genus/senegalemassilia |
| Senegalimassilia | https://lpsn.dsmz.de/genus/senegalimassilia |
| Seohaeicola | https://lpsn.dsmz.de/genus/seohaeicola |
| Seonamhaeicola | https://lpsn.dsmz.de/genus/seonamhaeicola |
| Seongchinamella | https://lpsn.dsmz.de/genus/seongchinamella |
| Seramator | https://lpsn.dsmz.de/genus/seramator |
| Serinibacter | https://lpsn.dsmz.de/genus/serinibacter |
| Serinicoccus | https://lpsn.dsmz.de/genus/serinicoccus |
| Serpens | https://lpsn.dsmz.de/genus/serpens |
| Serpentinicella | https://lpsn.dsmz.de/genus/serpentinicella |
| Serpentinimonas | https://lpsn.dsmz.de/genus/serpentinimonas |
| Serpula | https://lpsn.dsmz.de/genus/serpula |
| Serpulina | https://lpsn.dsmz.de/genus/serpulina |
| Serratia | https://lpsn.dsmz.de/genus/serratia |
| Sessilibacter | https://lpsn.dsmz.de/genus/sessilibacter |
| Sharpea | https://lpsn.dsmz.de/genus/sharpea |

| | |
|---|---|
| Sheuella | https://lpsn.dsmz.de/genus/sheuella |
| Shewanella | https://lpsn.dsmz.de/genus/shewanella |
| Shigella | https://lpsn.dsmz.de/genus/shigella |
| Shimazuella | https://lpsn.dsmz.de/genus/shimazuella |
| Shimia | https://lpsn.dsmz.de/genus/shimia |
| Shimwellia | https://lpsn.dsmz.de/genus/shimwellia |
| Shinella | https://lpsn.dsmz.de/genus/shinella |
| Shivajiella | https://lpsn.dsmz.de/genus/shivajiella |
| Shouchella | https://lpsn.dsmz.de/genus/shouchella |
| Shuttleworthia | https://lpsn.dsmz.de/genus/shuttleworthia |
| Siansivirga | https://lpsn.dsmz.de/genus/siansivirga |
| Siccibacter | https://lpsn.dsmz.de/genus/siccibacter |
| Siccirubricoccus | https://lpsn.dsmz.de/genus/siccirubricoccus |
| Siculibacillus | https://lpsn.dsmz.de/genus/siculibacillus |
| Sideroxyarcus | https://lpsn.dsmz.de/genus/sideroxyarcus |
| Silanimonas | https://lpsn.dsmz.de/genus/silanimonas |
| Silicibacter | https://lpsn.dsmz.de/genus/silicibacter |
| Silicimonas | https://lpsn.dsmz.de/genus/silicimonas |
| Silvanigrella | https://lpsn.dsmz.de/genus/silvanigrella |
| Silvibacterium | https://lpsn.dsmz.de/genus/silvibacterium |
| Silvimonas | https://lpsn.dsmz.de/genus/silvimonas |
| Simiduia | https://lpsn.dsmz.de/genus/simiduia |
| Siminovitchia | https://lpsn.dsmz.de/genus/siminovitchia |
| Simkania | https://lpsn.dsmz.de/genus/simkania |
| Simonsiella | https://lpsn.dsmz.de/genus/simonsiella |
| Simplicispira | https://lpsn.dsmz.de/genus/simplicispira |
| Simulacricoccus | https://lpsn.dsmz.de/genus/simulacricoccus |
| Singularimonas | https://lpsn.dsmz.de/genus/singularimonas |
| Singulisphaera | https://lpsn.dsmz.de/genus/singulisphaera |
| Sinibacillus | https://lpsn.dsmz.de/genus/sinibacillus |
| Sinimarinibacterium | https://lpsn.dsmz.de/genus/sinimarinibacterium |
| Sinirhodobacter | https://lpsn.dsmz.de/genus/sinirhodobacter |
| Sinisalibacter | https://lpsn.dsmz.de/genus/sinisalibacter |
| Sinobaca | https://lpsn.dsmz.de/genus/sinobaca |
| Sinobacter | https://lpsn.dsmz.de/genus/sinobacter |
| Sinobacterium | https://lpsn.dsmz.de/genus/sinobacterium |
| Sinococcus | https://lpsn.dsmz.de/genus/sinococcus |
| Sinomicrobium | https://lpsn.dsmz.de/genus/sinomicrobium |
| Sinomonas | https://lpsn.dsmz.de/genus/sinomonas |
| Sinorhizobium | https://lpsn.dsmz.de/genus/sinorhizobium |
| Sinorhodobacter | https://lpsn.dsmz.de/genus/sinorhodobacter |
| Sinosporangium | https://lpsn.dsmz.de/genus/sinosporangium |
| Siphonobacter | https://lpsn.dsmz.de/genus/siphonobacter |
| Skermanella | https://lpsn.dsmz.de/genus/skermanella |

| | |
|---|---|
| Skermania | https://lpsn.dsmz.de/genus/skermania |
| Slackia | https://lpsn.dsmz.de/genus/slackia |
| Smaragdicoccus | https://lpsn.dsmz.de/genus/smaragdicoccus |
| Smithella | https://lpsn.dsmz.de/genus/smithella |
| Sneathia | https://lpsn.dsmz.de/genus/sneathia |
| Sneathiella | https://lpsn.dsmz.de/genus/sneathiella |
| Snodgrassella | https://lpsn.dsmz.de/genus/snodgrassella |
| Snuella | https://lpsn.dsmz.de/genus/snuella |
| Sodaliphilus | https://lpsn.dsmz.de/genus/sodaliphilus |
| Sodalis | https://lpsn.dsmz.de/genus/sodalis |
| Soehngenia | https://lpsn.dsmz.de/genus/soehngenia |
| Solibacillus | https://lpsn.dsmz.de/genus/solibacillus |
| Solibaculum | https://lpsn.dsmz.de/genus/solibaculum |
| Solidesulfovibrio | https://lpsn.dsmz.de/genus/solidesulfovibrio |
| Solihabitans | https://lpsn.dsmz.de/genus/solihabitans |
| Solimicrobium | https://lpsn.dsmz.de/genus/solimicrobium |
| Solimonas | https://lpsn.dsmz.de/genus/solimonas |
| Solirhodobacter | https://lpsn.dsmz.de/genus/solirhodobacter |
| Solirubrobacter | https://lpsn.dsmz.de/genus/solirubrobacter |
| Solitalea | https://lpsn.dsmz.de/genus/solitalea |
| Solobacterium | https://lpsn.dsmz.de/genus/solobacterium |
| Soonwooa | https://lpsn.dsmz.de/genus/soonwooa |
| Soortia | https://lpsn.dsmz.de/genus/soortia |
| Sorangium | https://lpsn.dsmz.de/genus/sorangium |
| Spartinivicinus | https://lpsn.dsmz.de/genus/spartinivicinus |
| Specibacter | https://lpsn.dsmz.de/genus/specibacter |
| Spelaeibacter | https://lpsn.dsmz.de/genus/spelaeibacter |
| Spelaeicoccus | https://lpsn.dsmz.de/genus/spelaeicoccus |
| Sphaerimonospora | https://lpsn.dsmz.de/genus/sphaerimonospora |
| Sphaerisporangium | https://lpsn.dsmz.de/genus/sphaerisporangium |
| Sphaerobacter | https://lpsn.dsmz.de/genus/sphaerobacter |
| Sphaerochaeta | https://lpsn.dsmz.de/genus/sphaerochaeta |
| Sphaerosporangium | https://lpsn.dsmz.de/genus/sphaerosporangium |
| Sphaerotilus | https://lpsn.dsmz.de/genus/sphaerotilus |
| Sphingoaurantiacus | https://lpsn.dsmz.de/genus/sphingoaurantiacus |
| Sphingobacterium | https://lpsn.dsmz.de/genus/sphingobacterium |
| Sphingobium | https://lpsn.dsmz.de/genus/sphingobium |
| Sphingomicrobium | https://lpsn.dsmz.de/genus/sphingomicrobium |
| Sphingomonas | https://lpsn.dsmz.de/genus/sphingomonas |
| Sphingopyxis | https://lpsn.dsmz.de/genus/sphingopyxis |
| Sphingorhabdus | https://lpsn.dsmz.de/genus/sphingorhabdus |
| Sphingosinicella | https://lpsn.dsmz.de/genus/sphingosinicella |
| Sphingosinithalassobacter | https://lpsn.dsmz.de/genus/sphingosinithalassobacter |
| Spinactinospora | https://lpsn.dsmz.de/genus/spinactinospora |

| | |
|---|---|
| Spirabiliibacterium | https://lpsn.dsmz.de/genus/spirabiliibacterium |
| Spiractinospora | https://lpsn.dsmz.de/genus/spiractinospora |
| Spiribacter | https://lpsn.dsmz.de/genus/spiribacter |
| Spirilliplanes | https://lpsn.dsmz.de/genus/spirilliplanes |
| Spirillospora | https://lpsn.dsmz.de/genus/spirillospora |
| Spirillum | https://lpsn.dsmz.de/genus/spirillum |
| Spirochaeta | https://lpsn.dsmz.de/genus/spirochaeta |
| Spiroplasma | https://lpsn.dsmz.de/genus/spiroplasma |
| Spirosoma | https://lpsn.dsmz.de/genus/spirosoma |
| Spirulina | https://lpsn.dsmz.de/genus/spirulina |
| Spodiobacter | https://lpsn.dsmz.de/genus/spodiobacter |
| Spongiactinospora | https://lpsn.dsmz.de/genus/spongiactinospora |
| Spongiibacter | https://lpsn.dsmz.de/genus/spongiibacter |
| Spongiibacterium | https://lpsn.dsmz.de/genus/spongiibacterium |
| Spongiiferula | https://lpsn.dsmz.de/genus/spongiiferula |
| Spongiimicrobium | https://lpsn.dsmz.de/genus/spongiimicrobium |
| Spongiimonas | https://lpsn.dsmz.de/genus/spongiimonas |
| Spongiispira | https://lpsn.dsmz.de/genus/spongiispira |
| Spongiivirga | https://lpsn.dsmz.de/genus/spongiivirga |
| Sporacetigenium | https://lpsn.dsmz.de/genus/sporacetigenium |
| Sporanaerobacter | https://lpsn.dsmz.de/genus/sporanaerobacter |
| Sporichthya | https://lpsn.dsmz.de/genus/sporichthya |
| Sporobacter | https://lpsn.dsmz.de/genus/sporobacter |
| Sporobacterium | https://lpsn.dsmz.de/genus/sporobacterium |
| Sporocytophaga | https://lpsn.dsmz.de/genus/sporocytophaga |
| Sporofaciens | https://lpsn.dsmz.de/genus/sporofaciens |
| Sporohalobacter | https://lpsn.dsmz.de/genus/sporohalobacter |
| Sporolactobacillus | https://lpsn.dsmz.de/genus/sporolactobacillus |
| Sporolituus | https://lpsn.dsmz.de/genus/sporolituus |
| Sporomusa | https://lpsn.dsmz.de/genus/sporomusa |
| Sporosalibacterium | https://lpsn.dsmz.de/genus/sporosalibacterium |
| Sporosarcina | https://lpsn.dsmz.de/genus/sporosarcina |
| Sporotalea | https://lpsn.dsmz.de/genus/sporotalea |
| Sporotomaculum | https://lpsn.dsmz.de/genus/sporotomaculum |
| Stackebrandtia | https://lpsn.dsmz.de/genus/stackebrandtia |
| Stagnihabitans | https://lpsn.dsmz.de/genus/stagnihabitans |
| Stagnimonas | https://lpsn.dsmz.de/genus/stagnimonas |
| Stakelama | https://lpsn.dsmz.de/genus/stakelama |
| Staleya | https://lpsn.dsmz.de/genus/staleya |
| Stanierella | https://lpsn.dsmz.de/genus/stanierella |
| Staphylococcus | https://lpsn.dsmz.de/genus/staphylococcus |
| Staphylospora | https://lpsn.dsmz.de/genus/staphylospora |
| Staphylothermus | https://lpsn.dsmz.de/genus/staphylothermus |
| Stappia | https://lpsn.dsmz.de/genus/stappia |

| | |
|---|---|
| Starkeya | https://lpsn.dsmz.de/genus/starkeya |
| Stecheria | https://lpsn.dsmz.de/genus/stecheria |
| Stella | https://lpsn.dsmz.de/genus/stella |
| Stenothermobacter | https://lpsn.dsmz.de/genus/stenothermobacter |
| Stenotrophobacter | https://lpsn.dsmz.de/genus/stenotrophobacter |
| Stenotrophobium | https://lpsn.dsmz.de/genus/stenotrophobium |
| Stenotrophomonas | https://lpsn.dsmz.de/genus/stenotrophomonas |
| Stenoxybacter | https://lpsn.dsmz.de/genus/stenoxybacter |
| Steroidobacter | https://lpsn.dsmz.de/genus/steroidobacter |
| Sterolibacterium | https://lpsn.dsmz.de/genus/sterolibacterium |
| Stetteria | https://lpsn.dsmz.de/genus/stetteria |
| Stibiobacter | https://lpsn.dsmz.de/genus/stibiobacter |
| Stieleria | https://lpsn.dsmz.de/genus/stieleria |
| Stigmatella | https://lpsn.dsmz.de/genus/stigmatella |
| Stomatobaculum | https://lpsn.dsmz.de/genus/stomatobaculum |
| Stomatococcus | https://lpsn.dsmz.de/genus/stomatococcus |
| Streptacidiphilus | https://lpsn.dsmz.de/genus/streptacidiphilus |
| Streptantibioticus | https://lpsn.dsmz.de/genus/streptantibioticus |
| Streptimonospora | https://lpsn.dsmz.de/genus/streptimonospora |
| Streptoalloteichus | https://lpsn.dsmz.de/genus/streptoalloteichus |
| Streptobacillus | https://lpsn.dsmz.de/genus/streptobacillus |
| Streptococcus | https://lpsn.dsmz.de/genus/streptococcus |
| Streptohalobacillus | https://lpsn.dsmz.de/genus/streptohalobacillus |
| Streptomonospora | https://lpsn.dsmz.de/genus/streptomonospora |
| Streptomyces | https://lpsn.dsmz.de/genus/streptomyces |
| Streptosporangium | https://lpsn.dsmz.de/genus/streptosporangium |
| Streptoverticillium | https://lpsn.dsmz.de/genus/streptoverticillium |
| Stutzerimonas | https://lpsn.dsmz.de/genus/stutzerimonas |
| Stygiolobus | https://lpsn.dsmz.de/genus/stygiolobus |
| Subdoligranulum | https://lpsn.dsmz.de/genus/subdoligranulum |
| Subsaxibacter | https://lpsn.dsmz.de/genus/subsaxibacter |
| Subsaximicrobium | https://lpsn.dsmz.de/genus/subsaximicrobium |
| Subtercola | https://lpsn.dsmz.de/genus/subtercola |
| Succinatimonas | https://lpsn.dsmz.de/genus/succinatimonas |
| Succiniclasticum | https://lpsn.dsmz.de/genus/succiniclasticum |
| Succinimonas | https://lpsn.dsmz.de/genus/succinimonas |
| Succinispira | https://lpsn.dsmz.de/genus/succinispira |
| Succinivibrio | https://lpsn.dsmz.de/genus/succinivibrio |
| Suicoccus | https://lpsn.dsmz.de/genus/suicoccus |
| Suilimivivens | https://lpsn.dsmz.de/genus/suilimivivens |
| Suipraeoptans | https://lpsn.dsmz.de/genus/suipraeoptans |
| Suipraoptans | https://lpsn.dsmz.de/genus/suipraoptans |
| Sulfitobacter | https://lpsn.dsmz.de/genus/sulfitobacter |
| Sulfobacillus | https://lpsn.dsmz.de/genus/sulfobacillus |

| | |
|---|---|
| Sulfodiicoccus | https://lpsn.dsmz.de/genus/sulfodiicoccus |
| Sulfolobus | https://lpsn.dsmz.de/genus/sulfolobus |
| Sulfophobococcus | https://lpsn.dsmz.de/genus/sulfophobococcus |
| Sulfuracidifex | https://lpsn.dsmz.de/genus/sulfuracidifex |
| Sulfuricaulis | https://lpsn.dsmz.de/genus/sulfuricaulis |
| Sulfuricella | https://lpsn.dsmz.de/genus/sulfuricella |
| Sulfuricurvum | https://lpsn.dsmz.de/genus/sulfuricurvum |
| Sulfuriferula | https://lpsn.dsmz.de/genus/sulfuriferula |
| Sulfuriflexus | https://lpsn.dsmz.de/genus/sulfuriflexus |
| Sulfurifustis | https://lpsn.dsmz.de/genus/sulfurifustis |
| Sulfurihydrogenibium | https://lpsn.dsmz.de/genus/sulfurihydrogenibium |
| Sulfurimicrobium | https://lpsn.dsmz.de/genus/sulfurimicrobium |
| Sulfurimonas | https://lpsn.dsmz.de/genus/sulfurimonas |
| Sulfurirhabdus | https://lpsn.dsmz.de/genus/sulfurirhabdus |
| Sulfurisoma | https://lpsn.dsmz.de/genus/sulfurisoma |
| Sulfurisphaera | https://lpsn.dsmz.de/genus/sulfurisphaera |
| Sulfuritalea | https://lpsn.dsmz.de/genus/sulfuritalea |
| Sulfuritortus | https://lpsn.dsmz.de/genus/sulfuritortus |
| Sulfurivermis | https://lpsn.dsmz.de/genus/sulfurivermis |
| Sulfurivirga | https://lpsn.dsmz.de/genus/sulfurivirga |
| Sulfurococcus | https://lpsn.dsmz.de/genus/sulfurococcus |
| Sulfurospirillum | https://lpsn.dsmz.de/genus/sulfurospirillum |
| Sulfurovum | https://lpsn.dsmz.de/genus/sulfurovum |
| Sungkyunkwania | https://lpsn.dsmz.de/genus/sungkyunkwania |
| Sunxiuqinia | https://lpsn.dsmz.de/genus/sunxiuqinia |
| Suonthocola | https://lpsn.dsmz.de/genus/suonthocola |
| Sutcliffiella | https://lpsn.dsmz.de/genus/sutcliffiella |
| Sutterella | https://lpsn.dsmz.de/genus/sutterella |
| Suttonella | https://lpsn.dsmz.de/genus/suttonella |
| Swaminathania | https://lpsn.dsmz.de/genus/swaminathania |
| Swingsia | https://lpsn.dsmz.de/genus/swingsia |
| Swionibacillus | https://lpsn.dsmz.de/genus/swionibacillus |
| Symbiobacterium | https://lpsn.dsmz.de/genus/symbiobacterium |
| Symbiopectobacterium | https://lpsn.dsmz.de/genus/symbiopectobacterium |
| Symbiotes | https://lpsn.dsmz.de/genus/symbiotes |
| Symmachiella | https://lpsn.dsmz.de/genus/symmachiella |
| Synechococcus | https://lpsn.dsmz.de/genus/synechococcus |
| Synergistes | https://lpsn.dsmz.de/genus/synergistes |
| Syntrophaceticus | https://lpsn.dsmz.de/genus/syntrophaceticus |
| Syntrophobacter | https://lpsn.dsmz.de/genus/syntrophobacter |
| Syntrophobacterium | https://lpsn.dsmz.de/genus/syntrophobacterium |
| Syntrophobotulus | https://lpsn.dsmz.de/genus/syntrophobotulus |
| Syntrophococcus | https://lpsn.dsmz.de/genus/syntrophococcus |
| Syntrophomonas | https://lpsn.dsmz.de/genus/syntrophomonas |

| | |
|---|---|
| Syntrophorhabdus | https://lpsn.dsmz.de/genus/syntrophorhabdus |
| Syntrophospora | https://lpsn.dsmz.de/genus/syntrophospora |
| Syntrophotalea | https://lpsn.dsmz.de/genus/syntrophotalea |
| Syntrophothermus | https://lpsn.dsmz.de/genus/syntrophothermus |
| Syntrophus | https://lpsn.dsmz.de/genus/syntrophus |
| Szabonella | https://lpsn.dsmz.de/genus/szabonella |
| Tabrizicola | https://lpsn.dsmz.de/genus/tabrizicola |
| Taeania | https://lpsn.dsmz.de/genus/taeania |
| Taeseokella | https://lpsn.dsmz.de/genus/taeseokella |
| Tagaea | https://lpsn.dsmz.de/genus/tagaea |
| Tahibacter | https://lpsn.dsmz.de/genus/tahibacter |
| Taibaiella | https://lpsn.dsmz.de/genus/taibaiella |
| Tamaricihabitans | https://lpsn.dsmz.de/genus/tamaricihabitans |
| Tamilnaduibacter | https://lpsn.dsmz.de/genus/tamilnaduibacter |
| Tamlana | https://lpsn.dsmz.de/genus/tamlana |
| Tamlicoccus | https://lpsn.dsmz.de/genus/tamlicoccus |
| Tangfeifania | https://lpsn.dsmz.de/genus/tangfeifania |
| Tannerella | https://lpsn.dsmz.de/genus/tannerella |
| Tannockella | https://lpsn.dsmz.de/genus/tannockella |
| Tanticharoenia | https://lpsn.dsmz.de/genus/tanticharoenia |
| Taonella | https://lpsn.dsmz.de/genus/taonella |
| Tardiphaga | https://lpsn.dsmz.de/genus/tardiphaga |
| Tateyamaria | https://lpsn.dsmz.de/genus/tateyamaria |
| Tatlockia | https://lpsn.dsmz.de/genus/tatlockia |
| Tatumella | https://lpsn.dsmz.de/genus/tatumella |
| Tautonia | https://lpsn.dsmz.de/genus/tautonia |
| Taylorella | https://lpsn.dsmz.de/genus/taylorella |
| Tectibacter | https://lpsn.dsmz.de/genus/tectibacter |
| Teichococcus | https://lpsn.dsmz.de/genus/teichococcus |
| Telluria | https://lpsn.dsmz.de/genus/telluria |
| Telluribacter | https://lpsn.dsmz.de/genus/telluribacter |
| Tellurimicrobium | https://lpsn.dsmz.de/genus/tellurimicrobium |
| Tellurirhabdus | https://lpsn.dsmz.de/genus/tellurirhabdus |
| Telmatobacter | https://lpsn.dsmz.de/genus/telmatobacter |
| Telmatocola | https://lpsn.dsmz.de/genus/telmatocola |
| Telmatospirillum | https://lpsn.dsmz.de/genus/telmatospirillum |
| Temperatibacter | https://lpsn.dsmz.de/genus/temperatibacter |
| Tenacibaculum | https://lpsn.dsmz.de/genus/tenacibaculum |
| Tenebrionibacter | https://lpsn.dsmz.de/genus/tenebrionibacter |
| Tenebrionicola | https://lpsn.dsmz.de/genus/tenebrionicola |
| Tenggerimyces | https://lpsn.dsmz.de/genus/tenggerimyces |
| Tengunoibacter | https://lpsn.dsmz.de/genus/tengunoibacter |
| Tenuibacillus | https://lpsn.dsmz.de/genus/tenuibacillus |
| Tenuifilum | https://lpsn.dsmz.de/genus/tenuifilum |

| | |
|---|---|
| Tepidamorphus | https://lpsn.dsmz.de/genus/tepidamorphus |
| Tepidanaerobacter | https://lpsn.dsmz.de/genus/tepidanaerobacter |
| Tepidibacillus | https://lpsn.dsmz.de/genus/tepidibacillus |
| Tepidibacter | https://lpsn.dsmz.de/genus/tepidibacter |
| Tepidibaculum | https://lpsn.dsmz.de/genus/tepidibaculum |
| Tepidicaulis | https://lpsn.dsmz.de/genus/tepidicaulis |
| Tepidicella | https://lpsn.dsmz.de/genus/tepidicella |
| Tepidiforma | https://lpsn.dsmz.de/genus/tepidiforma |
| Tepidimicrobium | https://lpsn.dsmz.de/genus/tepidimicrobium |
| Tepidimonas | https://lpsn.dsmz.de/genus/tepidimonas |
| Tepidiphilus | https://lpsn.dsmz.de/genus/tepidiphilus |
| Tepidisphaera | https://lpsn.dsmz.de/genus/tepidisphaera |
| Tepiditoga | https://lpsn.dsmz.de/genus/tepiditoga |
| Terasakiella | https://lpsn.dsmz.de/genus/terasakiella |
| Terasakiispira | https://lpsn.dsmz.de/genus/terasakiispira |
| Teredinibacter | https://lpsn.dsmz.de/genus/teredinibacter |
| Terrabacter | https://lpsn.dsmz.de/genus/terrabacter |
| Terracidiphilus | https://lpsn.dsmz.de/genus/terracidiphilus |
| Terracoccus | https://lpsn.dsmz.de/genus/terracoccus |
| Terribacillus | https://lpsn.dsmz.de/genus/terribacillus |
| Terricaulis | https://lpsn.dsmz.de/genus/terricaulis |
| Terriglobus | https://lpsn.dsmz.de/genus/terriglobus |
| Terrihabitans | https://lpsn.dsmz.de/genus/terrihabitans |
| Terrilactibacillus | https://lpsn.dsmz.de/genus/terrilactibacillus |
| Terrimesophilobacter | https://lpsn.dsmz.de/genus/terrimesophilobacter |
| Terrimicrobium | https://lpsn.dsmz.de/genus/terrimicrobium |
| Terrimonas | https://lpsn.dsmz.de/genus/terrimonas |
| Terrisporobacter | https://lpsn.dsmz.de/genus/terrisporobacter |
| Tersicoccus | https://lpsn.dsmz.de/genus/tersicoccus |
| Tessaracoccus | https://lpsn.dsmz.de/genus/tessaracoccus |
| Testudinibacter | https://lpsn.dsmz.de/genus/testudinibacter |
| Tetragenococcus | https://lpsn.dsmz.de/genus/tetragenococcus |
| Tetrasphaera | https://lpsn.dsmz.de/genus/tetrasphaera |
| Tetrathiobacter | https://lpsn.dsmz.de/genus/tetrathiobacter |
| Texcoconibacillus | https://lpsn.dsmz.de/genus/texcoconibacillus |
| Thalassiella | https://lpsn.dsmz.de/genus/thalassiella |
| Thalassobacillus | https://lpsn.dsmz.de/genus/thalassobacillus |
| Thalassobacter | https://lpsn.dsmz.de/genus/thalassobacter |
| Thalassobaculum | https://lpsn.dsmz.de/genus/thalassobaculum |
| Thalassobius | https://lpsn.dsmz.de/genus/thalassobius |
| Thalassocella | https://lpsn.dsmz.de/genus/thalassocella |
| Thalassococcus | https://lpsn.dsmz.de/genus/thalassococcus |
| Thalassocola | https://lpsn.dsmz.de/genus/thalassocola |
| Thalassoglobus | https://lpsn.dsmz.de/genus/thalassoglobus |

| | |
|---|---|
| Thalassolituus | https://lpsn.dsmz.de/genus/thalassolituus |
| Thalassomonas | https://lpsn.dsmz.de/genus/thalassomonas |
| Thalassorhabdomicrobium | https://lpsn.dsmz.de/genus/thalassorhabdomicrobium |
| Thalassorhabdus | https://lpsn.dsmz.de/genus/thalassorhabdus |
| Thalassospira | https://lpsn.dsmz.de/genus/thalassospira |
| Thalassotalea | https://lpsn.dsmz.de/genus/thalassotalea |
| Thauera | https://lpsn.dsmz.de/genus/thauera |
| Thaumasiovibrio | https://lpsn.dsmz.de/genus/thaumasiovibrio |
| Thermacetogenium | https://lpsn.dsmz.de/genus/thermacetogenium |
| Thermaerobacter | https://lpsn.dsmz.de/genus/thermaerobacter |
| Thermanaeromonas | https://lpsn.dsmz.de/genus/thermanaeromonas |
| Thermanaerosceptrum | https://lpsn.dsmz.de/genus/thermanaerosceptrum |
| Thermanaerovibrio | https://lpsn.dsmz.de/genus/thermanaerovibrio |
| Thermasporomyces | https://lpsn.dsmz.de/genus/thermasporomyces |
| Thermaurantiacus | https://lpsn.dsmz.de/genus/thermaurantiacus |
| Thermaurantimonas | https://lpsn.dsmz.de/genus/thermaurantimonas |
| Thermicanus | https://lpsn.dsmz.de/genus/thermicanus |
| Thermincola | https://lpsn.dsmz.de/genus/thermincola |
| Thermithiobacillus | https://lpsn.dsmz.de/genus/thermithiobacillus |
| Thermoactinomyces | https://lpsn.dsmz.de/genus/thermoactinomyces |
| Thermoactinospora | https://lpsn.dsmz.de/genus/thermoactinospora |
| Thermoanaerobacter | https://lpsn.dsmz.de/genus/thermoanaerobacter |
| Thermoanaerobacterium | https://lpsn.dsmz.de/genus/thermoanaerobacterium |
| Thermoanaerobaculum | https://lpsn.dsmz.de/genus/thermoanaerobaculum |
| Thermoanaerobium | https://lpsn.dsmz.de/genus/thermoanaerobium |
| Thermoanaerosceptrum | https://lpsn.dsmz.de/genus/thermoanaerosceptrum |
| Thermobacillus | https://lpsn.dsmz.de/genus/thermobacillus |
| Thermobacteroides | https://lpsn.dsmz.de/genus/thermobacteroides |
| Thermobifida | https://lpsn.dsmz.de/genus/thermobifida |
| Thermobispora | https://lpsn.dsmz.de/genus/thermobispora |
| Thermobrachium | https://lpsn.dsmz.de/genus/thermobrachium |
| Thermocatellispora | https://lpsn.dsmz.de/genus/thermocatellispora |
| Thermochromatium | https://lpsn.dsmz.de/genus/thermochromatium |
| Thermocladium | https://lpsn.dsmz.de/genus/thermocladium |
| Thermoclostridium | https://lpsn.dsmz.de/genus/thermoclostridium |
| Thermococcoides | https://lpsn.dsmz.de/genus/thermococcoides |
| Thermococcus | https://lpsn.dsmz.de/genus/thermococcus |
| Thermocrinis | https://lpsn.dsmz.de/genus/thermocrinis |
| Thermocrispum | https://lpsn.dsmz.de/genus/thermocrispum |
| Thermodesulfatator | https://lpsn.dsmz.de/genus/thermodesulfatator |
| Thermodesulfitimonas | https://lpsn.dsmz.de/genus/thermodesulfitimonas |
| Thermodesulfobacterium | https://lpsn.dsmz.de/genus/thermodesulfobacterium |
| Thermodesulfobium | https://lpsn.dsmz.de/genus/thermodesulfobium |
| Thermodesulfomicrobium | https://lpsn.dsmz.de/genus/thermodesulfomicrobium |

| | |
|---|---|
| Thermodesulforhabdus | https://lpsn.dsmz.de/genus/thermodesulforhabdus |
| Thermodesulfovibrio | https://lpsn.dsmz.de/genus/thermodesulfovibrio |
| Thermodiscus | https://lpsn.dsmz.de/genus/thermodiscus |
| Thermofilum | https://lpsn.dsmz.de/genus/thermofilum |
| Thermoflavifilum | https://lpsn.dsmz.de/genus/thermoflavifilum |
| Thermoflavimicrobium | https://lpsn.dsmz.de/genus/thermoflavimicrobium |
| Thermoflexibacter | https://lpsn.dsmz.de/genus/thermoflexibacter |
| Thermoflexus | https://lpsn.dsmz.de/genus/thermoflexus |
| Thermogemmata | https://lpsn.dsmz.de/genus/thermogemmata |
| Thermogemmatispora | https://lpsn.dsmz.de/genus/thermogemmatispora |
| Thermogladius | https://lpsn.dsmz.de/genus/thermogladius |
| Thermogutta | https://lpsn.dsmz.de/genus/thermogutta |
| Thermogymnomonas | https://lpsn.dsmz.de/genus/thermogymnomonas |
| Thermohalobacter | https://lpsn.dsmz.de/genus/thermohalobacter |
| Thermohydrogenium | https://lpsn.dsmz.de/genus/thermohydrogenium |
| Thermoleophilum | https://lpsn.dsmz.de/genus/thermoleophilum |
| Thermolithobacter | https://lpsn.dsmz.de/genus/thermolithobacter |
| Thermolongibacillus | https://lpsn.dsmz.de/genus/thermolongibacillus |
| Thermomarinilinea | https://lpsn.dsmz.de/genus/thermomarinilinea |
| Thermomicrobium | https://lpsn.dsmz.de/genus/thermomicrobium |
| Thermomonas | https://lpsn.dsmz.de/genus/thermomonas |
| Thermomonospora | https://lpsn.dsmz.de/genus/thermomonospora |
| Thermonema | https://lpsn.dsmz.de/genus/thermonema |
| Thermophagus | https://lpsn.dsmz.de/genus/thermophagus |
| Thermophilibacter | https://lpsn.dsmz.de/genus/thermophilibacter |
| Thermoplasma | https://lpsn.dsmz.de/genus/thermoplasma |
| Thermopolyspora | https://lpsn.dsmz.de/genus/thermopolyspora |
| Thermoproteus | https://lpsn.dsmz.de/genus/thermoproteus |
| Thermorudis | https://lpsn.dsmz.de/genus/thermorudis |
| Thermosediminibacter | https://lpsn.dsmz.de/genus/thermosediminibacter |
| Thermosinus | https://lpsn.dsmz.de/genus/thermosinus |
| Thermosipho | https://lpsn.dsmz.de/genus/thermosipho |
| Thermosphaera | https://lpsn.dsmz.de/genus/thermosphaera |
| Thermosporothrix | https://lpsn.dsmz.de/genus/thermosporothrix |
| Thermostaphylospora | https://lpsn.dsmz.de/genus/thermostaphylospora |
| Thermostichus | https://lpsn.dsmz.de/genus/thermostichus |
| Thermostilla | https://lpsn.dsmz.de/genus/thermostilla |
| Thermosulfidibacter | https://lpsn.dsmz.de/genus/thermosulfidibacter |
| Thermosulfurimonas | https://lpsn.dsmz.de/genus/thermosulfurimonas |
| Thermosulfuriphilus | https://lpsn.dsmz.de/genus/thermosulfuriphilus |
| Thermosynergistes | https://lpsn.dsmz.de/genus/thermosynergistes |
| Thermosyntropha | https://lpsn.dsmz.de/genus/thermosyntropha |
| Thermotalea | https://lpsn.dsmz.de/genus/thermotalea |
| Thermoterrabacterium | https://lpsn.dsmz.de/genus/thermoterrabacterium |

| | |
|---|---|
| Thermothrix | https://lpsn.dsmz.de/genus/thermothrix |
| Thermotoga | https://lpsn.dsmz.de/genus/thermotoga |
| Thermotomaculum | https://lpsn.dsmz.de/genus/thermotomaculum |
| Thermotunica | https://lpsn.dsmz.de/genus/thermotunica |
| Thermovenabulum | https://lpsn.dsmz.de/genus/thermovenabulum |
| Thermovibrio | https://lpsn.dsmz.de/genus/thermovibrio |
| Thermovirga | https://lpsn.dsmz.de/genus/thermovirga |
| Thermovorax | https://lpsn.dsmz.de/genus/thermovorax |
| Thermovum | https://lpsn.dsmz.de/genus/thermovum |
| Thermus | https://lpsn.dsmz.de/genus/thermus |
| Thialkalicoccus | https://lpsn.dsmz.de/genus/thialkalicoccus |
| Thialkalimicrobium | https://lpsn.dsmz.de/genus/thialkalimicrobium |
| Thialkalivibrio | https://lpsn.dsmz.de/genus/thialkalivibrio |
| Thioalbus | https://lpsn.dsmz.de/genus/thioalbus |
| Thioalkalibacter | https://lpsn.dsmz.de/genus/thioalkalibacter |
| Thioalkalicoccus | https://lpsn.dsmz.de/genus/thioalkalicoccus |
| Thioalkalimicrobium | https://lpsn.dsmz.de/genus/thioalkalimicrobium |
| Thioalkalispira | https://lpsn.dsmz.de/genus/thioalkalispira |
| Thioalkalivibrio | https://lpsn.dsmz.de/genus/thioalkalivibrio |
| Thiobaca | https://lpsn.dsmz.de/genus/thiobaca |
| Thiobacillus | https://lpsn.dsmz.de/genus/thiobacillus |
| Thiobacimonas | https://lpsn.dsmz.de/genus/thiobacimonas |
| Thiobacter | https://lpsn.dsmz.de/genus/thiobacter |
| Thiobacterium | https://lpsn.dsmz.de/genus/thiobacterium |
| Thiocapsa | https://lpsn.dsmz.de/genus/thiocapsa |
| Thioclava | https://lpsn.dsmz.de/genus/thioclava |
| Thiococcus | https://lpsn.dsmz.de/genus/thiococcus |
| Thiocystis | https://lpsn.dsmz.de/genus/thiocystis |
| Thiodictyon | https://lpsn.dsmz.de/genus/thiodictyon |
| Thiofaba | https://lpsn.dsmz.de/genus/thiofaba |
| Thiofilum | https://lpsn.dsmz.de/genus/thiofilum |
| Thioflexithrix | https://lpsn.dsmz.de/genus/thioflexithrix |
| Thiogranum | https://lpsn.dsmz.de/genus/thiogranum |
| Thiohalobacter | https://lpsn.dsmz.de/genus/thiohalobacter |
| Thiohalocapsa | https://lpsn.dsmz.de/genus/thiohalocapsa |
| Thiohalomonas | https://lpsn.dsmz.de/genus/thiohalomonas |
| Thiohalophilus | https://lpsn.dsmz.de/genus/thiohalophilus |
| Thiohalorhabdus | https://lpsn.dsmz.de/genus/thiohalorhabdus |
| Thiohalospira | https://lpsn.dsmz.de/genus/thiohalospira |
| Thiolamprovum | https://lpsn.dsmz.de/genus/thiolamprovum |
| Thiolapillus | https://lpsn.dsmz.de/genus/thiolapillus |
| Thiolinea | https://lpsn.dsmz.de/genus/thiolinea |
| Thiomargarita | https://lpsn.dsmz.de/genus/thiomargarita |
| Thiomicrorhabdus | https://lpsn.dsmz.de/genus/thiomicrorhabdus |

| | |
|---|---|
| Thiomicrospira | https://lpsn.dsmz.de/genus/thiomicrospira |
| Thiomonas | https://lpsn.dsmz.de/genus/thiomonas |
| Thiopedia | https://lpsn.dsmz.de/genus/thiopedia |
| Thiophaeococcus | https://lpsn.dsmz.de/genus/thiophaeococcus |
| Thioploca | https://lpsn.dsmz.de/genus/thioploca |
| Thioprofundum | https://lpsn.dsmz.de/genus/thioprofundum |
| Thiopseudomonas | https://lpsn.dsmz.de/genus/thiopseudomonas |
| Thioreductor | https://lpsn.dsmz.de/genus/thioreductor |
| Thiorhodococcus | https://lpsn.dsmz.de/genus/thiorhodococcus |
| Thiorhodospira | https://lpsn.dsmz.de/genus/thiorhodospira |
| Thiorhodovibrio | https://lpsn.dsmz.de/genus/thiorhodovibrio |
| Thiosphaera | https://lpsn.dsmz.de/genus/thiosphaera |
| Thiospira | https://lpsn.dsmz.de/genus/thiospira |
| Thiospirillum | https://lpsn.dsmz.de/genus/thiospirillum |
| Thiospirochaeta | https://lpsn.dsmz.de/genus/thiospirochaeta |
| Thiosulfatihalobacter | https://lpsn.dsmz.de/genus/thiosulfatihalobacter |
| Thiosulfatimonas | https://lpsn.dsmz.de/genus/thiosulfatimonas |
| Thiosulfativibrio | https://lpsn.dsmz.de/genus/thiosulfativibrio |
| Thiothrix | https://lpsn.dsmz.de/genus/thiothrix |
| Thiovirga | https://lpsn.dsmz.de/genus/thiovirga |
| Thiovulum | https://lpsn.dsmz.de/genus/thiovulum |
| Thorsellia | https://lpsn.dsmz.de/genus/thorsellia |
| Tianweitania | https://lpsn.dsmz.de/genus/tianweitania |
| Tibeticola | https://lpsn.dsmz.de/genus/tibeticola |
| Tichowtungia | https://lpsn.dsmz.de/genus/tichowtungia |
| Timonella | https://lpsn.dsmz.de/genus/timonella |
| Tindallia | https://lpsn.dsmz.de/genus/tindallia |
| Tissierella | https://lpsn.dsmz.de/genus/tissierella |
| Tistlia | https://lpsn.dsmz.de/genus/tistlia |
| Tistrella | https://lpsn.dsmz.de/genus/tistrella |
| Tolumonas | https://lpsn.dsmz.de/genus/tolumonas |
| Tomitella | https://lpsn.dsmz.de/genus/tomitella |
| Tonsilliphilus | https://lpsn.dsmz.de/genus/tonsilliphilus |
| Toxothrix | https://lpsn.dsmz.de/genus/toxothrix |
| Trabulsiella | https://lpsn.dsmz.de/genus/trabulsiella |
| Tractidigestivibacter | https://lpsn.dsmz.de/genus/tractidigestivibacter |
| Tranquillimonas | https://lpsn.dsmz.de/genus/tranquillimonas |
| Trebonia | https://lpsn.dsmz.de/genus/trebonia |
| Treponema | https://lpsn.dsmz.de/genus/treponema |
| Trichlorobacter | https://lpsn.dsmz.de/genus/trichlorobacter |
| Trichococcus | https://lpsn.dsmz.de/genus/trichococcus |
| Trinickia | https://lpsn.dsmz.de/genus/trinickia |
| Tritonibacter | https://lpsn.dsmz.de/genus/tritonibacter |
| Tropicibacter | https://lpsn.dsmz.de/genus/tropicibacter |

| | |
|---|---|
| Tropicihabitans | https://lpsn.dsmz.de/genus/tropicihabitans |
| Tropicimonas | https://lpsn.dsmz.de/genus/tropicimonas |
| Truepera | https://lpsn.dsmz.de/genus/truepera |
| Trueperella | https://lpsn.dsmz.de/genus/trueperella |
| Tsukamurella | https://lpsn.dsmz.de/genus/tsukamurella |
| Tsuneonella | https://lpsn.dsmz.de/genus/tsuneonella |
| Tuberibacillus | https://lpsn.dsmz.de/genus/tuberibacillus |
| Tumebacillus | https://lpsn.dsmz.de/genus/tumebacillus |
| Tundrisphaera | https://lpsn.dsmz.de/genus/tundrisphaera |
| Tunicatimonas | https://lpsn.dsmz.de/genus/tunicatimonas |
| Turicella | https://lpsn.dsmz.de/genus/turicella |
| Turicibacter | https://lpsn.dsmz.de/genus/turicibacter |
| Turicimonas | https://lpsn.dsmz.de/genus/turicimonas |
| Turneriella | https://lpsn.dsmz.de/genus/turneriella |
| Tuwongella | https://lpsn.dsmz.de/genus/tuwongella |
| Uliginosibacterium | https://lpsn.dsmz.de/genus/uliginosibacterium |
| Ulvibacter | https://lpsn.dsmz.de/genus/ulvibacter |
| Ulvibacterium | https://lpsn.dsmz.de/genus/ulvibacterium |
| Umboniibacter | https://lpsn.dsmz.de/genus/umboniibacter |
| Umezawaea | https://lpsn.dsmz.de/genus/umezawaea |
| Undibacter | https://lpsn.dsmz.de/genus/undibacter |
| Undibacterium | https://lpsn.dsmz.de/genus/undibacterium |
| Urbifossiella | https://lpsn.dsmz.de/genus/urbifossiella |
| Ureaplasma | https://lpsn.dsmz.de/genus/ureaplasma |
| Urechidicola | https://lpsn.dsmz.de/genus/urechidicola |
| Ureibacillus | https://lpsn.dsmz.de/genus/ureibacillus |
| Ursidibacter | https://lpsn.dsmz.de/genus/ursidibacter |
| Uruburuella | https://lpsn.dsmz.de/genus/uruburuella |
| Usitatibacter | https://lpsn.dsmz.de/genus/usitatibacter |
| Vadicella | https://lpsn.dsmz.de/genus/vadicella |
| Vaginimicrobium | https://lpsn.dsmz.de/genus/vaginimicrobium |
| Vagococcus | https://lpsn.dsmz.de/genus/vagococcus |
| Vallicoccus | https://lpsn.dsmz.de/genus/vallicoccus |
| Vallitalea | https://lpsn.dsmz.de/genus/vallitalea |
| Vampirovibrio | https://lpsn.dsmz.de/genus/vampirovibrio |
| Vandammella | https://lpsn.dsmz.de/genus/vandammella |
| Vannielia | https://lpsn.dsmz.de/genus/vannielia |
| Varibaculum | https://lpsn.dsmz.de/genus/varibaculum |
| Variibacter | https://lpsn.dsmz.de/genus/variibacter |
| Variimorphobacter | https://lpsn.dsmz.de/genus/variimorphobacter |
| Variovorax | https://lpsn.dsmz.de/genus/variovorax |
| Varunaivibrio | https://lpsn.dsmz.de/genus/varunaivibrio |
| Vasilyevaea | https://lpsn.dsmz.de/genus/vasilyevaea |
| Veillonella | https://lpsn.dsmz.de/genus/veillonella |

| | |
|---|---|
| Velocimicrobium | https://lpsn.dsmz.de/genus/velocimicrobium |
| Venenivibrio | https://lpsn.dsmz.de/genus/venenivibrio |
| Ventosimonas | https://lpsn.dsmz.de/genus/ventosimonas |
| Verminephrobacter | https://lpsn.dsmz.de/genus/verminephrobacter |
| Veronia | https://lpsn.dsmz.de/genus/veronia |
| Verrucomicrobium | https://lpsn.dsmz.de/genus/verrucomicrobium |
| Verrucosispora | https://lpsn.dsmz.de/genus/verrucosispora |
| Verticia | https://lpsn.dsmz.de/genus/verticia |
| Verticiella | https://lpsn.dsmz.de/genus/verticiella |
| Vescimonas | https://lpsn.dsmz.de/genus/vescimonas |
| Vespertiliibacter | https://lpsn.dsmz.de/genus/vespertiliibacter |
| Vibrio | https://lpsn.dsmz.de/genus/vibrio |
| Vibrionimonas | https://lpsn.dsmz.de/genus/vibrionimonas |
| Vicinamibacter | https://lpsn.dsmz.de/genus/vicinamibacter |
| Vicingus | https://lpsn.dsmz.de/genus/vicingus |
| Victivallis | https://lpsn.dsmz.de/genus/victivallis |
| Vineibacter | https://lpsn.dsmz.de/genus/vineibacter |
| Virgibacillus | https://lpsn.dsmz.de/genus/virgibacillus |
| Virgisporangium | https://lpsn.dsmz.de/genus/virgisporangium |
| Virgosporangium | https://lpsn.dsmz.de/genus/virgosporangium |
| Viridibacillus | https://lpsn.dsmz.de/genus/viridibacillus |
| Vitellibacter | https://lpsn.dsmz.de/genus/vitellibacter |
| Vitiosangium | https://lpsn.dsmz.de/genus/vitiosangium |
| Vitreimonas | https://lpsn.dsmz.de/genus/vitreimonas |
| Vitreoscilla | https://lpsn.dsmz.de/genus/vitreoscilla |
| Vogesella | https://lpsn.dsmz.de/genus/vogesella |
| Volcaniella | https://lpsn.dsmz.de/genus/volcaniella |
| Volucribacter | https://lpsn.dsmz.de/genus/volucribacter |
| Vulcanibacillus | https://lpsn.dsmz.de/genus/vulcanibacillus |
| Vulcaniibacterium | https://lpsn.dsmz.de/genus/vulcaniibacterium |
| Vulcanisaeta | https://lpsn.dsmz.de/genus/vulcanisaeta |
| Vulcanithermus | https://lpsn.dsmz.de/genus/vulcanithermus |
| Vulgatibacter | https://lpsn.dsmz.de/genus/vulgatibacter |
| Waddlia | https://lpsn.dsmz.de/genus/waddlia |
| Wagnerdoeblera | https://lpsn.dsmz.de/genus/wagnerdoeblera |
| Waltera | https://lpsn.dsmz.de/genus/waltera |
| Wandonia | https://lpsn.dsmz.de/genus/wandonia |
| Wangia | https://lpsn.dsmz.de/genus/wangia |
| Wautersia | https://lpsn.dsmz.de/genus/wautersia |
| Wautersiella | https://lpsn.dsmz.de/genus/wautersiella |
| Weeksella | https://lpsn.dsmz.de/genus/weeksella |
| Weissella | https://lpsn.dsmz.de/genus/weissella |
| Weizmannia | https://lpsn.dsmz.de/genus/weizmannia |
| Wenjunlia | https://lpsn.dsmz.de/genus/wenjunlia |

| | |
|---|---|
| Wenxinia | https://lpsn.dsmz.de/genus/wenxinia |
| Wenyingzhuangia | https://lpsn.dsmz.de/genus/wenyingzhuangia |
| Wenzhouxiangella | https://lpsn.dsmz.de/genus/wenzhouxiangella |
| Wielerella | https://lpsn.dsmz.de/genus/wielerella |
| Wigglesworthia | https://lpsn.dsmz.de/genus/wigglesworthia |
| Williamsia | https://lpsn.dsmz.de/genus/williamsia |
| Williamsoniiplasma | https://lpsn.dsmz.de/genus/williamsoniiplasma |
| Williamwhitmania | https://lpsn.dsmz.de/genus/williamwhitmania |
| Winkia | https://lpsn.dsmz.de/genus/winkia |
| Winogradskyella | https://lpsn.dsmz.de/genus/winogradskyella |
| Wocania | https://lpsn.dsmz.de/genus/wocania |
| Woeseia | https://lpsn.dsmz.de/genus/woeseia |
| Wohlfahrtiimonas | https://lpsn.dsmz.de/genus/wohlfahrtiimonas |
| Wolbachia | https://lpsn.dsmz.de/genus/wolbachia |
| Wolinella | https://lpsn.dsmz.de/genus/wolinella |
| Woodsholea | https://lpsn.dsmz.de/genus/woodsholea |
| Wukongibacter | https://lpsn.dsmz.de/genus/wukongibacter |
| Xanthobacter | https://lpsn.dsmz.de/genus/xanthobacter |
| Xanthomarina | https://lpsn.dsmz.de/genus/xanthomarina |
| Xanthomonas | https://lpsn.dsmz.de/genus/xanthomonas |
| Xanthovirga | https://lpsn.dsmz.de/genus/xanthovirga |
| Xaviernesmea | https://lpsn.dsmz.de/genus/xaviernesmea |
| Xenophilus | https://lpsn.dsmz.de/genus/xenophilus |
| Xenorhabdus | https://lpsn.dsmz.de/genus/xenorhabdus |
| Xiamenia | https://lpsn.dsmz.de/genus/xiamenia |
| Xiangella | https://lpsn.dsmz.de/genus/xiangella |
| Xinfangfangia | https://lpsn.dsmz.de/genus/xinfangfangia |
| Xuhuaishuia | https://lpsn.dsmz.de/genus/xuhuaishuia |
| Xylanibacillus | https://lpsn.dsmz.de/genus/xylanibacillus |
| Xylanibacter | https://lpsn.dsmz.de/genus/xylanibacter |
| Xylanibacterium | https://lpsn.dsmz.de/genus/xylanibacterium |
| Xylanimicrobium | https://lpsn.dsmz.de/genus/xylanimicrobium |
| Xylanimonas | https://lpsn.dsmz.de/genus/xylanimonas |
| Xylanivirga | https://lpsn.dsmz.de/genus/xylanivirga |
| Xylella | https://lpsn.dsmz.de/genus/xylella |
| Xylophilus | https://lpsn.dsmz.de/genus/xylophilus |
| Yangia | https://lpsn.dsmz.de/genus/yangia |
| Yania | https://lpsn.dsmz.de/genus/yania |
| Yaniella | https://lpsn.dsmz.de/genus/yaniella |
| Yeosuana | https://lpsn.dsmz.de/genus/yeosuana |
| Yersinia | https://lpsn.dsmz.de/genus/yersinia |
| Yimella | https://lpsn.dsmz.de/genus/yimella |
| Yinghuangia | https://lpsn.dsmz.de/genus/yinghuangia |
| Yokenella | https://lpsn.dsmz.de/genus/yokenella |

| | |
|---|---|
| Yonghaparkia | https://lpsn.dsmz.de/genus/yonghaparkia |
| Yoonia | https://lpsn.dsmz.de/genus/yoonia |
| Youhaiella | https://lpsn.dsmz.de/genus/youhaiella |
| Youngiibacter | https://lpsn.dsmz.de/genus/youngiibacter |
| Youngimonas | https://lpsn.dsmz.de/genus/youngimonas |
| Youxingia | https://lpsn.dsmz.de/genus/youxingia |
| Yuhushiella | https://lpsn.dsmz.de/genus/yuhushiella |
| Zafaria | https://lpsn.dsmz.de/genus/zafaria |
| Zarconia | https://lpsn.dsmz.de/genus/zarconia |
| Zavarzinella | https://lpsn.dsmz.de/genus/zavarzinella |
| Zavarzinia | https://lpsn.dsmz.de/genus/zavarzinia |
| Zeaxanthinibacter | https://lpsn.dsmz.de/genus/zeaxanthinibacter |
| Zeimonas | https://lpsn.dsmz.de/genus/zeimonas |
| Zestosphaera | https://lpsn.dsmz.de/genus/zestosphaera |
| Zhangella | https://lpsn.dsmz.de/genus/zhangella |
| Zhaonella | https://lpsn.dsmz.de/genus/zhaonella |
| Zhengella | https://lpsn.dsmz.de/genus/zhengella |
| Zhenhengia | https://lpsn.dsmz.de/genus/zhenhengia |
| Zhihengliuella | https://lpsn.dsmz.de/genus/zhihengliuella |
| Zhizhongheella | https://lpsn.dsmz.de/genus/zhizhongheella |
| Zhongshania | https://lpsn.dsmz.de/genus/zhongshania |
| Zhouia | https://lpsn.dsmz.de/genus/zhouia |
| Zimmermannella | https://lpsn.dsmz.de/genus/zimmermannella |
| Zobellella | https://lpsn.dsmz.de/genus/zobellella |
| Zobellia | https://lpsn.dsmz.de/genus/zobellia |
| Zongyangia | https://lpsn.dsmz.de/genus/zongyangia |
| Zongyanglinia | https://lpsn.dsmz.de/genus/zongyanglinia |
| Zoogloea | https://lpsn.dsmz.de/genus/zoogloea |
| Zooshikella | https://lpsn.dsmz.de/genus/zooshikella |
| Zophobihabitans | https://lpsn.dsmz.de/genus/zophobihabitans |
| Zuelzera | https://lpsn.dsmz.de/genus/zuelzera |
| Zunongwangia | https://lpsn.dsmz.de/genus/zunongwangia |
| Zwartia | https://lpsn.dsmz.de/genus/zwartia |
| Zymobacter | https://lpsn.dsmz.de/genus/zymobacter |
| Zymomonas | https://lpsn.dsmz.de/genus/zymomonas |
| Zymophilus | https://lpsn.dsmz.de/genus/zymophilus |
| [Acidimicrobiales; not assigned to family] | https://lpsn.dsmz.de/family/acidimicrobiales-no-family |
| [Actinomycetales; not assigned to family] | https://lpsn.dsmz.de/family/actinomycetales-no-family |
| [Alteromonadales; not assigned to family] | https://lpsn.dsmz.de/family/alteromonadales-no-family |
| [Anaerolineae; not assigned to family] | https://lpsn.dsmz.de/family/anaerolineae-no-family |
| [Aquificales; not assigned to family] | https://lpsn.dsmz.de/family/aquificales-no-family |
| [Bacillota; not assigned to family] | https://lpsn.dsmz.de/family/bacillota-no-family |
| [Betaproteobacteria; not assigned to family] | https://lpsn.dsmz.de/family/betaproteobacteria-no-family |
| [Burkholderiales; not assigned to family] | https://lpsn.dsmz.de/family/burkholderiales-no-family |

| | |
|---|---|
| [Calditrichota; not assigned to family] | https://lpsn.dsmz.de/family/calditrichota-no-family |
| [Caryophanales; not assigned to family] | https://lpsn.dsmz.de/family/caryophanales-no-family |
| [Cytophagales; not assigned to family] | https://lpsn.dsmz.de/family/cytophagales-no-family |
| [Deltaproteobacteria; not assigned to family] | https://lpsn.dsmz.de/family/deltaproteobacteria-no-family |
| [Eubacteriales; not assigned to family] | https://lpsn.dsmz.de/family/eubacteriales-no-family |
| [Gammaproteobacteria; not assigned to family] | https://lpsn.dsmz.de/family/gammaproteobacteria-no-family |
| [Gemmatales; not assigned to family] | https://lpsn.dsmz.de/family/gemmatales-no-family |
| [Halanaerobiales; not assigned to family] | https://lpsn.dsmz.de/family/halanaerobiales-no-family |
| [Hydrogenophilia; not assigned to family] | https://lpsn.dsmz.de/family/hydrogenophilia-no-family |
| [Hyphomicrobiales; not assigned to family] | https://lpsn.dsmz.de/family/hyphomicrobiales-no-family |
| [Lysobacterales; not assigned to family] | https://lpsn.dsmz.de/family/lysobacterales-no-family |
| [Methylococcales; not assigned to family] | https://lpsn.dsmz.de/family/methylococcales-no-family |
| [Micrococcales; not assigned to family] | https://lpsn.dsmz.de/family/micrococcales-no-family |
| [Oceanospirillales; not assigned to family] | https://lpsn.dsmz.de/family/oceanospirillales-no-family |
| [Rickettsiales; not assigned to family] | https://lpsn.dsmz.de/family/rickettsiales-no-family |
| [Spirochaetia; not assigned to family] | https://lpsn.dsmz.de/family/spirochaetia-no-family |
| [Streptosporangiales; not assigned to family] | https://lpsn.dsmz.de/family/streptosporangiales-no-family |
| [Synergistota; not assigned to family] | https://lpsn.dsmz.de/family/synergistota-no-family |
| [Thermoanaerobacterales; not assigned to family] | https://lpsn.dsmz.de/family/thermoanaerobacterales-no-family |
| [Thermodesulfobacteriota; not assigned to family] | https://lpsn.dsmz.de/family/thermodesulfobacteriota-no-family |
| [Thermoplasmatales; not assigned to family] | https://lpsn.dsmz.de/family/thermoplasmatales-no-family |
| [Verrucomicrobiota; not assigned to family] | https://lpsn.dsmz.de/family/verrucomicrobiota-no-family |
| Abditibacteriaceae | https://lpsn.dsmz.de/family/abditibacteriaceae |
| Acanthopleuribacteraceae | https://lpsn.dsmz.de/family/acanthopleuribacteraceae |
| Acetobacteraceae | https://lpsn.dsmz.de/family/acetobacteraceae |
| Acholeplasmataceae | https://lpsn.dsmz.de/family/acholeplasmataceae |
| Achromatiaceae | https://lpsn.dsmz.de/family/thiotrichaceae |
| Achromobacteraceae | https://lpsn.dsmz.de/family/alcaligenaceae |
| Acidaminococcaceae | https://lpsn.dsmz.de/family/acidaminococcaceae |
| Acidiferrobacteraceae | https://lpsn.dsmz.de/family/acidiferrobacteraceae |
| Acidilobaceae | https://lpsn.dsmz.de/family/acidilobaceae |
| Acidimicrobiaceae | https://lpsn.dsmz.de/family/acidimicrobiaceae |
| Acidithiobacillaceae | https://lpsn.dsmz.de/family/acidithiobacillaceae |
| Acidobacteriaceae | https://lpsn.dsmz.de/family/acidobacteriaceae |
| Acidothermaceae | https://lpsn.dsmz.de/family/acidothermaceae |
| Actinocatenisporaceae | https://lpsn.dsmz.de/family/micromonosporaceae |
| Actinomycetaceae | https://lpsn.dsmz.de/family/actinomycetaceae |
| Actinoplanaceae | https://lpsn.dsmz.de/family/micromonosporaceae |
| Actinoplanetaceae | https://lpsn.dsmz.de/family/micromonosporaceae |
| Actinopolymorphaceae | https://lpsn.dsmz.de/family/actinopolymorphaceae |
| Actinopolysporaceae | https://lpsn.dsmz.de/family/pseudonocardiaceae |
| Actinospicaceae | https://lpsn.dsmz.de/family/actinospicaceae |
| Actinosynnemataceae | https://lpsn.dsmz.de/family/pseudonocardiaceae |
| Actinotaleaceae | https://lpsn.dsmz.de/family/cellulomonadaceae |

| | |
|---|---|
| Acuticoccaceae | https://lpsn.dsmz.de/family/amorphaceae |
| Aerococcaceae | https://lpsn.dsmz.de/family/aerococcaceae |
| Aeromonadaceae | https://lpsn.dsmz.de/family/aeromonadaceae |
| Aestuariirhabdaceae | https://lpsn.dsmz.de/family/aestuariirhabdaceae |
| Aestuariivirgaceae | https://lpsn.dsmz.de/family/aestuariivirgaceae |
| Afifellaceae | https://lpsn.dsmz.de/family/afifellaceae |
| Aggregatilineaceae | https://lpsn.dsmz.de/family/aggregatilineaceae |
| Ahrensiaceae | https://lpsn.dsmz.de/family/ahrensiaceae |
| Akkermansiaceae | https://lpsn.dsmz.de/family/akkermansiaceae |
| Alcaligenaceae | https://lpsn.dsmz.de/family/alcaligenaceae |
| Alcanivoracaceae | https://lpsn.dsmz.de/family/alcanivoracaceae |
| Alcanivoraceae | https://lpsn.dsmz.de/family/alcanivoracaceae |
| Algiphilaceae | https://lpsn.dsmz.de/family/algiphilaceae |
| Alicyclobacillaceae | https://lpsn.dsmz.de/family/alicyclobacillaceae |
| Allostreptomycetaceae | https://lpsn.dsmz.de/family/streptomycetaceae |
| Alsobacteraceae | https://lpsn.dsmz.de/family/alsobacteraceae |
| Alteromonadaceae | https://lpsn.dsmz.de/family/alteromonadaceae |
| Amorphaceae | https://lpsn.dsmz.de/family/amorphaceae |
| Anaerohalosphaeraceae | https://lpsn.dsmz.de/family/anaerohalosphaeraceae |
| Anaerolineaceae | https://lpsn.dsmz.de/family/anaerolineaceae |
| Anaeromyxobacteraceae | https://lpsn.dsmz.de/family/anaeromyxobacteraceae |
| Anaeroplasmataceae | https://lpsn.dsmz.de/family/anaeroplasmataceae |
| Anaplasmataceae | https://lpsn.dsmz.de/family/ehrlichiaceae |
| Ancalomicrobiaceae | https://lpsn.dsmz.de/family/ancalomicrobiaceae |
| Antricoccaceae | https://lpsn.dsmz.de/family/antricoccaceae |
| Anulomycetaceae | https://lpsn.dsmz.de/family/metamycoplasmataceae |
| Aphanothecaceae | https://lpsn.dsmz.de/family/aphanothecaceae |
| Aquaspirillaceae | https://lpsn.dsmz.de/family/chromobacteriaceae |
| Aquificaceae | https://lpsn.dsmz.de/family/aquificaceae |
| Aquipuribacteraceae | https://lpsn.dsmz.de/family/intrasporangiaceae |
| Arcanobacteriaceae | https://lpsn.dsmz.de/family/actinomycetaceae |
| Archaeoglobaceae | https://lpsn.dsmz.de/family/archaeoglobaceae |
| Archangiaceae | https://lpsn.dsmz.de/family/archangiaceae |
| Arcobacteraceae | https://lpsn.dsmz.de/family/arcobacteraceae |
| Ardenticatenaceae | https://lpsn.dsmz.de/family/ardenticatenaceae |
| Arenicellaceae | https://lpsn.dsmz.de/family/arenicellaceae |
| Arenimicrobiaceae | https://lpsn.dsmz.de/family/arenimicrobiaceae |
| Armatimonadaceae | https://lpsn.dsmz.de/family/armatimonadaceae |
| Arsenicicoccaceae | https://lpsn.dsmz.de/family/dermatophilaceae |
| Atopobiaceae | https://lpsn.dsmz.de/family/atopobiaceae |
| Atribacteraceae | https://lpsn.dsmz.de/family/atribacteraceae |
| Aurantimonadaceae | https://lpsn.dsmz.de/family/aurantimonadaceae |
| Azonexaceae | https://lpsn.dsmz.de/family/azonexaceae |
| Azospirillaceae | https://lpsn.dsmz.de/family/azospirillaceae |

| | |
|---|---|
| Azotobacteraceae | https://lpsn.dsmz.de/family/pseudomonadaceae |
| Bacillaceae | https://lpsn.dsmz.de/family/bacillaceae |
| Bacteriovoracaceae | https://lpsn.dsmz.de/family/bacteriovoracaceae |
| Bacteroidaceae | https://lpsn.dsmz.de/family/bacteroidaceae |
| Baekduiaceae | https://lpsn.dsmz.de/family/baekduiaceae |
| Balneatricaceae | https://lpsn.dsmz.de/family/balneatricaceae |
| Balneatrichaceae | https://lpsn.dsmz.de/family/balneatricaceae |
| Balneicellaceae | https://lpsn.dsmz.de/family/balneicellaceae |
| Balneolaceae | https://lpsn.dsmz.de/family/balneolaceae |
| Barnesiellaceae | https://lpsn.dsmz.de/family/barnesiellaceae |
| Bartonellaceae | https://lpsn.dsmz.de/family/bartonellaceae |
| Bdellovibrionaceae | https://lpsn.dsmz.de/family/pseudobdellovibrionaceae |
| Beggiatoaceae | https://lpsn.dsmz.de/family/beggiatoaceae |
| Beijerinckiaceae | https://lpsn.dsmz.de/family/beijerinckiaceae |
| Bernardetiaceae | https://lpsn.dsmz.de/family/bernardetiaceae |
| Beutenbergiaceae | https://lpsn.dsmz.de/family/beutenbergiaceae |
| Bifidobacteriaceae | https://lpsn.dsmz.de/family/bifidobacteriaceae |
| Blastocatellaceae | https://lpsn.dsmz.de/family/blastocatellaceae |
| Blastochloridaceae | https://lpsn.dsmz.de/family/blastochloridaceae |
| Blattabacteriaceae | https://lpsn.dsmz.de/family/blattabacteriaceae |
| Bogoriellaceae | https://lpsn.dsmz.de/family/bogoriellaceae |
| Borreliaceae | https://lpsn.dsmz.de/family/borreliaceae |
| Borrelomycetaceae | https://lpsn.dsmz.de/family/metamycoplasmataceae |
| Boseaceae | https://lpsn.dsmz.de/family/boseaceae |
| Brachyspiraceae | https://lpsn.dsmz.de/family/brachyspiraceae |
| Bradymonadaceae | https://lpsn.dsmz.de/family/bradymonadaceae |
| Bradyrhizobiaceae | https://lpsn.dsmz.de/family/nitrobacteraceae |
| Branhamaceae | https://lpsn.dsmz.de/family/moraxellaceae |
| Breoghaniaceae | https://lpsn.dsmz.de/family/breoghaniaceae |
| Brevibacteriaceae | https://lpsn.dsmz.de/family/brevibacteriaceae |
| Brevinemataceae | https://lpsn.dsmz.de/family/brevinemataceae |
| Breznakiellaceae | https://lpsn.dsmz.de/family/breznakiellaceae |
| Brucellaceae | https://lpsn.dsmz.de/family/brucellaceae |
| Bruguierivoracaceae | https://lpsn.dsmz.de/family/pectobacteriaceae |
| Bryobacteraceae | https://lpsn.dsmz.de/family/bryobacteraceae |
| Budviciaceae | https://lpsn.dsmz.de/family/budviciaceae |
| Burkholderiaceae | https://lpsn.dsmz.de/family/burkholderiaceae |
| Caedimonadaceae | https://lpsn.dsmz.de/family/caedimonadaceae |
| Calderihabitantaceae | https://lpsn.dsmz.de/family/thermoanaerobacteraceae |
| Caldicoprobacteraceae | https://lpsn.dsmz.de/family/caldicoprobacteraceae |
| Caldilineaceae | https://lpsn.dsmz.de/family/caldilineaceae |
| Caldisericaceae | https://lpsn.dsmz.de/family/caldisericaceae |
| Caldisphaeraceae | https://lpsn.dsmz.de/family/caldisphaeraceae |
| Calditerrivibrionaceae | https://lpsn.dsmz.de/family/deferribacteraceae |

| | |
|---|---|
| Calditrichaceae | https://lpsn.dsmz.de/family/calditrichaceae |
| Campylobacteraceae | https://lpsn.dsmz.de/family/campylobacteraceae |
| Capillimicrobiaceae | https://lpsn.dsmz.de/family/capillimicrobiaceae |
| Capsulimonadaceae | https://lpsn.dsmz.de/family/capsulimonadaceae |
| Carbonactinosporaceae | https://lpsn.dsmz.de/family/streptomycetaceae |
| Cardiobacteriaceae | https://lpsn.dsmz.de/family/cardiobacteriaceae |
| Carnobacteriaceae | https://lpsn.dsmz.de/family/carnobacteriaceae |
| Caryophanaceae | https://lpsn.dsmz.de/family/caryophanaceae |
| Casimicrobiaceae | https://lpsn.dsmz.de/family/casimicrobiaceae |
| Catabacteraceae | https://lpsn.dsmz.de/family/christensenellaceae |
| Catabacteriaceae | https://lpsn.dsmz.de/family/christensenellaceae |
| Catalimonadaceae | https://lpsn.dsmz.de/family/catalinimonadaceae |
| Catalinimonadaceae | https://lpsn.dsmz.de/family/catalinimonadaceae |
| Catenulisporaceae | https://lpsn.dsmz.de/family/catenulisporaceae |
| Caulobacteraceae | https://lpsn.dsmz.de/family/caulobacteraceae |
| Celerinatantimonadaceae | https://lpsn.dsmz.de/family/celerinatantimonadaceae |
| Cellulomonadaceae | https://lpsn.dsmz.de/family/cellulomonadaceae |
| Cellvibrionaceae | https://lpsn.dsmz.de/family/cellvibrionaceae |
| Cesiribacteraceae | https://lpsn.dsmz.de/family/cesiribacteraceae |
| Chelatococcaceae | https://lpsn.dsmz.de/family/chelatococcaceae |
| Chitinibacteraceae | https://lpsn.dsmz.de/family/chromobacteriaceae |
| Chitinispirillaceae | https://lpsn.dsmz.de/family/chitinispirillaceae |
| Chitinivibrionaceae | https://lpsn.dsmz.de/family/chitinivibrionaceae |
| Chitinophagaceae | https://lpsn.dsmz.de/family/chitinophagaceae |
| Chlamydiaceae | https://lpsn.dsmz.de/family/chlamydiaceae |
| Chlamydozoaceae | https://lpsn.dsmz.de/family/chlamydiaceae |
| Chloracidobacteriaceae | https://lpsn.dsmz.de/family/acidobacteriaceae |
| Chlorobiaceae | https://lpsn.dsmz.de/family/chlorobiaceae |
| Chloroflexaceae | https://lpsn.dsmz.de/family/chloroflexaceae |
| Christensenellaceae | https://lpsn.dsmz.de/family/christensenellaceae |
| Chromatiaceae | https://lpsn.dsmz.de/family/chromatiaceae |
| Chromobacteriaceae | https://lpsn.dsmz.de/family/chromobacteriaceae |
| Chroococcaceae | https://lpsn.dsmz.de/family/chroococcaceae |
| Chrysiogenaceae | https://lpsn.dsmz.de/family/chrysiogenaceae |
| Chthonomonadaceae | https://lpsn.dsmz.de/family/chthonomonadaceae |
| Clostridiaceae | https://lpsn.dsmz.de/family/clostridiaceae |
| Cohaesibacteraceae | https://lpsn.dsmz.de/family/cohaesibacteraceae |
| Colwelliaceae | https://lpsn.dsmz.de/family/colwelliaceae |
| Comamonadaceae | https://lpsn.dsmz.de/family/comamonadaceae |
| Conexibacteraceae | https://lpsn.dsmz.de/family/conexibacteraceae |
| Conexivisphaeraceae | https://lpsn.dsmz.de/family/thermoproteaceae |
| Coprobacillaceae | https://lpsn.dsmz.de/family/coprobacillaceae |
| Coprothermobacteraceae | https://lpsn.dsmz.de/family/coprothermobacteraceae |
| Coriobacteriaceae | https://lpsn.dsmz.de/family/coriobacteriaceae |

| | |
|---|---|
| Corynebacteriaceae | https://lpsn.dsmz.de/family/corynebacteriaceae |
| Coxiellaceae | https://lpsn.dsmz.de/family/coxiellaceae |
| Crenotrichaceae | https://lpsn.dsmz.de/family/crenotrichaceae |
| Crocinitomicaceae | https://lpsn.dsmz.de/family/crocinitomicaceae |
| Cryomorphaceae | https://lpsn.dsmz.de/family/cryomorphaceae |
| Cryptosporangiaceae | https://lpsn.dsmz.de/family/cryptosporangiaceae |
| Culicoidibacteraceae | https://lpsn.dsmz.de/family/culicoidibacteraceae |
| Cuniculiplasmataceae | https://lpsn.dsmz.de/family/cuniculiplasmataceae |
| Cyanobacteriaceae | https://lpsn.dsmz.de/family/geminocystaceae |
| Cyanobiaceae | https://lpsn.dsmz.de/family/synechococcaceae |
| Cyanothecaceae | https://lpsn.dsmz.de/family/cyanothecaceae |
| Cyclobacteriaceae | https://lpsn.dsmz.de/family/cyclobacteriaceae |
| Cystobacteraceae | https://lpsn.dsmz.de/family/archangiaceae |
| Cytophagaceae | https://lpsn.dsmz.de/family/cytophagaceae |
| Deferribacteraceae | https://lpsn.dsmz.de/family/deferribacteraceae |
| Deferrisomataceae | https://lpsn.dsmz.de/family/deferrisomataceae |
| Defluviitaleaceae | https://lpsn.dsmz.de/family/defluviitaleaceae |
| Dehalococcoidaceae | https://lpsn.dsmz.de/family/dehalococcoidaceae |
| Deinococcaceae | https://lpsn.dsmz.de/family/deinococcaceae |
| Demequinaceae | https://lpsn.dsmz.de/family/demequinaceae |
| Dermabacteraceae | https://lpsn.dsmz.de/family/dermabacteraceae |
| Dermacoccaceae | https://lpsn.dsmz.de/family/dermacoccaceae |
| Dermatophilaceae | https://lpsn.dsmz.de/family/dermatophilaceae |
| Desulfallaceae | https://lpsn.dsmz.de/family/peptococcaceae |
| Desulfallantaceae | https://lpsn.dsmz.de/family/peptococcaceae |
| Desulfarculaceae | https://lpsn.dsmz.de/family/desulfarculaceae |
| Desulfatibacillaceae | https://lpsn.dsmz.de/family/desulfobacteraceae |
| Desulfatiglandaceae | https://lpsn.dsmz.de/family/desulfobacteraceae |
| Desulfatirhabdiaceae | https://lpsn.dsmz.de/family/desulfobacteraceae |
| Desulfitibacteraceae | https://lpsn.dsmz.de/family/peptococcaceae |
| Desulfitobacteriaceae | https://lpsn.dsmz.de/family/desulfitobacteriaceae |
| Desulfobaccaceae | https://lpsn.dsmz.de/family/syntrophaceae |
| Desulfobacteraceae | https://lpsn.dsmz.de/family/desulfobacteraceae |
| Desulfobacteriaceae | https://lpsn.dsmz.de/family/desulfobacteraceae |
| Desulfobulbaceae | https://lpsn.dsmz.de/family/desulfobulbaceae |
| Desulfocapsaceae | https://lpsn.dsmz.de/family/desulfobulbaceae |
| Desulfococcaceae | https://lpsn.dsmz.de/family/desulfobacteraceae |
| Desulfocucumaceae | https://lpsn.dsmz.de/family/peptococcaceae |
| Desulfocucumeraceae | https://lpsn.dsmz.de/family/peptococcaceae |
| Desulfofabaceae | https://lpsn.dsmz.de/family/desulfobacteraceae |
| Desulfofarciminaceae | https://lpsn.dsmz.de/family/peptococcaceae |
| Desulfohalobiaceae | https://lpsn.dsmz.de/family/desulfohalobiaceae |
| Desulfohalotomaculaceae | https://lpsn.dsmz.de/family/peptococcaceae |
| Desulfolunaceae | https://lpsn.dsmz.de/family/desulfobacteraceae |

| | |
|---|---|
| Desulfomicrobiaceae | https://lpsn.dsmz.de/family/desulfomicrobiaceae |
| Desulfomonilaceae | https://lpsn.dsmz.de/family/syntrophaceae |
| Desulfonatronaceae | https://lpsn.dsmz.de/family/desulfonatronaceae |
| Desulfonatronovibrionaceae | https://lpsn.dsmz.de/family/desulfohalobiaceae |
| Desulfonatronumaceae | https://lpsn.dsmz.de/family/desulfonatronaceae |
| Desulfonauticaceae | https://lpsn.dsmz.de/family/desulfohalobiaceae |
| Desulfoplanaceae | https://lpsn.dsmz.de/family/desulfomicrobiaceae |
| Desulfoplanetaceae | https://lpsn.dsmz.de/family/desulfomicrobiaceae |
| Desulforegulaceae | https://lpsn.dsmz.de/family/desulfobacteraceae |
| Desulfosalsimonadaceae | https://lpsn.dsmz.de/family/desulfobacteraceae |
| Desulfosarcinaceae | https://lpsn.dsmz.de/family/desulfobacteraceae |
| Desulfosudaceae | https://lpsn.dsmz.de/family/desulfosudaceae |
| Desulfothermaceae | https://lpsn.dsmz.de/family/desulfohalobiaceae |
| Desulfotomaculaceae | https://lpsn.dsmz.de/family/peptococcaceae |
| Desulfovibrionaceae | https://lpsn.dsmz.de/family/desulfovibrionaceae |
| Desulfurellaceae | https://lpsn.dsmz.de/family/desulfurellaceae |
| Desulfuribacillaceae | https://lpsn.dsmz.de/family/desulfuribacillaceae |
| Desulfurisporaceae | https://lpsn.dsmz.de/family/peptococcaceae |
| Desulfurivibrionaceae | https://lpsn.dsmz.de/family/desulfobulbaceae |
| Desulfurobacteriaceae | https://lpsn.dsmz.de/family/desulfurobacteriaceae |
| Desulfurococcaceae | https://lpsn.dsmz.de/family/desulfurococcaceae |
| Desulfuromonaceae | https://lpsn.dsmz.de/family/desulfuromonadaceae |
| Desulfuromonadaceae | https://lpsn.dsmz.de/family/desulfuromonadaceae |
| Devosiaceae | https://lpsn.dsmz.de/family/devosiaceae |
| Dictyobacteraceae | https://lpsn.dsmz.de/family/dictyobacteraceae |
| Dictyoglomaceae | https://lpsn.dsmz.de/family/dictyoglomeraceae |
| Dictyoglomeraceae | https://lpsn.dsmz.de/family/dictyoglomeraceae |
| Dietziaceae | https://lpsn.dsmz.de/family/dietziaceae |
| Dissulfuribacteraceae | https://lpsn.dsmz.de/family/dissulfuribacteraceae |
| Dissulfurirhabdaceae | https://lpsn.dsmz.de/family/dissulfurirhabdaceae |
| Dissulfurispiraceae | https://lpsn.dsmz.de/family/dissulfurispiraceae |
| Draconibacteriaceae | https://lpsn.dsmz.de/family/prolixibacteraceae |
| Dysgonomonadaceae | https://lpsn.dsmz.de/family/dysgonomonadaceae |
| Ectothiorhodospiraceae | https://lpsn.dsmz.de/family/ectothiorhodospiraceae |
| Eggerthellaceae | https://lpsn.dsmz.de/family/eggerthellaceae |
| Egibacteraceae | https://lpsn.dsmz.de/family/egibacteraceae |
| Egicoccaceae | https://lpsn.dsmz.de/family/egicoccaceae |
| Ehrlichiaceae | https://lpsn.dsmz.de/family/ehrlichiaceae |
| Elioraeaceae | https://lpsn.dsmz.de/family/acetobacteraceae |
| Elusimicrobiaceae | https://lpsn.dsmz.de/family/elusimicrobiaceae |
| Emcibacteraceae | https://lpsn.dsmz.de/family/emcibacteraceae |
| Endomicrobiaceae | https://lpsn.dsmz.de/family/endomicrobiaceae |
| Endozoicomonadaceae | https://lpsn.dsmz.de/family/endozoicomonadaceae |
| Enterobacteraceae | https://lpsn.dsmz.de/family/enterobacteriaceae |

| | |
|---|---|
| Enterobacteriaceae | https://lpsn.dsmz.de/family/enterobacteriaceae |
| Enterococcaceae | https://lpsn.dsmz.de/family/enterococcaceae |
| Entomoplasmataceae | https://lpsn.dsmz.de/family/entomoplasmataceae |
| Erwiniaceae | https://lpsn.dsmz.de/family/erwiniaceae |
| Erysipelatoclostridiaceae | https://lpsn.dsmz.de/family/erysipelotrichaceae |
| Erysipelotrichaceae | https://lpsn.dsmz.de/family/erysipelotrichaceae |
| Erythrobacteraceae | https://lpsn.dsmz.de/family/erythrobacteraceae |
| Eubacteriaceae | https://lpsn.dsmz.de/family/eubacteriaceae |
| Euzebyaceae | https://lpsn.dsmz.de/family/euzebyaceae |
| Fastidiosibacteraceae | https://lpsn.dsmz.de/family/fastidiosibacteraceae |
| Ferrimonadaceae | https://lpsn.dsmz.de/family/ferrimonadaceae |
| Ferroplasmaceae | https://lpsn.dsmz.de/family/ferroplasmataceae |
| Ferroplasmataceae | https://lpsn.dsmz.de/family/ferroplasmataceae |
| Fervidicoccaceae | https://lpsn.dsmz.de/family/fervidicoccaceae |
| Fervidobacteriaceae | https://lpsn.dsmz.de/family/fervidobacteriaceae |
| Fibrobacteraceae | https://lpsn.dsmz.de/family/fibrobacteraceae |
| Filobacteriaceae | https://lpsn.dsmz.de/family/filobacteriaceae |
| Fimbriimonadaceae | https://lpsn.dsmz.de/family/fimbriimonadaceae |
| Flammeovirgaceae | https://lpsn.dsmz.de/family/flammeovirgaceae |
| Flavobacteriaceae | https://lpsn.dsmz.de/family/flavobacteriaceae |
| Flexibacteraceae | https://lpsn.dsmz.de/family/flexibacteraceae |
| Flexistipitaceae | https://lpsn.dsmz.de/family/deferribacteraceae |
| Fluviibacteraceae | https://lpsn.dsmz.de/family/fluviibacteraceae |
| Francisellaceae | https://lpsn.dsmz.de/family/francisellaceae |
| Frankiaceae | https://lpsn.dsmz.de/family/frankiaceae |
| Fulvivirgaceae | https://lpsn.dsmz.de/family/fulvivirgaceae |
| Fusobacteriaceae | https://lpsn.dsmz.de/family/fusobacteriaceae |
| Gaiellaceae | https://lpsn.dsmz.de/family/gaiellaceae |
| Gallionellaceae | https://lpsn.dsmz.de/family/gallionellaceae |
| Geminicoccaceae | https://lpsn.dsmz.de/family/geminicoccaceae |
| Geminocystaceae | https://lpsn.dsmz.de/family/geminocystaceae |
| Gemmataceae | https://lpsn.dsmz.de/family/gemmataceae |
| Gemmatimonadaceae | https://lpsn.dsmz.de/family/gemmatimonadaceae |
| Geoalkalibacteraceae | https://lpsn.dsmz.de/family/desulfuromonadaceae |
| Geobacteraceae | https://lpsn.dsmz.de/family/geobacteraceae |
| Geodermatophilaceae | https://lpsn.dsmz.de/family/geodermatophilaceae |
| Geopsychrobacteraceae | https://lpsn.dsmz.de/family/geobacteraceae |
| Geothermobacteraceae | https://lpsn.dsmz.de/family/geobacteraceae |
| Geovibrionaceae | https://lpsn.dsmz.de/family/geovibrionaceae |
| Gloeocapsaceae | https://lpsn.dsmz.de/family/chroococcaceae |
| Glycomycetaceae | https://lpsn.dsmz.de/family/glycomycetaceae |
| Gordoniaceae | https://lpsn.dsmz.de/family/gordoniaceae |
| Gottschalkiaceae | https://lpsn.dsmz.de/family/gottschalkiaceae |
| Gracilibacteraceae | https://lpsn.dsmz.de/family/gracilibacteraceae |

| | |
|---|---|
| Granulosicoccaceae | https://lpsn.dsmz.de/family/granulosicoccaceae |
| Gynuellaceae | https://lpsn.dsmz.de/family/gynuellaceae |
| Hafniaceae | https://lpsn.dsmz.de/family/hafniaceae |
| Hahellaceae | https://lpsn.dsmz.de/family/hahellaceae |
| Halanaerobiaceae | https://lpsn.dsmz.de/family/halanaerobiaceae |
| Haliangiaceae | https://lpsn.dsmz.de/family/kofleriaceae |
| Haliangiaceae | https://lpsn.dsmz.de/family/kofleriaceae |
| Halieaceae | https://lpsn.dsmz.de/family/halieaceae |
| Haliscomenobacteraceae | https://lpsn.dsmz.de/family/lewinellaceae |
| Haloanaerobiaceae | https://lpsn.dsmz.de/family/halanaerobiaceae |
| Haloarculaceae | https://lpsn.dsmz.de/family/haloarculaceae |
| Halobacteriaceae | https://lpsn.dsmz.de/family/halobacteriaceae |
| Halobacteriovoracaceae | https://lpsn.dsmz.de/family/halobacteriovoracaceae |
| Halobacteriovoraceae | https://lpsn.dsmz.de/family/halobacteriovoracaceae |
| Halobacteroidaceae | https://lpsn.dsmz.de/family/halobacteroidaceae |
| Halococcaceae | https://lpsn.dsmz.de/family/halococcaceae |
| Haloferacaceae | https://lpsn.dsmz.de/family/haloferacaceae |
| Halomonadaceae | https://lpsn.dsmz.de/family/halomonadaceae |
| Haloplasmataceae | https://lpsn.dsmz.de/family/haloplasmataceae |
| Halorhodospiraceae | https://lpsn.dsmz.de/family/halorhodospiraceae |
| Halorubraceae | https://lpsn.dsmz.de/family/halorubraceae |
| Halothiobacillaceae | https://lpsn.dsmz.de/family/halothiobacillaceae |
| Helicobacteraceae | https://lpsn.dsmz.de/family/helicobacteraceae |
| Heliobacteriaceae | https://lpsn.dsmz.de/family/heliobacteriaceae |
| Herpetosiphonaceae | https://lpsn.dsmz.de/family/herpetosiphonaceae |
| Hippeaceae | https://lpsn.dsmz.de/family/desulfurellaceae |
| Holophagaceae | https://lpsn.dsmz.de/family/holophagaceae |
| Holosporaceae | https://lpsn.dsmz.de/family/holosporaceae |
| Hoyosellaceae | https://lpsn.dsmz.de/family/nocardiaceae |
| Hungateiclostridiaceae | https://lpsn.dsmz.de/family/oscillospiraceae |
| Hydrogenimonaceae | https://lpsn.dsmz.de/family/hydrogenimonadaceae |
| Hydrogenimonadaceae | https://lpsn.dsmz.de/family/hydrogenimonadaceae |
| Hydrogenophilaceae | https://lpsn.dsmz.de/family/hydrogenophilaceae |
| Hydrogenothermaceae | https://lpsn.dsmz.de/family/hydrogenothermaceae |
| Hymenobacteraceae | https://lpsn.dsmz.de/family/hymenobacteraceae |
| Hyphomicrobiaceae | https://lpsn.dsmz.de/family/hyphomicrobiaceae |
| Hyphomonadaceae | https://lpsn.dsmz.de/family/hyphomonadaceae |
| Iamiaceae | https://lpsn.dsmz.de/family/iamiaceae |
| Ichthyobacteriaceae | https://lpsn.dsmz.de/family/ichthyobacteriaceae |
| Idiomarinaceae | https://lpsn.dsmz.de/family/idiomarinaceae |
| Ignatzschineriaceae | https://lpsn.dsmz.de/family/ignatzschineriaceae |
| Ignavibacteriaceae | https://lpsn.dsmz.de/family/ignavibacteriaceae |
| Ilumatobacteraceae | https://lpsn.dsmz.de/family/ilumatobacteraceae |
| Immundisolibacteraceae | https://lpsn.dsmz.de/family/immundisolibacteraceae |

| | |
|---|---|
| Intrasporangiaceae | https://lpsn.dsmz.de/family/intrasporangiaceae |
| Iodidimonadaceae | https://lpsn.dsmz.de/family/iodidimonadaceae |
| Isosphaeraceae | https://lpsn.dsmz.de/family/isosphaeraceae |
| Jatrophihabitantaceae | https://lpsn.dsmz.de/family/jatrophihabitantaceae |
| Jiangellaceae | https://lpsn.dsmz.de/family/jiangellaceae |
| Jonesiaceae | https://lpsn.dsmz.de/family/jonesiaceae |
| Kaistiaceae | https://lpsn.dsmz.de/family/kaistiaceae |
| Kallotenuaceae | https://lpsn.dsmz.de/family/kallotenuaceae |
| Kangiellaceae | https://lpsn.dsmz.de/family/kangiellaceae |
| Kiloniellaceae | https://lpsn.dsmz.de/family/kiloniellaceae |
| Kineosporiaceae | https://lpsn.dsmz.de/family/kineosporiaceae |
| Kiritimatiellaceae | https://lpsn.dsmz.de/family/kiritimatiellaceae |
| Kofleriaceae | https://lpsn.dsmz.de/family/kofleriaceae |
| Koleobacteraceae | https://lpsn.dsmz.de/family/koleobacteraceae |
| Kordiimonadaceae | https://lpsn.dsmz.de/family/temperatibacteraceae |
| Kosmotogaceae | https://lpsn.dsmz.de/family/kosmotogaceae |
| Kribbellaceae | https://lpsn.dsmz.de/family/kribbellaceae |
| Ktedobacteraceae | https://lpsn.dsmz.de/family/ktedonobacteraceae |
| Ktedonobacteraceae | https://lpsn.dsmz.de/family/ktedonobacteraceae |
| Ktedonosporobacteraceae | https://lpsn.dsmz.de/family/ktedonosporobacteraceae |
| Kytococcaceae | https://lpsn.dsmz.de/family/kytococcaceae |
| Labilitrichaceae | https://lpsn.dsmz.de/family/labilitrichaceae |
| Lachnospiraceae | https://lpsn.dsmz.de/family/lachnospiraceae |
| Lacipirellulaceae | https://lpsn.dsmz.de/family/lacipirellulaceae |
| Lactobacillaceae | https://lpsn.dsmz.de/family/lactobacillaceae |
| Lawsonellaceae | https://lpsn.dsmz.de/family/lawsonellaceae |
| Leeiaceae | https://lpsn.dsmz.de/family/chromobacteriaceae |
| Legionellaceae | https://lpsn.dsmz.de/family/legionellaceae |
| Lentimicrobiaceae | https://lpsn.dsmz.de/family/lentimicrobiaceae |
| Lentisphaeraceae | https://lpsn.dsmz.de/family/lentisphaeraceae |
| Leptolyngbyaceae | https://lpsn.dsmz.de/family/leptolyngbyaceae |
| Leptospiraceae | https://lpsn.dsmz.de/family/leptospiraceae |
| Leptotrichiaceae | https://lpsn.dsmz.de/family/leptotrichiaceae |
| Leuconostocaceae | https://lpsn.dsmz.de/family/leuconostocaceae |
| Leucotrichaceae | https://lpsn.dsmz.de/family/thiotrichaceae |
| Leucotrichaceae | https://lpsn.dsmz.de/family/thiotrichaceae |
| Lewinellaceae | https://lpsn.dsmz.de/family/lewinellaceae |
| Lichenibacteriaceae | https://lpsn.dsmz.de/family/lichenibacteriaceae |
| Lichenihabitantaceae | https://lpsn.dsmz.de/family/lichenihabitantaceae |
| Limnochordaceae | https://lpsn.dsmz.de/family/limnochordaceae |
| Listeriaceae | https://lpsn.dsmz.de/family/listeriaceae |
| Litoricolaceae | https://lpsn.dsmz.de/family/litoricolaceae |
| Longimicrobiaceae | https://lpsn.dsmz.de/family/longimicrobiaceae |
| Lujinxingiaceae | https://lpsn.dsmz.de/family/bradymonadaceae |

| | |
|---|---|
| Luteibaculaceae | https://lpsn.dsmz.de/family/cryomorphaceae |
| Lyngbyaceae | https://lpsn.dsmz.de/family/oscillatoriaceae |
| Lysobacteraceae | https://lpsn.dsmz.de/family/lysobacteraceae |
| Mabikibacteraceae | https://lpsn.dsmz.de/family/notoacmeibacteraceae |
| Magnetococcaceae | https://lpsn.dsmz.de/family/magnetococcaceae |
| Mangrovivirgaceae | https://lpsn.dsmz.de/family/mangrovivirgaceae |
| Maricaulaceae | https://lpsn.dsmz.de/family/maricaulaceae |
| Marinifilaceae | https://lpsn.dsmz.de/family/marinifilaceae |
| Marinilabiliaceae | https://lpsn.dsmz.de/family/marinilabiliaceae |
| Marinobacteraceae | https://lpsn.dsmz.de/family/alteromonadaceae |
| Mariprofundaceae | https://lpsn.dsmz.de/family/mariprofundaceae |
| Marivirgaceae | https://lpsn.dsmz.de/family/marivirgaceae |
| Meganemaceae | https://lpsn.dsmz.de/family/neomegalonemataceae |
| Meganemataceae | https://lpsn.dsmz.de/family/neomegalonemataceae |
| Melioribacteraceae | https://lpsn.dsmz.de/family/melioribacteraceae |
| Mesoaciditogaceae | https://lpsn.dsmz.de/family/mesoaciditogaceae |
| Metamycoplasmataceae | https://lpsn.dsmz.de/family/metamycoplasmataceae |
| Methanobacteriaceae | https://lpsn.dsmz.de/family/methanobacteriaceae |
| Methanocalculaceae | https://lpsn.dsmz.de/family/methanocalculaceae |
| Methanocaldococcaceae | https://lpsn.dsmz.de/family/methanocaldococcaceae |
| Methanocellaceae | https://lpsn.dsmz.de/family/methanocellaceae |
| Methanococcaceae | https://lpsn.dsmz.de/family/methanococcaceae |
| Methanocorpusculaceae | https://lpsn.dsmz.de/family/methanocorpusculaceae |
| Methanomassiliicoccaceae | https://lpsn.dsmz.de/family/methanomassiliicoccaceae |
| Methanomicrobiaceae | https://lpsn.dsmz.de/family/methanomicrobiaceae |
| Methanonatronarchaeaceae | https://lpsn.dsmz.de/family/methanonatronarchaeaceae |
| Methanoplanaceae | https://lpsn.dsmz.de/family/methanomicrobiaceae |
| Methanopyraceae | https://lpsn.dsmz.de/family/methanopyraceae |
| Methanoregulaceae | https://lpsn.dsmz.de/family/methanoregulaceae |
| Methanosaetaceae | https://lpsn.dsmz.de/family/methanosaetaceae |
| Methanosarcinaceae | https://lpsn.dsmz.de/family/methanosarcinaceae |
| Methanospirillaceae | https://lpsn.dsmz.de/family/methanospirillaceae |
| Methanothermaceae | https://lpsn.dsmz.de/family/methanothermaceae |
| Methanotrichaceae | https://lpsn.dsmz.de/family/methanosaetaceae |
| Methanotrichaceae | https://lpsn.dsmz.de/family/methanosaetaceae |
| Methermicoccaceae | https://lpsn.dsmz.de/family/methermicoccaceae |
| Methylobacteriaceae | https://lpsn.dsmz.de/family/methylobacteriaceae |
| Methylococcaceae | https://lpsn.dsmz.de/family/methylococcaceae |
| Methylocystaceae | https://lpsn.dsmz.de/family/methylocystaceae |
| Methylomonadaceae | https://lpsn.dsmz.de/family/methylococcaceae |
| Methylophilaceae | https://lpsn.dsmz.de/family/methylophilaceae |
| Methylothermaceae | https://lpsn.dsmz.de/family/methylothermaceae |
| Microbacteriaceae | https://lpsn.dsmz.de/family/microbacteriaceae |
| Microbulbiferaceae | https://lpsn.dsmz.de/family/microbulbiferaceae |

| | |
|---|---|
| Micrococcaceae | https://lpsn.dsmz.de/family/micrococcaceae |
| Micromonosporaceae | https://lpsn.dsmz.de/family/micromonosporaceae |
| Micropepsaceae | https://lpsn.dsmz.de/family/micropepsaceae |
| Microscillaceae | https://lpsn.dsmz.de/family/microscillaceae |
| Microsphaeraceae | https://lpsn.dsmz.de/family/nakamurellaceae |
| Microvenatoraceae | https://lpsn.dsmz.de/family/microvenatoraceae |
| Miltoncostaeaceae | https://lpsn.dsmz.de/family/miltoncostaeaceae |
| Minwuiaceae | https://lpsn.dsmz.de/family/minwuiaceae |
| Mogibacteriaceae | https://lpsn.dsmz.de/family/mogibacteriaceae |
| Mooreiaceae | https://lpsn.dsmz.de/family/mooreiaceae |
| Moorellaceae | https://lpsn.dsmz.de/family/thermoanaerobacteraceae |
| Moraxellaceae | https://lpsn.dsmz.de/family/moraxellaceae |
| Morganellaceae | https://lpsn.dsmz.de/family/morganellaceae |
| Moritellaceae | https://lpsn.dsmz.de/family/moritellaceae |
| Motilibacteraceae | https://lpsn.dsmz.de/family/motilibacteraceae |
| Mucispirillaceae | https://lpsn.dsmz.de/family/deferribacteraceae |
| Muribaculaceae | https://lpsn.dsmz.de/family/muribaculaceae |
| Mycobacteriaceae | https://lpsn.dsmz.de/family/mycobacteriaceae |
| Mycoplasmataceae | https://lpsn.dsmz.de/family/mycoplasmataceae |
| Mycoplasmoidaceae | https://lpsn.dsmz.de/family/mycoplasmoidaceae |
| Myxococcaceae | https://lpsn.dsmz.de/family/myxococcaceae |
| Mzabimycetaceae | https://lpsn.dsmz.de/family/pseudonocardiaceae |
| Nakamurellaceae | https://lpsn.dsmz.de/family/nakamurellaceae |
| Nannocystaceae | https://lpsn.dsmz.de/family/nannocystaceae |
| Nanobdellaceae | https://lpsn.dsmz.de/family/nanobdellaceae |
| Natranaerobiaceae | https://lpsn.dsmz.de/family/natranaerobiaceae |
| Natranaerofabaceae | https://lpsn.dsmz.de/family/natranaerofabaceae |
| Natrialbaceae | https://lpsn.dsmz.de/family/natrialbaceae |
| Natronoarchaeaceae | https://lpsn.dsmz.de/family/halobacteriaceae |
| Natronospirillaceae | https://lpsn.dsmz.de/family/natronospirillaceae |
| Nautiliaceae | https://lpsn.dsmz.de/family/nautiliaceae |
| Neisseriaceae | https://lpsn.dsmz.de/family/neisseriaceae |
| Neomegalonemataceae | https://lpsn.dsmz.de/family/neomegalonemataceae |
| Nevskiaceae | https://lpsn.dsmz.de/family/nevskiaceae |
| Nitratiruptoraceae | https://lpsn.dsmz.de/family/nitratiruptoraceae |
| Nitriliruptoraceae | https://lpsn.dsmz.de/family/nitriliruptoraceae |
| Nitrobacteraceae | https://lpsn.dsmz.de/family/nitrobacteraceae |
| Nitrobacteriaceae | https://lpsn.dsmz.de/family/nitrobacteraceae |
| Nitrosomonadaceae | https://lpsn.dsmz.de/family/nitrosomonadaceae |
| Nitrosopumilaceae | https://lpsn.dsmz.de/family/nitrosopumilaceae |
| Nitrososphaeraceae | https://lpsn.dsmz.de/family/nitrososphaeraceae |
| Nitrospinaceae | https://lpsn.dsmz.de/family/nitrospinaceae |
| Nitrospiraceae | https://lpsn.dsmz.de/family/nitrospiraceae |
| Nocardiaceae | https://lpsn.dsmz.de/family/nocardiaceae |

| | |
|---|---|
| Nocardioidaceae | https://lpsn.dsmz.de/family/nocardioidaceae |
| Nocardiopsaceae | https://lpsn.dsmz.de/family/nocardiopsidaceae |
| Nocardiopsidaceae | https://lpsn.dsmz.de/family/nocardiopsidaceae |
| Notoacmeibacteraceae | https://lpsn.dsmz.de/family/notoacmeibacteraceae |
| Novibacillaceae | https://lpsn.dsmz.de/family/thermoactinomycetaceae |
| Oceanospirillaceae | https://lpsn.dsmz.de/family/oceanospirillaceae |
| Odoribacteraceae | https://lpsn.dsmz.de/family/odoribacteraceae |
| Oerskoviaceae | https://lpsn.dsmz.de/family/promicromonosporaceae |
| Oleiphilaceae | https://lpsn.dsmz.de/family/oleiphilaceae |
| Oligoflexaceae | https://lpsn.dsmz.de/family/oligoflexaceae |
| Oligosphaeraceae | https://lpsn.dsmz.de/family/oligosphaeraceae |
| Opitutaceae | https://lpsn.dsmz.de/family/opitutaceae |
| Orbaceae | https://lpsn.dsmz.de/family/orbaceae |
| Ornithinimicrobiaceae | https://lpsn.dsmz.de/family/ornithinimicrobiaceae |
| Oscillariaceae | https://lpsn.dsmz.de/family/oscillatoriaceae |
| Oscillatoriaceae | https://lpsn.dsmz.de/family/oscillatoriaceae |
| Oscillochloridaceae | https://lpsn.dsmz.de/family/oscillochloridaceae |
| Oscillospiraceae | https://lpsn.dsmz.de/family/oscillospiraceae |
| Ostreibacteriaceae | https://lpsn.dsmz.de/family/ostreibacteriaceae |
| Owenweeksiaceae | https://lpsn.dsmz.de/family/schleiferiaceae |
| Oxalobacteraceae | https://lpsn.dsmz.de/family/oxalobacteraceae |
| Paenibacillaceae | https://lpsn.dsmz.de/family/paenibacillaceae |
| Paludibacteraceae | https://lpsn.dsmz.de/family/paludibacteraceae |
| Parachlamydiaceae | https://lpsn.dsmz.de/family/parachlamydiaceae |
| Paracoccaceae | https://lpsn.dsmz.de/family/paracoccaceae |
| Paraconexibacteraceae | https://lpsn.dsmz.de/family/paraconexibacteraceae |
| Parvibaculaceae | https://lpsn.dsmz.de/family/parvibaculaceae |
| Parvicellaceae | https://lpsn.dsmz.de/family/parvicellaceae |
| Parviterribacteraceae | https://lpsn.dsmz.de/family/parviterribacteraceae |
| Parvularculaceae | https://lpsn.dsmz.de/family/parvularculaceae |
| Pasteurellaceae | https://lpsn.dsmz.de/family/pasteurellaceae |
| Pasteuriaceae | https://lpsn.dsmz.de/family/pasteuriaceae |
| Patulibacteraceae | https://lpsn.dsmz.de/family/patulibacteraceae |
| Pectobacteriaceae | https://lpsn.dsmz.de/family/pectobacteriaceae |
| Pelobacteraceae | https://lpsn.dsmz.de/family/desulfuromonadaceae |
| Pelotomaculaceae | https://lpsn.dsmz.de/family/peptococcaceae |
| Peptococcaceae | https://lpsn.dsmz.de/family/peptococcaceae |
| Peptoniphilaceae | https://lpsn.dsmz.de/family/peptoniphilaceae |
| Peptostreptococcaceae | https://lpsn.dsmz.de/family/peptostreptococcaceae |
| Peredibacteraceae | https://lpsn.dsmz.de/family/peredibacteraceae |
| Perlucidibacaceae | https://lpsn.dsmz.de/family/moraxellaceae |
| Persicobacteraceae | https://lpsn.dsmz.de/family/persicobacteraceae |
| Petrotogaceae | https://lpsn.dsmz.de/family/petrotogaceae |
| Phaeocystidibacteraceae | https://lpsn.dsmz.de/family/cryomorphaceae |

| | |
|---|---|
| Phaselicystaceae | https://lpsn.dsmz.de/family/phaselicystaceae |
| Phaselicystidaceae | https://lpsn.dsmz.de/family/phaselicystaceae |
| Phreatobacteraceae | https://lpsn.dsmz.de/family/phreatobacteraceae |
| Phycisphaeraceae | https://lpsn.dsmz.de/family/phycisphaeraceae |
| Phyllobacteriaceae | https://lpsn.dsmz.de/family/phyllobacteriaceae |
| Phytomonosporaceae | https://lpsn.dsmz.de/family/micromonosporaceae |
| Picrophilaceae | https://lpsn.dsmz.de/family/picrophilaceae |
| Pillotinaceae | https://lpsn.dsmz.de/family/spirochaetaceae |
| Pirellulaceae | https://lpsn.dsmz.de/family/pirellulaceae |
| Piscirickettsiaceae | https://lpsn.dsmz.de/family/piscirickettsiaceae |
| Planctomycetaceae | https://lpsn.dsmz.de/family/planctomycetaceae |
| Planococcaceae | https://lpsn.dsmz.de/family/caryophanaceae |
| Plectonemataceae | https://lpsn.dsmz.de/family/oscillatoriaceae |
| Pleioneaceae | https://lpsn.dsmz.de/family/kangiellaceae |
| Pleomorphomonadaceae | https://lpsn.dsmz.de/family/pleomorphomonadaceae |
| Pleurococcaceae | https://lpsn.dsmz.de/family/chroococcaceae |
| Polyangiaceae | https://lpsn.dsmz.de/family/polyangiaceae |
| Pontiellaceae | https://lpsn.dsmz.de/family/pontiellaceae |
| Porphyromonadaceae | https://lpsn.dsmz.de/family/porphyromonadaceae |
| Porticoccaceae | https://lpsn.dsmz.de/family/porticoccaceae |
| Prevotellaceae | https://lpsn.dsmz.de/family/prevotellaceae |
| Prochloraceae | https://lpsn.dsmz.de/family/prochloraceae |
| Prochlorotrichaceae | https://lpsn.dsmz.de/family/pseudanabaenaceae |
| Prolixibacteraceae | https://lpsn.dsmz.de/family/prolixibacteraceae |
| Promicromonosporaceae | https://lpsn.dsmz.de/family/promicromonosporaceae |
| Propionibacteriaceae | https://lpsn.dsmz.de/family/propionibacteriaceae |
| Proteinivoracaceae | https://lpsn.dsmz.de/family/proteinivoracaceae |
| Proteinivoraceae | https://lpsn.dsmz.de/family/proteinivoracaceae |
| Pseudanabaenaceae | https://lpsn.dsmz.de/family/pseudanabaenaceae |
| Pseudoalteromonadaceae | https://lpsn.dsmz.de/family/pseudoalteromonadaceae |
| Pseudobacteriovoracaceae | https://lpsn.dsmz.de/family/pseudobacteriovoracaceae |
| Pseudobdellovibrionaceae | https://lpsn.dsmz.de/family/pseudobdellovibrionaceae |
| Pseudomonadaceae | https://lpsn.dsmz.de/family/pseudomonadaceae |
| Pseudonocardiaceae | https://lpsn.dsmz.de/family/pseudonocardiaceae |
| Pseudopelobacteraceae | https://lpsn.dsmz.de/family/desulfuromonadaceae |
| Pseudoxanthobacteraceae | https://lpsn.dsmz.de/family/pseudoxanthobacteraceae |
| Psittacicellaceae | https://lpsn.dsmz.de/family/psittacicellaceae |
| Psychromonadaceae | https://lpsn.dsmz.de/family/psychromonadaceae |
| Puniceicoccaceae | https://lpsn.dsmz.de/family/puniceicoccaceae |
| Pyrinomonadaceae | https://lpsn.dsmz.de/family/pyrinomonadaceae |
| Pyrodictiaceae | https://lpsn.dsmz.de/family/pyrodictiaceae |
| Raineyaceae | https://lpsn.dsmz.de/family/raineyaceae |
| Rarobacteraceae | https://lpsn.dsmz.de/family/rarobacteraceae |
| Rectinemataceae | https://lpsn.dsmz.de/family/treponemataceae |

| | |
|---|---|
| Reichenbachiellaceae | https://lpsn.dsmz.de/family/reichenbachiellaceae |
| Reticulibacteraceae | https://lpsn.dsmz.de/family/reticulibacteraceae |
| Reyranellaceae | https://lpsn.dsmz.de/family/reyranellaceae |
| Rhabdaerophilaceae | https://lpsn.dsmz.de/family/rhabdaerophilaceae |
| Rhizobiaceae | https://lpsn.dsmz.de/family/rhizobiaceae |
| Rhodanobacteraceae | https://lpsn.dsmz.de/family/rhodanobacteraceae |
| Rhodobacteraceae | https://lpsn.dsmz.de/family/paracoccaceae |
| Rhodobiaceae | https://lpsn.dsmz.de/family/rhodobiaceae |
| Rhodocyclaceae | https://lpsn.dsmz.de/family/rhodocyclaceae |
| Rhodospirillaceae | https://lpsn.dsmz.de/family/rhodospirillaceae |
| Rhodothalassiaceae | https://lpsn.dsmz.de/family/rhodothalassiaceae |
| Rhodothermaceae | https://lpsn.dsmz.de/family/rhodothermaceae |
| Rhodovibrionaceae | https://lpsn.dsmz.de/family/rhodovibrionaceae |
| Rickettsiaceae | https://lpsn.dsmz.de/family/rickettsiaceae |
| Rikenellaceae | https://lpsn.dsmz.de/family/rikenellaceae |
| Ristellaceae | https://lpsn.dsmz.de/family/bacteroidaceae |
| Robiginitomaculaceae | https://lpsn.dsmz.de/family/hyphomonadaceae |
| Roseiarcaceae | https://lpsn.dsmz.de/family/roseiarcaceae |
| Roseiflexaceae | https://lpsn.dsmz.de/family/roseiflexaceae |
| Roseivirgaceae | https://lpsn.dsmz.de/family/roseivirgaceae |
| Roseobacteraceae | https://lpsn.dsmz.de/family/roseobacteraceae |
| Ruaniaceae | https://lpsn.dsmz.de/family/ruaniaceae |
| Rubricoccaceae | https://lpsn.dsmz.de/family/rubricoccaceae |
| Rubritaleaceae | https://lpsn.dsmz.de/family/rubritaleaceae |
| Rubrobacteraceae | https://lpsn.dsmz.de/family/rubrobacteraceae |
| Ruminococcaceae | https://lpsn.dsmz.de/family/oscillospiraceae |
| Saccharospirillaceae | https://lpsn.dsmz.de/family/saccharospirillaceae |
| Salibacteraceae | https://lpsn.dsmz.de/family/cryomorphaceae |
| Salinarimonadaceae | https://lpsn.dsmz.de/family/salinarimonadaceae |
| Salinibacteraceae | https://lpsn.dsmz.de/family/salinibacteraceae |
| Salinisphaeraceae | https://lpsn.dsmz.de/family/salinisphaeraceae |
| Salinivirgaceae | https://lpsn.dsmz.de/family/salinivirgaceae |
| Salisaetaceae | https://lpsn.dsmz.de/family/salisaetaceae |
| Salsipaludibacteraceae | https://lpsn.dsmz.de/family/salsipaludibacteraceae |
| Sandaracinaceae | https://lpsn.dsmz.de/family/sandaracinaceae |
| Sanguibacteraceae | https://lpsn.dsmz.de/family/jonesiaceae |
| Sapromycetaceae | https://lpsn.dsmz.de/family/acholeplasmataceae |
| Saprospiraceae | https://lpsn.dsmz.de/family/saprospiraceae |
| Sarcinaceae | https://lpsn.dsmz.de/family/clostridiaceae |
| Schleiferiaceae | https://lpsn.dsmz.de/family/schleiferiaceae |
| Sedimentisphaeraceae | https://lpsn.dsmz.de/family/sedimentisphaeraceae |
| Segniliparaceae | https://lpsn.dsmz.de/family/segniliparaceae |
| Segnochrobactraceae | https://lpsn.dsmz.de/family/segnochrobactraceae |
| Selenomonadaceae | https://lpsn.dsmz.de/family/selenomonadaceae |

| | |
|---|---|
| Shewanellaceae | https://lpsn.dsmz.de/family/shewanellaceae |
| Silvanigrellaceae | https://lpsn.dsmz.de/family/silvanigrellaceae |
| Simkaniaceae | https://lpsn.dsmz.de/family/simkaniaceae |
| Simonsiellaceae | https://lpsn.dsmz.de/family/neisseriaceae |
| Sinobacteraceae | https://lpsn.dsmz.de/family/nevskiaceae |
| Smithellaceae | https://lpsn.dsmz.de/family/syntrophaceae |
| Sneathiellaceae | https://lpsn.dsmz.de/family/sneathiellaceae |
| Sokoloviaceae | https://lpsn.dsmz.de/family/pseudanabaenaceae |
| Solimonadaceae | https://lpsn.dsmz.de/family/nevskiaceae |
| Solirubrobacteraceae | https://lpsn.dsmz.de/family/solirubrobacteraceae |
| Soortiaceae | https://lpsn.dsmz.de/family/soortiaceae |
| Sphaerobacteraceae | https://lpsn.dsmz.de/family/sphaerobacteraceae |
| Sphaerochaetaceae | https://lpsn.dsmz.de/family/sphaerochaetaceae |
| Sphaerotilaceae | https://lpsn.dsmz.de/family/comamonadaceae |
| Sphingobacteriaceae | https://lpsn.dsmz.de/family/sphingobacteriaceae |
| Sphingomonadaceae | https://lpsn.dsmz.de/family/sphingomonadaceae |
| Sphingosinicellaceae | https://lpsn.dsmz.de/family/sphingosinicellaceae |
| Spirillaceae | https://lpsn.dsmz.de/family/spirillaceae |
| Spirochaetaceae | https://lpsn.dsmz.de/family/spirochaetaceae |
| Spiroplasmataceae | https://lpsn.dsmz.de/family/spiroplasmataceae |
| Spirosomaceae | https://lpsn.dsmz.de/family/spirosomataceae |
| Spirosomataceae | https://lpsn.dsmz.de/family/spirosomataceae |
| Spirulinaceae | https://lpsn.dsmz.de/family/spirulinaceae |
| Spongiibacteraceae | https://lpsn.dsmz.de/family/spongiibacteraceae |
| Sporanaerobacteraceae | https://lpsn.dsmz.de/family/tissierellaceae |
| Sporichthyaceae | https://lpsn.dsmz.de/family/sporichthyaceae |
| Sporolactobacillaceae | https://lpsn.dsmz.de/family/sporolactobacillaceae |
| Sporomusaceae | https://lpsn.dsmz.de/family/sporomusaceae |
| Staphylococcaceae | https://lpsn.dsmz.de/family/staphylococcaceae |
| Stappiaceae | https://lpsn.dsmz.de/family/stappiaceae |
| Stellaceae | https://lpsn.dsmz.de/family/stellaceae |
| Steroidobacteraceae | https://lpsn.dsmz.de/family/steroidobacteraceae |
| Sterolibacteriaceae | https://lpsn.dsmz.de/family/sterolibacteriaceae |
| Streptococcaceae | https://lpsn.dsmz.de/family/streptococcaceae |
| Streptomycetaceae | https://lpsn.dsmz.de/family/streptomycetaceae |
| Streptosporangiaceae | https://lpsn.dsmz.de/family/streptosporangiaceae |
| Succinivibrionaceae | https://lpsn.dsmz.de/family/succinivibrionaceae |
| Sulfolobaceae | https://lpsn.dsmz.de/family/sulfolobaceae |
| Sulfuricellaceae | https://lpsn.dsmz.de/family/sulfuricellaceae |
| Sulfurimonadaceae | https://lpsn.dsmz.de/family/helicobacteraceae |
| Sulfurospirillaceae | https://lpsn.dsmz.de/family/campylobacteraceae |
| Sulfurovaceae | https://lpsn.dsmz.de/family/helicobacteraceae |
| Sutterellaceae | https://lpsn.dsmz.de/family/sutterellaceae |
| Symbiobacteriaceae | https://lpsn.dsmz.de/family/symbiobacteriaceae |

| | |
|---|---|
| Synechococcaceae | https://lpsn.dsmz.de/family/synechococcaceae |
| Synergistaceae | https://lpsn.dsmz.de/family/synergistaceae |
| Syntrophaceae | https://lpsn.dsmz.de/family/syntrophaceae |
| Syntrophobacteraceae | https://lpsn.dsmz.de/family/syntrophobacteraceae |
| Syntrophomonadaceae | https://lpsn.dsmz.de/family/syntrophomonadaceae |
| Syntrophorhabdaceae | https://lpsn.dsmz.de/family/syntrophorhabdaceae |
| Syntrophotaleaceae | https://lpsn.dsmz.de/family/syntrophotaleaceae |
| Tannerellaceae | https://lpsn.dsmz.de/family/tannerellaceae |
| Temperatibacteraceae | https://lpsn.dsmz.de/family/temperatibacteraceae |
| Tenuifilaceae | https://lpsn.dsmz.de/family/tenuifilaceae |
| Tepidamorphaceae | https://lpsn.dsmz.de/family/tepidamorphaceae |
| Tepidanaerobacteraceae | https://lpsn.dsmz.de/family/tepidanaerobacteraceae |
| Tepidiformaceae | https://lpsn.dsmz.de/family/tepidiformaceae |
| Tepidimicrobiaceae | https://lpsn.dsmz.de/family/tissierellaceae |
| Tepidisphaeraceae | https://lpsn.dsmz.de/family/tepidisphaeraceae |
| Terasakiellaceae | https://lpsn.dsmz.de/family/terasakiellaceae |
| Terminosporaceae | https://lpsn.dsmz.de/family/clostridiaceae |
| Terrimicrobiaceae | https://lpsn.dsmz.de/family/terrimicrobiaceae |
| Thalassobaculaceae | https://lpsn.dsmz.de/family/thalassobaculaceae |
| Thalassospiraceae | https://lpsn.dsmz.de/family/thalassospiraceae |
| Thermaceae | https://lpsn.dsmz.de/family/thermaceae |
| Thermincolaceae | https://lpsn.dsmz.de/family/peptococcaceae |
| Thermithiobacillaceae | https://lpsn.dsmz.de/family/thermithiobacillaceae |
| Thermoactinomycetaceae | https://lpsn.dsmz.de/family/thermoactinomycetaceae |
| Thermoanaerobacteraceae | https://lpsn.dsmz.de/family/thermoanaerobacteraceae |
| Thermoanaerobaculaceae | https://lpsn.dsmz.de/family/thermoanaerobaculaceae |
| Thermococcaceae | https://lpsn.dsmz.de/family/thermococcaceae |
| Thermodesulfatatoraceae | https://lpsn.dsmz.de/family/thermodesulfobacteriaceae |
| Thermodesulfobacteriaceae | https://lpsn.dsmz.de/family/thermodesulfobacteriaceae |
| Thermodesulfobiaceae | https://lpsn.dsmz.de/family/thermodesulfobiaceae |
| Thermodesulforhabdaceae | https://lpsn.dsmz.de/family/syntrophobacteraceae |
| Thermodesulfovibrionaceae | https://lpsn.dsmz.de/family/thermodesulfovibrionaceae |
| Thermofilaceae | https://lpsn.dsmz.de/family/thermofilaceae |
| Thermoflexaceae | https://lpsn.dsmz.de/family/thermoflexaceae |
| Thermoflexibacteraceae | https://lpsn.dsmz.de/family/thermoflexibacteraceae |
| Thermogemmatisporaceae | https://lpsn.dsmz.de/family/thermogemmatisporaceae |
| Thermoguttaceae | https://lpsn.dsmz.de/family/thermoguttaceae |
| Thermohalobacteraceae | https://lpsn.dsmz.de/family/clostridiaceae |
| Thermoleophilaceae | https://lpsn.dsmz.de/family/thermoleophilaceae |
| Thermolithobacteraceae | https://lpsn.dsmz.de/family/thermolithobacteraceae |
| Thermomicrobiaceae | https://lpsn.dsmz.de/family/thermomicrobiaceae |
| Thermomonosporaceae | https://lpsn.dsmz.de/family/thermomonosporaceae |
| Thermonemataceae | https://lpsn.dsmz.de/family/thermonemataceae |
| Thermoplasmataceae | https://lpsn.dsmz.de/family/thermoplasmataceae |

| | |
|---|---|
| Thermoproteaceae | https://lpsn.dsmz.de/family/thermoproteaceae |
| Thermosediminibacteraceae | https://lpsn.dsmz.de/family/thermosediminibacteraceae |
| Thermosporotrichaceae | https://lpsn.dsmz.de/family/thermosporotrichaceae |
| Thermostichaceae | https://lpsn.dsmz.de/family/[thermostichaceae]-no-family |
| Thermosynergistaceae | https://lpsn.dsmz.de/family/thermosynergistaceae |
| Thermotogaceae | https://lpsn.dsmz.de/family/thermotogaceae |
| Thermotomaculaceae | https://lpsn.dsmz.de/family/thermotomaculaceae |
| Thioalkalibacteraceae | https://lpsn.dsmz.de/family/thioalkalibacteraceae |
| Thioalkalispiraceae | https://lpsn.dsmz.de/family/thioalkalispiraceae |
| Thiobacillaceae | https://lpsn.dsmz.de/family/thiobacillaceae |
| Thiocapsaceae | https://lpsn.dsmz.de/family/chromatiaceae |
| Thiofilaceae | https://lpsn.dsmz.de/family/thiofilaceae |
| Thiolineaceae | https://lpsn.dsmz.de/family/thiolineaceae |
| Thioprofundaceae | https://lpsn.dsmz.de/family/thioprofundaceae |
| Thioreductoraceae | https://lpsn.dsmz.de/family/nautiliaceae |
| Thiotrichaceae | https://lpsn.dsmz.de/family/thiotrichaceae |
| Thiovulaceae | https://lpsn.dsmz.de/family/helicobacteraceae |
| Thorselliaceae | https://lpsn.dsmz.de/family/thorselliaceae |
| Tichowtungiaceae | https://lpsn.dsmz.de/family/tichowtungiaceae |
| Tissierellaceae | https://lpsn.dsmz.de/family/tissierellaceae |
| Tomitellaceae | https://lpsn.dsmz.de/family/nocardiaceae |
| Treboniaceae | https://lpsn.dsmz.de/family/treboniaceae |
| Treponemataceae | https://lpsn.dsmz.de/family/treponemataceae |
| Trichloromonadaceae | https://lpsn.dsmz.de/family/desulfuromonadaceae |
| Trueperaceae | https://lpsn.dsmz.de/family/trueperaceae |
| Tsukamurellaceae | https://lpsn.dsmz.de/family/tsukamurellaceae |
| Turicibacteraceae | https://lpsn.dsmz.de/family/turicibacteraceae |
| Usitatibacteraceae | https://lpsn.dsmz.de/family/usitatibacteraceae |
| Vallicoccaceae | https://lpsn.dsmz.de/family/vallicoccaceae |
| Vallitaleaceae | https://lpsn.dsmz.de/family/vallitaleaceae |
| Veillonellaceae | https://lpsn.dsmz.de/family/veillonellaceae |
| Ventosimonadaceae | https://lpsn.dsmz.de/family/ventosimonadaceae |
| Verrucomicrobiaceae | https://lpsn.dsmz.de/family/verrucomicrobiaceae |
| Vibrionaceae | https://lpsn.dsmz.de/family/vibrionaceae |
| Vicinamibacteraceae | https://lpsn.dsmz.de/family/vicinamibacteraceae |
| Vicingaceae | https://lpsn.dsmz.de/family/cryomorphaceae |
| Victivallaceae | https://lpsn.dsmz.de/family/victivallaceae |
| Vitreoscillaceae | https://lpsn.dsmz.de/family/neisseriaceae |
| Vulgatibacteraceae | https://lpsn.dsmz.de/family/vulgatibacteraceae |
| Waddliaceae | https://lpsn.dsmz.de/family/waddliaceae |
| Weeksellaceae | https://lpsn.dsmz.de/family/weeksellaceae |
| Wenzhouxiangellaceae | https://lpsn.dsmz.de/family/wenzhouxiangellaceae |
| Williamwhitmaniaceae | https://lpsn.dsmz.de/family/williamwhitmaniaceae |
| Woeseiaceae | https://lpsn.dsmz.de/family/woeseiaceae |

| | |
|---|---|
| Xanthobacteraceae | https://lpsn.dsmz.de/family/xanthobacteraceae |
| Xanthomonadaceae | https://lpsn.dsmz.de/family/lysobacteraceae |
| Xylanivirgaceae | https://lpsn.dsmz.de/family/xylanivirgaceae |
| Yaniaceae | https://lpsn.dsmz.de/family/micrococcaceae |
| Yaniellaceae | https://lpsn.dsmz.de/family/micrococcaceae |
| Yersiniaceae | https://lpsn.dsmz.de/family/yersiniaceae |
| Zavarziniaceae | https://lpsn.dsmz.de/family/zavarziniaceae |
| Zhaonellaceae | https://lpsn.dsmz.de/family/zhaonellaceae |
| Zoogloeaceae | https://lpsn.dsmz.de/family/zoogloeaceae |
| Zooshikellaceae | https://lpsn.dsmz.de/family/zooshikellaceae |
| Zymomonadaceae | https://lpsn.dsmz.de/family/zymomonadaceae |
| [Anaerolineae; not assigned to order] | https://lpsn.dsmz.de/order/anaerolineae-no-order |
| [Bacillota; not assigned to order] | https://lpsn.dsmz.de/order/bacillota-no-order |
| [Betaproteobacteria; not assigned to order] | https://lpsn.dsmz.de/order/betaproteobacteria-no-order |
| [Calditrichota; not assigned to order] | https://lpsn.dsmz.de/order/calditrichota-no-order |
| [Deltaproteobacteria; not assigned to order] | https://lpsn.dsmz.de/order/deltaproteobacteria-no-order |
| [Epsilonproteobacteria; not assigned to order] | https://lpsn.dsmz.de/order/epsilonproteobacteria-no-order |
| [Gammaproteobacteria; not assigned to order] | https://lpsn.dsmz.de/order/gammaproteobacteria-no-order |
| [Hydrogenophilia; not assigned to order] | https://lpsn.dsmz.de/order/hydrogenophilia-no-order |
| [Spirochaetia; not assigned to order] | https://lpsn.dsmz.de/order/spirochaetia-no-order |
| [Synergistota; not assigned to order] | https://lpsn.dsmz.de/order/synergistota-no-order |
| [Thermodesulfobacteriota; not assigned to order] | https://lpsn.dsmz.de/order/thermodesulfobacteriota-no-order |
| [Verrucomicrobiota; not assigned to order] | https://lpsn.dsmz.de/order/verrucomicrobiota-no-order |
| Abditibacteriales | https://lpsn.dsmz.de/order/abditibacteriales |
| Acanthopleuribacterales | https://lpsn.dsmz.de/order/acanthopleuribacterales |
| Acholeplasmatales | https://lpsn.dsmz.de/order/acholeplasmatales |
| Acidaminococcales | https://lpsn.dsmz.de/order/acidaminococcales |
| Acidiferrobacterales | https://lpsn.dsmz.de/order/acidiferrobacterales |
| Acidilobales | https://lpsn.dsmz.de/order/acidilobales |
| Acidimicrobiales | https://lpsn.dsmz.de/order/acidimicrobiales |
| Acidithiobacillales | https://lpsn.dsmz.de/order/acidithiobacillales |
| Acidobacteriales | https://lpsn.dsmz.de/order/terriglobales |
| Acidothermales | https://lpsn.dsmz.de/order/acidothermales |
| Actinocatenisporales | https://lpsn.dsmz.de/order/micromonosporales |
| Actinomycetales | https://lpsn.dsmz.de/order/actinomycetales |
| Actinoplanales | https://lpsn.dsmz.de/order/micromonosporales |
| Actinopolysporales | https://lpsn.dsmz.de/order/pseudonocardiales |
| Aeromonadales | https://lpsn.dsmz.de/order/aeromonadales |
| Aggregatilineales | https://lpsn.dsmz.de/order/aggregatilineales |
| Alteromonadales | https://lpsn.dsmz.de/order/alteromonadales |
| Anaerolineales | https://lpsn.dsmz.de/order/anaerolineales |
| Anaeroplasmatales | https://lpsn.dsmz.de/order/anaeroplasmatales |
| Antricoccales | https://lpsn.dsmz.de/order/geodermatophilales |
| Anulomycetales | https://lpsn.dsmz.de/order/mycoplasmoidales |

| | |
|---|---|
| Aquificales | https://lpsn.dsmz.de/order/aquificales |
| Aquipuribacterales | https://lpsn.dsmz.de/order/micrococcales |
| Archaeoglobales | https://lpsn.dsmz.de/order/archaeoglobales |
| Ardenticatenales | https://lpsn.dsmz.de/order/ardenticatenales |
| Arenicellales | https://lpsn.dsmz.de/order/arenicellales |
| Armatimonadales | https://lpsn.dsmz.de/order/armatimonadales |
| Atribacterales | https://lpsn.dsmz.de/order/atribacterales |
| Bacillales | https://lpsn.dsmz.de/order/caryophanales |
| Bacteriovoracales | https://lpsn.dsmz.de/order/bacteriovoracales |
| Bacteroidales | https://lpsn.dsmz.de/order/bacteroidales |
| Balneolales | https://lpsn.dsmz.de/order/balneolales |
| Bdellovibrionales | https://lpsn.dsmz.de/order/bdellovibrionales |
| Beggiatoales | https://lpsn.dsmz.de/order/thiotrichales |
| Beutenbergiales | https://lpsn.dsmz.de/order/micrococcales |
| Bifidobacteriales | https://lpsn.dsmz.de/order/bifidobacteriales |
| Blastocatellales | https://lpsn.dsmz.de/order/blastocatellales |
| Bogoriellales | https://lpsn.dsmz.de/order/micrococcales |
| Borrelomycetales | https://lpsn.dsmz.de/order/mycoplasmoidales |
| Brachyspirales | https://lpsn.dsmz.de/order/brachyspirales |
| Brachyspiriales | https://lpsn.dsmz.de/order/brachyspirales |
| Bradymonadales | https://lpsn.dsmz.de/order/bradymonadales |
| Brevibacteriales | https://lpsn.dsmz.de/order/micrococcales |
| Brevinematales | https://lpsn.dsmz.de/order/brevinematales |
| Bryobacterales | https://lpsn.dsmz.de/order/bryobacterales |
| Burkholderiales | https://lpsn.dsmz.de/order/burkholderiales |
| Caldilineales | https://lpsn.dsmz.de/order/caldilineales |
| Caldisericales | https://lpsn.dsmz.de/order/caldisericales |
| Calditrichales | https://lpsn.dsmz.de/order/calditrichales |
| Campylobacterales | https://lpsn.dsmz.de/order/campylobacterales |
| Capsulimonadales | https://lpsn.dsmz.de/order/capsulimonadales |
| Cardiobacteriales | https://lpsn.dsmz.de/order/cardiobacteriales |
| Caryophanales | https://lpsn.dsmz.de/order/caryophanales |
| Catenulisporales | https://lpsn.dsmz.de/order/catenulisporales |
| Caulobacterales | https://lpsn.dsmz.de/order/caulobacterales |
| Cellulomonadales | https://lpsn.dsmz.de/order/micrococcales |
| Cellvibrionales | https://lpsn.dsmz.de/order/cellvibrionales |
| Chitinispirillales | https://lpsn.dsmz.de/order/chitinispirillales |
| Chitinivibrionales | https://lpsn.dsmz.de/order/chitinivibrionales |
| Chitinophagales | https://lpsn.dsmz.de/order/chitinophagales |
| Chlamydiales | https://lpsn.dsmz.de/order/chlamydiales |
| Chlorobiales | https://lpsn.dsmz.de/order/chlorobiales |
| Chloroflexales | https://lpsn.dsmz.de/order/chloroflexales |
| Chromatiales | https://lpsn.dsmz.de/order/chromatiales |
| Chroococcales | https://lpsn.dsmz.de/order/chroococcales |

| | |
|---|---|
| Chrysiogenales | https://lpsn.dsmz.de/order/chrysiogenales |
| Chthoniobacterales | https://lpsn.dsmz.de/order/terrimicrobiales |
| Chthonomonadales | https://lpsn.dsmz.de/order/chthonomonadales |
| Clostridiales | https://lpsn.dsmz.de/order/eubacteriales |
| Conexivisphaerales | https://lpsn.dsmz.de/order/thermoproteales |
| Coprothermobacterales | https://lpsn.dsmz.de/order/coprothermobacterales |
| Coriobacteriales | https://lpsn.dsmz.de/order/coriobacteriales |
| Corynebacteriales | https://lpsn.dsmz.de/order/mycobacteriales |
| Cryptosporangiales | https://lpsn.dsmz.de/order/cryptosporangiales |
| Culicoidibacterales | https://lpsn.dsmz.de/order/culicoidibacterales |
| Cytophagales | https://lpsn.dsmz.de/order/cytophagales |
| Deferribacterales | https://lpsn.dsmz.de/order/deferribacterales |
| Deferrisomatales | https://lpsn.dsmz.de/order/deferrisomatales |
| Dehalococcoidales | https://lpsn.dsmz.de/order/dehalococcoidales |
| Deinococcales | https://lpsn.dsmz.de/order/deinococcales |
| Demequinales | https://lpsn.dsmz.de/order/micrococcales |
| Dermabacterales | https://lpsn.dsmz.de/order/micrococcales |
| Dermatophilales | https://lpsn.dsmz.de/order/micrococcales |
| Desulfarcales | https://lpsn.dsmz.de/order/desulfarculales |
| Desulfarculales | https://lpsn.dsmz.de/order/desulfarculales |
| Desulfatiglandales | https://lpsn.dsmz.de/order/desulfobacterales |
| Desulfobaccales | https://lpsn.dsmz.de/order/syntrophobacterales |
| Desulfobacterales | https://lpsn.dsmz.de/order/desulfobacterales |
| Desulfobulbales | https://lpsn.dsmz.de/order/desulfobacterales |
| Desulfomonilales | https://lpsn.dsmz.de/order/syntrophobacterales |
| Desulfotomaculales | https://lpsn.dsmz.de/order/eubacteriales |
| Desulfovibrionales | https://lpsn.dsmz.de/order/desulfovibrionales |
| Desulfurellales | https://lpsn.dsmz.de/order/desulfurellales |
| Desulfuribacillales | https://lpsn.dsmz.de/order/caryophanales |
| Desulfurobacteriales | https://lpsn.dsmz.de/order/desulfurobacteriales |
| Desulfurococcales | https://lpsn.dsmz.de/order/desulfurococcales |
| Desulfuromonadales | https://lpsn.dsmz.de/order/desulfuromonadales |
| Desulfuromonales | https://lpsn.dsmz.de/order/desulfuromonadales |
| Dictyoglomales | https://lpsn.dsmz.de/order/dictyoglomerales |
| Dictyoglomerales | https://lpsn.dsmz.de/order/dictyoglomerales |
| Dissulfuribacterales | https://lpsn.dsmz.de/order/dissulfuribacterales |
| Eggerthellales | https://lpsn.dsmz.de/order/eggerthellales |
| Egibacterales | https://lpsn.dsmz.de/order/egibacterales |
| Egicoccales | https://lpsn.dsmz.de/order/egicoccales |
| Elusimicrobiales | https://lpsn.dsmz.de/order/elusimicrobiales |
| Emcibacterales | https://lpsn.dsmz.de/order/emcibacterales |
| Endomicrobiales | https://lpsn.dsmz.de/order/endomicrobiales |
| Enterobacterales | https://lpsn.dsmz.de/order/enterobacterales |
| Enterobacteriales | https://lpsn.dsmz.de/order/enterobacterales |

| | |
|---|---|
| Entomoplasmatales | https://lpsn.dsmz.de/order/entomoplasmatales |
| Erysipelotrichales | https://lpsn.dsmz.de/order/erysipelotrichales |
| Eubacteriales | https://lpsn.dsmz.de/order/eubacteriales |
| Euzebyales | https://lpsn.dsmz.de/order/euzebyales |
| Fervidicoccales | https://lpsn.dsmz.de/order/fervidicoccales |
| Fibrobacterales | https://lpsn.dsmz.de/order/fibrobacterales |
| Fimbriimonadales | https://lpsn.dsmz.de/order/fimbriimonadales |
| Flavobacteriales | https://lpsn.dsmz.de/order/flavobacteriales |
| Frankiales | https://lpsn.dsmz.de/order/frankiales |
| Fusobacteriales | https://lpsn.dsmz.de/order/fusobacteriales |
| Gaiellales | https://lpsn.dsmz.de/order/gaiellales |
| Gemmatales | https://lpsn.dsmz.de/order/gemmatales |
| Gemmatimonadales | https://lpsn.dsmz.de/order/gemmatimonadales |
| Geobacterales | https://lpsn.dsmz.de/order/desulfuromonadales |
| Geodermatophilales | https://lpsn.dsmz.de/order/geodermatophilales |
| Geovibriales | https://lpsn.dsmz.de/order/deferribacterales |
| Glycomycetales | https://lpsn.dsmz.de/order/glycomycetales |
| Halanaerobiales | https://lpsn.dsmz.de/order/halanaerobiales |
| Haliangiales | https://lpsn.dsmz.de/order/myxococcales |
| Haloanaerobiales | https://lpsn.dsmz.de/order/halanaerobiales |
| Halobacteriales | https://lpsn.dsmz.de/order/halobacteriales |
| Haloferacales | https://lpsn.dsmz.de/order/haloferacales |
| Haloplasmatales | https://lpsn.dsmz.de/order/haloplasmatales |
| Heliobacteriales | https://lpsn.dsmz.de/order/eubacteriales |
| Herpetosiphonales | https://lpsn.dsmz.de/order/herpetosiphonales |
| Holophagales | https://lpsn.dsmz.de/order/holophagales |
| Holosporales | https://lpsn.dsmz.de/order/holosporales |
| Hydrogenophilales | https://lpsn.dsmz.de/order/hydrogenophilales |
| Hyphomicrobiales | https://lpsn.dsmz.de/order/hyphomicrobiales |
| Hyphomonadales | https://lpsn.dsmz.de/order/caulobacterales |
| Iamiales | https://lpsn.dsmz.de/order/acidimicrobiales |
| Ignavibacteriales | https://lpsn.dsmz.de/order/ignavibacteriales |
| Igneococcales | https://lpsn.dsmz.de/order/desulfurococcales |
| Immundisolibacterales | https://lpsn.dsmz.de/order/immundisolibacterales |
| Iodidimonadales | https://lpsn.dsmz.de/order/iodidimonadales |
| Isosphaerales | https://lpsn.dsmz.de/order/isosphaerales |
| Jatrophihabitantales | https://lpsn.dsmz.de/order/jatrophihabitantales |
| Jiangellales | https://lpsn.dsmz.de/order/jiangellales |
| Kallotenuales | https://lpsn.dsmz.de/order/kallotenuales |
| Kangiellales | https://lpsn.dsmz.de/order/oceanospirillales |
| Kiloniellales | https://lpsn.dsmz.de/order/rhodospirillales |
| Kineosporiales | https://lpsn.dsmz.de/order/kineosporiales |
| Kiritimatiellales | https://lpsn.dsmz.de/order/kiritimatiellales |
| Kitasatosporales | https://lpsn.dsmz.de/order/kitasatosporales |

| | |
|---|---|
| Koleobacterales | https://lpsn.dsmz.de/order/koleobacterales |
| Kordiimonadales | https://lpsn.dsmz.de/order/kordiimonadales |
| Kosmotogales | https://lpsn.dsmz.de/order/kosmotogales |
| Ktedobacterales | https://lpsn.dsmz.de/order/ktedonobacterales |
| Ktedonobacterales | https://lpsn.dsmz.de/order/ktedonobacterales |
| Lactobacillales | https://lpsn.dsmz.de/order/lactobacillales |
| Legionellales | https://lpsn.dsmz.de/order/legionellales |
| Lentisphaerales | https://lpsn.dsmz.de/order/lentisphaerales |
| Leptospirales | https://lpsn.dsmz.de/order/leptospirales |
| Leptospiriales | https://lpsn.dsmz.de/order/leptospirales |
| Limnochordales | https://lpsn.dsmz.de/order/limnochordales |
| Longimicrobiales | https://lpsn.dsmz.de/order/longimicrobiales |
| Lysobacterales | https://lpsn.dsmz.de/order/lysobacterales |
| Magnetococcales | https://lpsn.dsmz.de/order/magnetococcales |
| Maricaulales | https://lpsn.dsmz.de/order/caulobacterales |
| Marinilabiliales | https://lpsn.dsmz.de/order/bacteroidales |
| Mariprofundales | https://lpsn.dsmz.de/order/mariprofundales |
| Mesoaciditogales | https://lpsn.dsmz.de/order/mesoaciditogales |
| Methanobacteriales | https://lpsn.dsmz.de/order/methanobacteriales |
| Methanocellales | https://lpsn.dsmz.de/order/methanocellales |
| Methanococcales | https://lpsn.dsmz.de/order/methanococcales |
| Methanomassiliicoccales | https://lpsn.dsmz.de/order/methanomassiliicoccales |
| Methanomicrobiales | https://lpsn.dsmz.de/order/methanomicrobiales |
| Methanonatronarchaeales | https://lpsn.dsmz.de/order/methanonatronarchaeales |
| Methanopyrales | https://lpsn.dsmz.de/order/methanopyrales |
| Methanosarcinales | https://lpsn.dsmz.de/order/methanosarcinales |
| Methanotrichales | https://lpsn.dsmz.de/order/methanosarcinales |
| Methylococcales | https://lpsn.dsmz.de/order/methylococcales |
| Methylophilales | https://lpsn.dsmz.de/order/nitrosomonadales |
| Microbacteriales | https://lpsn.dsmz.de/order/micrococcales |
| Micrococcales | https://lpsn.dsmz.de/order/micrococcales |
| Micromonosporales | https://lpsn.dsmz.de/order/micromonosporales |
| Micropepsales | https://lpsn.dsmz.de/order/micropepsales |
| Miltoncostaeales | https://lpsn.dsmz.de/order/miltoncostaeales |
| Minwuiales | https://lpsn.dsmz.de/order/minwuiales |
| Moorellales | https://lpsn.dsmz.de/order/thermoanaerobacterales |
| Moraxellales | https://lpsn.dsmz.de/order/pseudomonadales |
| Motilibacterales | https://lpsn.dsmz.de/order/frankiales |
| Motilibacterales | https://lpsn.dsmz.de/order/frankiales |
| Mycobacteriales | https://lpsn.dsmz.de/order/mycobacteriales |
| Mycoplasmatales | https://lpsn.dsmz.de/order/mycoplasmatales |
| Mycoplasmoidales | https://lpsn.dsmz.de/order/mycoplasmoidales |
| Myxococcales | https://lpsn.dsmz.de/order/myxococcales |
| Nakamurellales | https://lpsn.dsmz.de/order/nakamurellales |

| | |
|---|---|
| Nannocystales | https://lpsn.dsmz.de/order/myxococcales |
| Nanobdellales | https://lpsn.dsmz.de/order/nanobdellales |
| Natranaerobiales | https://lpsn.dsmz.de/order/natranaerobiales |
| Natrialbales | https://lpsn.dsmz.de/order/natrialbales |
| Nautiliales | https://lpsn.dsmz.de/order/nautiliales |
| Neisseriales | https://lpsn.dsmz.de/order/neisseriales |
| Nevskiales | https://lpsn.dsmz.de/order/nevskiales |
| Nitriliruptorales | https://lpsn.dsmz.de/order/nitriliruptorales |
| Nitrosomonadales | https://lpsn.dsmz.de/order/nitrosomonadales |
| Nitrosopumilales | https://lpsn.dsmz.de/order/nitrosopumilales |
| Nitrososphaerales | https://lpsn.dsmz.de/order/nitrososphaerales |
| Nitrospinales | https://lpsn.dsmz.de/order/nitrospinales |
| Nitrospirales | https://lpsn.dsmz.de/order/nitrospirales |
| Oceanospirillales | https://lpsn.dsmz.de/order/oceanospirillales |
| Oligoflexales | https://lpsn.dsmz.de/order/oligoflexales |
| Oligosphaerales | https://lpsn.dsmz.de/order/oligosphaerales |
| Opitutales | https://lpsn.dsmz.de/order/opitutales |
| Orbales | https://lpsn.dsmz.de/order/orbales |
| Oscillatoriales | https://lpsn.dsmz.de/order/oscillatoriales |
| Parachlamydiales | https://lpsn.dsmz.de/order/chlamydiales |
| Parvularculales | https://lpsn.dsmz.de/order/parvularculales |
| Pasteurellales | https://lpsn.dsmz.de/order/pasteurellales |
| Petrotogales | https://lpsn.dsmz.de/order/petrotogales |
| Phycisphaerales | https://lpsn.dsmz.de/order/phycisphaerales |
| Phytomonosporales | https://lpsn.dsmz.de/order/micromonosporales |
| Picrophilales | https://lpsn.dsmz.de/order/thermoplasmatales |
| Pirellulales | https://lpsn.dsmz.de/order/pirellulales |
| Planctomycetales | https://lpsn.dsmz.de/order/planctomycetales |
| Plectridiales | https://lpsn.dsmz.de/order/caryophanales |
| Polyangiales | https://lpsn.dsmz.de/order/myxococcales |
| Prochlorales | https://lpsn.dsmz.de/order/synechococcales |
| Propionibacteriales | https://lpsn.dsmz.de/order/propionibacteriales |
| Pseudomonadales | https://lpsn.dsmz.de/order/pseudomonadales |
| Pseudonocardiales | https://lpsn.dsmz.de/order/pseudonocardiales |
| Puniceicoccales | https://lpsn.dsmz.de/order/puniceicoccales |
| Rhizobiales | https://lpsn.dsmz.de/order/hyphomicrobiales |
| Rhodobacterales | https://lpsn.dsmz.de/order/rhodobacterales |
| Rhodocyclales | https://lpsn.dsmz.de/order/rhodocyclales |
| Rhodospirillales | https://lpsn.dsmz.de/order/rhodospirillales |
| Rhodothalassiales | https://lpsn.dsmz.de/order/rhodothalassiales |
| Rhodothermales | https://lpsn.dsmz.de/order/rhodothermales |
| Rickettsiales | https://lpsn.dsmz.de/order/rickettsiales |
| Ruaniales | https://lpsn.dsmz.de/order/micrococcales |
| Rubrobacterales | https://lpsn.dsmz.de/order/rubrobacterales |

| | |
|---|---|
| Salsipaludibacterales | https://lpsn.dsmz.de/order/salsipaludibacterales |
| Saprospirales | https://lpsn.dsmz.de/order/saprospirales |
| Sedimentisphaerales | https://lpsn.dsmz.de/order/sedimentisphaerales |
| Selenomonadales | https://lpsn.dsmz.de/order/selenomonadales |
| Silvanigrellales | https://lpsn.dsmz.de/order/silvanigrellales |
| Sneathiellales | https://lpsn.dsmz.de/order/sneathiellales |
| Solirubrobacterales | https://lpsn.dsmz.de/order/solirubrobacterales |
| Sphaerobacterales | https://lpsn.dsmz.de/order/sphaerobacterales |
| Sphingobacteriales | https://lpsn.dsmz.de/order/sphingobacteriales |
| Sphingomonadales | https://lpsn.dsmz.de/order/sphingomonadales |
| Spirillales | https://lpsn.dsmz.de/order/nitrosomonadales |
| Spirochaetales | https://lpsn.dsmz.de/order/spirochaetales |
| Spirulinales | https://lpsn.dsmz.de/order/spirulinales |
| Sporichthyales | https://lpsn.dsmz.de/order/sporichthyales |
| Sporomusales | https://lpsn.dsmz.de/order/selenomonadales |
| Streptomycetales | https://lpsn.dsmz.de/order/kitasatosporales |
| Streptomycetales | https://lpsn.dsmz.de/order/kitasatosporales |
| Streptosporangiales | https://lpsn.dsmz.de/order/streptosporangiales |
| Sulfolobales | https://lpsn.dsmz.de/order/sulfolobales |
| Sulfuricellales | https://lpsn.dsmz.de/order/nitrosomonadales |
| Synechococcales | https://lpsn.dsmz.de/order/synechococcales |
| Synergistales | https://lpsn.dsmz.de/order/synergistales |
| Syntrophales | https://lpsn.dsmz.de/order/syntrophobacterales |
| Syntrophobacterales | https://lpsn.dsmz.de/order/syntrophobacterales |
| Syntrophorhabdales | https://lpsn.dsmz.de/order/syntrophorhabdales |
| Tepidiformales | https://lpsn.dsmz.de/order/tepidiformales |
| Tepidisphaerales | https://lpsn.dsmz.de/order/tepidisphaerales |
| Terriglobales | https://lpsn.dsmz.de/order/terriglobales |
| Terrimicrobiales | https://lpsn.dsmz.de/order/terrimicrobiales |
| Thermales | https://lpsn.dsmz.de/order/thermales |
| Thermoanaerobacterales | https://lpsn.dsmz.de/order/thermoanaerobacterales |
| Thermoanaerobaculales | https://lpsn.dsmz.de/order/thermoanaerobaculales |
| Thermococcales | https://lpsn.dsmz.de/order/thermococcales |
| Thermodesulfobacteriales | https://lpsn.dsmz.de/order/thermodesulfobacteriales |
| Thermodesulfobiales | https://lpsn.dsmz.de/order/thermoanaerobacterales |
| Thermodesulfovibrionales | https://lpsn.dsmz.de/order/thermodesulfovibrionales |
| Thermofilales | https://lpsn.dsmz.de/order/thermoproteales |
| Thermoflexales | https://lpsn.dsmz.de/order/thermoflexales |
| Thermogemmatisporales | https://lpsn.dsmz.de/order/thermogemmatisporales |
| Thermoleophilales | https://lpsn.dsmz.de/order/thermoleophilales |
| Thermolithobacterales | https://lpsn.dsmz.de/order/thermolithobacterales |
| Thermomicrobiales | https://lpsn.dsmz.de/order/thermomicrobiales |
| Thermoplasmatales | https://lpsn.dsmz.de/order/thermoplasmatales |
| Thermoproteales | https://lpsn.dsmz.de/order/thermoproteales |

| | |
|---|---|
| Thermosediminibacterales | https://lpsn.dsmz.de/order/thermosediminibacterales |
| Thermostichales | https://lpsn.dsmz.de/order/thermostichales |
| Thermotogales | https://lpsn.dsmz.de/order/thermotogales |
| Thermotomaculales | https://lpsn.dsmz.de/order/thermotomaculales |
| Thiotrichales | https://lpsn.dsmz.de/order/thiotrichales |
| Tichowtungiales | https://lpsn.dsmz.de/order/tichowtungiales |
| Tissierellales | https://lpsn.dsmz.de/order/tissierellales |
| Trueperales | https://lpsn.dsmz.de/order/trueperales |
| Veillonellales | https://lpsn.dsmz.de/order/veillonellales |
| Verrucomicrobiales | https://lpsn.dsmz.de/order/verrucomicrobiales |
| Vibrionales | https://lpsn.dsmz.de/order/vibrionales |
| Vicinamibacterales | https://lpsn.dsmz.de/order/vicinamibacterales |
| Victivallales | https://lpsn.dsmz.de/order/victivallales |
| Xanthomonadales | https://lpsn.dsmz.de/order/lysobacterales |
| [Bacillota; not assigned to class] | https://lpsn.dsmz.de/class/bacillota-no-class |
| [Calditrichota; not assigned to class] | https://lpsn.dsmz.de/class/calditrichota-no-class |
| [Synergistota; not assigned to class] | https://lpsn.dsmz.de/class/synergistota-no-class |
| [Thermodesulfobacteriota; not assigned to class] | https://lpsn.dsmz.de/class/thermodesulfobacteriota-no-class |
| [Verrucomicrobiota; not assigned to class] | https://lpsn.dsmz.de/class/verrucomicrobiota-no-class |
| Abditibacteria | https://lpsn.dsmz.de/class/abditibacteriia |
| Abditibacteriia | https://lpsn.dsmz.de/class/abditibacteriia |
| Acidimicrobiia | https://lpsn.dsmz.de/class/acidimicrobiia |
| Acidithiobacillia | https://lpsn.dsmz.de/class/acidithiobacillia |
| Acidobacteria | https://lpsn.dsmz.de/class/terriglobia |
| Acidobacteriae | https://lpsn.dsmz.de/class/terriglobia |
| Acidobacteriia | https://lpsn.dsmz.de/class/terriglobia |
| Actinobacteria | https://lpsn.dsmz.de/class/actinomycetes |
| Actinomycetes | https://lpsn.dsmz.de/class/actinomycetes |
| Actinomycetia | https://lpsn.dsmz.de/class/actinomycetes |
| Alphabacteria | https://lpsn.dsmz.de/class/alphaproteobacteria |
| Alphaproteobacteria | https://lpsn.dsmz.de/class/alphaproteobacteria |
| Anaerolineae | https://lpsn.dsmz.de/class/anaerolineae |
| Anoxyphotobacteria | https://lpsn.dsmz.de/class/alphaproteobacteria |
| Aquificae | https://lpsn.dsmz.de/class/aquificae |
| Aquificia | https://lpsn.dsmz.de/class/aquificae |
| Arabobacteria | https://lpsn.dsmz.de/class/actinomycetes |
| Archaeobacteria | https://lpsn.dsmz.de/class/methanobacteria |
| Archaeoglobea | https://lpsn.dsmz.de/class/archaeoglobi |
| Archaeoglobi | https://lpsn.dsmz.de/class/archaeoglobi |
| Ardenticatenia | https://lpsn.dsmz.de/class/ardenticatenia |
| Armatimonadia | https://lpsn.dsmz.de/class/armatimonadia |
| Arthrobacteria | https://lpsn.dsmz.de/class/actinomycetes |
| Atribacteria | https://lpsn.dsmz.de/class/atribacteria |
| Bacilli | https://lpsn.dsmz.de/class/bacilli |

| | |
|---|---|
| Bacillia | https://lpsn.dsmz.de/class/bacilli |
| Bacteriovoracia | https://lpsn.dsmz.de/class/oligoflexia |
| Bacteroidetia | https://lpsn.dsmz.de/class/bacteroidia |
| Bacteroidia | https://lpsn.dsmz.de/class/bacteroidia |
| Balneolia | https://lpsn.dsmz.de/class/balneolia |
| Bdellovibrionia | https://lpsn.dsmz.de/class/oligoflexia |
| Betaproteobacteria | https://lpsn.dsmz.de/class/betaproteobacteria |
| Blastocatellia | https://lpsn.dsmz.de/class/blastocatellia |
| Caldilineae | https://lpsn.dsmz.de/class/caldilineae |
| Caldilineia | https://lpsn.dsmz.de/class/caldilineae |
| Caldisericia | https://lpsn.dsmz.de/class/caldisericia |
| Calditrichae | https://lpsn.dsmz.de/class/calditrichia |
| Calditrichia | https://lpsn.dsmz.de/class/calditrichia |
| Campylobacteria | https://lpsn.dsmz.de/class/epsilonproteobacteria |
| Caulobacteria | https://lpsn.dsmz.de/class/alphaproteobacteria |
| Chitinispirillia | https://lpsn.dsmz.de/class/chitinispirillia |
| Chitinivibrionia | https://lpsn.dsmz.de/class/chitinivibrionia |
| Chitinophagia | https://lpsn.dsmz.de/class/chitinophagia |
| Chlamydiae | https://lpsn.dsmz.de/class/chlamydiia |
| Chlamydiia | https://lpsn.dsmz.de/class/chlamydiia |
| Chlorobacteria | https://lpsn.dsmz.de/class/chloroflexia |
| Chlorobea | https://lpsn.dsmz.de/class/chlorobiia |
| Chlorobia | https://lpsn.dsmz.de/class/chlorobiia |
| Chlorobia | https://lpsn.dsmz.de/class/chlorobiia |
| Chlorobiia | https://lpsn.dsmz.de/class/chlorobiia |
| Chloroflexi | https://lpsn.dsmz.de/class/chloroflexia |
| Chloroflexia | https://lpsn.dsmz.de/class/chloroflexia |
| Chromatibacteria | https://lpsn.dsmz.de/class/gammaproteobacteria |
| Chromatiia | https://lpsn.dsmz.de/class/gammaproteobacteria |
| Chroobacteria | https://lpsn.dsmz.de/class/cyanophyceae |
| Chrysiogenetes | https://lpsn.dsmz.de/class/chrysiogenetes |
| Chthonomonadetes | https://lpsn.dsmz.de/class/chthonomonadetes |
| Clostridia | https://lpsn.dsmz.de/class/clostridia |
| Clostridiia | https://lpsn.dsmz.de/class/clostridia |
| Conexivisphaeria | https://lpsn.dsmz.de/class/thermoprotei |
| Coprothermobacteria | https://lpsn.dsmz.de/class/coprothermobacteria |
| Coriobacteriia | https://lpsn.dsmz.de/class/coriobacteriia |
| Crenarchaeota | https://lpsn.dsmz.de/class/thermoprotei |
| Culicoidibacteria | https://lpsn.dsmz.de/class/culicoidibacteria |
| Cyanophyceae | https://lpsn.dsmz.de/class/cyanophyceae |
| Cytophagia | https://lpsn.dsmz.de/class/cytophagia |
| Deferribacteres | https://lpsn.dsmz.de/class/deferribacteres |
| Deferribacteria | https://lpsn.dsmz.de/class/deferribacteres |
| Deferrisomatia | https://lpsn.dsmz.de/class/deltaproteobacteria |

| | |
|---|---|
| Dehalococcoidia | https://lpsn.dsmz.de/class/dehalococcoidia |
| Deinococci | https://lpsn.dsmz.de/class/deinococci |
| Deinococcia | https://lpsn.dsmz.de/class/deinococci |
| Deltabacteria | https://lpsn.dsmz.de/class/deltaproteobacteria |
| Deltaproteobacteria | https://lpsn.dsmz.de/class/deltaproteobacteria |
| Desulfarculia | https://lpsn.dsmz.de/class/deltaproteobacteria |
| Desulfobaccia | https://lpsn.dsmz.de/class/deltaproteobacteria |
| Desulfobacteria | https://lpsn.dsmz.de/class/deltaproteobacteria |
| Desulfobulbia | https://lpsn.dsmz.de/class/deltaproteobacteria |
| Desulfomonilia | https://lpsn.dsmz.de/class/deltaproteobacteria |
| Desulfotomaculia | https://lpsn.dsmz.de/class/clostridia |
| Desulfovibrionia | https://lpsn.dsmz.de/class/deltaproteobacteria |
| Desulfurellia | https://lpsn.dsmz.de/class/deltaproteobacteria |
| Desulfuribacillia | https://lpsn.dsmz.de/class/bacilli |
| Desulfuromonadia | https://lpsn.dsmz.de/class/deltaproteobacteria |
| Dictyoglomeria | https://lpsn.dsmz.de/class/dictyoglomeria |
| Dictyoglomia | https://lpsn.dsmz.de/class/dictyoglomeria |
| Dissulfuribacteria | https://lpsn.dsmz.de/class/deltaproteobacteria |
| Elusimicrobia | https://lpsn.dsmz.de/class/elusimicrobia |
| Elusimicrobiia | https://lpsn.dsmz.de/class/elusimicrobia |
| Endomicrobia | https://lpsn.dsmz.de/class/endomicrobiia |
| Endomicrobiia | https://lpsn.dsmz.de/class/endomicrobiia |
| Epsilobacteria | https://lpsn.dsmz.de/class/aquificae |
| Epsilonproteobacteria | https://lpsn.dsmz.de/class/epsilonproteobacteria |
| Erysipelotrichia | https://lpsn.dsmz.de/class/erysipelotrichia |
| Ferrobacteria | https://lpsn.dsmz.de/class/deferribacteres |
| Fibrobacteria | https://lpsn.dsmz.de/class/fibrobacteria |
| Fimbriimonadia | https://lpsn.dsmz.de/class/fimbriimonadia |
| Firmibacteria | https://lpsn.dsmz.de/class/bacilli |
| Flavobacteria | https://lpsn.dsmz.de/class/cytophagia |
| Flavobacteriia | https://lpsn.dsmz.de/class/flavobacteriia |
| Fusobacteria | https://lpsn.dsmz.de/class/fusobacteriia |
| Fusobacteriia | https://lpsn.dsmz.de/class/fusobacteriia |
| Gammaproteobacteria | https://lpsn.dsmz.de/class/gammaproteobacteria |
| Gemmatimonadetes | https://lpsn.dsmz.de/class/gemmatimonadetes |
| Gemmatimonadia | https://lpsn.dsmz.de/class/gemmatimonadetes |
| Gloeobacteria | https://lpsn.dsmz.de/class/cyanophyceae |
| Hadobacteria | https://lpsn.dsmz.de/class/deinococci |
| Halanaerobiia | https://lpsn.dsmz.de/class/clostridia |
| Halobacteria | https://lpsn.dsmz.de/class/halobacteria |
| Halomebacteria | https://lpsn.dsmz.de/class/halobacteria |
| Holophagae | https://lpsn.dsmz.de/class/holophagae |
| Hormogoneae | https://lpsn.dsmz.de/class/cyanophyceae |
| Hydrogenophilalia | https://lpsn.dsmz.de/class/hydrogenophilia |

| | |
|---|---|
| Hydrogenophilia | https://lpsn.dsmz.de/class/hydrogenophilia |
| Ignavibacteria | https://lpsn.dsmz.de/class/ignavibacteria |
| Ignavibacteriia | https://lpsn.dsmz.de/class/ignavibacteria |
| Kiritimatiellae | https://lpsn.dsmz.de/class/kiritimatiellia |
| Kiritimatiellia | https://lpsn.dsmz.de/class/kiritimatiellia |
| Ktedobacteria | https://lpsn.dsmz.de/class/ktedonobacteria |
| Ktedonobacteria | https://lpsn.dsmz.de/class/ktedonobacteria |
| Lentisphaeria | https://lpsn.dsmz.de/class/lentisphaeria |
| Leptospiria | https://lpsn.dsmz.de/class/spirochaetia |
| Limnochordia | https://lpsn.dsmz.de/class/limnochordia |
| Longimicrobia | https://lpsn.dsmz.de/class/longimicrobiia |
| Longimicrobiia | https://lpsn.dsmz.de/class/longimicrobiia |
| Mariprofundia | https://lpsn.dsmz.de/class/alphaproteobacteria |
| Methanobacteria | https://lpsn.dsmz.de/class/methanobacteria |
| Methanococci | https://lpsn.dsmz.de/class/methanococci |
| Methanomicrobia | https://lpsn.dsmz.de/class/methanomicrobia |
| Methanonatronarchaeia | https://lpsn.dsmz.de/class/methanonatronarchaeia |
| Methanopyri | https://lpsn.dsmz.de/class/methanopyri |
| Methanothermea | https://lpsn.dsmz.de/class/methanococci |
| Mollicutes | https://lpsn.dsmz.de/class/mollicutes |
| Myxococcia | https://lpsn.dsmz.de/class/deltaproteobacteria |
| Myxococcia | https://lpsn.dsmz.de/class/deltaproteobacteria |
| Nanobdellia | https://lpsn.dsmz.de/class/nanobdellia |
| Natranaerobiia | https://lpsn.dsmz.de/class/clostridia |
| Nautiliia | https://lpsn.dsmz.de/class/epsilonproteobacteria |
| Negativicutes | https://lpsn.dsmz.de/class/negativicutes |
| Nitriliruptoria | https://lpsn.dsmz.de/class/nitriliruptoria |
| Nitrososphaeria | https://lpsn.dsmz.de/class/nitrososphaeria |
| Nitrospinia | https://lpsn.dsmz.de/class/nitrospinia |
| Nitrospira | https://lpsn.dsmz.de/class/nitrospiria |
| Nitrospiria | https://lpsn.dsmz.de/class/nitrospiria |
| Oligoflexia | https://lpsn.dsmz.de/class/oligoflexia |
| Oligosphaeria | https://lpsn.dsmz.de/class/oligosphaeria |
| Opitutae | https://lpsn.dsmz.de/class/opitutae |
| Opitutia | https://lpsn.dsmz.de/class/opitutae |
| Oxyphotobacteria | https://lpsn.dsmz.de/class/cyanophyceae |
| Phycisphaerae | https://lpsn.dsmz.de/class/phycisphaerae |
| Picrophilea | https://lpsn.dsmz.de/class/thermoplasmata |
| Planctomycea | https://lpsn.dsmz.de/class/planctomycetia |
| Planctomycetacia | https://lpsn.dsmz.de/class/planctomycetia |
| Planctomycetes | https://lpsn.dsmz.de/class/planctomycetia |
| Planctomycetia | https://lpsn.dsmz.de/class/planctomycetia |
| Polyangia | https://lpsn.dsmz.de/class/deltaproteobacteria |
| Polyangiia | https://lpsn.dsmz.de/class/deltaproteobacteria |

| | |
|---|---|
| Protoarchaea | https://lpsn.dsmz.de/class/thermococci |
| Rhodothermia | https://lpsn.dsmz.de/class/rhodothermia |
| Rubrobacteria | https://lpsn.dsmz.de/class/rubrobacteria |
| Saprospiria | https://lpsn.dsmz.de/class/saprospiria |
| Selenobacteria | https://lpsn.dsmz.de/class/negativicutes |
| Selenomonadia | https://lpsn.dsmz.de/class/negativicutes |
| Spartobacteria | https://lpsn.dsmz.de/class/terrimicrobiia |
| Sphingobacteriia | https://lpsn.dsmz.de/class/sphingobacteriia |
| Spirochaetes | https://lpsn.dsmz.de/class/spirochaetia |
| Spirochaetia | https://lpsn.dsmz.de/class/spirochaetia |
| Streptomycetes | https://lpsn.dsmz.de/class/actinomycetes |
| Synergistia | https://lpsn.dsmz.de/class/synergistia |
| Syntrophia | https://lpsn.dsmz.de/class/deltaproteobacteria |
| Syntrophobacteria | https://lpsn.dsmz.de/class/deltaproteobacteria |
| Syntrophorhabdia | https://lpsn.dsmz.de/class/deltaproteobacteria |
| Teichobacteria | https://lpsn.dsmz.de/class/bacilli |
| Tepidiformia | https://lpsn.dsmz.de/class/tepidiformia |
| Terriglobia | https://lpsn.dsmz.de/class/terriglobia |
| Terrimicrobia | https://lpsn.dsmz.de/class/terrimicrobiia |
| Terrimicrobiia | https://lpsn.dsmz.de/class/terrimicrobiia |
| Thermia | https://lpsn.dsmz.de/class/deinococci |
| Thermoanaerobaculia | https://lpsn.dsmz.de/class/thermoanaerobaculia |
| Thermococci | https://lpsn.dsmz.de/class/thermococci |
| Thermodesulfobacteria | https://lpsn.dsmz.de/class/thermodesulfobacteria |
| Thermodesulfobacteriia | https://lpsn.dsmz.de/class/thermodesulfobacteria |
| Thermodesulfobiia | https://lpsn.dsmz.de/class/clostridia |
| Thermodesulfovibrionia | https://lpsn.dsmz.de/class/thermodesulfovibrionia |
| Thermoflexia | https://lpsn.dsmz.de/class/thermoflexia |
| Thermoleophilia | https://lpsn.dsmz.de/class/thermoleophilia |
| Thermolithobacteria | https://lpsn.dsmz.de/class/thermolithobacteria |
| Thermomicrobia | https://lpsn.dsmz.de/class/thermomicrobia |
| Thermoplasmata | https://lpsn.dsmz.de/class/thermoplasmata |
| Thermoprotea | https://lpsn.dsmz.de/class/thermoprotei |
| Thermoprotei | https://lpsn.dsmz.de/class/thermoprotei |
| Thermotogae | https://lpsn.dsmz.de/class/thermotogae |
| Thermotogia | https://lpsn.dsmz.de/class/thermotogae |
| Tichowtungiia | https://lpsn.dsmz.de/class/tichowtungiia |
| Tissierellia | https://lpsn.dsmz.de/class/tissierellia |
| Togobacteria | https://lpsn.dsmz.de/class/thermotogae |
| Verrucomicrobiae | https://lpsn.dsmz.de/class/verrucomicrobiae |
| Verrucomicrobiia | https://lpsn.dsmz.de/class/verrucomicrobiae |
| Vicinamibacteria | https://lpsn.dsmz.de/class/vicinamibacteria |
| Zetaproteobacteria | https://lpsn.dsmz.de/class/alphaproteobacteria |
| Abditibacteriota | https://lpsn.dsmz.de/phylum/abditibacteriota |

| | |
|---|---|
| Acidobacteraeota | https://lpsn.dsmz.de/phylum/acidobacteriota |
| Acidobacteria | https://lpsn.dsmz.de/phylum/acidobacteriota |
| Acidobacteriota | https://lpsn.dsmz.de/phylum/acidobacteriota |
| Actinobacteraeota | https://lpsn.dsmz.de/phylum/actinomycetota |
| Actinobacteria | https://lpsn.dsmz.de/phylum/actinomycetota |
| Actinobacteria | https://lpsn.dsmz.de/phylum/actinomycetota |
| Actinobacteriota | https://lpsn.dsmz.de/phylum/actinomycetota |
| Actinomycetota | https://lpsn.dsmz.de/phylum/actinomycetota |
| Alphaproteobacteraeota | https://lpsn.dsmz.de/phylum/pseudomonadota |
| Alphaproteobacteriota | https://lpsn.dsmz.de/phylum/pseudomonadota |
| Aquificae | https://lpsn.dsmz.de/phylum/aquificota |
| Aquificaeota | https://lpsn.dsmz.de/phylum/aquificota |
| Aquificota | https://lpsn.dsmz.de/phylum/aquificota |
| Aquithermota | https://lpsn.dsmz.de/phylum/aquificota |
| Archaebacteria | https://lpsn.dsmz.de/phylum/euryarchaeota |
| Armatimonadaeota | https://lpsn.dsmz.de/phylum/armatimonadota |
| Armatimonadetes | https://lpsn.dsmz.de/phylum/armatimonadota |
| Armatimonadota | https://lpsn.dsmz.de/phylum/armatimonadota |
| Atribacterota | https://lpsn.dsmz.de/phylum/atribacterota |
| Bacillaeota | https://lpsn.dsmz.de/phylum/bacillota |
| Bacillota | https://lpsn.dsmz.de/phylum/bacillota |
| Bacteroidaeota | https://lpsn.dsmz.de/phylum/bacteroidota |
| Bacteroidetes | https://lpsn.dsmz.de/phylum/bacteroidota |
| Bacteroidota | https://lpsn.dsmz.de/phylum/bacteroidota |
| Balneolaeota | https://lpsn.dsmz.de/phylum/balneolota |
| Balneolota | https://lpsn.dsmz.de/phylum/balneolota |
| Bdellovibrionota | https://lpsn.dsmz.de/phylum/pseudomonadota |
| Bdellovibrionota | https://lpsn.dsmz.de/phylum/pseudomonadota |
| Caldiserica | https://lpsn.dsmz.de/phylum/caldisericota |
| Caldisericaeota | https://lpsn.dsmz.de/phylum/caldisericota |
| Caldisericota | https://lpsn.dsmz.de/phylum/caldisericota |
| Calditrichaeota | https://lpsn.dsmz.de/phylum/calditrichota |
| Calditrichota | https://lpsn.dsmz.de/phylum/calditrichota |
| Campylobacterota | https://lpsn.dsmz.de/phylum/pseudomonadota |
| Campylobacterota | https://lpsn.dsmz.de/phylum/pseudomonadota |
| Chlamydaeota | https://lpsn.dsmz.de/phylum/chlamydiota |
| Chlamydiae | https://lpsn.dsmz.de/phylum/chlamydiota |
| Chlamydiota | https://lpsn.dsmz.de/phylum/chlamydiota |
| Chlorobacteria | https://lpsn.dsmz.de/phylum/chloroflexota |
| Chlorobiota | https://lpsn.dsmz.de/phylum/chlorobiota |
| Chloroflexaeota | https://lpsn.dsmz.de/phylum/chloroflexota |
| Chloroflexi | https://lpsn.dsmz.de/phylum/chloroflexota |
| Chloroflexota | https://lpsn.dsmz.de/phylum/chloroflexota |
| Chrysiogenaeota | https://lpsn.dsmz.de/phylum/chrysiogenota |

| | |
|---|---|
| Chrysiogenetes | https://lpsn.dsmz.de/phylum/chrysiogenota |
| Chrysiogenetota | https://lpsn.dsmz.de/phylum/chrysiogenota |
| Chrysiogenota | https://lpsn.dsmz.de/phylum/chrysiogenota |
| Coprothermobacterota | https://lpsn.dsmz.de/phylum/coprothermobacterota |
| Crenarchaeota | https://lpsn.dsmz.de/phylum/thermoproteota |
| Cyanobacteria | https://lpsn.dsmz.de/phylum/cyanobacteriota |
| Cyanobacteriota | https://lpsn.dsmz.de/phylum/cyanobacteriota |
| Deferribacteraeota | https://lpsn.dsmz.de/phylum/deferribacterota |
| Deferribacteres | https://lpsn.dsmz.de/phylum/deferribacterota |
| Deferribacterota | https://lpsn.dsmz.de/phylum/deferribacterota |
| Deinococcaeota | https://lpsn.dsmz.de/phylum/deinococcota |
| Deinococcota | https://lpsn.dsmz.de/phylum/deinococcota |
| Deinococcus-Thermus | https://lpsn.dsmz.de/phylum/deinococcota |
| Desulfobacterota | https://lpsn.dsmz.de/phylum/pseudomonadota |
| Desulfotomaculota | https://lpsn.dsmz.de/phylum/bacillota |
| Dictyoglomaeota | https://lpsn.dsmz.de/phylum/dictyoglomerota |
| Dictyoglomera | https://lpsn.dsmz.de/phylum/dictyoglomerota |
| Dictyoglomerota | https://lpsn.dsmz.de/phylum/dictyoglomerota |
| Dictyoglomi | https://lpsn.dsmz.de/phylum/dictyoglomerota |
| Dictyoglomota | https://lpsn.dsmz.de/phylum/dictyoglomerota |
| Dictyoglomota | https://lpsn.dsmz.de/phylum/dictyoglomerota |
| Elusimicrobaeota | https://lpsn.dsmz.de/phylum/elusimicrobiota |
| Elusimicrobia | https://lpsn.dsmz.de/phylum/elusimicrobiota |
| Elusimicrobiota | https://lpsn.dsmz.de/phylum/elusimicrobiota |
| Endobacteria | https://lpsn.dsmz.de/phylum/bacillota |
| Eobacteria | https://lpsn.dsmz.de/phylum/chloroflexota |
| Epsilonbacteraeota | https://lpsn.dsmz.de/phylum/pseudomonadota |
| Euryarchaeota | https://lpsn.dsmz.de/phylum/euryarchaeota |
| Eurybacteria | https://lpsn.dsmz.de/phylum/bacillota |
| Eurybacteria | https://lpsn.dsmz.de/phylum/bacillota |
| Fibrobacteraeota | https://lpsn.dsmz.de/phylum/fibrobacterota |
| Fibrobacteres | https://lpsn.dsmz.de/phylum/fibrobacterota |
| Fibrobacterota | https://lpsn.dsmz.de/phylum/fibrobacterota |
| Firmacutes | https://lpsn.dsmz.de/phylum/bacillota |
| Firmicutes | https://lpsn.dsmz.de/phylum/bacillota |
| Fusobacteraeota | https://lpsn.dsmz.de/phylum/fusobacteriota |
| Fusobacteria | https://lpsn.dsmz.de/phylum/fusobacteriota |
| Fusobacteriota | https://lpsn.dsmz.de/phylum/fusobacteriota |
| Gemmatimonadaeota | https://lpsn.dsmz.de/phylum/gemmatimonadota |
| Gemmatimonadetes | https://lpsn.dsmz.de/phylum/gemmatimonadota |
| Gemmatimonadota | https://lpsn.dsmz.de/phylum/gemmatimonadota |
| Hadobacteria | https://lpsn.dsmz.de/phylum/deinococcota |
| Ignavibacteriota | https://lpsn.dsmz.de/phylum/chlorobiota |
| Kiritimatiellaeota | https://lpsn.dsmz.de/phylum/kiritimatiellota |

| | |
|---|---|
| Kiritimatiellota | https://lpsn.dsmz.de/phylum/kiritimatiellota |
| Lentisphaerae | https://lpsn.dsmz.de/phylum/lentisphaerota |
| Lentisphaeraeota | https://lpsn.dsmz.de/phylum/lentisphaerota |
| Lentisphaerota | https://lpsn.dsmz.de/phylum/lentisphaerota |
| Mendosicutes | https://lpsn.dsmz.de/phylum/euryarchaeota |
| Methanobacteraeota | https://lpsn.dsmz.de/phylum/euryarchaeota |
| Methanobacteriota | https://lpsn.dsmz.de/phylum/euryarchaeota |
| Mollicutaeota | https://lpsn.dsmz.de/phylum/mycoplasmatota |
| Mollicutota | https://lpsn.dsmz.de/phylum/mycoplasmatota |
| Mycoplasmatota | https://lpsn.dsmz.de/phylum/mycoplasmatota |
| Myxococcota | https://lpsn.dsmz.de/phylum/pseudomonadota |
| Myxococcota | https://lpsn.dsmz.de/phylum/pseudomonadota |
| Nanoarchaeota | https://lpsn.dsmz.de/phylum/nanoarchaeota |
| Nitrososphaeraeota | https://lpsn.dsmz.de/phylum/nitrososphaerota |
| Nitrososphaerota | https://lpsn.dsmz.de/phylum/nitrososphaerota |
| Nitrospinae | https://lpsn.dsmz.de/phylum/nitrospinota |
| Nitrospinota | https://lpsn.dsmz.de/phylum/nitrospinota |
| Nitrospirae | https://lpsn.dsmz.de/phylum/nitrospirota |
| Nitrospiraeota | https://lpsn.dsmz.de/phylum/nitrospirota |
| Nitrospirota | https://lpsn.dsmz.de/phylum/nitrospirota |
| Planctobacteria | https://lpsn.dsmz.de/phylum/planctomycetota |
| Planctomycetaeota | https://lpsn.dsmz.de/phylum/planctomycetota |
| Planctomycetes | https://lpsn.dsmz.de/phylum/planctomycetota |
| Planctomycetota | https://lpsn.dsmz.de/phylum/planctomycetota |
| Posibacteria | https://lpsn.dsmz.de/phylum/bacillota |
| Proteobacteria | https://lpsn.dsmz.de/phylum/pseudomonadota |
| Proteobacteria | https://lpsn.dsmz.de/phylum/pseudomonadota |
| Proteobacteria | https://lpsn.dsmz.de/phylum/pseudomonadota |
| Pseudomonadota | https://lpsn.dsmz.de/phylum/pseudomonadota |
| Rhodothermaeota | https://lpsn.dsmz.de/phylum/rhodothermota |
| Rhodothermota | https://lpsn.dsmz.de/phylum/rhodothermota |
| Sphingobacteria | https://lpsn.dsmz.de/phylum/bacteroidota |
| Spirochaetae | https://lpsn.dsmz.de/phylum/spirochaetota |
| Spirochaetae | https://lpsn.dsmz.de/phylum/spirochaetota |
| Spirochaetaeota | https://lpsn.dsmz.de/phylum/spirochaetota |
| Spirochaetes | https://lpsn.dsmz.de/phylum/spirochaetota |
| Spirochaetota | https://lpsn.dsmz.de/phylum/spirochaetota |
| Synergistaeota | https://lpsn.dsmz.de/phylum/synergistota |
| Synergistetes | https://lpsn.dsmz.de/phylum/synergistota |
| Synergistota | https://lpsn.dsmz.de/phylum/synergistota |
| Synthermota | https://lpsn.dsmz.de/phylum/thermotogota |
| Tenericutes | https://lpsn.dsmz.de/phylum/mycoplasmatota |
| Thaumarchaeota | https://lpsn.dsmz.de/phylum/nitrososphaerota |
| Thermodesulfobacteraeota | https://lpsn.dsmz.de/phylum/thermodesulfobacteriota |

| | |
|---|---|
| Thermodesulfobacteria | https://lpsn.dsmz.de/phylum/thermodesulfobacteriota |
| Thermodesulfobacteriota | https://lpsn.dsmz.de/phylum/thermodesulfobacteriota |
| Thermomicrobaeota | https://lpsn.dsmz.de/phylum/thermomicrobiota |
| Thermomicrobia | https://lpsn.dsmz.de/phylum/thermomicrobiota |
| Thermomicrobiota | https://lpsn.dsmz.de/phylum/thermomicrobiota |
| Thermoproteaeota | https://lpsn.dsmz.de/phylum/thermoproteota |
| Thermoproteota | https://lpsn.dsmz.de/phylum/thermoproteota |
| Thermotogae | https://lpsn.dsmz.de/phylum/thermotogota |
| Thermotogaeota | https://lpsn.dsmz.de/phylum/thermotogota |
| Thermotogota | https://lpsn.dsmz.de/phylum/thermotogota |
| Verrucomicrobaeota | https://lpsn.dsmz.de/phylum/verrucomicrobiota |
| Verrucomicrobia | https://lpsn.dsmz.de/phylum/verrucomicrobiota |
| Verrucomicrobia | https://lpsn.dsmz.de/phylum/verrucomicrobiota |
| Verrucomicrobia | https://lpsn.dsmz.de/phylum/verrucomicrobiota |
| Verrucomicrobiota | https://lpsn.dsmz.de/phylum/verrucomicrobiota |
